# Genome-wide mapping of autonomously replicating sequences in the marine diatom *Phaeodactylum tricornutum*

**DOI:** 10.1101/2024.07.07.602421

**Authors:** Hyun-Sik Yun, Kohei Yoneda, Takehito Sugasawa, Iwane Suzuki, Yoshiaki Maeda

## Abstract

Autonomously replicating sequences (ARSs) are important accessories in episomal vectors that allow them to be replicated and stably maintained within transformants. Despite their importance, no information on ARSs in diatoms have been reported. Therefore, we attempted to identify ARS candidates in the model diatom, *Phaeodactylum tricornutum*, via chromatin immunoprecipitation sequencing. In this study, subunits of the origin recognition complex (ORC), ORC2 and ORC4, were used to screen for ARS candidates. ORC2 and ORC4 bound to 355 sites on the *P. tricornutum* genome, of which 69 were constantly screened after multiple attempts. The screened ARS candidates had an AT-richness of approximately 50% (44.39–52.92%) and did not have conserved sequences. In addition, ARS candidates were distributed randomly but had a dense distribution pattern at several sites. Their positions tended to overlap with those of the genetic region (73.91%). Compared to the ARSs of several other eukaryotic organisms, the characteristics of the screened ARS candidates are complex. Thus, our findings suggest that the diatom has a distinct and unique native ARSs.

## Introduction

Diatoms, a microalgal taxon, are photosynthetic organisms that play an important role as producers in aquatic ecosystems (B-Béres *et al.,* 2023). Their ability to produce various types of bioproducts using light energy and inorganic nutrients showcases their potential as a bioresource (Saxena *et al*., 2021). Moreover, the superior biomass productivity of diatoms compared to that of terrestrial plants further highlights their value as a bioresource (Marella *et al*., 2020). To effectively utilise the value of diatoms, efforts have been made to apply genetic engineering to algal biotechnology (Mosey *et al*., 2021; Sproles *et al*., 2021). The development of molecular tools aids in these efforts to enhance the value of diatoms as a resource and provide access to deeper knowledge regarding them (Butler *et al*., 2020; Mosey *et al*., 2021). As a vector for introducing genes of external origin, plasmids randomly integrated into the chromosomes (Chr) in diatoms are one such useful molecular tool that has been applied to various organisms (D’Adamo *et al*., 2019). Attempts have been made to apply this useful tool to diatoms as well, resulting in the successful development of a method for introducing plasmids into them (Dunahay *et al*., 1995; D’Adamo *et al*., 2019; Butler *et al*., 2020). Therefore, it has become possible to introduce a plasmid containing a target gene using various methods, such as biolistics (particle bombardment) and electroporation, to create a diatom transformant (Miyahara *et al*., 2013; Karas *et al*., 2015; Velmurugan & Deka, 2018).

Although methods for producing diatom transformants have been established, limitations remain. In the biolistics and electroporation of a diatom, the target gene is introduced by inserting a plasmid sequences into the genomic DNA of the diatom (Miyahara *et al*., 2013; Velmurugan & Deka, 2018; George *et al*., 2020). In this case, the location wherein the plasmid is inserted is random, and due to random integration, each transformant clone exhibits a different expression pattern for the introduced gene (George *et al*., 2020; Kassaw *et al*., 2022). In addition, there is a possibility that expression of the introduced gene may be silenced as the transformant is sub-cultured (Leon & Fernandez, 2007; Lee *et al*., 2020). Unstable expression of the introduced gene can be a critical handicap when conducting research using transformants (Leon & Fernandez, 2007). In addition, there are no reports of the introduction of large-sized plasmids into diatoms via electroporation (Karas *et al*., 2015; Muñoz *et al*., 2018). Restrictions on the size of plasmids that can be introduced affect their design, and it is difficult to effectively introduce a large number and large size of genes to take advantage of mega-plasmids when using electroporation (Karas *et al*., 2015; Neerathilingam *et al*., 2019; George *et al*., 2020). Therefore, although it is possible to create transformants via electroporation, there are clear limitations; thus, an alternative approach is necessary.

Bacterial conjugation has emerged as a novel method to generate diatom transformants (Karas *et al*., 2015). By adding an origin of transfer (oriT) to the plasmid design, plasmids of a relatively larger size (e.g., 24–94 kbp) than those that could be introduced by using electroporation can then be introduced via conjugation (Karas *et al*., 2015). One of the major advantages is that as the size of the plasmid can be increased, restrictions on plasmid design can be reduced (Garza *et al*., 2023). Furthermore, the fact that plasmids are transferred to diatoms in the form of episomal DNA via conjugation distinguishes it from electroporation (Kassaw *et al*., 2022; Garza *et al*., 2023). The phenomenon where a transgene inserted into genomic DNA is silenced due to various mechanisms, such as chromatin modification or transcriptional regulation, is a clear limitation of electroporation (Neupert *et al*., 2020). In contrast, the ability to introduce plasmids in the form of episomal DNA is a great advantage of conjugation when used to create diatom transformants (Garza *et al*., 2023).

However, diatom transformation via conjugation also has limitations that must be resolved. In order for the episomal DNA of a eukaryotic organism to be maintained stably, it must contain sequences that can function as a centromere (CEN) and autonomously replicating sequence (ARS) (Cao *et al*., 2017). During the initial studies when conjugation was attempted for diatom transformation, it was hoped that the plasmid could be stably maintained in the transformant as episomal DNA by including yeast CEN/ARS in the plasmid design (Karas *et al*., 2015; Diner *et al*., 2016). Previous studies have reported that yeast CEN/ARS provides stability to the episomal DNA introduced in diatom transformants (Garza *et al*., 2023), but it was revealed that the functionality provided by it had limitations in maintaining episomal DNA (Diner *et al*., 2017; Kumar *et al*., 2020). Therefore, for improved stability of episomal DNA, sequence information of the CEN and ARSs of diatoms is essential (Diner *et al*., 2017; Kumar *et al*., 2020). In previous research, information about the sequence of CEN in the marine diatom, *Phaeodactylum tricornutum*, was revealed, and it was also reported that the stability of episomal DNA can be strengthened through the identified sequences (Diner *et al*., 2017). However, there is currently no information on ARSs in diatoms.

In this study, we attempted to elucidate information on the ARSs of *P. tricornutum*. Using ORC2 and ORC4, subunits of the origin recognition complex (ORC; Figs. 1, S1, S2), a protein complex that recognises ARSs (Cheng *et al*., 2020), nucleotide sequences of the regions where ORC interacts in the genomic DNA of the diatom were analysed via chromatin immunoprecipitation sequencing (ChIP-seq) (Schepers & Papior, 2010). Furthermore, ARS candidates were predicted from sequence information obtained from the ChIP-seq analysis of ORC2 and ORC4. Based on these results, this study provides unique information and insight into the ARSs in diatoms.

**Fig. 1.**
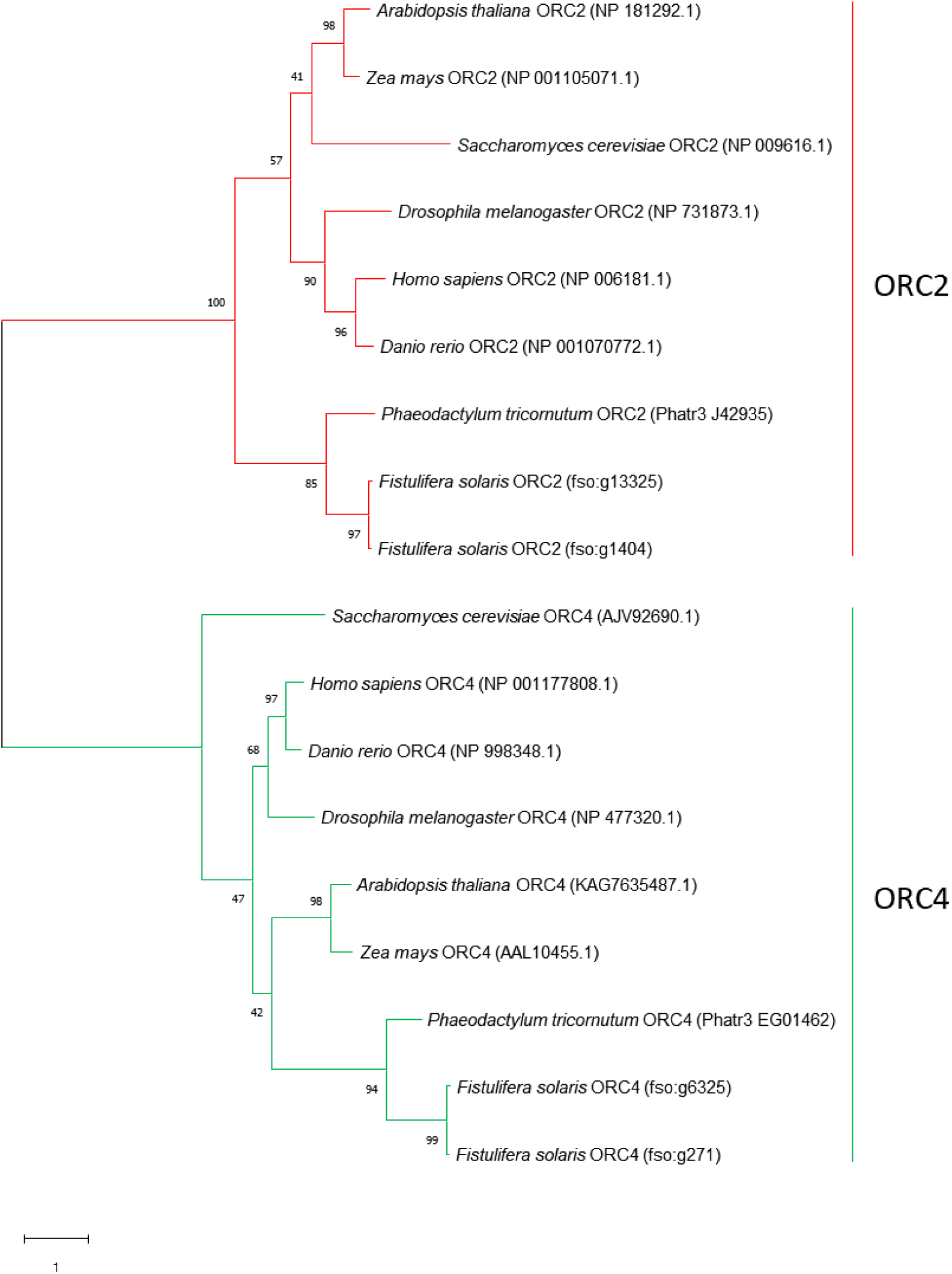
Phylogenetic tree visualising the relationship between the ORC2 and OCR4 of eight eukaryotic organisms. A phylogenetic tree was created based on amino acid sequences of the ORC2 and ORC4 from eight eukaryotic organisms (*Phaeodactylum tricornutum, Fistulifera solaris, Homo sapiens, Danio rerio, Drosophila melanogaster, Arabidopsis thaliana, Zea mays,* and *Saccharomyces cerevisiae*). Branches related to ORC2 are indicated by red lines, and those related to ORC4 indicated by green lines.

**Fig. 2.**
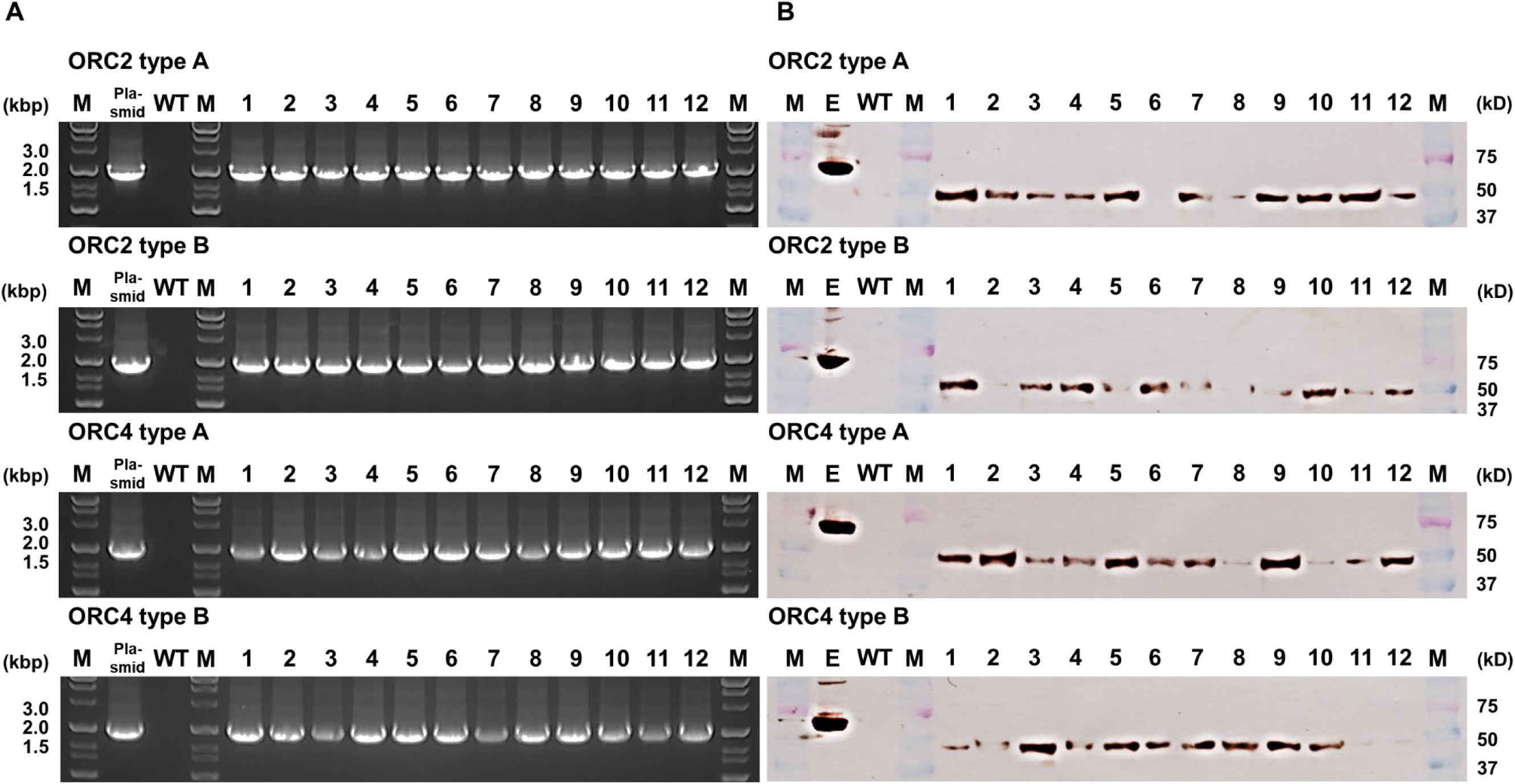
Existence and expression of the FLAG-tag-fused ORC subunit cassette. (A) To confirm the presence of the gene cassette introduced in the transformant, the region from the nitrate reductase promoter to the end of the ORC subunit was amplified via PCR. (B) Western blotting confirmed that the FLAG-tag-fused ORC subunit was expressed from the introduced cassette in the transformant. Lanes 1∼2: transformant clones, M: ladder, WT: wild type, E: Epitope Tag Protein Marker Lysate.

## Materials and Methods

### Algal strain and culturing conditions

The *P. tricornutum* CCAP 1055/1 strain was obtained from the Culture Collection of Algae and Protozoa (CCAP, Oban, UK). The *P. tricornutum* was cultivated and preserved on f/2 medium agar plates (1.5%, Bacto agar) (Guillard, 1975). The f/2 medium was made with artificial seawater (Marine Art SF-1) as a base. For the experiment, colonies on agar plates were picked, inoculated into the liquid medium (150 ml f/2 medium in a 300 ml flask), and cultured on a shaker (130 rpm). The agar plates and flasks were placed in an illuminated (with cool, white fluorescent lights) growth chamber, with the temperature and light set at 20°C and 40 µmol photons m^−2^ s^−1^, respectively.

### Plasmid construction and transformant creation for FLAG-tag-fused ORC2 and ORC4

For *P. tricornutum* transformants expressing FLAG-tag (DYKDDDDK)-fused ORC2 and ORC4, plasmids were constructed based on the pPha-T1 vector (Zaslavskaia *et al*., 2000). The promoter for expression of the insert (FLAG-tag-fused ORC2 and ORC4) was replaced with the nitrate reductase (NR) promoter from the fcpA promoter (Fig. S3). The ORC2 and ORC4 genes inserted downstream of the NR promoter were amplified using primer sets designed to fuse the FLAG-tag at the N-terminus; ORC2: ORC2_Fw and ORC2_Rev; ORC4: ORC4_Fw and ORC4_Rev (Table S1). The linearized plasmids were prepared via PCR with the primer set, pPha_Fw and pPha_Rev (Table S1), and the insert fragments fused to the linearized plasmids using the In-Fusion Cloning Kit (Takara Bio Inc., Shiga, Japan), according to the protocol provided by the manufacturer with the *Escherichia coli* JM109 strain. Entire sequences of the constructed plasmids were confirmed with nanopore sequencing (Eurofins Genomics K.K., Tokyo, Japan).

The plasmid containing the correct sequence of inserts was introduced into *P. tricornutum* with the electroporation method, as previously reported (Miyahara *et al*., 2013). For electroporation, vegetative cells in the growth phase (optical density measured at 750 nm [OD_750_]: 0.3–0.4, 3.24 million cells/ml) were collected via centrifugation (2000 × *g*, 2 min). The collected cell pellets were washed with 0.77 M mannitol solution. Electroporation was performed using NEPA21 (Nepa Gene, Chiba, Japan) after 10 μg plasmid DNA was added to the mixture for each attempt. Subsequently, the cells were suspended in fresh f/2 medium and cultured for recovery in a growth chamber at 20°C and the light set at 40 µmol photons m^−2^ s^−1^ for 1–2 days. Cells that completed recovery culture were collected via centrifugation and spread onto plates for selection (f/2 medium agar plates with zeocin: 1.5% [w/v] agar and 50 μg/ml zeocin). Colonies of the transformant were observed on the plates 1–2 months after spreading, and the observed colonies then picked and transferred to a well plate containing f/2 medium with 50 μg/ml zeocin.

### Chromatin immunoprecipitation sequencing

For ChIP, approximately 100 mg (wet weight) frozen samples (OD_750_: 0.3–0.4, 3.24 × 10^6^ cells/ml) were homogenized (2500 rpm, 60 s) with a multi-beads shocker (Yasui-Kikai, Osaka, Japan). The homogenized samples were fixed with 1% paraformaldehyde (PFA) solution (1 ml) for 10 min (cross-linking). To neutralize PFA, 2 M glycine solution (ChIP Reagents #318–07131; Nippon Gene, Tokyo, Japan) was added (212 µl per 1 ml sample, final concentration: 350 mM). The pellets of cross-linked samples were collected through centrifugation (2000 × *g*, 4°C, 10 min) and washed twice with NP40 buffer (1 ml) (ChIP Reagents #318–07131; Nippon Gene). The washed pellets were suspended by adding 60 µl SDS lysis buffer (ChIP Reagent #318–07131; Nippon Gene), and then 540 µl ChIP dilution buffer (ChIP Reagent #318–07131; Nippon Gene) added. The suspended samples were sheared by using a Bioruptor Plus sonicator (Diagenode, Denville, NJ, USA). The sonication conditions were set to a 1-min cycle (30 s active, 30 s pause) and conducted for 2 h (target length: 200–1000 bp). The sheared samples were centrifuged (13000 × *g*, 4°C, 10 min), and the supernatants separated. A portion of the supernatant (100 µl) was stored separately as an input (Inp) sample (negative control sample), and the remaining supernatant used as an immunoprecipitation (IP) sample. The protein concentration of the supernatant for immunoprecipitation was measured via the bicinchoninic acid assay (Thermo Scientific, Rockford, IL, USA) (Walker, 2009), and 1× RIPA buffer-150 mM (ChIP Reagents #318–07131; Nippon Gene) added to produce a final protein concentration of the supernatant of 400–500 µg/ml. The diluted samples (500 µl) were mixed with antibody (anti-DYKDDDDK tag, 66008-4-Ig; Proteintech, Rosemont, IL, USA)-coated beads (Dynabeads protein G; Invitrogen, Waltham, MA, USA) and incubated on a rotator overnight at 4°C. To prepare the antibody-coated beads, beads (30 µl) were washed using PBST (500 µl) and collected using a magnetic stand. The washed beads were incubated with the antibodies (10 µg/300 µl in PBST) at room temperature for 10 min using a rotator. After incubation, the beads were collected using a magnetic stand and washed with PBST (500 µl) twice. From the incubated samples (with antibody-coated beads), the beads were collected using a magnetic stand. The collected beads were washed with 1× RIPA buffer-150 mM (500 µl), 1× RIPA buffer-500 mM (500 µl) (ChIP Reagent #318–07131; Nippon Gene), and TE buffer (500 µl) (pH 8.0). ChIP direct elution buffer (300 µl) (ChIP Reagent #318–07131; Nippon Gene) was added to the washed beads and incubated at 95°C for 15 min (reverse-cross-linking). For the input samples, ChIP direct elution buffer (200 µl) was added, and the same process as that described below performed. The eluted DNA samples (250 µl) were separated from the beads using a magnetic stand, whereafter 0.5 µl RNase A (5 mg/ml) was added and incubated (37°C, 30 min). To the incubated samples, 6 µl proteinase K (20 mg/ml) was added and incubated (56°C, 1 h). The incubated DNA samples were purified with the NucleoSpin Gel and PCR Clean-up kit (#740609.250; Takara Bio Inc.). The purified DNA samples were used to prepare the library. The DNA concentrations of the samples were measured via a Nanodrop (Thermo Scientific), and the samples diluted with Mili-Q to adjust the DNA concentrations to 1 ng/µl. Libraries were created using the SMARTer ThruPLEX DNA-Seq Kit (Takara Bio Inc.). DNA samples were repaired, had adapters added, and then used to synthesize libraries. The synthesized libraries were amplified using primers including an index. The amplified libraries were purified using AMPure XP beads (Beckman Coulter Diagnostics, Brea, CA, USA) and quantified using a DNA 7500 Bioanalyzer System (Agilent, Palo Alto, CA, USA). Subsequently, sequencing data were obtained using the MiSeq platform (Illumina, San Diego, CA, USA).

### Bioinformatic analyses

Bowtie2 (ver. 2.2.5) (Langmead & Salzberg, 2012) was used as a software tool to align FASTQ files to the genome (Bowler *et al.,* 2008). The mapped read files (SAM files) were converted to a BAM file format through the samtools view function. BAM files were sorted using the samtools sort function, and the sorted BAM files indexed using the samtools index function. MACS2 (ver. 2.2.9.1) was used as a software tool for peak-calling, and the BAM files peak-called via MACS2 (Feng *et al*., 2012). Peak results were visualised via Integrative Genomics Viewer (IGV) (Robinson *et al*., 2011). Motif analysis was performed using MEME Suite (Bailey *et al*., 2015).

## Results

### Selection of *Phaeodactylum tricornutum* transformants for ChIP-seq

DNA fragments of ORC subunits (ORC2 and ORC4) were successfully amplified from genomic DNA using primer sets designed to add a FLAG-tag to the N-terminus (Fig. S3; Table S1). DNA fragments of 1824 bp in length (FLAG with methionine: 27 bp, ORC2: 1767 bp, sequence for In-Fusion Cloning: 15 bp + 15 bp) were obtained by targeting the ORC2 gene, and DNA fragments of 1860 bp in length (FLAG with methionine: 27 bp, ORC4: 1803 bp, sequence for In-Fusion Cloning: 15 bp + 15 bp) obtained by targeting the ORC4 gene. The amplified DNA fragments were introduced into the expression plasmids as inserts, which were confirmed via Sanger sequencing. Interestingly, from the sequencing results, it was confirmed that two types of sequences with single nucleotide polymorphisms (SNPs) exist for ORC2 and ORC4 (Fig. S2, S4, S5), and all confirmed SNPs were distributed at the positions of missense and synonymous variants reported in the reference database (Ensembl Genomes; http://www.ensemblgenomes.org). In the present study, the two types of ORC2 and ORC4 genes were tentatively designated as type A and type B. In the case of ORC2, compared to the registered sequence (Phatr3_J42935), the DNA sequence of type A had a 6-bp mismatch, and two amino acids had been replaced in the translated amino acid sequence, whereas the DNA sequence of type B had a 2-bp mismatch with two amino acids that had been replaced in the translated amino acid sequence (Fig. S4). In the case of ORC4, compared to the registered sequence (Phatr3_EG01462), the DNA sequence of type A had a 3-bp mismatch, but no amino acids had been replaced in the translated amino acid sequence, whereas the DNA sequence of type B had a 6-bp mismatch with four amino acids that had been replaced in the translated amino acid sequence (Fig. S5). For transformation of *P. tricornutum*, plasmids were prepared separately for both types A and B of ORC2 and ORC4. We confirmed that the plasmids were constructed as we designed by using nanopore sequencing.

Electroporation was performed to introduce the prepared plasmids into *P. tricornutum*, and the transformant clones were selected on agar plates containing zeocin. To confirm the presence of the introduced cassette in the transformants, the region from the end of the NR promoter to the end of the FLAG-tag-fused ORC subunit was amplified via PCR (primer sets: PtNRpseq_Fw/ORC2_Rev, PtNRpseq_Fw/ORC4_Rev; Table S1). DNA fragments of the expected size (ORC2: 1916 bp, ORC4: 1952 bp) could be amplified from genomic DNA of the transformants (Fig. 2). Among transformants with the correct cassette, western blotting was performed to select transformant lines showing the highest expression of the FLAG-tag-fused ORC subunits for use in ChIP-seq. The selected transformant lines (ORC2 A: 1, 5; ORC2 B: 1, 4; ORC4 A 1, 2; ORC4 B: 3, 5) showed relatively inhibited growth compared to that of the wild type (WT) (Fig. S6). Based on the observed growth pattern, the log phase in which the ORC is expected to actively recognise ARSs was identified (Dillin & Rine, 1998), and each transformant line evaluated once again for stable expression of the FLAG-tag-fused ORC subunit in log phase. According to the western blotting results, all selected transformant lines showed stable expression levels throughout the log phase interval (Fig. S7). Based on these results, we selected the transformant lines ORC2 A 1, ORC2 B 1, ORC4 A 1, ORC4 B 3 for ChIP-seq.

### Distribution of screened ORC2- and ORC4-binding sites

Raw data of ChIP-seq were obtained from quality-checked produced library samples, and the results are shown in figures and tables (Figs. S8, S9; Table 1, S2). Information on peak-called sites for each Chr was summarized in Table S3, and based on this, the sites shared by the results obtained via each type of ORC subunits (namely, ORC2 A, ORC2 B, ORC4 A, and ORC4 B) were visualised using a Venn diagram (Fig. 3). Furthermore, the distribution of each site on each Chr was visualised (Fig. 4; Fig. S10). A total of 355 sites were screened from ORC2 and ORC4 (both A and B types). From each Chr (Chr 1–33), as few as 0 sites (Chr 28 and 32) and as many as 23 sites (Chr 3) were screened. Among the total 355 sites, 69 were peak-called by all four ChIP-seq experiments. These sites were not found on Chr 2, 6, 8, 9, 11, 13, 21, 27, 28, 30, and 32 (0 sites), and the largest number of sites was distributed on Chr 1 (nine sites). The uniquely screened sites from each subunit included 43 in ORC2 A, 86 in ORC2 B, 32 in ORC4 A, and 48 in ORC4 B. Interestingly, the screened sites showed a dense distribution (Fig. S10). Among these, results for as many as 14 sites (Chr 25, 0–100 kbp) that were closely distributed had been included. The densely distributed sites accounted for 40% of all sites screened. The remaining 60% (213 sites) were randomly distributed throughout the genome, excluding Chr 28 and 32.

**Table 1.** Statistical information about ChIP-seq in this study.

|  | ORC2 A |  | ORC2 B |  | ORC4 A |  | ORC4 B |  |
| --- | --- | --- | --- | --- | --- | --- | --- | --- |
|  | IP<sup>a</sup> | Inp<sup>b</sup> | IP<sup>a</sup> | Inp<sup>b</sup> | IP<sup>a</sup> | Inp<sup>b</sup> | IP<sup>a</sup> | Inp<sup>b</sup> |
| Protein concentration (µg/ml) | 440 | 440 | 440 | 440 | 440 | 440 | 440 | 440 |
| Library concentration (nM) | 149.50 | 109.25 | 303.70 | 312.35 | 132.10 | 71.00 | 242.20 | 388.60 |
| Number of reads obtained | 1596109 | 1445983 | 1815371 | 1949374 | 1665248 | 1332361 | 1715721 | 1903497 |
| Number of reads mapped to genome | 1366003 | 1264133 | 1539561 | 1714406 | 1429425 | 1165235 | 1402937 | 1569338 |
| Number of peak called | 163 | 163 | 206 | 206 | 140 | 140 | 162 | 162 |

<sup>a</sup>IP: immunoprecipitation sample. <sup>b</sup>Inp: input sample (negative control)

**Fig. 3.**
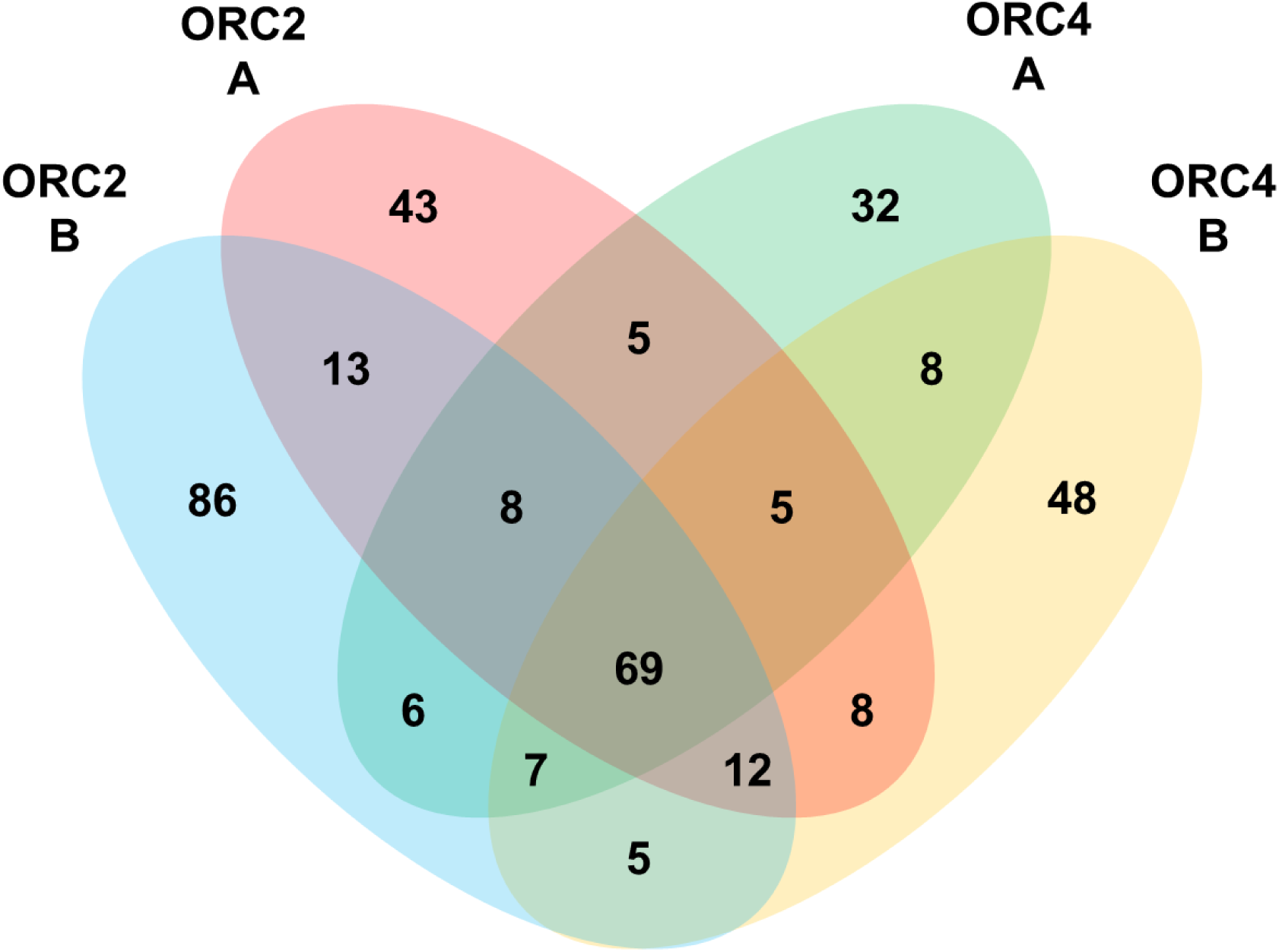
Venn diagram of the number of sites screened from the ORC subunits. The number of sites screened from ORC2 A (red), ORC2 B (blue), ORC4 A (green), and ORC4 B (orange) was visualised through a Venn diagram. The number of sites shared between each result is displayed in the overlapping area.

**Fig. 4.**
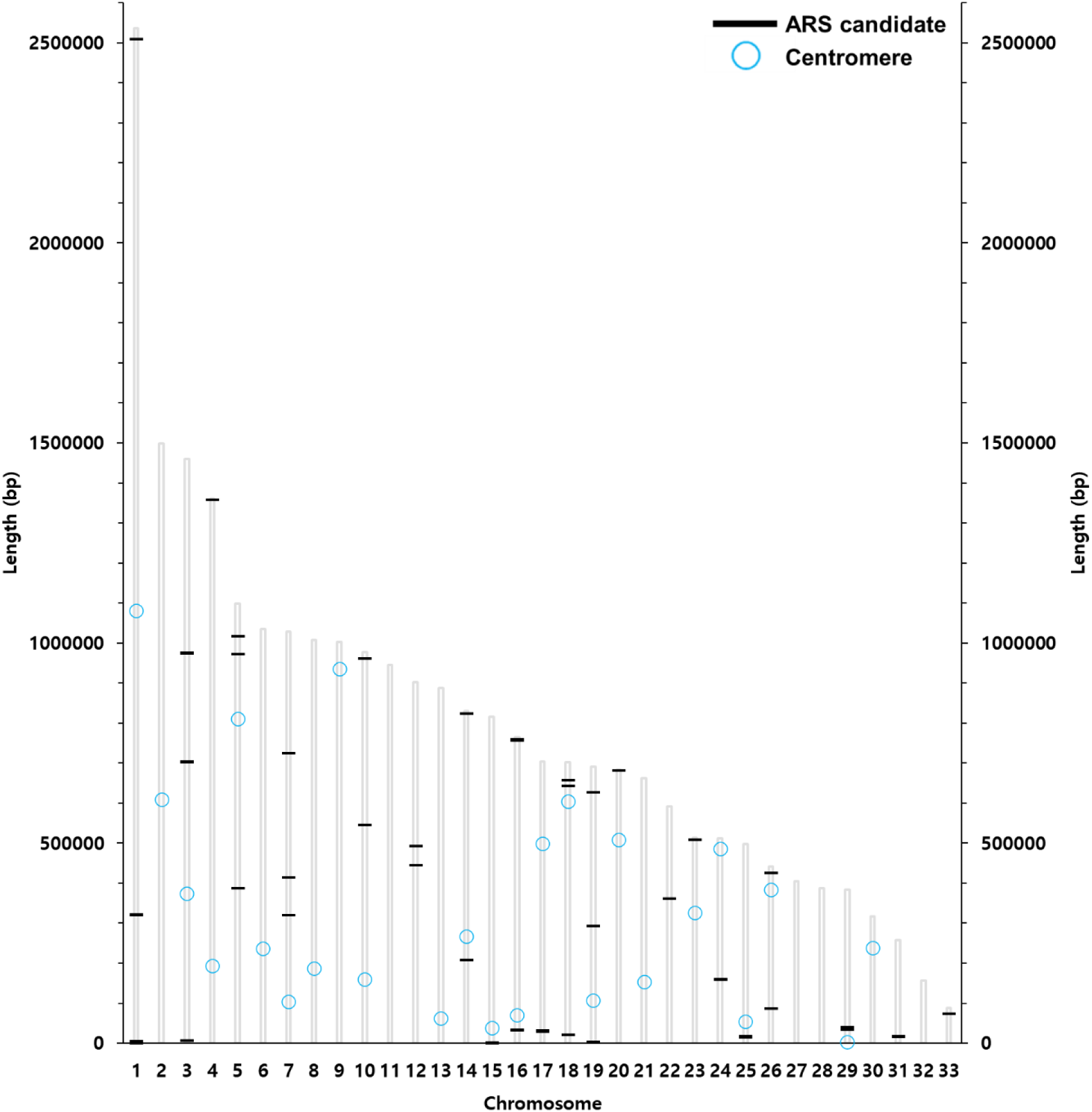
Distribution of shared sites on the genome of *Phaeodactylum tricornutum*. The locations of the 69 sites screened from all ORC subunits (ORC2 A, ORC2 B, ORC4 A, and ORC4 B) are indicated by black lines on the genome of *Phaeodactylum tricornutum*. The locations of centromeres are indicated by light blue circles (Diner *et al.,* 2017). ARS: autonomously replicating sequence.

**Fig. 5.**
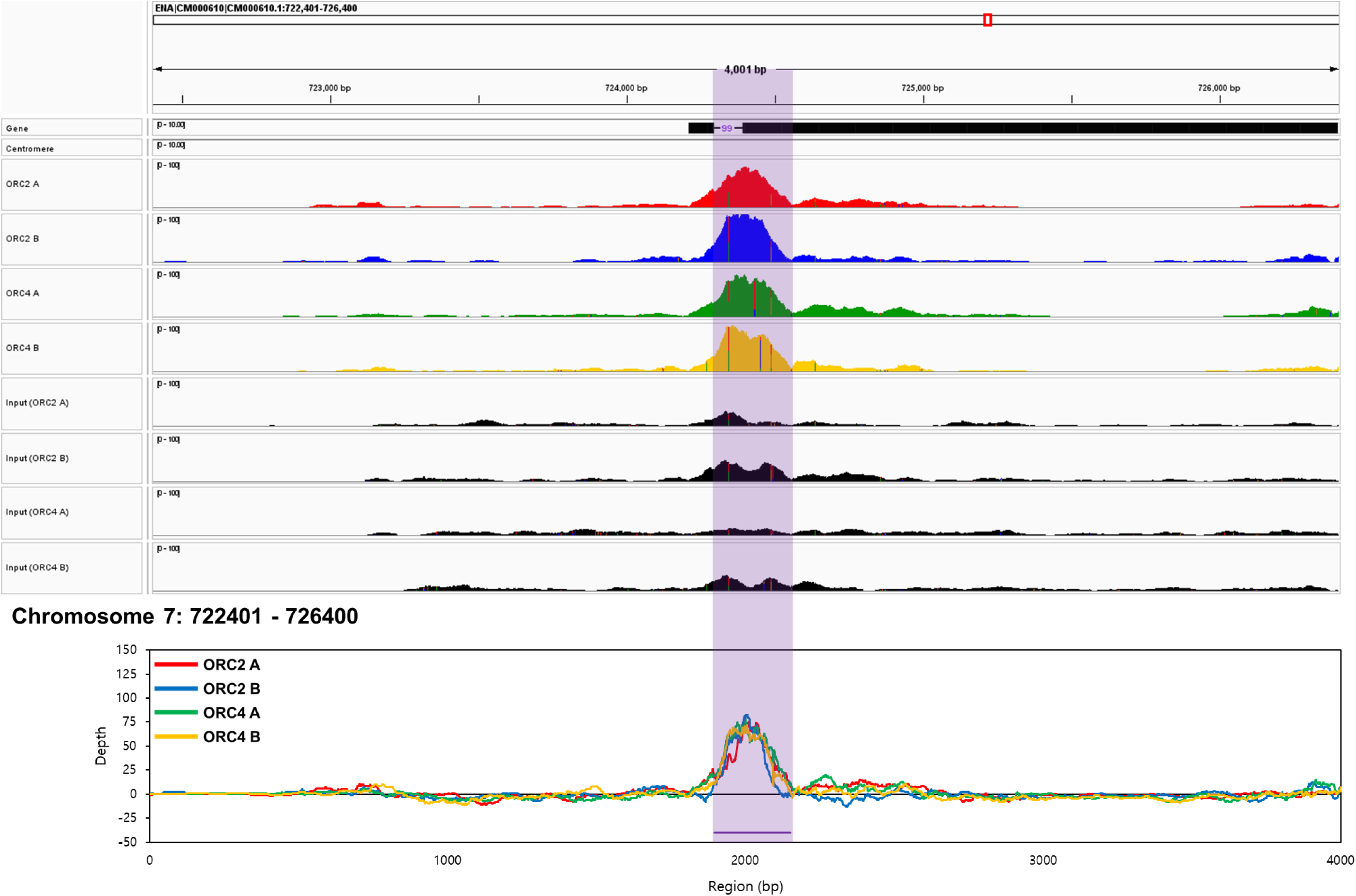
Peak data and depth values at the locations (chr 7: 722401-726400) of shared site number 21 (chr 7: 724295-724550) on the genome of *Phaeodactylum tricornutum*. The results obtained from ORC2 A (red peaks and lines), ORC2 B (blue peaks and lines), ORC4 A (green peaks and lines), and ORC4 B (orange peaks and lines) were visualised through peak (IGV image) and depth images, and shared sites that are autonomously replicating sequence candidates were indicated by a purple line over the peak and depth images.

In the distribution of the 69 shared sites, a tendency for the distributed locations to be concentrated was confirmed (Fig. 4). Overall, except for the concentrated sites, shared sites were randomly distributed throughout the genome. Therefore, our results revealed that 355 sites were screened, of which 58.87% were unshared and 19.44% shared. The screened sites were randomly distributed, but up to 40% of the sites showed a tendency to be densely distributed. In particular, a more concentrated distribution was confirmed for the 69 shared sites.

### Characteristics of the common binding sites of ORC2 and ORC4

Information on the 69 shared sites where type A and B genes of ORC2 and ORC4 commonly bound is summarized in Table 2. In addition, the peak data of the shared 69 sites obtained via ChIP-seq and the depth calculated based on the peak data are visualised in Figs. 5 and S11. The 69 shared sites ranged from 191–1284 bp in length, with an average length of 392.46 bp. The AT-richness of the 69 shared sites was distributed in the range of 44.39–52.92% (average: 49.01%). The depth value was calculated based on data obtained from the IP and Inp samples, and the average value derived from the four results corresponding to each sequence. As a result of comparing the maximum depth values of each shared site, the values were as low as 23.00 (site number 19, located at 319325–319758 bp in Chr 7) and as high as 179.00 (site number 49, located at 627717–627982 bp in Chr 19), and on average, the maximum depth value of the 69 sites was 59.77. For the minimum depth value, the values were as low as −11.75 (site number 26, located at 207738–207944 bp in Chr 14) and as high as 39.75 (site number 49, located at 627717–627982 bp in Chr 19), and on average, the minimum depth value of the 69 sites was 9.29. Therefore, from the results, it was revealed that the shared sites had the common characteristic of having an AT-richness of 49.01 ± 1.70% (the AT-richness of each results, ORC2A: 48.24 ± 2.19%, ORC2B: 47.74 ± 2.35%, ORC4A: 48.04 ± 2.31%, ORC4B: 47.77 ± 2.14%; the entire genome: 51.5%) (Scala *et al*., 2002). However, the length and depth of the sequence were not limited to a narrow range, and the differences between sites were clear.

**Table 2.**
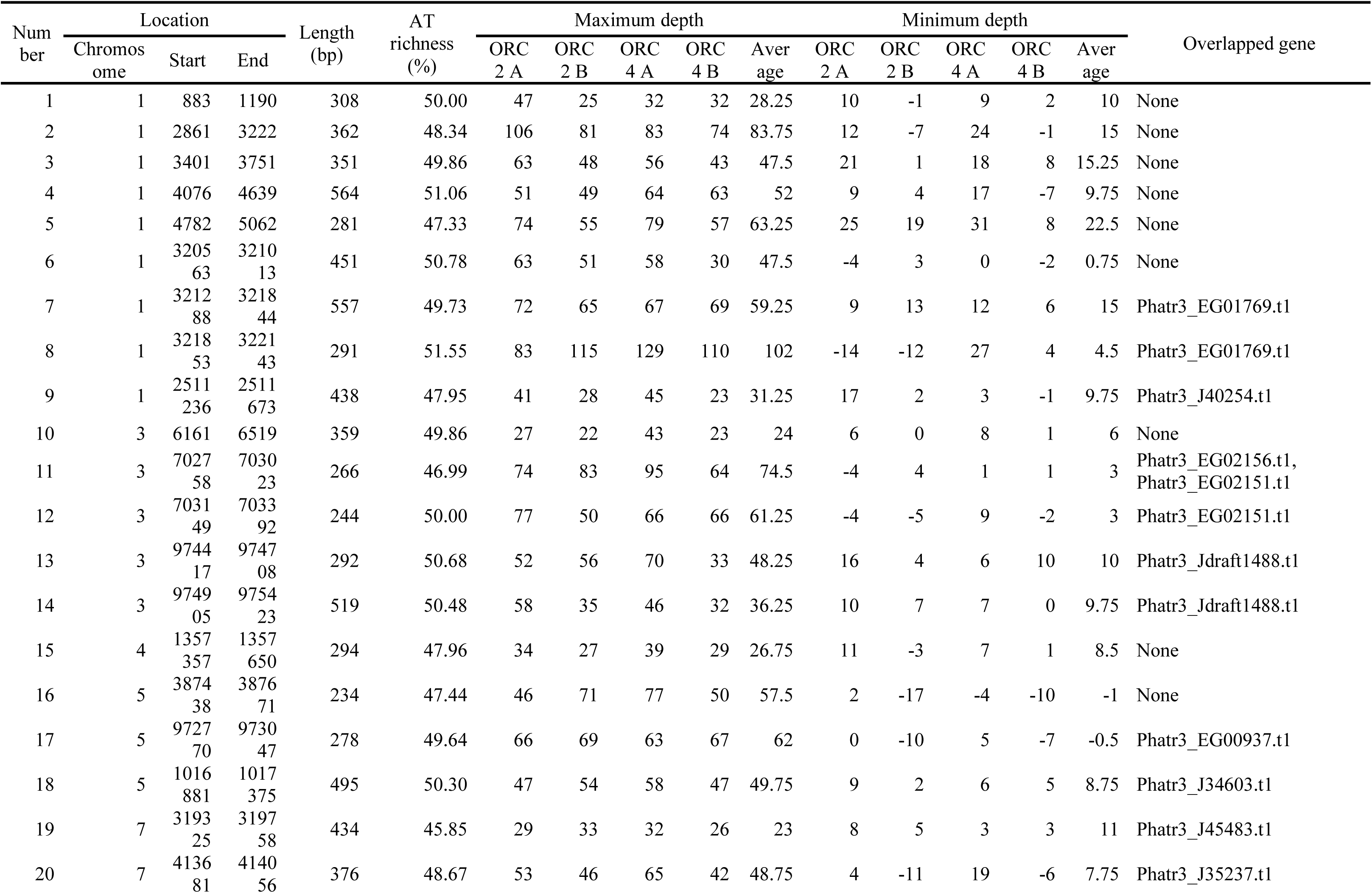
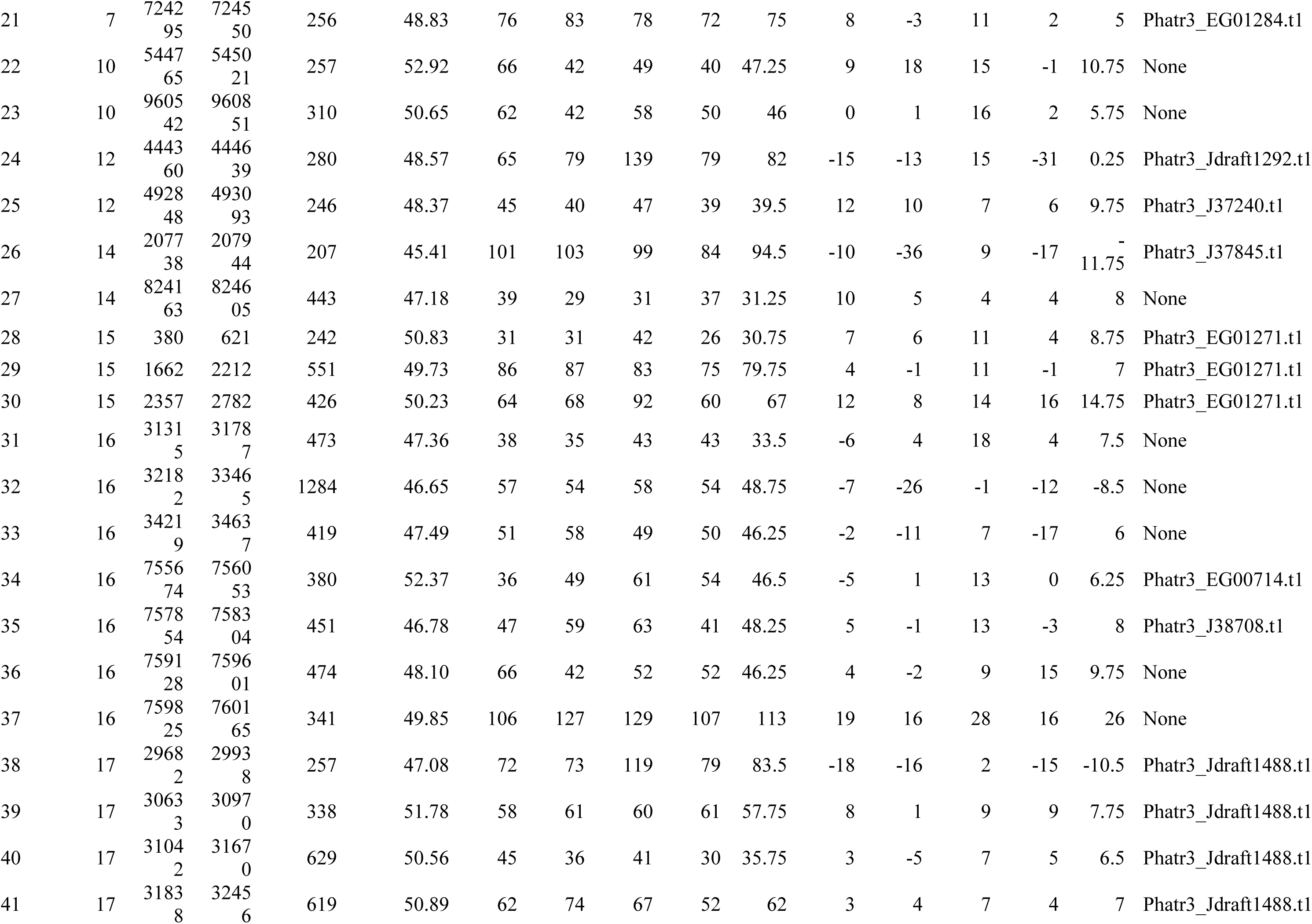
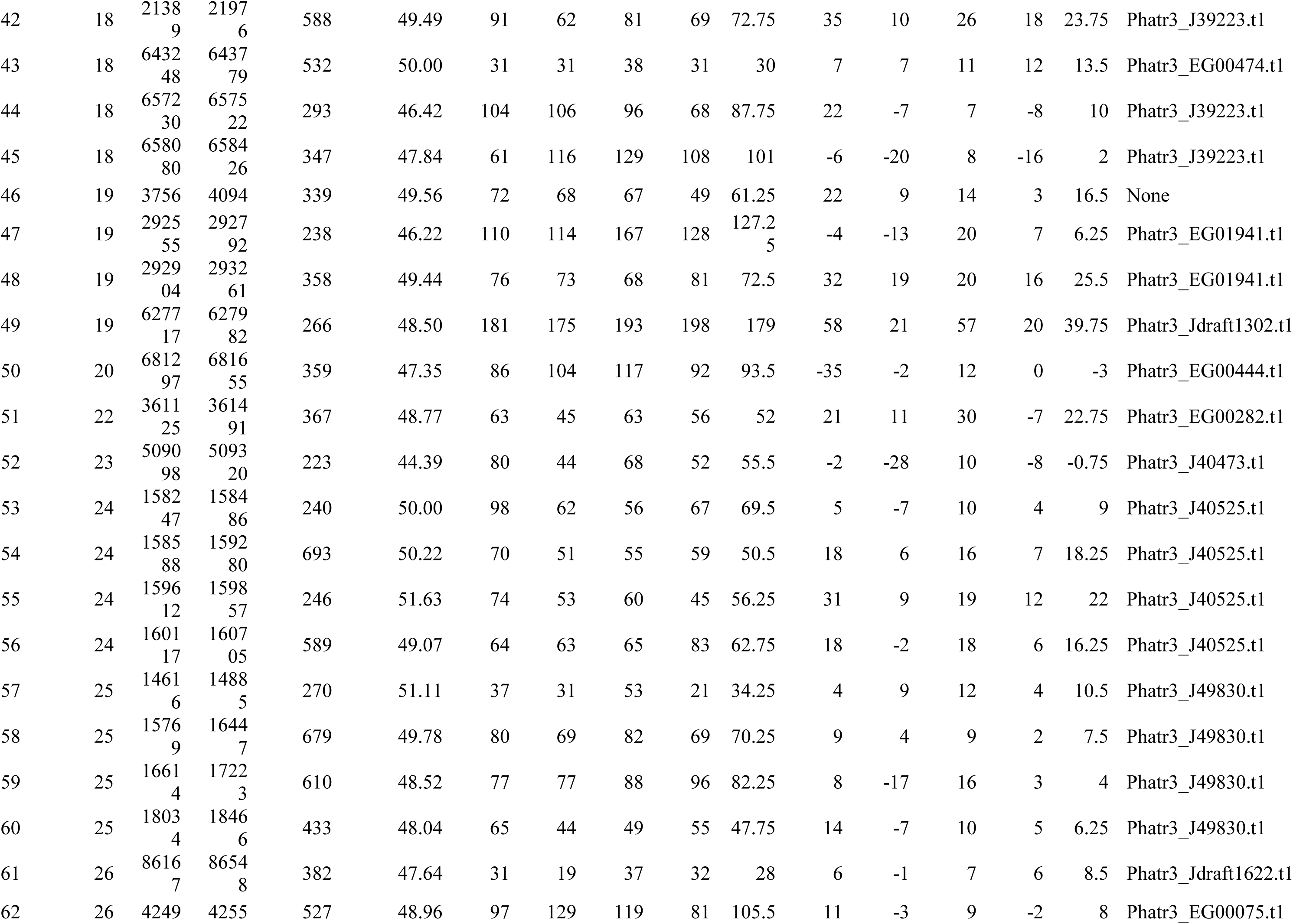
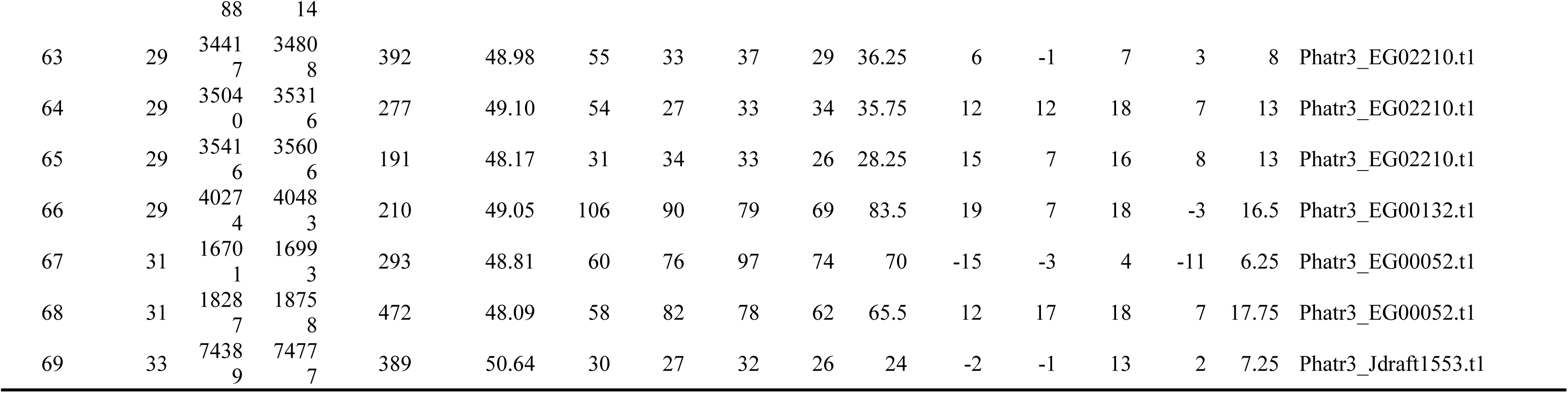
Information on shared sites.

Using the MEME Suite tool, motifs of shared sites were analysed, and the results are summarized in Fig. 6 and S12. Ten patterns of sequences expected to be motifs (motifs 1–10) were predicted. The AT-richness of the predicted motifs within the shared sites reached 54.85%, showing a relatively high value compared to that of the 69 shared sites (44.39–52.92%). Each predicted motif could be found in 4∼11 sites of the 69 shared sites. Although the motifs were shared by a relatively small number of sites, the sites sharing the motifs showed six patterns (patterns A–F) in its arrangement. In the case of pattern A, motifs 2, 3, and 5 in the forward direction (5′–3′) were followed by motifs 1, 4, and 9 in the reverse direction (3′–5′). Site numbers 24, 34, 45, and 67 corresponded to pattern A, and for site number 34, motif 9 was not included. In the case of pattern B, motifs 4 and 8 were arranged in the forward direction, followed by motif 10 in the forward direction and motif 7 in the reverse direction. Located between motifs 8 and 10 were motif 6 (number 43) in the forward direction, motif 2 (number 29, 32) in the reverse direction, or motif 7 (number 32) in the reverse direction. Site number 32 had two additional motifs 6 and 7, and sites number 65 and 68 showed an arrangement containing only motifs 4 and 8 as core motifs. Sites number 10 and 46, classified as pattern C, had an arrangement consisting of motif 1 in the forward direction, motif 3 in the reverse direction, motif 4 in the forward direction, and motif 2 in the reverse direction. Patterns D, E, and F had two motifs, each consisting of motif 9 in the forward direction and motif 6 in the reverse direction (pattern D: number 52, 63), motifs 9 and 3 in the reverse direction (pattern E: number 27, 36), and motif 1 in the forward direction and motif 3 in the reverse direction (pattern F: number 13, 40). Sites number 27 and 40 had motifs 4 (forward direction) and 3 (reverse direction) added to the core motif, respectively. Other sites that did not belong to patterns A–F had 1–4 motifs, but no pattern of motif arrangement could be grouped from them. These results revealed that sequences with relatively high AT-richness were predicted as motifs in shared sites, and that the predicted motifs were not randomly located. Furthermore, there were several patterns in the arrangement of motifs, which could be classified into six patterns.

**Fig. 6.**
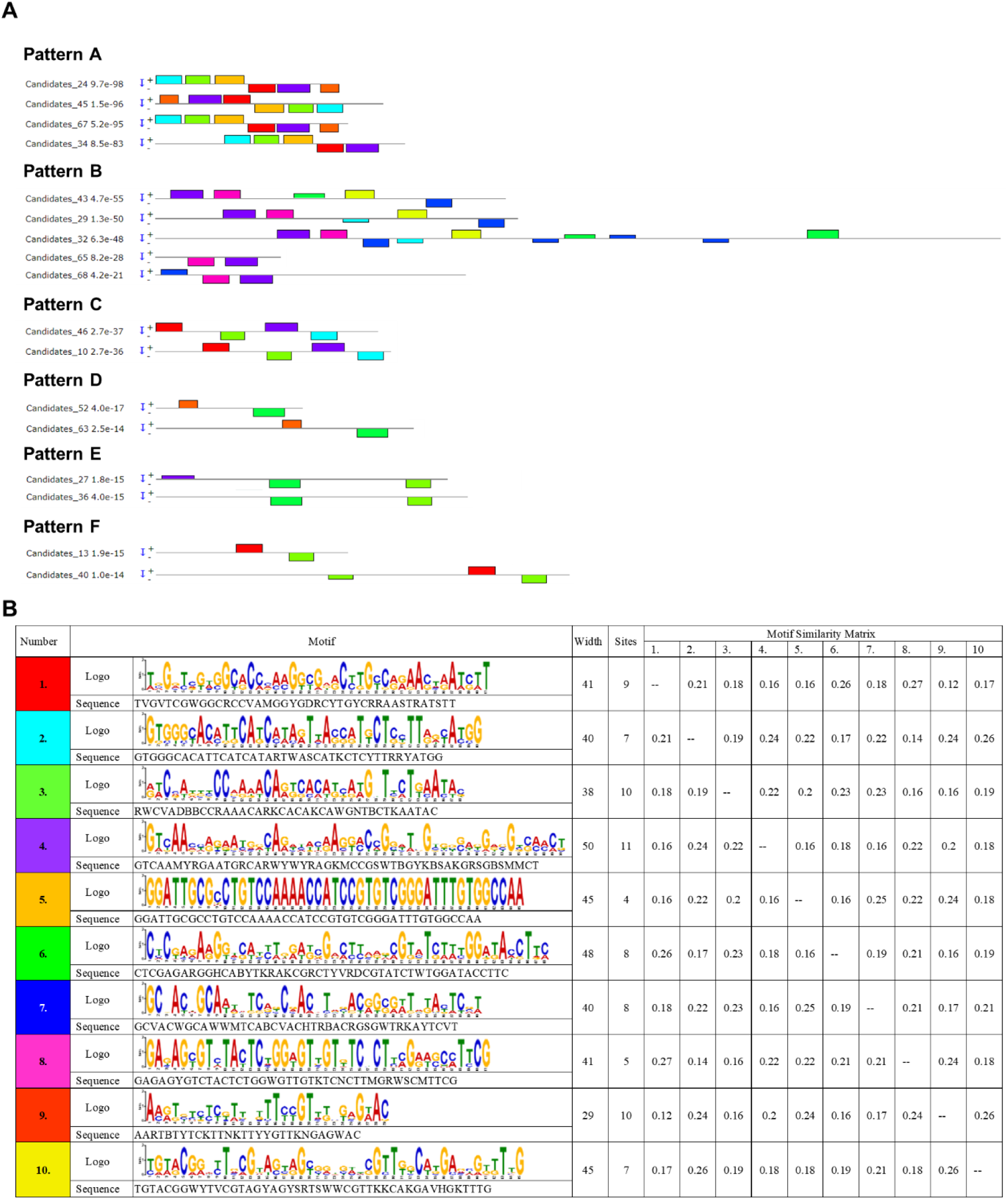
Arrangement patterns of motifs predicted from shared sites. Among the shared sites, there were those with the same arrangement pattern of predicted motifs, (A) which were classified into six types. (B) The sequence information of the arranged motifs and the similarity between motifs were organised.

In the distribution of the 69 shared sites on the genome of *P. tricornutum*, it was confirmed from the obtained results that 73.91% (51 sites) existed on the gene-coding region and overlapped in whole or in part (Fig. 5, S11, Table 2). However, none of the 355 sites, including the 69 shared sites, were distributed or overlapped with the centromere regions (Figs. 7, S13) (Diner *et al*., 2017). The results for each of the 25 centromere regions showed that the maximum depth value ranged from 0.75–14.75 and the minimum depth value ranged from −12 to −0.5 (except for the centromere of Chr 23, −105.25). Compared to the 69 shared sites (maximum depth value: 23–179), the highest maximum depth (14.75) obtained in the centromere region was relatively lower. In the case of the centromeres in Chr 7, 15, 16, and 23, there was an area where the gene-coding regions overlapped and peaks clearly detected. However, among the centromeres that did not contain the gene-coding region, that in Chr 4 was the only one in which a clear peak was detected. For the minimum depth value, that of the centromere converged to 0. The centromere in Chr 23 was an exception, where strong peaks were detected only from Inp samples in regions where the gene-coding region did not overlap, resulting in a minimum depth value of −105.25. Although relatively high depth values were obtained in regions where the centromere and gene-coding regions overlapped, the values were clearly lower compared to those of the 69 shared sites. In addition, depth values converging to 0 were obtained above the centromere region. Therefore, ORC2 and ORC4 are not expected to bind to centromeres. We also investigated whether the 69 shared sites and motifs within them were found in the genomes of other diatom species using D-genies and FIMO, respectively; however, none were identified.

**Fig. 7.**
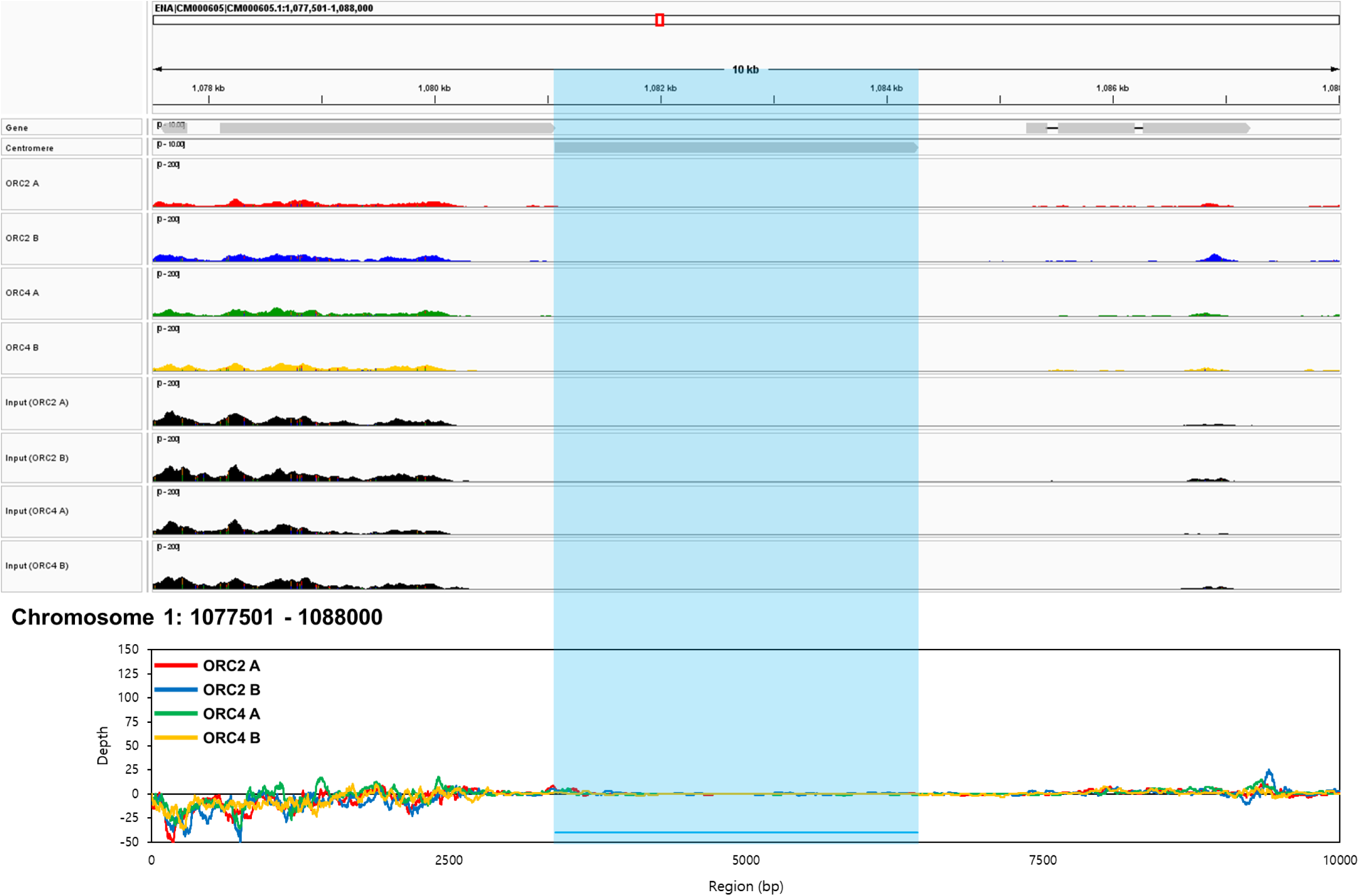
Peak data and depth values at the centromere region and surroundings of chromosome 1 in *Phaeodactylum tricornutum*. The results obtained from ORC2 A (red peaks and lines), ORC2 B (blue peaks and lines), ORC4 A (green peaks and lines), and ORC4 B (orange peaks and lines) were visualised through peak (IGV image) and depth images, and the centromere region indicated by a light blue line over the peak and depth images.

## Discussion

Based on data obtained via ChIP-seq, it was confirmed that the length of the ARS candidates screened in this study was mainly in the range of 100–399 bp (Fig. S9). This result shows a similar pattern to most human ARSs that have a length of less than 500 bp (Yu *et al*., 2022). In addition, for other eukaryotic organisms, including yeast, the length of ARSs was reported to be approximately 100–600 bp (Dao *et al*., 2021; Hu Yixin & Stillman, 2023). Referring to previous studies conducted on other eukaryotic organisms, the ARS of *P. tricornutum* is expected to have a length of 100–399 bp. In addition, ARSs over 400 bp in length are also expected. In humans, although there are many short ARSs present (less than 500 bp), long ARSs (500–2000 bp) also exist (Yu *et al*., 2022). Yeast (*S. pombe*) has ARSs ranging in length from 500–1500 bp (Hu & Stillman, 2023). Among the 69 shared sites, site number 32 (Chr 16: 32182–33465 bp) had a length of 1284 bp (Table 2). The peak and depth pattern of site number 32 was relatively dynamic, rising and falling throughout the region of the site (Fig. 5, S11). This dynamic pattern is in sharp contrast to that observed for site number 21 (724295–724550 bp in Chr 7), where only one clear peak was identified. In the case of sites number 7 (321288–321844 bp in Chr 1) and 8 (321853–322143 bp in Chr 1), they were screened individually through peak-calling, but a small gap (8 bp) was observed between the two sites. Considering all of these results, it seems likely that in the case of site number 32, which has a length longer than 1 kbp, several ARS candidates were closely packed and peak-called as one candidate (Nakato & Sakata, 2021). In addition, referring to previous research reports on the structure of the DNA-binding modules, as the length of the sequence that an ORC binds to is very limited, it is possible that a 1-kbp ARS contains many non-essential sequences (Feng *et al*., 2021). Based on our results, we suggest that the ARSs of *P. tricornutum* are mainly less than 500 bp in length, and the small number of ARSs with lengths more than 500 bp are likely to be clusters of closely distributed ARSs.

The 355 sites screened in this study are distributed randomly on the genome, with some sites showing a dense distribution (Fig. S10). In previous studies, a random distribution of ARSs has been observed in the genomes of eukaryotic organisms, including humans, with patterns in which they are sparse in some regions and extremely concentrated in others (Yu *et al*., 2022). Among these distribution patterns, the concentrated distribution of ARSs in specific regions is referred to as a zone in DNA replication, and the characteristics of each zone can be observed in the S phase during cell division (Gaboriaud & Wu, 2019; Guilbaud *et al*., 2022; Hu & Stillman, 2023). When DNA replication begins in cells that have entered the S phase from the G1 phase, the entire genome is not replicated simultaneously, but instead replicated sequentially in each region (Gaboriaud & Wu, 2019; Hu & Stillman, 2023). In relation to this, each zone is called an early-replicating, mid-replicating, and late-replicating zone, depending on when replication begins from the beginning to the end of the S phase (Gaboriaud & Wu, 2019; Hu & Stillman, 2023). Even if ARSs are grouped in the same zone, there are differences in their activity within the zone, and they can be classified into efficient and inefficient ARSs (Gaboriaud & Wu, 2019; Guilbaud *et al*., 2022). Among our results, shared sites number 1–5 are expected to represent examples of a replicating zone composed of efficient and inefficient ARSs (Table 2). In this expected zone, sites with different levels of ORC subunit communication are densely distributed, with a maximum depth ranging from 28.25–83.75 (number 1: 28.25, number 2: 83.75, number 3: 47.50, number 4: 52.00, number 5: 63.25) (Fig. S11). Although each ARS has the functionality of being the origin of DNA replication, it has been suggested that their dense distribution may act as a factor to enhance their functionality (Rhind, 2014). However, there is an opposite case in which yeast episomal vectors can be stabilised by only one efficient ARS (Lopez *et al*., 2021). Therefore, the distribution of ARSs in *P. tricornutum* and their significance for DNA replication are considered to require further study. In summary, as with other eukaryotic organisms, the ARSs of *P. tricornutum* tend to be distributed randomly and include those that are clustered together closely. A dense group of ARSs is expected to represent a replicating zone and be composed of ARSs at various levels in communication with the ORC subunit.

The 69 shared sites showed a commonality of AT-richness close to 50% (44.39–52.92%, 49.01 ± 1.70%) (Table 2). According to previous reports, AT-richness is an important factor in ARSs (Hu & Stillman, 2023). Yeast (*S. pombe* and *S. cerevisiae*) ARSs are rich in AT, whereas metazoan ARSs are rich in GC, and each eukaryotic organism has unique characteristics (Hu & Stillman, 2023). Referring to this, the characteristic of not being biased towards either AT or GC is expected to be a point that distinguishes ARSs of *P. tricornutum* from those of other organisms. Ironically, the yeast ARS was applied in an attempt to design an episomal vector for microalgae, which reported some successful results (Karas *et al*., 2015). This fact suggests that an AT-rich yeast ARS can function in *P. tricornutum* (Karas *et al*., 2015), which is contrary to the 50% AT-richness of ARS candidates revealed in our study. However, although the yeast ARS, which has a relatively high AT-richness, provides an ARS function to episomal vectors, it is limited in performing its complete function; therefore, it is assumed that at least the native ARS of *P. tricornutum* will have characteristics (including AT-richness) that are distinct from those of the yeast ARS (Kumar *et al*., 2020).

As episomal DNA containing yeast CEN/ARS could be stably replicated in *P. tricornutum* (Diner *et al*., 2016), we predicted that the ARSs of *P. tricornutum* would also contain the ARS consensus sequence (ACS) similar to that of yeast ARSs (Hu *et al*., 2020; Tan *et al*., 2021). Therefore, we expected that ACS candidates would be found in the 69 shared sites via motif analysis (Fig. 6, S12). However, the predicted motif results were contrary to our expectations. The predicted motifs were shared at 4–11 sites among the 69 sites (Fig. 6). Furthermore, there were no conserved or shared sequences in all 69 sites. Considering these results, it is highly likely that an ACS does not exist in the ARS of *P. tricornutum*, and it is expected that it is not included in the factors that enable yeast CEN/ARS to be recognised by the ORC of *P. tricornutum*. Previous studies have reported that the yeast species, *S. cerevisiae* and *S. pombe*, have AT-rich ARSs, but that the ORC of *S. cerevisiae* recognises a specific sequence (e.g., ACS), whereas that of *S. pombe* recognises a non-specific AT-rich sequence (Hu & Stillman, 2023; Lee *et al*., 2023). Additionally, the metazoan ARS does not have an ACS that is found in the *S. cerevisiae* ARS; therefore, non-specific sequences are recognised by the ORC (Hu & Stillman, 2023; Lee *et al*., 2023). In light of our results and previously reported cases, the ARSs of *P. tricornutum* are expected to have non-specific sequences recognised by an ORC. Our results revealed the existence of a pattern in the arrangement of predicted motifs (Fig. 6). Among the 69 sites, several (2–5 sites) were grouped according to the motifs they contained. Each of these pattern groups had similar sequences, at least in terms of the predicted motif sequences. Interestingly, although a motif sequence was shared by each group, the maximum depth value of each site belonging to the same group was not unified (Table 2). For the type A pattern group, where most of the site consisted of motif sequences (sequences shared at a relatively high rate), maximum depth values ranged from 46.5 (number 34) to 101 (number 45). Similarly, for pattern B (28.25–79.75), C (24–61.25), D (36.25–55.5), E (31.25–46.25), and F (35.75–48.25) types, there was no tendency for maximum depth values to be unified among sites belonging to the same group. Although the sequences were relatively shared, differences in depth values existed in each group, suggesting that the specific sequence might not be a factor in determining the ARS of *P. tricornutum*, or at least not significantly so. Based on this, we propose that the ORC2 and ORC4 of *P. tricornutum* do not require specific sequences to recognise ARSs, and further support that the sequences of ARSs in *P. tricornutum* are not conservative.

Our study revealed that the ORC2 and ORC4 of *P. tricornutum* did not bind to centromeric regions (Fig. 7, S13). Previous studies on the non-replicative functions of the ORC have revealed that it performs various roles in addition to the function of recognising replication origins, such as binding to centromeric regions (Popova *et al*., 2018). Interestingly, *Drosophila* and mammalian ORCs interact with centromeric regions, whereas yeast ORCs do not (Popova *et al*., 2018). Furthermore, a centromeric replication origin was reported to exist in the mammalian centromere (Pelletier *et al*., 1999; Massey & Koren, 2022). In other words, there is a centromere that can be recognised by ORC, which contains a region that can play the role of an ARS (Popova *et al*., 2018; Massey & Koren, 2022). Comparing our results with those of previous studies, it is expected that the centromeric regions of *P. tricornutum* do not contain ARSs and do not exhibit non-replicative functions (Popova *et al*., 2018; Massey & Koren, 2022).

In summary, the ORC2 and ORC4 from *P. tricornutum* used in this study, and the results obtained from them, were heterogeneous when compared to those of eukaryotic organisms. Moreover, their amino acid sequences were distinct from those of other eukaryotic organisms. Additionally, in *P. tricornutum*, it is notable that the sites screened from ORC2 and ORC4 were not unified and shared at a low level. The sequences of the screened sites were neither AT-rich nor GC-rich, and neither were they conserved nor unified. Furthermore, the screened sites had the mixed characteristics of yeast and metazoan ARSs. Ultimately, the ARS candidates we screened showed complex and mixed characteristics of ARSs, as revealed in previous research. In addition to screening ARS candidates, our study presents those with potential for use in combination with centromeres in episomal plasmids. Overall, these findings suggest candidates that could be used as native ARSs in episomal plasmids for *P. tricornutum* and reveal their characteristics.

## Conclusion

In this study, we screened potential ARS candidates using the native ORC2 and ORC4 from *P. tricornutum*. ORC2 and ORC4 bound to 355 sites on the *P. tricornutum* genome, and 69 of these were commonly recognised. Binding sites were mainly less than 500 bp in length with an AT-richness of 49.01 ± 1.70%. There were no conserved sequences or shared motifs observed between sites. Furthermore, because the binding frequency of ORCs differs between sites with similar sequences, the native ORC of *P. tricornutum* was expected to bind sequences in a non-specific manner. The screened sites were randomly distributed but contained a dense distribution in some regions. Interestingly, ORC2 and ORC4 did not bind to the centromere; therefore, no interactions between them were expected. From our results, we were able to observe characteristics that were similar in some parts to those of yeast ARSs and in other parts to metazoan ARSs. Based on these results, we suggest that the native ARS of *P. tricornutum* has complex and unique characteristics compared to those of other eukaryotic organisms. Furthermore, our study provides unique information and insight into the ARS of episomal vectors for diatoms.

## Acknowledgments

This work was supported by a JSPS KAKENHI Grant-in-Aid for Scientific Research (grant number 21K04784 (Y.M. serves as a principal investigator) and 24H00392 (Y.M. serves as a co-investigator) and H.-S.Y. thanks to the financial support of JST SPRING (grant number JPMJSP2124).

## Author contributions

H-SY: investigation, methodology, formal analysis, writing original draft, visualization

KY: investigation, methodology, formal analysis, writing–review & editing

TS: investigation, methodology, formal analysis, writing–review & editing

IS: methodology, formal analysis, writing–review & editing

YM: conceptualization, methodology, formal analysis, writing–review & editing, funding acquisition.

All authors read and approved the final manuscript.

## Data availability

The ChIP-seq project data were deposited in DDBJ under BioProject accession number PRJDB18238 and BioSample accession numbers SAMD00791632 (ORC2A, IP), SAMD00791633 (ORC2A, Inp), SAMD00791634 (ORC2B, IP), SAMD00791635 (ORC2B, Inp), SAMD00791636 (ORC4A, IP), SAMD00791637 (ORC4A, Inp), SAMD00791638 (ORC4B, IP), and SAMD00791639 (ORC4B, Inp). The reads data are available under DRA Run accession numbers DRR571401 (ORC2A, IP), DRR571402 (ORC2A, Inp), DRR571403 (ORC2B, IP), DRR571404 (ORC2B, Inp), DRR571405 (ORC4A, IP), DRR571406 (ORC4A, Inp), DRR571407 (ORC4B, IP), and DRR571408 (ORC4B, Inp).

