## Supplementary for "Genome-wide mapping of autonomously replicating sequences in the marine diatom *Phaeodactylum tricornutum*"

**Supplementary Information Appendix: Materials and Methods**

**Estimation of the diatom genes encoding ORC2 and ORC4**

We performed a BLASTp search to estimate the genes encoding ORC2 and ORC4 using yeast ORC2 (NP 009616.1) and ORC4 (AJV92690.1) as query sequences, respectively. The amino acid sequences of ORC2 and ORC4 were used for phylogenetic analysis, and a phylogenetic tree constructed using MEGAX (Kumar *et al.,* 2018). Based on the LG+G+F model, the phylogenetic tree was generated using maximum likelihood parameters with 1000 bootstrap replications. Regarding ORC and ARS candidates, amino acid and DNA sequence motifs were analysed through a web-based tool (MEME Suite; https://meme-suite.org) (Bailey *et al.,* 2015).

**Western blotting for confirmation of expression levels in transformants**

For western blotting, *P. tricornutum* transformant clones from well plates were transferred to flasks containing fresh f/2 medium with 50 µg/ml zeocin. Vegetative cells in the growth phase ($OD_{750}$: 0.3–0.4, 3.24 × $10^6$ cells/mL) were collected via centrifugation, and the medium components washed with water. Transformant pellets (1 × $10^7$ cells) were used for western blotting. Pellets were suspended in 75 µl of 1% (w/v) sodium dodecyl sulphate (SDS) solution, and 25 µl of 4× Laemmli loading buffer (250 mM Tris-HCl [pH 6.8], 40% [v/v] glycerol, 0.01% [w/v] bromophenol blue, 8% [w/v] SDS, and 20% [v/v] β-mercaptoethanol) added thereafter. The mixtures were heated at 100°C for 10 min and centrifuged (20500 × *g*, 5 min). The denatured proteins contained in the supernatant were separated through SDS-polyacrylamide gel electrophoresis (350 V, 25 mA, 3 h) with a 15% (w/v) gel (e-PAGEL; ATTO, Tokyo, Japan). The separated proteins were transferred to a membrane (Immobilon-P Transfer Membrane; Sigma, Billerica, MA, USA) and soaked in transfer buffer (40 mM Tris, 242 mM glycine, 0.1% [w/v] SDS, 5% [v/v] methanol) via electroblotting (150 V, 1 h) using a blotting system (WSE-4040 HorizeBLOT 4M-R; ATTO). The protein-transferred membrane was coated with generic proteins using a blocking solution (phosphate-buffered saline [PBS] with 5% [w/v] skim milk) for 1 h at room temperature (25°C). The coated membrane was treated with a primary antibody (anti-DYKDDDDK tag, 66008-4-Ig; Proteintech, Rosemont, IL, USA) diluted in PBS at a ratio of 1:5000 and incubated for 1 h at room temperature. Excessive primary antibodies that did not attach to the target protein were removed via washing with PBST (PBS with 0.1% Tween20) three times for 10 min. The washed membrane was then treated with a secondary antibody (horseradish peroxidase-conjugated anti-mouse IgG [H+L]; Promega, Madison, WI, USA) diluted in PBS at a ratio of 1:2500 and incubated for 1 h at room temperature. Excessive secondary antibodies that did not attach to the primary antibody were removed via washing with PBST three times for 10 min. To visualise the expression of FLAG-tag-fused ORC2 and ORC4, the membrane was treated with a substrate solution (Metal Enhanced DAB Substrate Kit; Thermo Fisher Scientific, Waltham, MA, USA) and incubated for 1 min.

**Supplementary Information Appendix: Results**

**Phylogenetic relationships and similarities of ORCs**

Our BLASTp search predicted Phatr3_J42935 and Phatr3_EG01462 as the genes encoding ORC2 and ORC4, respectively. The phylogenetic relationship between eight species harbouring ORC2 and ORC4 is visualised in Fig. 1. On the phylogenetic tree, the ORC2 and ORC4 groups are clearly separated, and ORCs of metazoan (*Homo sapiens*, *Danio rerio*, *Drosophila melanogaster*), plant (*Arabidopsis thaliana*, *Zea mays*), diatom (*Phaeodactylum tricornutum*, *Fistulifera solaris* [allopolyploid]) (Maeda *et al.,* 2022), and yeast (*Saccharomyces cerevisiae*) species are grouped respectively. In the aligned amino acid sequences of the ORCs (Fig. S1, S2), the results showed that although phylogenetic differences existed between the groups, the sequences were conserved. In particular, the sequences where the winged helix and AAA+ domains of ORC2 and ORC4 are located were more conserved when compared to other parts of the sequence (Bleichert *et al.,* 2015; Li *et al.,* 2018; Cheng *et al.,* 2020; Schmidt & Bleichert, 2020). Furthermore, the motif sequences predicted from eight species of ORCs were also mainly located in conserved regions. Similar to the grouping results in the phylogenetic tree, the motif arrangement pattern was relatively similar among metazoan, plant, and diatom species, and that of the yeast ORCs relatively distant from the other seven species. There were variations in the sequences of ORC2 and ORC4 depending on the taxonomy, but they showed an overall conservative trend. The close relationship of the predicted diatom ORC2 and ORC4 with those of other organisms, as well as the conservation of functionally important domains, encouraged us to use Phatr3_J42935 and Phatr3_EG01462 as the diatom ORC2- and ORC4-encording genes for subsequent investigation.

**Raw data obtained via ChIP-seq with an Illumina sequencer**

Genomic DNA was extracted from transformants, sheared during sonication, and subjected to ChIP treatment, from which library samples were created. Prior to analysis using the Illumina sequencer, it was confirmed via Bioanalyzer Systems that the library samples mainly consisted of fragments of approximately 200–1000 bp in size (Fig. S8). Information on the prepared library samples and results analysed using the Illumina sequencer are summarized in Table 1. The concentrations of the prepared library samples were variously distributed within the range of 71.00–388.60 nM, and thus the samples with different concentrations were diluted to a concentration of 2 nM and analysed using an Illumina sequencer. The number of reads obtained from each library sample at a unified concentration exceeded one million, with as few as 1445983 reads (ORC2 A Inp) and as many as 1949374 reads (ORC2 B Inp) obtained. Via Bowtie2, among the obtained reads, those that were not mapped onto the reference genome data of *P. tricornutum* were excluded. Ultimately, over one million reads mapped to the genome were obtained from all samples, with as few as 1264133 reads (ORC2 A Inp) and as many as 1714406 reads (ORC2 B Inp) mapped. Peak-calling was performed through MACS2 from the mapped reads, and by pairing IP and Inp samples, it was confirmed that 140–206 sites (ORC2 A: 163 sites, ORC2 B: 206 sites, ORC4 A: 140 sites, and ORC4 B: 162 sites) were peak-called on the genome of *P. tricornutum*. More than 80% of peak-called sites (ORC2 A: 86.54%, ORC2 B: 92.27%, ORC4 A: 81.45%, and ORC4 B: 87.50%) were distributed in the length range of 101–400 bp (Fig. S9), and in particular, sites with a length of 101–200 bp (ORC2 A: 30.13%, ORC2 B: 45.89%, ORC4 A: 27.42%, and ORC4 B: 24.38%) or 201–300 bp (ORC2 A: 39.74%, ORC2 B: 38.16%, ORC4 A: 37.10%, and ORC4 B: 50.63%) were the most abundant. The average length of the sites was 241.02–317.58 bp (ORC2 A: 295.28 bp, ORC2 B: 241.02 bp, ORC4 A: 317.58 bp, and ORC4 B: 277.87 bp).

**Supplementary Information Appendix: Discussion**

In previous studies conducted on the yeast ORC, transformants designed to express an ORC subunit with additional mutations resulted in a more suppressed growth pattern than that in the WT (Klemm & Bell, 2001; Hu *et al.,* 2020). Similar to studies conducted on yeast, growth of the *P. tricornutum* transformants used in this study showed a pattern of inhibition when compared to that in the WT (Fig. S6). Overexpression of the FLAG-tag-fused ORC subunit was suggested to be the likely cause of the growth inhibition observed in the transformants. However, the N- and C-terminus tag proteins fused for ChIP-seq did not appear to cause significant changes in ORC function (Klemm & Bell, 2001; Amin *et al.,* 2020; Hu *et al.,* 2020; Li *et al.,* 2022). Considering that ORCs have non-replicative functions, such as gene silencing and transcriptional regulation, in addition to the function of recognising replication origins (Chesnokov, 2007; Sasaki & Gilbert, 2007), there is a possibility that overexpression of ORC2 and ORC4 in *P. tricornutum* is involved in similar functions. Unfortunately, due to the absence of previous studies on native ORC subunits in *P. tricornutum*, information on the impact of overexpressed ORC2 and ORC4 is unclear. We propose that the presence of an ORC subunit expression cassette in the introduced vector is likely to be related to the growth inhibition pattern observed for the transformant.

In the present study, we found that the *P. tricornutum* CCAP 1055/1 strain had two allelic gene types (types A and B) of ORC2 and ORC4 (Fig. S3, S4). We expected that the four types of ORC subunits could interact with the same locations on the genome of *P. tricornutum*. However, among the 355 sites identified, only 69 (19.43%) were commonly mapped in our ChIP-seq analyses for these four subunits. By contrast, previous ChIP-seq analyses in a budding yeast (*Torulaspora delbrueckii*) revealed that the recognition sites of ORC subunits (namely, ORC1, ORC2, and ORC4) were shared at a high rate (64.63%) (Maria *et al.,* 2021). Therefore, the proportion of the shared sites in *P. tricornutum* was relatively low compared to that in *T. delbrueckii*. As each ORC subunit cannot independently interact with the ARS, divergence of the mapped sites of ORC2 and ORC4 in the present study suggest that we could not perfectly eliminate non-specific detection during the ChIP-seq experiments. Along with non-specific detection, divergence of the mapped sites between the two types of each subunit might be attributable to the functional divergence between the allelic genes encoding ORC subunits that had been reported previously in plants. González et al. (2020) reported that knocking out one allelic gene of the ORC1 subunit in *A. thaliana* interferes with normal development (González *et al.,* 2020), indicating that allelic ORC1 genes do not completely complement each other. It remains elusive whether similar functional differences exist in *P. tricornutum*. At least both ORC2 and ORC4 alleles are expected to be expressed (Hoguin *et al.,* 2021). According to previous studies on allele-specific expression in *P. tricornutum*, 1% of genes show biased expression (Hoguin *et al.,* 2021). When referring to the data of Hoguin et al. (2021) (NCBI accession ID: SRX2578671), it was confirmed that alignments matching the sequences of both alleles of ORC2 and ORC4 were included.

Although ORC2 and ORC4 bind to ARSs and form a complex with other subunits, it is ironic that only 69 of the 355 sites screened are shared (Fig. 3). A and B types exist in each ORC subunit, and the differences in amino acid sequence between allelic genes are small. There was one amino acid difference in the region expected to comprise a WH domain, and two (ORC2) and five (ORC4) amino acid differences in the region expected to represent an AAA+ domain. However, depending on the position of the amino acid sequence, the substitution of only 1–2 amino acids could affect the function of the ORC subunit, which can also lead to changes in the motif of the DNA sequence recognised by an ORC (Chesnokov *et al.,* 2001; Hu *et al.,* 2020). Therefore, it is possible that the binding sites of each A and B type of ORC2 and ORC4 do not match perfectly due to differences in amino acid sequence between the allelic genes. However, proteins binding to ARSs may not always attach to common sites (Masuda *et al.,* 2020). ORC and minichromosome maintenance (MCM) proteins have the common characteristic of binding to ARSs, but in previous studies, sites recognised by *Schizosaccharomyces pombe* ORC4 and MCM2 were compared with those recognised only by ORC4 (Masuda *et al.,* 2020). Interestingly, there were differences in characteristics, including motifs, between groups of binding sites (Masuda *et al.,* 2020). Similarly, because sequences within the ARSs where each ORC subunit binds are different (Feng *et al.,* 2021), there may be a site that can be screened only from ORC2 or ORC4. Furthermore, even in the S phase, when DNA replication occurs, the ARSs screened from early and late S phases may be different (Guilbaud *et al.,* 2022). According to previous research on human ARSs, there is a clear difference in the number of ARS sites that can be screened between human cells 15 min after entering the S phase from the G1 phase and cells 3 h after entering the S phase (Guilbaud *et al.,* 2022). In addition, it is notable that there are many sites that are screened only in cells that have passed for 15 min and in cells that have passed for 3 h (Guilbaud *et al.,* 2022). Furthermore, another previous study reported that differences may occur between ChIP-seq results obtained from synchronized and unsynchronized cultures (Petrie *et al.,* 2023). Therefore, it is expected that the cell cycle may influence the sites that can be screened. In this study, cells in log phase ($OD_{750}$: 0.3–0.4) were used for analysis without being synchronized; therefore, it is likely that cells in the G1, S, G2, and M phases were mixed. It is possible that the differences observed in results were derived from unsynchronized samples. Lastly, the peak-calling process may be one of the causes related to the inaccuracy of ChIP-seq results (Chen *et al.,* 2021; Nakato & Sakata, 2021). MACS2, one of the popular peak-calling tools, has limitations in accuracy, and the need for a more improved tool has been suggested (Chen *et al.,* 2021; Nakato & Sakata, 2021). In the present study, ARS candidates were screened via peak-calling using MACS2. The peak-calling process may also be a factor causing differences between results. To summarize the expected causes of differences observed between results, we found differences in the amino acid sequence between each allele type (including SNPs), differences in the sequences of DNA binding to ORC2 and ORC4, the effects of an unsynchronized sample, and errors occurring from peak-calling. For these reasons, we suggest that shared results were obtained for only 19.44% of the sites screened.

In metazoan ARSs, there is no ACS or shared motif, and G-rich (or GC-rich) regions were recognised by an ORC (Hyrien, 2015; Ganier *et al.,* 2019). Furthermore, histone modifications are involved in the accessibility and activity of replication origins (Hyrien, 2015; Ganier *et al.,* 2019). In other words, ARSs recognised by an ORC, which recognises non-specific sequences, have site-specific characteristics (Shen *et al.,* 2010; Wang *et al.,* 2021). In the genome of a eukaryotic organism, regions that satisfy these elements are mainly where genes and promoters are located (Prioleau & MacAlpine, 2016; Ganier *et al.,* 2019; Hu & Stillman, 2023). Interestingly, our results showed that 73.91% of the 69 shared sites overlapped with the region where the gene was located (Table 2), and several sites that did not overlap with the gene region were located upstream of the gene, where the gene promoter is likely to be located (Fig. S11). These results are expected to show a similar trend compared to the phenomenon where the ORCs of other eukaryotic organisms recognise their native ARSs.

For yeast species that do not have a centromeric replication origin, in order to create a transformant by introducing an episomal vector, the vector must contain sequences that play the role of the ARS and centromere (Liachko *et al.,* 2011; Lefrançois *et al.,* 2013). The absence of either an ARS or centromere makes it impossible or difficult to achieve vector stability (Liachko *et al.,* 2011; Lefrançois *et al.,* 2013). As with yeast, *P. tricornutum*, which is not expected to have a centromeric replication origin, is expected to require both ARS and centromere sequences for stably maintaining episomal plasmids (Diner *et al.,* 2016; Diner *et al.,* 2017). As mentioned earlier, although the yeast CEN/ARS sequence is not ideal, it successfully provided stability to an episomal plasmid in *P. tricornutum* (Diner *et al.,* 2016). Furthermore, episomal vectors with the native centromeres (e.g., centromeres of Chr 25 and 26) of *P. tricornutum* showed complete stability (Diner *et al.,* 2017). In a previous study, only plasmids containing fragments (from Chr 25 and 26) expected to contain the centromere were stable (Diner *et al.,* 2017). Unfortunately, only the stability of plasmids derived from the centromere was mentioned, and there was no discussion about the ARS (Diner *et al.,* 2017). Interestingly, our results screened ARS candidates (Chr 25: site number 57–60, Chr 26: site number 62) near the centromere regions of Chr 25 and 26 (Fig. 4, S10; Table 2). According to our results, centromeres of Chr 25 and 26 do not interact with the ORC that recognises ARSs (Fig. 7, S13). Therefore, we suggest that ARS sequences (including the screened shared sites), as well as centromeres of Chr 25 and 26 (absence of a centromeric replication origin), may have contributed to the plasmid stability observed in the previous study (Diner *et al.,* 2017). In addition, from previous research results, we assumed that plasmids containing only ARSs cannot be stable (Diner *et al.,* 2017). In particular, Chr 26 contained a shared site number 61 located far from the centromere (Table 2). From this, it is expected that stable maintenance cannot be expected if the episomal plasmid introduced into *P. tricornutum* contains only an ARS. Based on our results, we suggest that the combination of a centromere and ARS, rather than an ARS alone, is required to provide stability to episomal plasmids in *P. tricornutum*. Furthermore, ARS candidates were found near the centromeres of Chr 25 and 26 in *P. tricornutum*, and we suggest that these candidates may be useful ARS resources for episomal plasmids expressed in *P. tricornutum*.

**Supplementary Information**

**Supplementary Figures**

**>ORC2**

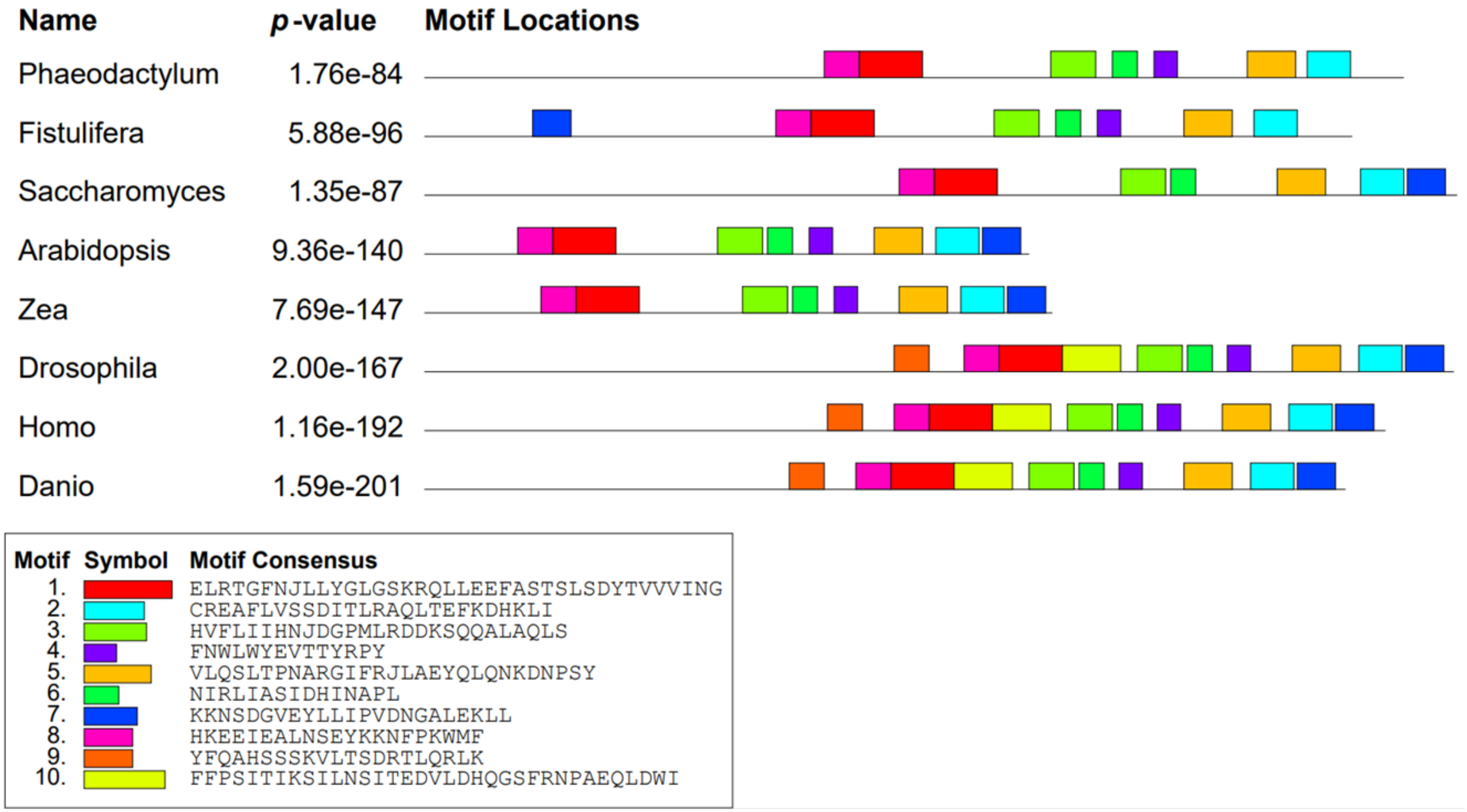

**1.**

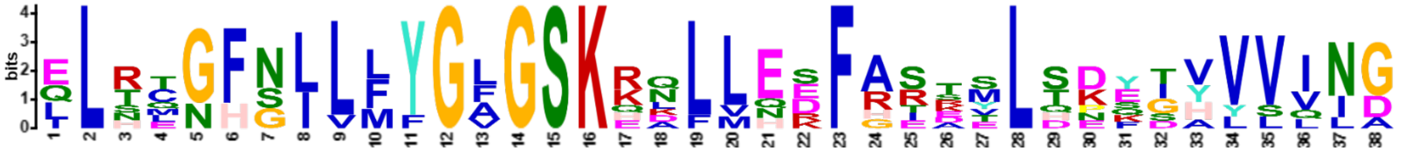

| Name | Start | *p*-value | | Sites | |
|---|---|---|---|---|---|
| 5. Zea | 92 | 1.58e-39 | ykdqyrnwlf | ELRCGFGLLMYGFGSKKQLLEDFASTTLTNFTVVVING | ylpavnlkqv |
| 4. Arabidopsis | 78 | 3.15e-37 | yktmyskwvf | ELRCGFGLLMYGFGSKKALVEDFASASLTDYSVVVING | ylpsvnlkqv |
| 8. Danio | 281 | 4.48e-35 | heknfrqwml | QLRMGFNILLYGLGSKQLLLEKFRSSMLSDYDHVVVNG | ffpsitlksi |
| 7. Homo | 304 | 6.64e-34 | yeklfhkwml | QLHLGFNIVLYGLGSKRDLLERFRTTMLQDSIHVVING | ffpgisvksv |
| 6. Drosophila | 346 | 3.67e-30 | yrsyfpkwmc | ILNEGFNILLYGLGSKHQLLQSFHREVLHKQTVLVVNG | ffpsltikdm |
| 1. Phaeodactylum | 262 | 1.72e-28 | LQERFPEWRF | LLSTNHSLLFYGAGSKRNLMNEFARRELDKEGYVVIID | STDPDVNINA |
| 3. Saccharomyces | 307 | 2.63e-25 | qkkmfpqywf | ELTQGFSLLFYGVGSKRNFLEEFAIDYLSPKIAYSQLA | yenelqqnkp |
| 2. Fistulifera | 233 | 1.09e-24 | LERNFDEWRF | LLTTNHNLLFFGAGSKEKLLHSFGEKSLSEEGYVLLID | GFAKDVSVTA |

**2.**

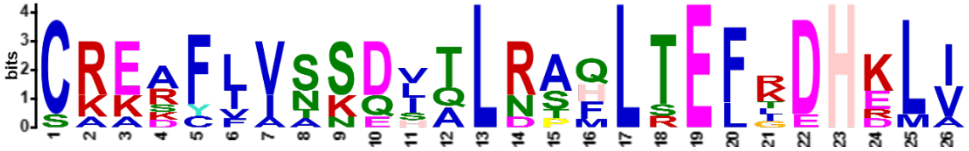

| Name | Start | *p*-value | | Sites | |
|---|---|---|---|---|---|
| 7. Homo | 520 | 2.60e-29 | glsfqdfyqq | CREAFLVNSDLTLRAQLTEFRDHKLI | rtkkgtdgve |
| 8. Danio | 497 | 5.29e-28 | glsfqdfyqr | CRESFLVNSDITLRTQLTEFRDHKLI | rnkkgadgve |
| 6. Drosophila | 562 | 1.03e-27 | gmpfrdlyss | CREAFLVSSDLALRAQLTEFLDHKLV | kskrsvdgse |
| 5. Zea | 323 | 3.67e-27 | gmpvsslytk | CRERFLVSSQVTLNAHLTEFKDHDLV | kvrkhsdgqd |
| 4. Arabidopsis | 308 | 4.32e-22 | gmptddlysa | SRERFFVSSQVTLNSHLTEFKDHELV | ktkrnsdgqe |
| 2. Fistulifera | 499 | 1.86e-19 | WVEFSSFRDQ | CKKKYIITKDHQLDPFLTELGDHRLI | QKKIIKGVNY |
| 1. Phaeodactylum | 531 | 3.45e-19 | WIEYRDFCNA | CKKACVIAKDSQLRAFLSELTDHELI | AMNTTANANR |
| 3. Saccharomyces | 563 | 3.72e-18 | gvelklfnhl | CAADFIASNEIALRSMLREFIEHKMA | nitknnsgme |

**3.**

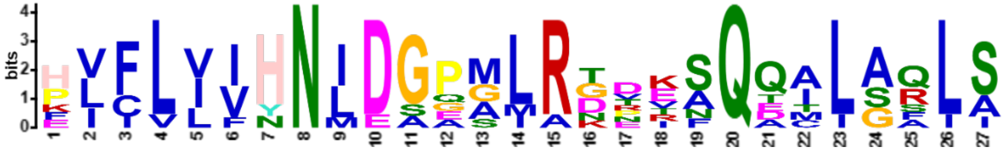

| Name | Start | *p*-value | | Sites | |
|---|---|---|---|---|---|
| 5. Zea | 192 | 1.85e-27 | mrqtsddvdd | HVCLLIHNIDGPALRDTESQQCLAQIS | ccpqirvvas |
| 8. Danio | 364 | 6.00e-27 | trtlrgdpsn | HVFLIINNIDGPMLRGDRNQQALGQLA | alpnmhllas |
| 4. Arabidopsis | 177 | 1.29e-24 | hgpqsgdkdc | FICVVVHNIDGPALRDPESQQTLARLS | scshirlvas |
| 2. Fistulifera | 343 | 2.28e-23 | IAYDAMETLE | PVFLIIHNIEGEGLRTDVSQEALAALS | VNSRVANGTA |
| 6. Drosophila | 429 | 4.82e-23 | eeefalipet | HLFLIVHNLDGAMLRNVKAQAILSRLA | ripnihllas |
| 7. Homo | 387 | 7.71e-22 | vnkfkedssl | ELFLLIHNLDSQMLRGEKSQQIIGQLS | slhniylias |
| 3. Saccharomyces | 419 | 1.71e-19 | dfyknqpldi | KLILVVHNLDGPSIRKNTFQTMLSFLS | virqiaivas |
| 1. Phaeodactylum | 377 | 3.88e-17 | ATIGDNPTYT | PIFLVFYNMDAGGMATRIAQDALASLI | VNSTVANGIQ |

**4.**

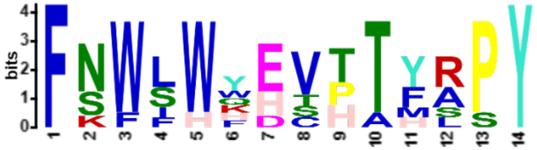

| Name | Start | *p*-value | | Sites | |
|---|---|---|---|---|---|
| 4. Arabidopsis | 232 | 1.14e-17 | lwdkkmvhkq | FNWLWHHVPTFAPY | nvegvffplv |
| 7. Homo | 441 | 1.47e-17 | lmwdhakqsl | FNWLWYETTTYSPY | teetsyensl |
| 5. Zea | 247 | 7.34e-17 | lwdkkmvhkq | FKWSWYHVPTFAPY | kvegvfypli |
| 8. Danio | 418 | 1.93e-15 | lvwdhakmsm | FNWLWFESTTYRSY | teetsyensl |
| 2. Fistulifera | 405 | 2.13e-15 | KLWRLPVMAS | FSWIWKEVHAYRPY | DNEFVMLETD |
| 1. Phaeodactylum | 439 | 2.60e-13 | QMWSSSTAAN | FSWFHQEVHTHRPY | VEELTTLRDQ |
| 6. Drosophila | 483 | 4.25e-13 | llwdqgklcs | FNFSWWDCTTMLPY | tnetafensl |

5.

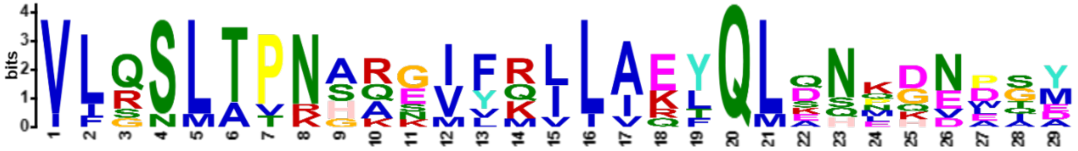

| Name | Start | *p*-value | | Sites | |
|---|---|---|---|---|---|
| 7. Homo | 480 | 1.73e-30 | gslplsslth | VLRSLTPNARGIFRLLIKYQLDNQDNPSY | iglsfqdfyq |
| 8. Danio | 457 | 1.12e-29 | galalsslth | VLRSLTPNARGIFRLLAEFQLENKDNPAY | sglsfqdfyq |
| 5. Zea | 286 | 6.45e-26 | haqttktalv | VLQSLTPNAQSVFRVLAEYQLANEKEEGM | pvsslytkcr |
| 4. Arabidopsis | 271 | 2.50e-25 | taqtaktaai | VLQSLTPNGQNVFKILAEYQLSHPDEDGM | ptddlysasr |
| 6. Drosophila | 522 | 2.04e-22 | gelalssmrs | VFSSLTTNSRGIYMLIVKYQLKNKGNATY | qgmpfrdlys |
| 3. Saccharomyces | 513 | 1.11e-20 | tssgaegaky | VLQSLTVNSKKMYKLLIETQMQNMGNLSA | ntgpkrgtqr |
| 1. Phaeodactylum | 495 | 5.07e-18 | SQASSDRILR | VLQNLAPRHAEVVQILARLQLDSNQDWIE | YRDFCNACKK |
| 2. Fistulifera | 457 | 2.94e-16 | AETKKARVLS | IIGSMAPKHAEILQILAQLQLQQLHVDQD | KEGWVEFSSF |

6.

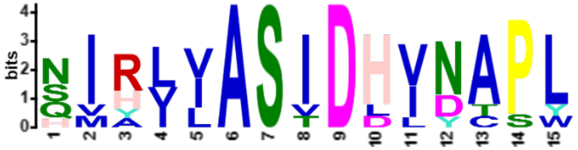

| Name | Start | *p*-value | | Sites | |
|---|---|---|---|---|---|
| 5. Zea | 222 | 1.13e-17 | qclaqisccp | QIRVVASIDHVNAPL | lwdkkmvhkq |
| 4. Arabidopsis | 207 | 1.13e-17 | qtlarlsscs | HIRLVASIDHVNAPL | lwdkkmvhkq |
| 7. Homo | 417 | 1.52e-16 | qiigqlsslh | NIYLIASIDHLNAPL | mwdhakqslf |
| 8. Danio | 394 | 5.27e-16 | qalgqlaalp | NMHLLASIDHINAPL | vwdhakmsmf |
| 6. Drosophila | 459 | 6.33e-16 | ailsrlarip | NIHLLASIDHINTPL | lwdqgklcsf |
| 3. Saccharomyces | 449 | 3.68e-14 | tmlsflsvir | QIAIVASTDHIYAPL | lwdnmkaqny |
| 1. Phaeodactylum | 414 | 7.63e-13 | VNSTVANGIQ | SVRVIASVDLVDAPW | QMWSSSTAAN |
| 2. Fistulifera | 380 | 1.17e-12 | VNSRVANGTA | SIRIIASIDDVDCSY | KLWRLPVMAS |

**7.**

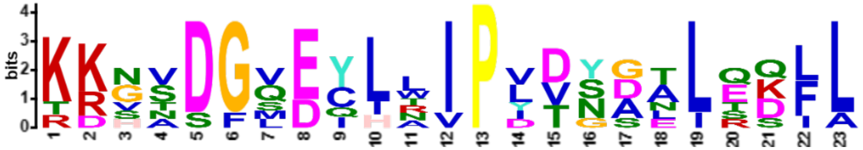

| Name | Start | *p*-value | | Sites | |
|---|---|---|---|---|---|
| 7. Homo | 548 | 8.58e-24 | efrdhklirt | KKGTDGVEYLLIPVDNGTLTDFL | ekeeeea |
| 8. Danio | 525 | 6.22e-23 | efrdhklirn | KKGADGVEYLLIPVDNGTLSDFL | ekndve |
| 4. Arabidopsis | 336 | 2.65e-19 | efkdhelvkt | KRNSDGQECLNIPLTSDAIRQLL | ldlnq |
| 5. Zea | 351 | 2.39e-18 | efkdhdlvkv | RKHSDGQDCLRIPLVSDALEKLL | qela |
| 6. Drosophila | 590 | 3.23e-18 | efldhklvks | KRSVDGSEQLTIPIDGALLQQFL | eeqekk |
| 3. Saccharomyces | 591 | 4.32e-15 | efiehkmani | TKNNSGMEIIWVPYTYAELEKLL | ktvlntl |
| 2. Fistulifera | 66 | 2.36e-13 | APTGRSFKTG | KDVVDFLDYHAIPDVYSNLQSIA | KPEYEASEEQ |

**8.**

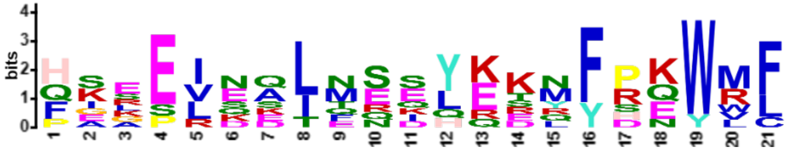

| Name | Start | *p*-value | | Sites | |
|---|---|---|---|---|---|
| 4. Arabidopsis | 57 | 1.72e-17 | retastiemk | HSKEISELMSDYKTMYSKWVF | elrcgfgllm |
| 6. Drosophila | 325 | 1.97e-17 | llseiktsae | HEGSINAIMEEYRSYFPKWMC | ilnegfnill |
| 8. Danio | 260 | 3.86e-16 | lgllgkntpr | FAEEINQLNSKHEKNFRQWML | qlrmgfnill |
| 5. Zea | 71 | 2.95e-15 | raslaqippk | HKEEVESLTRSYKDQYRNWLF | elrcgfgllm |
| 7. Homo | 283 | 1.18e-14 | rnllskvsps | FSAELKQLNQQYEKLFHKWML | qlhlgfnivl |
| 2. Fistulifera | 212 | 3.29e-13 | EKCRQISREH | QGLELDAIQSELERNFDEWRF | LLTTNHNLLF |
| 1. Phaeodactylum | 241 | 1.40e-12 | DECAALISSY | PISEVEDTENSLQERFPEWRF | LLSTNHSLLF |
| 3. Saccharomyces | 286 | 3.35e-12 | lvsnffnenf | QKRPRQKLFEIQKKMFPQYWF | eltqgfsllf |

**9.**

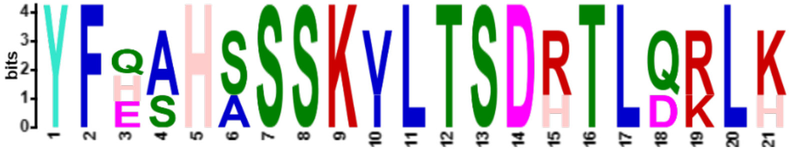


| Name | Start | *p*-value | | Sites | |
|---|---|---|---|---|---|
| 8. Danio | 220 | 3.22e-25 | ktaesdlvee | YFQAHSSSKVLTSDRTLQRLH | tpqldretll |
| 7. Homo | 243 | 1.29e-23 | rdktsdlvee | YFEAHSSSKVLTSDRTLQKLK | rakldqqtlr |
| 6. Drosophila | 283 | 1.32e-21 | snefvpesdg | YFHSHASSKILTSDHTLDRLK | nprlaadrvf |

**10.**

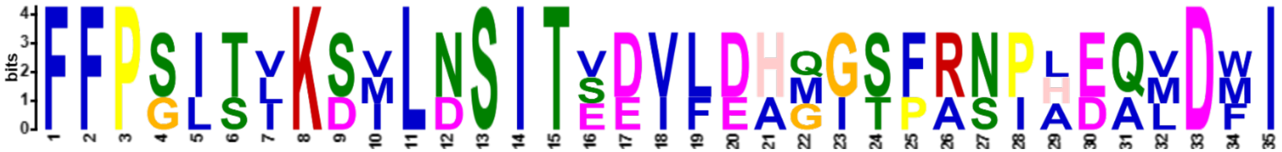


| Name | Start | *p*-value | | Sites | |
|---|---|---|---|---|---|
| 8. Danio | 319 | 1.22e-38 | sdydhvvvng | FFPSITLKSILNSITVDVFEHQGSFRNPAEQMDFI | trtlrgdpsn |
| 7. Homo | 342 | 1.55e-33 | qdsihvving | FFPGISVKSVLNSITEEVLDHMGTFRSILDQLDWI | vnkfkedssl |
| 6. Drosophila | 384 | 3.37e-30 | hkqtvlvvng | FFPSLTIKDMLDSITSDILDAGISPANPHEAVDMI | eeefalipet |

**>ORC4**

| Name | *p*-value |
|---|---|
| Phaeodactylum | 5.62e-100 |
| Fistulifera | 6.47e-85 |
| Saccharomyces | 2.70e-60 |
| Arabidopsis | 2.54e-170 |
| Zea | 1.94e-167 |
| Drosophila | 9.38e-171 |
| Homo | 5.00e-262 |
| Danio | 5.52e-256 |

Motif Locations

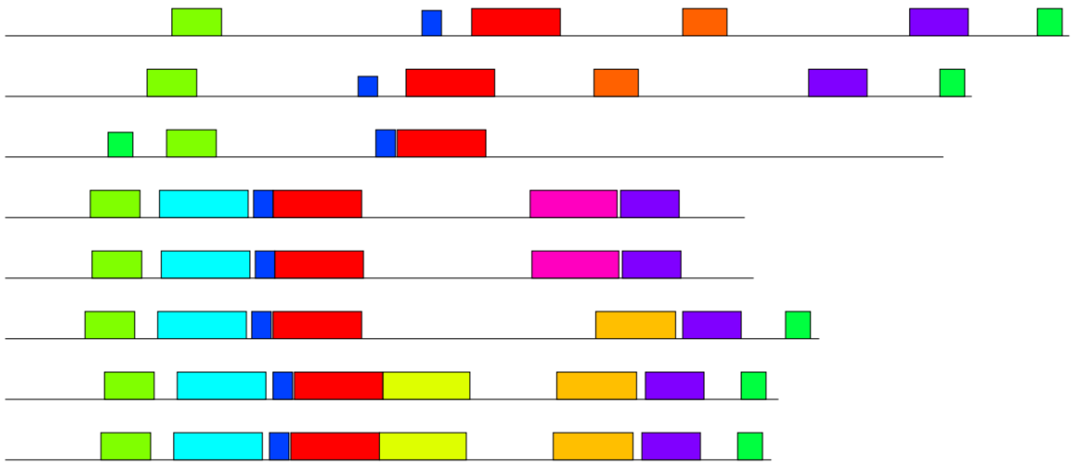

| Motif | Symbol | Motif Consensus |
|---|---|---|
| 1. | | FAAQKNQTLLYNLLDVVQSAQSPVCVVGLTCRLDVLELLEKRVKSRFSHR |
| 2. | | HLNGLLHTDDKIALKEITRQLHLENVVGDKVFGSFAENLAFLJEALKKGG |
| 3. | | EGENNSVLLLGPRGSGKTLLLBQVLRDL |
| 4. | | HSSNKYEKPVVLKAFEHLLQLELIRPAENRGGN |
| 5. | | DSKANILHGLSVLELCLIIAMKHLNDIYDGEPFNFQMVYNEFKKF |
| 6. | | QRYPQCPTDVRQWA |
| 7. | | KPVIFILDEFD |
| 8. | | MQRQPKLEALRDCSILELYJLVCMRRLEVKEQSSYNFIRIMKEYKAIQD |
| 9. | | CRLGRDVRWFSRVLSFAJINYRDDC |
| 10. | | QIHLFNSFGFPQYVDIFKEQLSLPQEFPDKRFAEKWNZNVQKLCEDKSV |

**1.**

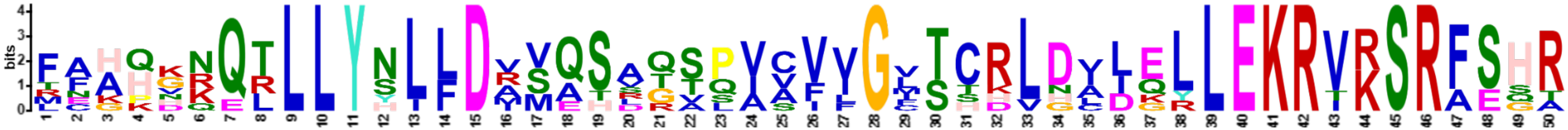

| | Name | Start | *p*-value | Sites | | |
|---|---|---|---|---|---|---|
| 8. | Danio | 162 | 2.21e-55 | vlflldefdl | FAHHKNQTLLYNLLDVSQSAQTPVAVVGFTCRLDVLELLEKRVKSRFSHR | qihlfsslsf |
| 7. | Homo | 164 | 4.49e-54 | vifildefdl | FAHHKNQTLLYNLFDISQSAQTPIAVIGLTCRLDILELLEKRVKSRFSHR | qihlmnsfgf |
| 6. | Drosophila | 152 | 4.05e-52 | vifileefdl | FCAHHNQTLLYNLFDVSQSAQAPICVLGVTCRLDVIELLEKRVKSRFSHR | qvflfpslrr |
| 5. | Zea | 153 | 8.40e-43 | tilfileefd | LFAQGKQRLLYSLLDAMQSLTSQAVVIGVSCRLDADQLLEKRVRSRFSHR | kllfispsld |
| 4. | Arabidopsis | 152 | 1.38e-41 | tiifvldefd | MFAQGKQRLLYSLLDAMQSVTSQAVVVGISSRLDADQLLEKRVRSRFSHR | kflflppsre |
| 3. | Saccharomyces | 222 | 7.08e-39 | vvfifdeidt | FAGPVRQTLLYNLFDMVEHSRVPVCIFGCTTKLNILEYLEKRVKSRFSQR | viympqiqnl |
| 1. | Phaeodactylum | 264 | 2.31e-34 | ATANSLDSGL | TEHKDRQLLLYHLLDRVASDGSSVCFVGMTCHLGLIGLLEKRIRSRAEGT | SQFLQFGPCT |
| 2. | Fistulifera | 227 | 7.19e-27 | GHSLQQPLDQ | RNKQNQELLLYYILDRVATRGSLVSFVGLSHDVHCLKRLEKRTRSRAESA | LFISCESFPS |

**2.**

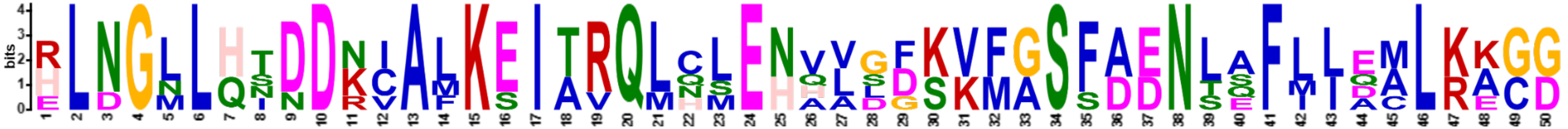

| Name | Start | *p*-value | | Sites | |
|---|---|---|---|---|---|
| 8. Danio | 96 | 3.63e-53 | kevqknvllv | ELNGLLQTDDKIALKEITRQLHLENVVGDKVFGSFAENLAFLLEALKKGD | ksssrpvlfl |
| 7. Homo | 98 | 2.74e-49 | eevsenvlqv | HLNGLLQINDKIALKEITRQLNLENVVGDKVFGSFAENLSFLLEALKKGD | rtsscpvifi |
| 4. Arabidopsis | 88 | 1.19e-43 | eqfpdsvsvi | RLNGLLHSDDNCAFKEIARQLCMEHHLLFSKMASFDDNSQFIIAMLRACG | lahktiifvl |
| 6. Drosophila | 87 | 2.37e-43 | ksfgentliv | HLDGNLHTDDRVALKSITVQMQLENAADGKVFGSFAENLAFLLQCLKAGG | khsksvifil |
| 5. Zea | 89 | 1.27e-41 | eehpdaisvi | RLNGMLHNDDNCAMKEIARQLCSEHQLSFSKMASSDDNTEFMIDMLRECG | lahktilfil |

**3.**

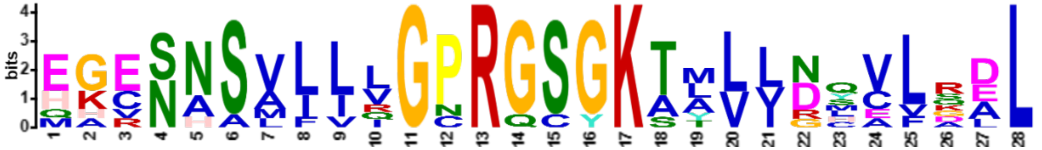

| Name | Start | *p*-value | | Sites | |
|---|---|---|---|---|---|
| 8. Danio | 55 | 6.75e-29 | llellrrtav | HGESNSVLIVGPRGSGKTMLLGCVLREL | mslkevqknv |
| 7. Homo | 57 | 2.57e-28 | lsellkrtal | HGESNSVLIIGPRGSGKTMLINHALKEL | meieevsenv |
| 5. Zea | 50 | 6.78e-26 | lkylvassvs | EACNNSVLLLGPRGCGKAAVVDMVLDDL | keehpdaisv |
| 6. Drosophila | 46 | 9.42e-26 | vrlllqrtae | MGESNSLLLLGPRGSGKTTLINSVLADL | lpnksfgent |
| 4. Arabidopsis | 49 | 1.30e-25 | lkfivstsit | EGCNNSMLLLGPRGSGKAAVLDLVVGDL | leqfpdsvsv |
| 1. Phaeodactylum | 95 | 6.41e-21 | MFRPCLVQRD | EHVNASAIVQGCRGSGKTLLVNQCFQAL | RDELDAQSHA |
| 3. Saccharomyces | 92 | 1.57e-19 | idriikqsii | QKESHSVILVGPRQSYKTYLLDYELSLL | qqsykeqfit |
| 2. Fistulifera | 81 | 6.67e-19 | YRLLRTHLLH | EKRNAAAFLRGNRGSGKSLVLRQCLRAL | QDESEHIQGS |

**4.**

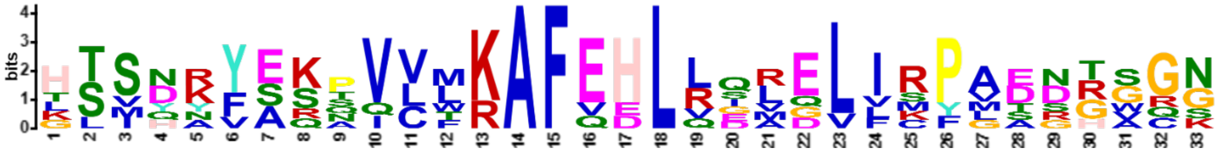

| Name | Start | *p*-value | | Sites | |
|---|---|---|---|---|---|
| 7. Homo | 362 | 5.19e-30 | efqkfvqrka | HSVYNFEKPVVMKAFEHLQQLELIKPMERTSGN | sqreyqlmkl |
| 8. Danio | 360 | 5.15e-28 | efkkfiqrks | HSIHKFEKPVVMKAFEHLLQLELVRPVDSGVCK | vqreyqlmrl |
| 5. Zea | 349 | 3.07e-25 | keyrsiqday | KTSDKYASTVCFRAFEHLLDRELISFGDNRWRN | qaleyrpvkl |
| 4. Arabidopsis | 348 | 9.85e-25 | keykaihdsf | HTSDYYAQNVCLRAFEHLRERQVICYAENRGQS | qtgeyrlqkl |
| 6. Drosophila | 383 | 1.21e-23 | arfskfakvs | TTMQAVERSIVLKAFEHLRIAELIMPLTGGAGG | gvgkvqkefe |
| 1. Phaeodactylum | 511 | 8.82e-23 | MLEEYLGSYK | GSSNRYSKQVLTKAFVDLLSVGLLRPASDHSGG | APLQYQHDRS |
| 2. Fistulifera | 454 | 3.82e-21 | EYNTAIQKGS | LLSNRYSSAQLWKAFQELVGMDLFRPAADTGGN | LPFQYMHLDS |

**5.**

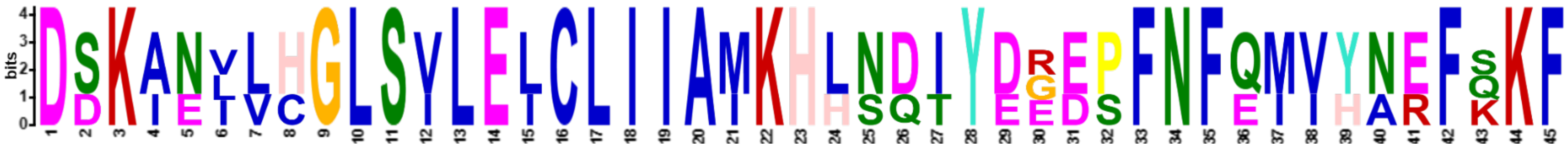

| Name | Start | *p*-value | | Sites | |
|---|---|---|---|---|---|
| 7. Homo | 312 | 3.35e-54 | lmeasqlcsm | DSKANIVHGLSVLEICLIIAMKHLNDIYEEEPFNFQMVYNEFQKF | vqrkahsvyn |
| 8. Danio | 310 | 5.37e-54 | flesgrlisa | DSKANVLHGLSILELCLIIAMKHLNDTYDGEPFNFQMVHNEFKKF | iqrkshsihk |
| 6. Drosophila | 334 | 8.00e-39 | maavgsqfeg | DDKIELLCGLSVLELCLIIAIKHHSQIYDRDSFNFEIIYARFSKF | akvsttmqav |

**6.**

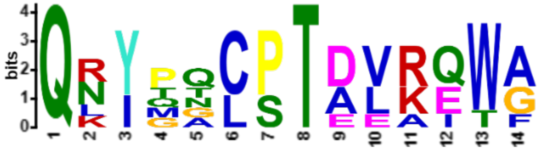

| Name | Start | *p*-value | Sites | | |
|---|---|---|---|---|---|
| 8. Danio | 414 | 1.27e-19 | lehgqvmeal | QRYPQCPTDVKQWA | lsafa |
| 7. Homo | 416 | 1.82e-19 | ldntqimnal | QKYPNCPTDVRQWA | tsslswl |
| 6. Drosophila | 441 | 3.27e-14 | ltysqihhcm | QRYQALPTEVAQWA | qssli |
| 2. Fistulifera | 528 | 1.63e-13 | VHREVKVALE | QNITQCSTALREWG | LKTN |
| 1. Phaeodactylum | 583 | 1.97e-13 | VYRELEGAIA | QNIMGCSTALREWG | RKMN |
| 3. Saccharomyces | 59 | 8.86e-10 | fgslqrrllq | QLYGTLPTDEKITF | tylqdcqqei |

**7.**

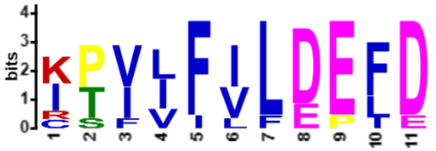

| Name | Start | *p*-value | Sites | | |
|---|---|---|---|---|---|
| 7. Homo | 152 | 5.18e-14 | alkkgdrtss | CPVIFILDEFD | lfahhknqtl |
| 4. Arabidopsis | 141 | 7.87e-13 | amlracglah | KTIIFVLDEFD | mfaqgkqrll |
| 5. Zea | 142 | 2.28e-11 | dmlrecglah | KTILFILEEFD | lfaqgkqrll |
| 6. Drosophila | 140 | 3.51e-11 | qclkaggkhs | KSVIFILEEFD | lfcahhnqtl |
| 8. Danio | 150 | 3.83e-11 | alkkgdksss | RPVLFLLDEFD | lfahhknqtl |
| 3. Saccharomyces | 210 | 5.27e-11 | gevdresitk | ITVVFIFDEID | tfagpvrqtl |
| 1. Phaeodactylum | 236 | 5.00e-10 | EVLQIAKDDN | IPIVIVLDELD | LFLGQQKATA |
| 2. Fistulifera | 200 | 4.42e-8 | EALEIARGDQ | IPFLFVLDPLE | AFIATGGHSL |

8.

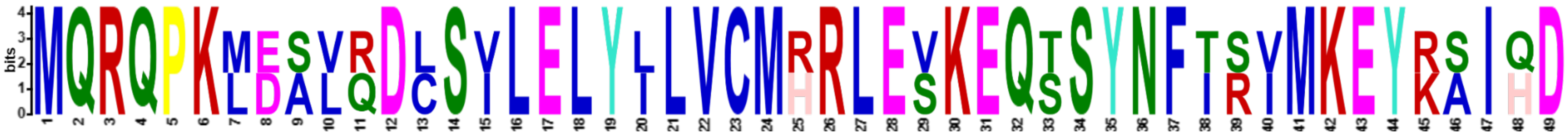

| Name | Start | *p*-value | Sites | | |
|---|---|---|---|---|---|
| 4. Arabidopsis | 297 | 4.93e-57 | lenfkaalss | MQRQPKLEAVRDCSVLELYLLVCMRRLEVKEQSSYNFISVMKEYKAIHD | sfhtsdyyaq |
| 5. Zea | 298 | 3.63e-55 | vesflkalss | MQRQPKMDSLQDLSILELYILVCMHRLESKEQTSYNFTRIMKEYRSIQD | ayktsdkyas |

9.

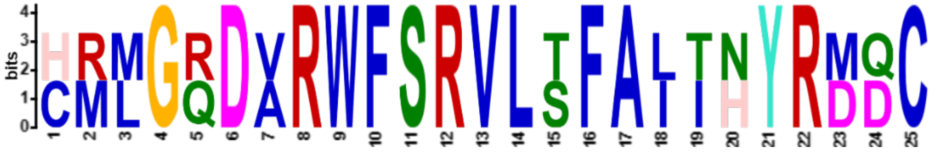

| Name | Start | *p*-value | Sites | | |
|---|---|---|---|---|---|
| 1. Phaeodactylum | 383 | 1.67e-31 | RRVYNAFVRD | CMLGQDVRWFSRVLTFAITNYRMDC | LAAMEGDASS |
| 2. Fistulifera | 333 | 1.19e-29 | ESVVKCLQRE | HRMGRDARWFSRVLSFALIHYRDQC | SHGLNSDDAD |

10.

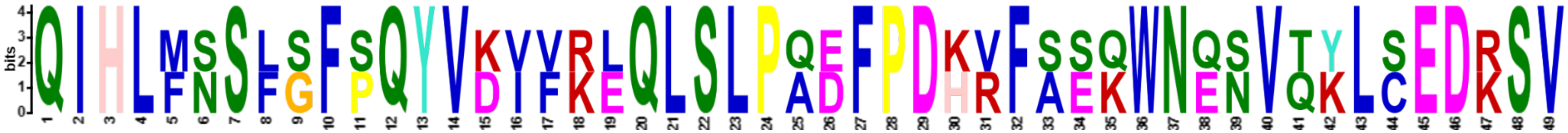

| Name | Start | *p*-value | Sites | | |
|---|---|---|---|---|---|
| 7. Homo | 214 | 2.30e-57 | krvksrfshr | QIHLMNSFGFPQYVKIFKEQLSLPAEFPDKVFAEKWNENVQYLSEDRSV | qevlqkhfni |
| 8. Danio | 212 | 8.23e-50 | krvksrfshr | QIHLFSSLSFSQYVDVVRLQLSLPQDFPDHRFSSQWNQSVTKLCEDKSV | levlkrsfns |

**Fig. S1: Arrangement and distribution of motifs predicted from the ORC2 and ORC4 of eight eukaryotic organisms.**

The locations of motifs predicted using MEME Suite are displayed on the ORC2 and ORC4 sequences of each eukaryotic organism. The sequence information of predicted motifs was aligned and organised.

>ORC2

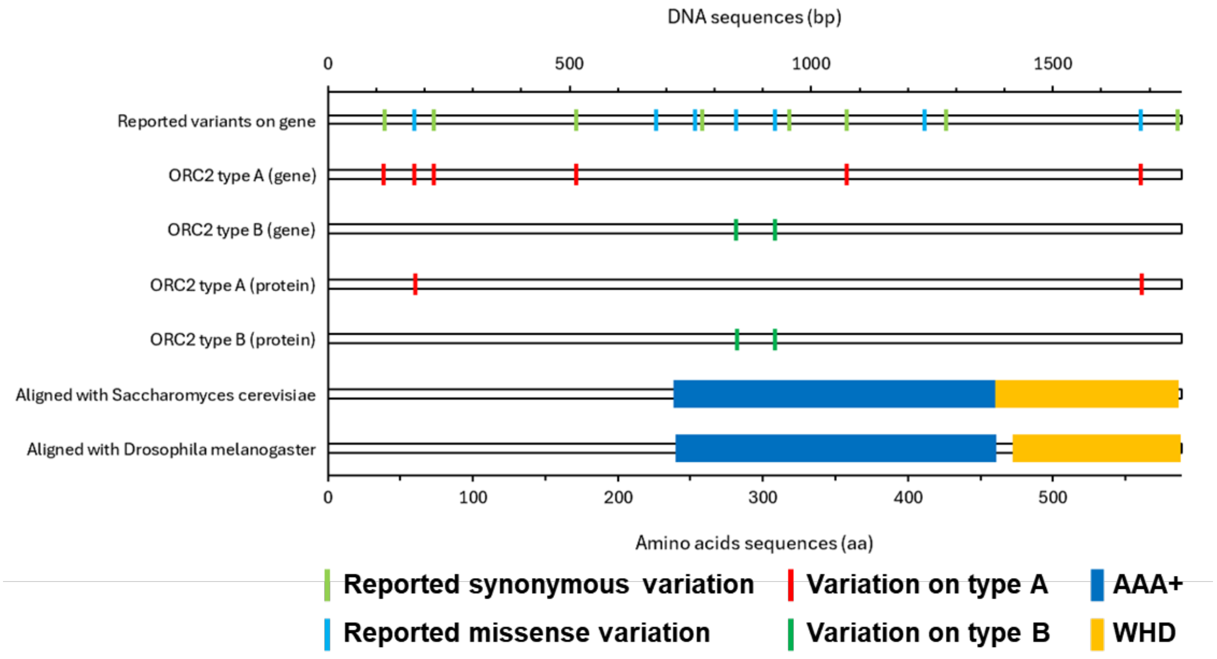
DNA sequences (bp)
0
500
1000
1500
Reported variants on gene
ORC2 type A (gene)
ORC2 type B (gene)
ORC2 type A (protein)
ORC2 type B (protein)
Aligned with Saccharomyces cerevisiae
Aligned with Drosophila melanogaster
0
100
200
300
400
500
Amino acids sequences (aa)
Reported synonymous variation
Reported missense variation
Variation on type A
Variation on type B
AAA+
WHD

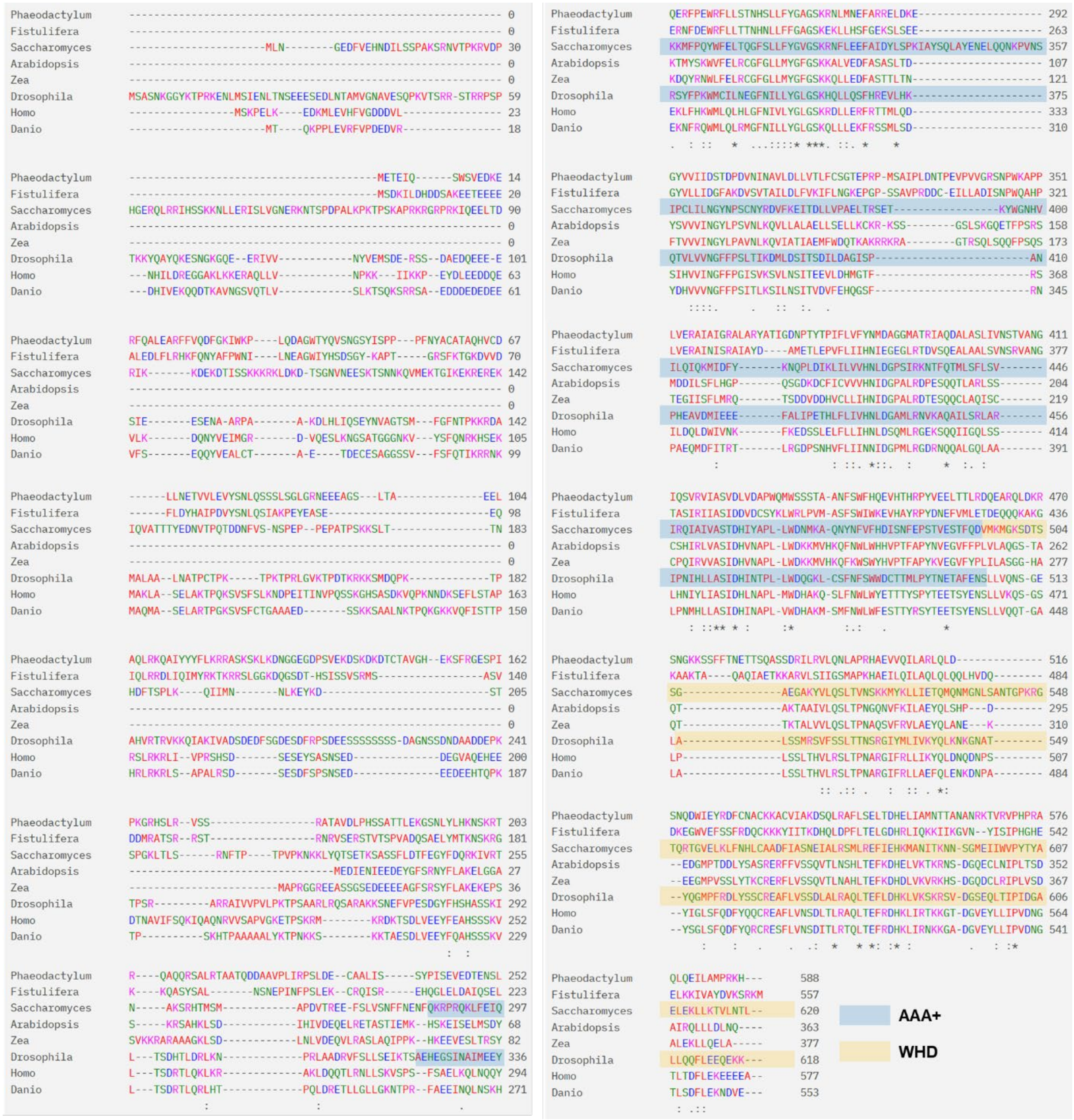
Phaeodactylum
Fistulifera
Saccharomyces
Arabidopsis
Zea
Drosophila
Homo
Danio
AAA+
WHD

**>ORC4**

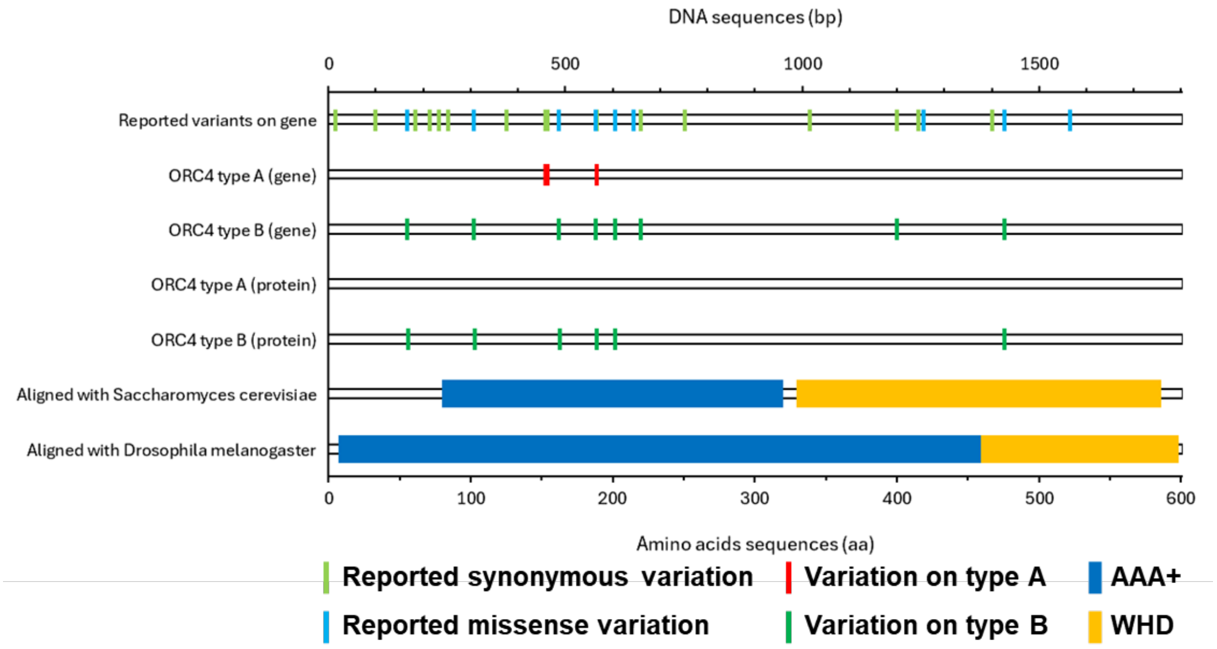

DNA sequences (bp)
0
500
1000
1500
Reported variants on gene
ORC4 type A (gene)
ORC4 type B (gene)
ORC4 type A (protein)
ORC4 type B (protein)
Aligned with Saccharomyces cerevisiae
Aligned with Drosophila melanogaster
0
100
200
300
400
500
600
Amino acids sequences (aa)
Reported synonymous variation
Variation on type A
AAA+
Reported missense variation
Variation on type B
WHD


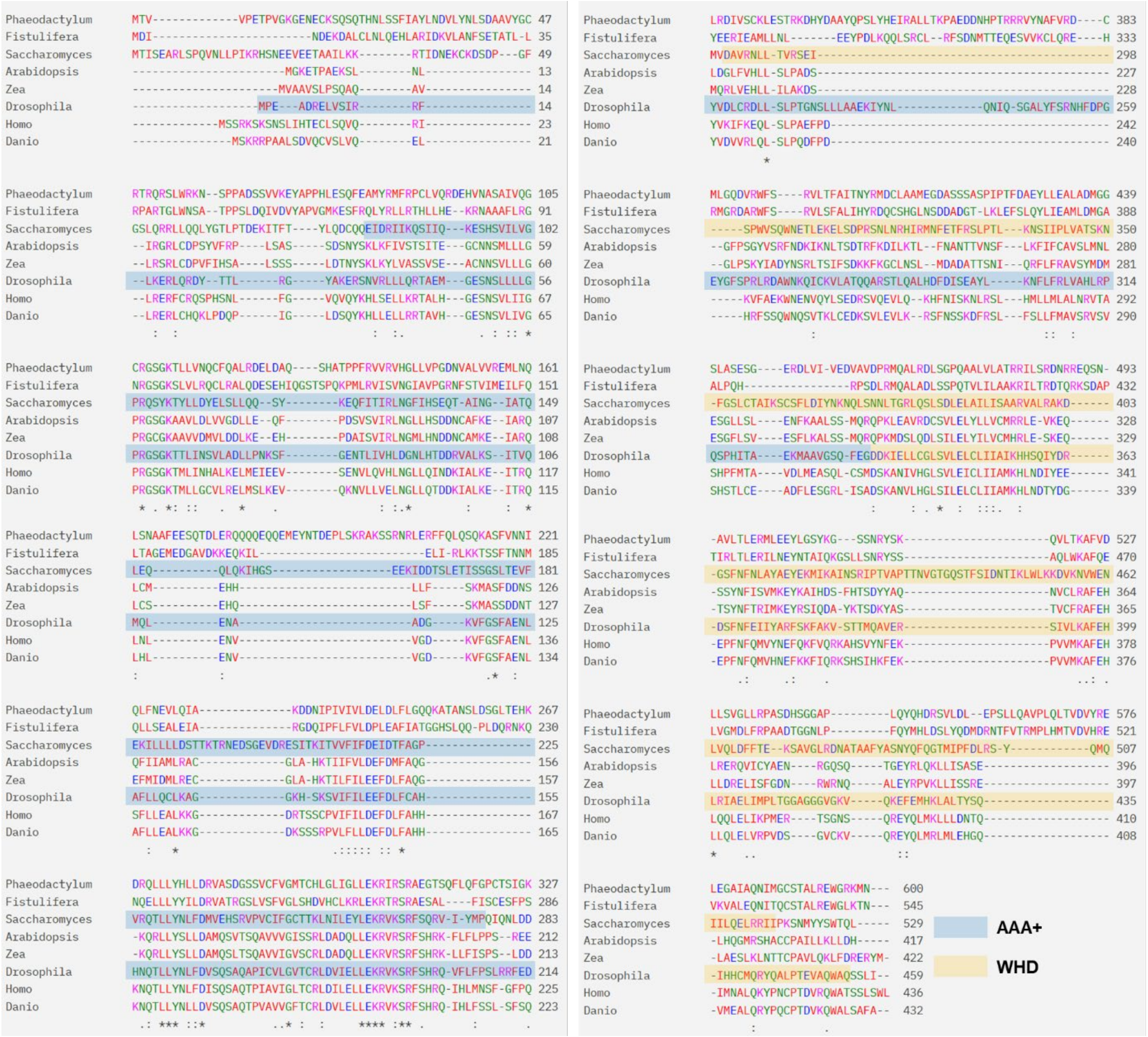

Phaeodactylum
Fistulifera
Saccharomyces
Arabidopsis
Zea
Drosophila
Homo
Danio
AAA+
WHD

**Fig. S2: Multiple sequence alignment result of the ORC2 and ORC4 sequences from eight eukaryotic organisms.**

ORC2 and ORC4 sequences from eight eukaryotic organisms were aligned and compared via multiple sequence alignment (Clustal Omega; https://www.ebi.ac.uk/). The positions of the AAA+ (ATPases associated with diverse cellular activities) domain and WHD (winged helix domain) are indicated by blue and orange lines, respectively, on the sequences of *Saccharomyces cerevisiae* (Li *et al.,* 2018) and *Drosophila melanogaster* (Bleichert *et al.,* 2015).

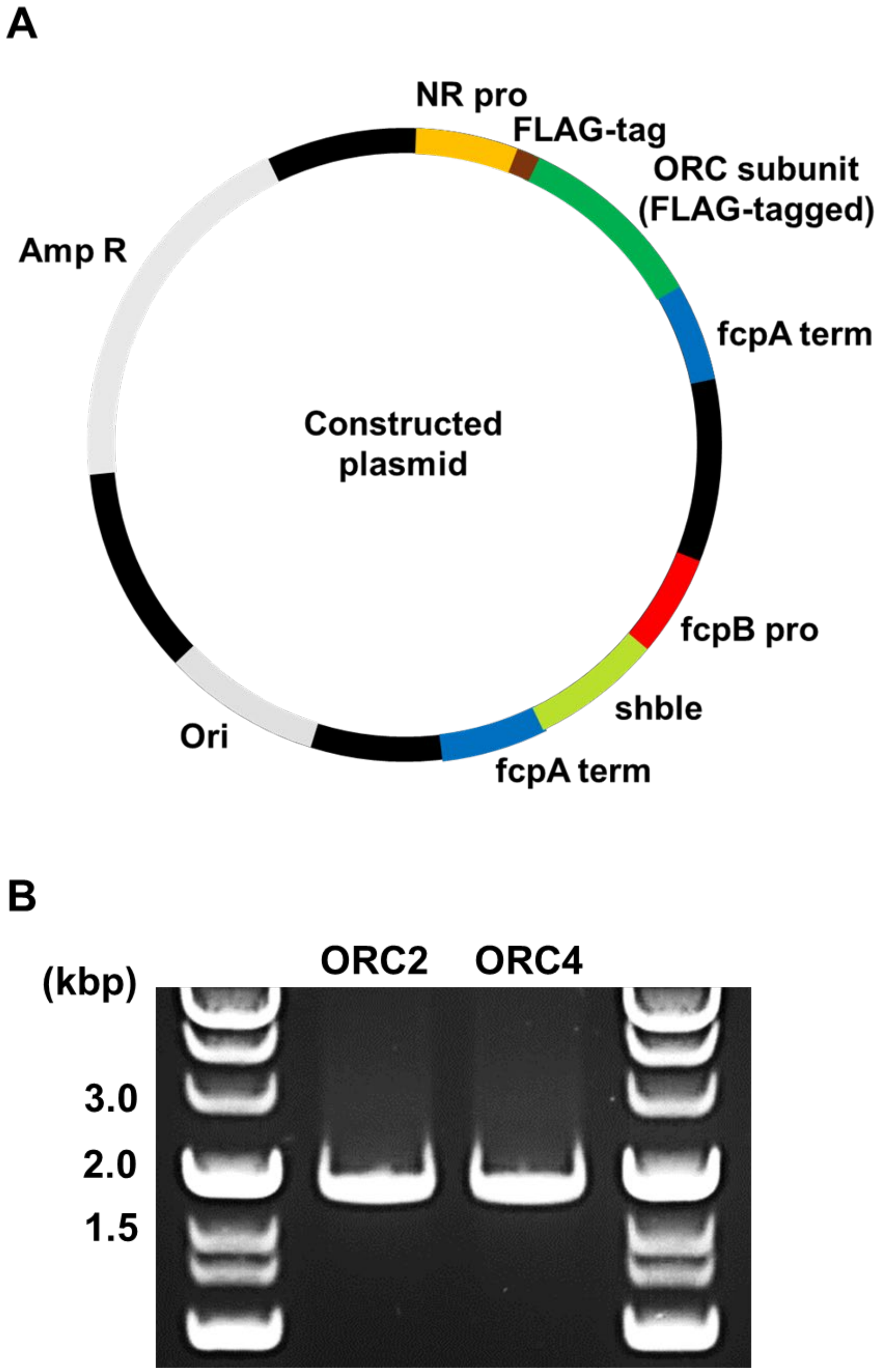


**Fig. S3: Design of plasmids used in this study to express FLAG-tag-fused ORC subunits in transformants and DNA fragments of native ORC2 and ORC4 amplified from *Phaeodactylum tricornutum*.**

(A) Using pPha-T1 as a backbone plasmid, plasmids were designed to express FLAG-tagged ORC subunits (ORC2 and ORC4) driven by the nitrate reductase promoter (NR pro). (B) DNA fragments of ORC2 and ORC4 amplified with primers designed to locate the FLAG-tag at the N-terminus had a size of less than 2 kbp (ORC2: 1824 bp, ORC4: 1860 bp).

>ORC2 (Phatr3_J42935; Chr1: 1565023-1566789)

```
atggagacagagattcagtcctggagcgttgaagataaggaacgtttccaagcacttgag
 M  E  T  E  I  Q  S  W  S  V  E  D  K  E  R  F  Q  A  L  E
gcccggttctttgtacaagatttggaaagatatggaaaccgttgcaagatgcaggttgg
 A  R  F  F  V  Q  D  F  G  K  I  W  K  P  L  Q  D  A  G  W
acctaccaagtctccaatggatcctacatatccccgccaccattcaactatgcctgtgca
 T  Y  Q  V  S  N  G  S  Y  I  S  P  P  P  F  N  Y  A  C  A
acggcgcagcatgtctgtgatctcttaaacgagacagtggtattggaagtctacagtaat
 T  A  Q  H  V  C  D  L  L  N  E  T  V  V  L  E  V  Y  S  N
ttgcaaagtagtagtttatctggacttggcaggaacgaggaagaagctggcagtctaact
 L  Q  S  S  S  L  S  G  L  G  R  N  E  E  E  A  G  S  L  T
gcggaggagcttgctcaacttcgcaaacaagcaatttactactacttcctaaagcgtaga
 A  E  E  L  A  Q  L  R  K  Q  A  I  Y  Y  Y  F  L  K  R  R
gcttcaaaatcaaagcttaaagacaatggaggagaaggagacccttccgtcgaaaaagac
 A  S  K  S  K  L  K  D  N  G  G  E  G  D  P  S  V  E  K  D
tccaaagacaaagacacgtgtaccgccgttggtcatgagaagagttttcgcggggaatcc
 S  K  D  K  D  T  C  T  A  V  G  H  E  K  S  F  R  G  E  S
ccgatcccaaaaggacgacattcactccgagtatcaagtcgcgcaacagccgtagacctc
 P  I  P  K  G  R  H  S  L  R  V  S  S  R  A  T  A  V  D  L
cctcattcatccgcgacaacactggaaaagggatccaatctctatttgcacaagaattcc
 P  H  S  S  A  T  T  L  E  K  G  S  N  L  Y  L  H  K  N  S
aaaagaactcgccaggcacagcaacggtctgcgttacgaaccgcagcaactcaggatgac
 K  R  T  R  Q  A  Q  Q  R  S  A  L  R  T  A  A  T  Q  D  D
gctgcagtacctttgattcgacctagtctcgatgaatgtgcggcgttgatatcttcctac
 A  A  V  P  L  I  R  P  S  L  D  E  C  A  A  L  I  S  S  Y
cctatatctgaagtagaggatacggagaattctcttcaggaaaggtttccggaatggcgg
 P  I  S  E  V  E  D  T  E  N  S  L  Q  E  R  F  P  E  W  R
tttttactgtctaccaaccattccctttttgttctacggggcagggtcgaaacggaatctc
 F  L  L  S  T  N  H  S  L  L  F  Y  G  A  G  S  K  R  N  L
atgaatgaattcgcgcggcgggagctggacaaggaagggtacgtggtgattatcgacagc
 M  N  E  F  A  R  R  E  L  D  K  E  G  Y  V  V  I  I  D  S
accgacccggacgtcaacattaatgccgtgttggatttgttggtgactcttttctgcagc
 T  D  P  D  V  N  I  N  A  V  L  D  L  L  V  T  L  F  C  S
ggcactgaaccacggcccatgtccgctattcctttggacaatacaccagaagtgccagtc
 G  T  E  P  R  P  M  S  A  I  P  L  D  N  T  P  E  V  P  V
gtgggacggtccaatccttggaaggccccaccgttggtggaacgggccattgcgattggg
 V  G  R  S  N  P  W  K  A  P  P  L  V  E  R  A  I  A  I  G
cgagcattggccagatatgctacgattggcgataatcccacgtatacgcctatctttctc
 R  A  L  A  R  Y  A  T  I  G  D  N  P  T  Y  T  P  I  F  L
gtgttttacaatatggatgcgggaggcatggcaactcgcatagcgcaagacgctttggct
 V  F  Y  N  M  D  A  G  G  M  A  T  R  I  A  Q  D  A  L  A
tcattgatcgtcaacagcaccgtcgcgaatgggattcagtccgtccgagtgatcgccagc
 S  L  I  V  N  S  T  V  A  N  G  I  Q  S  V  R  V  I  A  S
gttgacctggtcgatgcaccatggcaaatgtggtcgtcatcgaccgccgccaatttttca
 V  D  L  V  D  A  P  W  Q  M  W  S  S  S  T  A  A  N  F  S
tggtttcatcaggaagtccacacccaccgtccgtatgtagaggaactgactacacttcga
 W  F  H  Q  E  V  H  T  H  R  P  Y  V  E  E  L  T  T  L  R
gaccaagaggcacggcaattagataaacgttcaaacggtaagaaaagtagcttttttcaca
 D  Q  E  A  R  Q  L  D  K  R  S  N  G  K  K  S  S  F  F  T
aacgagacgacatcacaagcgtcgagcgatcgaatcttgcgggtattacaaaatctggcc
 N  E  T  T  S  Q  A  S  S  D  R  I  L  R  V  L  Q  N  L  A
ccccgacatgccgaagtggtacaaatacttgctagacttcaactcgacagcaaccaggat
 P  R  H  A  E  V  V  Q  I  L  A  R  L  Q  L  D  S  N  Q  D
tggatcgaatatcgtgatttttgcaacgcttgcaaaaaggcgtgtgtgattgccaaagac
 W  I  E  Y  R  D  F  C  N  A  C  K  K  A  C  V  I  A  K  D
tctcagttgcgcgcctttttatcggagttaacggaccatgagttgatcgcaatgaacacg
 S  Q  L  R  A  F  L  S  E  L  T  D  H  E  L  I  A  M  N  T
actgcaaatgcgaacaggaaaacggtgcgggtgccgcatccacgagcgcaactacaagaa
 T  A  N  A  N  R  K  T  V  R  V  P  H  P  R  A  Q  L  Q  E
atcttggctatgccacgtaaacattga
 I  L  A  M  P  R  K  H  -
```

>ORC2 type A

```
atggagacagagattcagtcctggagcgttgaagataaggaacgtttccaagcacttgag
 M  E  T  E  I  Q  S  W  S  V  E  D  K  E  R  F  Q  A  L  E
gcccggttctttgtacaagattttggaaagatatggaaaccgttgcaagatgcaggatgg
 A  R  F  F  V  Q  D  F  G  K  I  W  K  P  L  Q  D  A  G  W
acctaccaagtctccaatggatcctacatatccccgccaccattcaactatgcctgtgaa
 T  Y  Q  V  S  N  G  S  Y  I  S  P  P  P  F  N  Y  A  C  E
acggcgcagcatgtctgtgatctcttaaacgagacagtagtattggaagtctacagtaat
 T  A  Q  H  V  C  D  L  L  N  E  T  V  V  L  E  V  Y  S  N
ttgcaaagtagtagtttatctggacttggcaggaacgaggaagaagctggcagtctaact
 L  Q  S  S  S  L  S  G  L  G  R  N  E  E  E  A  G  S  L  T
gcggaggagcttgctcaacttcgcaaacaagcaatttactactacttcctaaagcgtaga
 A  E  E  L  A  Q  L  R  K  Q  A  I  Y  Y  Y  F  L  K  R  R
gcttcaaaatcaaagcttaaagacaatggaggagaaggagacccttccgtcgaaaaagac
 A  S  K  S  K  L  K  D  N  G  G  E  G  D  P  S  V  E  K  D
tccaaagacaaagacacgtgtaccgccgttggtcatgagaagagttttcgcggggaatcc
 S  K  D  K  D  T  C  T  A  V  G  H  E  K  S  F  R  G  E  S
ccgatcccaaaaggacgacattcactccgagtttcaagtcgcgcaacagccgtagacctc
 P  I  P  K  G  R  H  S  L  R  V  S  S  R  A  T  A  V  D  L
cctcattcatccgcgacaacactggaaaagggatccaatctctatttgcacaagaattcc
 P  H  S  S  A  T  T  L  E  K  G  S  N  L  Y  L  H  K  N  S
aaaagaactcgccaggcacagcaacggtctgcgttacgaaccgcagcaactcaggatgac
 K  R  T  R  Q  A  Q  Q  R  S  A  L  R  T  A  A  T  Q  D  D
gctgcagtacctttgattcgacctagtctcgatgaatgtgcggcgttgatatcttcctac
 A  A  V  P  L  I  R  P  S  L  D  E  C  A  A  L  I  S  S  Y
cctatatctgaagtagaggatacggagaattctcttcaggaaaggtttccggaatggcgg
 P  I  S  E  V  E  D  T  E  N  S  L  Q  E  R  F  P  E  W  R
tttttactgtctaccaaccattccctttttgttctacggggcagggtcgaaacggaatctc
 F  L  L  S  T  N  H  S  L  L  F  Y  G  A  G  S  K  R  N  L
atgaatgaattcgcgcggcgggagctggacaaggaagggtacgtggtgattatcgacagc
 M  N  E  F  A  R  R  E  L  D  K  E  G  Y  V  V  I  I  D  S
accgacccggacgtcaacattaatgccgtgttggatttgttggtgactcttttctgcagc
 T  D  P  D  V  N  I  N  A  V  L  D  L  L  V  T  L  F  C  S
ggcactgaaccacggcccatgtccgctattcctttggacaatacaccagaagtgccagtc
 G  T  E  P  R  P  M  S  A  I  P  L  D  N  T  P  E  V  P  V
gtgggacggtccaatccttggaaggccccaccgttggtggaacgggccattgcaattggg
 V  G  R  S  N  P  W  K  A  P  P  L  V  E  R  A  I  A  I  G
cgagcattggccagatatgctacgattggcgataatcccacgtatacgcctatctttctc
 R  A  L  A  R  Y  A  T  I  G  D  N  P  T  Y  T  P  I  F  L
gtgttttacaatatggatgcgggaggcatggcaactcgcatagcgcaagacgctttggct
 V  F  Y  N  M  D  A  G  G  M  A  T  R  I  A  Q  D  A  L  A
tcattgatcgtcaacagcaccgtcgcgaatgggattcagtccgtccgagtgatcgccagc
 S  L  I  V  N  S  T  V  A  N  G  I  Q  S  V  R  V  I  A  S
gttgacctggtcgatgcaccatggcaaatgtggtcgtcatcgaccgccgccaatttttca
 V  D  L  V  D  A  P  W  Q  M  W  S  S  S  T  A  A  N  F  S
tggtttcatcaggaagtccacacccaccgtccgtatgtagaggaactgactacacttcga
 W  F  H  Q  E  V  H  T  H  R  P  Y  V  E  E  L  T  T  L  R
gaccaagaggcacggcaattagataaacgttcaaacggtaagaaaagtagcttttttcaca
 D  Q  E  A  R  Q  L  D  K  R  S  N  G  K  K  S  S  F  F  T
aacgagacgacatcacaagcgtcgagcgatcgaatcttgcgggtattacaaaatctggcc
 N  E  T  T  S  Q  A  S  S  D  R  I  L  R  V  L  Q  N  L  A
ccccgacatgccgaagtggtacaaatacttgctagacttcaactcgacagcaaccaggat
 P  R  H  A  E  V  V  Q  I  L  A  R  L  Q  L  D  S  N  Q  D
tggatcgaatatcgtgatttttgcaacgcttgcaaaaaggcgtgtgtgattgccaaagac
 W  I  E  Y  R  D  F  C  N  A  C  K  K  A  C  V  I  A  K  D
tctcagttgcgcgcctttttatcggagttaacggaccatgagttgatcgcaatgaacacg
 S  Q  L  R  A  F  L  S  E  L  T  D  H  E  L  I  A  M  N  T
agtgcaaatgcgaacaggaaaacggtgcgggtgccgcatccacgagcgcaactacaagaa
 S  A  N  A  N  R  K  T  V  R  V  P  H  P  R  A  Q  L  Q  E
atcttggctatgccacgtaaacattga
 I  L  A  M  P  R  K  H  -
```

>ORC2 type B

```
atggagacagagattcagtcctggagcgttgaagataaggaacgtttccaagcacttgag
 M  E  T  E  I  Q  S  W  S  V  E  D  K  E  R  F  Q  A  L  E 
gcccggttctttgtacaagattttggaaagatatggaaaccgttgcaagatgcaggttgg
 A  R  F  F  V  Q  D  F  G  K  I  W  K  P  L  Q  D  A  G  W 
acctaccaagtctccaatggatcctacatatccccgccaccattcaactatgcctgtgca
 T  Y  Q  V  S  N  G  S  Y  I  S  P  P  P  F  N  Y  A  C  A 
acggcgcagcatgtctgtgatctcttaaacgagacagtggtattggaagtctacagtaat
 T  A  Q  H  V  C  D  L  L  N  E  T  V  V  L  E  V  Y  S  N 
ttgcaaagtagtagtttatctggacttggcaggaacgaggaagaagctggcagtctaact
 L  Q  S  S  S  L  S  G  L  G  R  N  E  E  E  A  G  S  L  T 
gcggaggagcttgctcaacttcgcaaacaagcaatttactactacttcctaaagcgtaga
 A  E  E  L  A  Q  L  R  K  Q  A  I  Y  Y  Y  F  L  K  R  R 
gcttcaaaatcaaagcttaaagacaatggaggagaaggagacccttccgtcgaaaaagac
 A  S  K  S  K  L  K  D  N  G  G  E  G  D  P  S  V  E  K  D 
tccaaagacaaagacacgtgtaccgccgttggtcatgagaagagttttcgcggggaatcc
 S  K  D  K  D  T  C  T  A  V  G  H  E  K  S  F  R  G  E  S 
ccgatcccaaaaggacgacattcactccgagtatcaagtcgcgcaacagccgtagacctc
 P  I  P  K  G  R  H  S  L  R  V  S  S  R  A  T  A  V  D  L 
cctcattcatccgcgacaacactggaaaagggatccaatctctatttgcacaagaattcc
 P  H  S  S  A  T  T  L  E  K  G  S  N  L  Y  L  H  K  N  S 
aaaagaactcgccaggcacagcaacggtctgcgttacgaaccgcagcaactcaggatgac
 K  R  T  R  Q  A  Q  Q  R  S  A  L  R  T  A  A  T  Q  D  D 
gctgcagtacctttgattcgacctagtctcgatgaatgtgcggcgttgatatcttcctac
 A  A  V  P  L  I  R  P  S  L  D  E  C  A  A  L  I  S  S  Y 
cctatatctgaagtagaggatacggagaattctcttcaggaaaggtttccggaatggcgg
 P  I  S  E  V  E  D  T  E  N  S  L  Q  E  R  F  P  E  W  R 
tttttactgtctaccaaccattccctttttgttctacggggcagggtcgaaacggaatctc
 F  L  L  S  T  N  H  S  L  L  F  Y  G  A  G  S  K  R  N  L 
atggatgaattcgcgcggcgggagctggacaaggaagggtacgtggtgattatcgacagc
 M  D  E  F  A  R  R  E  L  D  K  E  G  Y  V  V  I  I  D  S 
accgacccggacgtcaacattaaggccgtgttggatttgttggtgactcttttctgcagc
 T  D  P  D  V  N  I  K  A  V  L  D  L  L  V  T  L  F  C  S 
ggcactgaaccacggcccatgtccgctattcctttggacaatacaccagaagtgccagtc
 G  T  E  P  R  P  M  S  A  I  P  L  D  N  T  P  E  V  P  V 
gtgggacggtccaatccttggaaggccccaccgttggtggaacgggccattgcgattggg
 V  G  R  S  N  P  W  K  A  P  P  L  V  E  R  A  I  A  I  G 
cgagcattggccagatatgctacgattggcgataatcccacgtatacgcctatctttctc
 R  A  L  A  R  Y  A  T  I  G  D  N  P  T  Y  T  P  I  F  L 
gtgttttacaatatggatgcgggaggcatggcaactcgcatagcgcaagacgctttggct
 V  F  Y  N  M  D  A  G  G  M  A  T  R  I  A  Q  D  A  L  A 
tcattgatcgtcaacagcaccgtcgcgaatgggattcagtccgtccgagtgatcgccagc
 S  L  I  V  N  S  T  V  A  N  G  I  Q  S  V  R  V  I  A  S 
gttgacctggtcgatgcaccatggcaaatgtggtcgtcatcgaccgccgccaatttttca
 V  D  L  V  D  A  P  W  Q  M  W  S  S  S  T  A  A  N  F  S 
tggtttcatcaggaagtccacacccaccgtccgtatgtagaggaactgactacacttcga
 W  F  H  Q  E  V  H  T  H  R  P  Y  V  E  E  L  T  T  L  R 
gaccaagaggcacggcaattagataaacgttcaaacggtaagaaaagtagcttttttcaca
 D  Q  E  A  R  Q  L  D  K  R  S  N  G  K  K  S  S  F  F  T 
aacgagacgacatcacaagcgtcgagcgatcgaatcttgcgggtattacaaaatctggcc
 N  E  T  T  S  Q  A  S  S  D  R  I  L  R  V  L  Q  N  L  A 
ccccgacatgccgaagtggtacaaatacttgctagacttcaactcgacagcaaccaggat
 P  R  H  A  E  V  V  Q  I  L  A  R  L  Q  L  D  S  N  Q  D 
tggatcgaatatcgtgattttgcaacgcttgcaaaaaggcgtgtgtgattgccaaagac
 W  I  E  Y  R  D  F  C  N  A  C  K  K  A  C  V  I  A  K  D 
tctcagttgcgcgcctttttatcggagttaacggaccatgagttgatcgcaatgaacacg
 S  Q  L  R  A  F  L  S  E  L  T  D  H  E  L  I  A  M  N  T 
actgcaaatgcgaacaggaaaacggtgcgggtgccgcatccacgagcgcaactacaagaa
 T  A  N  A  N  R  K  T  V  R  V  P  H  P  R  A  Q  L  Q  E 
atcttggctatgccacgtaaacattga
 I  L  A  M  P  R  K  H  - 
```

**Fig. S4: DNA and amino acid sequence information of native ORC2 from *Phaeodactylum tricornutum*.**

The DNA sequence, translation, and genome coordinates of ORC2 from *Phaeodactylum tricornutum* in the genome database (assembly ASM15095v2) and the DNA sequence, translation of ORC2 single nucleotide polymorphisms used for cloning related to expression of FLAG-tag-fused ORC2 in transformants for chromatin immunoprecipitation sequencing.

>ORC4 (Phatr3_EG01462; Chr2: 24707-25948, 24149-24652)

```
atgactgttgttccggagactcccgtcggtaagggagagaacgagtgcaagtcccaatca
 M  T  V  V  P  E  T  P  V  G  K  G  E  N  E  C  K  S  Q  S 
cagacacacaatctatccagcttcattgcctacttaaatgatgttctgtacaacttatcc
 Q  T  H  N  L  S  S  F  I  A  Y  L  N  D  V  L  Y  N  L  S 
gatgctgccgtttacggctgtcgaactcgccaaagatcgctctggcggaaaaattcgccg
 D  A  A  V  Y  G  C  R  T  R  Q  R  S  L  W  R  K  N  S  P 
cctgcggattcttcagtcgtcaaggagtacgcaccacctcatttagagtcgcagtttgaa
 P  A  D  S  S  V  V  K  E  Y  A  P  P  H  L  E  S  Q  F  E 
gcaatgtaccgtatgttccgtccctgcctcgtccaacgtgacgagcacgtgaacgcttca
 A  M  Y  R  M  F  R  P  C  L  V  Q  R  D  E  H  V  N  A  S 
gccattgtgcaaggctgccgaggcagtggcaagaccttgctggtgaatcagtgctttcag
 A  I  V  Q  G  C  R  G  S  G  K  T  L  L  V  N  Q  C  F  Q 
gctttgcgagatgagctagatgcacagtctcacgccacaccaccctttcgagtcgtgcgc
 A  L  R  D  E  L  D  A  Q  S  H  A  T  P  P  F  R  V  V  R 
gtccacgggttgctcgtgccgggtgacaatgttgcattggtcgtacgagaaatgctaaac
 V  H  G  L  L  V  P  G  D  N  V  A  L  V  V  R  E  M  L  N 
cagctttccaacgcagcattcgaggagagtcaaacagatctggaacgacagcaacaacaa
 Q  L  S  N  A  A  F  E  E  S  Q  T  D  L  E  R  Q  Q  Q  Q 
gaacaacaggaaatggaatacaataccgatgaaccattatccaaacgtgccaaatcttct
 E  Q  Q  E  M  E  Y  N  T  D  E  P  L  S  K  R  A  K  S  S 
cgaaatcgactggaacgcttcttccagctacagtcgcaaaaggcatcgttcgtcaacaac
 R  N  R  L  E  R  F  F  Q  L  Q  S  Q  K  A  S  F  V  N  N 
attcaacttttcaacgaagtgttgcagattgccaaggacgacaatattcctatcgtaatt
 I  Q  L  F  N  E  V  L  Q  I  A  K  D  D  N  I  P  I  V  I 
gttctggatgagctcgatctttttctgggacaacagaaagcgacggcaaattctttggat
 V  L  D  E  L  D  L  F  L  G  Q  Q  K  A  T  A  N  S  L  D 
agtggtttgacggagcataaagatcgacaactgttactttatcaccttctcgatcgcgta
 S  G  L  T  E  H  K  D  R  Q  L  L  L  Y  H  L  L  D  R  V 
gcttccgacggttcatctgtgtgctttgtcggtatgacctgccatctaggcctcatcggt
 A  S  D  G  S  S  V  C  F  V  G  M  T  C  H  L  G  L  I  G 
ttactggaaaaacgaattcgtagtcgagccgagggtacatctcaatttcttcaatttgga
 L  L  E  K  R  I  R  S  R  A  E  G  T  S  Q  F  L  Q  F  G 
ccgtgcacgtccatcggcaaacttcgcgatattgtgtcgtgcaagttggaatcgacacga
 P  C  T  S  I  G  K  L  R  D  I  V  S  C  K  L  E  S  T  R 
aaagaccactatgatgcggcttatcagccatcgctttaccacgaaattcgcgcactcctg
 K  D  H  Y  D  A  A  Y  Q  P  S  L  Y  H  E  I  R  A  L  L 
accaaacctgcagaagacgacaatcatcccacaaggagacgagtttataatgctttcgtg
 T  K  P  A  E  D  D  N  H  P  T  R  R  R  V  Y  N  A  F  V 
cgggattgcatgttgggtcaagatgttcgctggttcagccgcgtcttgacattcgcgatc
 R  D  C  M  L  G  Q  D  V  R  W  F  S  R  V  L  T  F  A  I 
acaaactatcggatggattgcttggccgccatggaaggtgatgctagctcatcggcatct
 T  N  Y  R  M  D  C  L  A  A  M  E  G  D  A  S  S  S  A  S 
ccgattccaacatttgatgccgaatacctcttggaagcattggccgatatgggaggatcc
 P  I  P  T  F  D  A  E  Y  L  L  E  A  L  A  D  M  G  G  S 
ttggcatctgaatcgggagaacgcgacctggtgatagtagaggatgtggctgtcgatccc
 L  A  S  E  S  G  E  R  D  L  V  I  V  E  D  V  A  V  D  P 
cgtatgcaagccttgcgagatttgtcgggcccgcaagcagcgctggtattggctacgagg
 R  M  Q  A  L  R  D  L  S  G  P  Q  A  A  L  V  L  A  T  R 
cgaattctatcccgggacaatcgccgggagcaaagcaacgccgttttgacgcttgaacgc
 R  I  L  S  R  D  N  R  R  E  Q  S  N  A  V  L  T  L  E  R 
atgttggaagagtaccttggctcgtacaaaggtagttcaaatcggtattccaaacaagtt
 M  L  E  E  Y  L  G  S  Y  K  G  S  S  N  R  Y  S  K  Q  V 
ctgaccaaggcttttgttgacttgcttagtgttggcctgctacgtcccgcttcagatcat
 L  T  K  A  F  V  D  L  L  S  V  G  L  L  R  P  A  S  D  H 
agcggtgggcacccctgcagtaccaacatgatcgttccgtcttggatttggagccgtct
 S  G  G  A  P  L  Q  Y  Q  H  D  R  S  V  L  D  L  E  P  S 
cttttgcaagctgttcctttgcagctaacggttgacgtgtatcgcgaattggaaggagcg
 L  L  Q  A  V  P  L  Q  L  T  V  D  V  Y  R  E  L  E  G  A 
atagctcagaacattatgggttgttcgactgctctccgagagtgggggcgaaagatgaac
 I  A  Q  N  I  M  G  C  S  T  A  L  R  E  W  G  R  K  M  N 
tag
 - 
```

>ORC4 type A

```
atgactgttgttccggagactcccgtcggtaagggagagaacgagtgcaagtcccaatca
 M  T  V  V  P  E  T  P  V  G  K  G  E  N  E  C  K  S  Q  S
cagacacacaatctatccagcttcattgcctacttaaatgatgttctgtacaacttatcc
 Q  T  H  N  L  S  S  F  I  A  Y  L  N  D  V  L  Y  N  L  S
gatgctgccgtttacggctgtcgaactcgccaaagatcgctctggcggaaaaattcgccg
 D  A  A  V  Y  G  C  R  T  R  Q  R  S  L  W  R  K  N  S  P
cctgcggattcttcagtcgtcaaggagtacgcaccacctcatttagagtcgcagtttgaa
 P  A  D  S  S  V  V  K  E  Y  A  P  P  H  L  E  S  Q  F  E
gcaatgtaccgtatgttccgtccctgcctcgtccaacgtgacgagcacgtgaacgcttca
 A  M  Y  R  M  F  R  P  C  L  V  Q  R  D  E  H  V  N  A  S
gccattgtgcaaggctgccgaggcagtggcaagaccttgctggtgaatcagtgctttcag
 A  I  V  Q  G  C  R  G  S  G  K  T  L  L  V  N  Q  C  F  Q
gctttgcgagatgagctagatgcacagtctcacgccacaccaccctttcgagtcgtgcgc
 A  L  R  D  E  L  D  A  Q  S  H  A  T  P  P  F  R  V  V  R
gtccacgggttgctcgtgccgggtgacaatgttgcattggttgtacgagaaatgctaaac
 V  H  G  L  L  V  P  G  D  N  V  A  L  V  V  R  E  M  L  N
cagcttgccaacgcagcattcgaggagagtcaaacagatctggaacgacagcaacaacaa
 Q  L  A  N  A  A  F  E  E  S  Q  T  D  L  E  R  Q  Q  Q  Q
gaacaacaggaaatggaatacaatactgatgaaccattatccaaacgtgccaaatcttct
 E  Q  Q  E  M  E  Y  N  T  D  E  P  L  S  K  R  A  K  S  S
cgaagtcgactggaacgcttcttccagctacagtcgcaaaaggcatcgttcgtcaacaac
 R  S  R  L  E  R  F  F  Q  L  Q  S  Q  K  A  S  F  V  N  N
attcaacttttcaacgaagtgttgcagattgccaaggacgacaatattcctatcgtaatt
 I  Q  L  F  N  E  V  L  Q  I  A  K  D  D  N  I  P  I  V  I
gttctggatgagctcgatctttttctgggacaacagaaagcgacggcaaattctttggat
 V  L  D  E  L  D  L  F  L  G  Q  Q  K  A  T  A  N  S  L  D
agtggtttgacggagcataaagatcgacaactgttactttatcaccttctcgatcgcgta
 S  G  L  T  E  H  K  D  R  Q  L  L  L  Y  H  L  L  D  R  V
gcttccgacggttcatctgtgtgctttgtcggtatgacctgccatctaggcctcatcggt
 A  S  D  G  S  S  V  C  F  V  G  M  T  C  H  L  G  L  I  G
ttactggaaaaacgaattcgtagtcgagccgagggtacatctcaatttcttcaatttgga
 L  L  E  K  R  I  R  S  R  A  E  G  T  S  Q  F  L  Q  F  G
ccgtgcacgtccatcggcaaacttcgcgatattgtgtcgtgcaagttggaatcgacacga
 P  C  T  S  I  G  K  L  R  D  I  V  S  C  K  L  E  S  T  R
aaagaccactatgatgcggcttatcagccatcgctttaccacgaaattcgcgcactcctg
 K  D  H  Y  D  A  A  Y  Q  P  S  L  Y  H  E  I  R  A  L  L
accaaacctgcagaagacgacaatcatcccacaaggagacgagtttataatgctttcgtg
 T  K  P  A  E  D  D  N  H  P  T  R  R  R  V  Y  N  A  F  V
cgggattgcatgttgggtcaagatgttcgctggttcagccgcgtcttgacattcgcgatt
 R  D  C  M  L  G  Q  D  V  R  W  F  S  R  V  L  T  F  A  I
acaaactatcggatggattgcttggccgccatggaaggtgatgctagctcatcggcatct
 T  N  Y  R  M  D  C  L  A  A  M  E  G  D  A  S  S  S  A  S
ccgattccaacatttgatgccgaatacctcttggaagcattggccgatatgggaggatcc
 P  I  P  T  F  D  A  E  Y  L  L  E  A  L  A  D  M  G  G  S
ttggcatctgaatcgggagaacgcgacctggtgatagtagaggatgtggctgtcgatccc
 L  A  S  E  S  G  E  R  D  L  V  I  V  E  D  V  A  V  D  P
cgtatgcaagccttgcgagatttgtcgggcccgcaagcagcgctggtattggctacgagg
 R  M  Q  A  L  R  D  L  S  G  P  Q  A  A  L  V  L  A  T  R
cgaattctatcccgggacaatcgccgggagcaaagcaacgccgttttgacgcttgaacgc
 R  I  L  S  R  D  N  R  R  E  Q  S  N  A  V  L  T  L  E  R
atgttggaagagtaccttggctcgtacaaaggtagttcaaatcggtattccaaacaagtt
 M  L  E  E  Y  L  G  S  Y  K  G  S  S  N  R  Y  S  K  Q  V
ctgaccaaggcttttgttgacttgcttagtgttggcctgctacgtcccgcttcagatcat
 L  T  K  A  F  V  D  L  L  S  V  G  L  L  R  P  A  S  D  H
agcggtggggcacccctgcagtaccaacatgatcgttccgtcttggatttggagccgtct
 S  G  G  A  P  L  Q  Y  Q  H  D  R  S  V  L  D  L  E  P  S
cttttgcaagctgttcctttgcagctaacggttgacgtgtatcgcgaattggaaggagcg
 L  L  Q  A  V  P  L  Q  L  T  V  D  V  Y  R  E  L  E  G  A
atagctcagaacattatgggttgttcgactgctctccgagagtgggggcgaaagatgaac
 I  A  Q  N  I  M  G  C  S  T  A  L  R  E  W  G  R  K  M  N
tag
 -
```

>ORC4 type B

```
atgactgttgttccggagactcccgtcggtaagggagagaacgagtgcaagtcccaatca
 M  T  V  V  P  E  T  P  V  G  K  G  E  N  E  C  K  S  Q  S
cagacacacaatctatccagcttcattgcctacttaaatgatgttctgtacaacttatcc
 Q  T  H  N  L  S  S  F  I  A  Y  L  N  D  V  L  Y  N  L  S
gatgctgccgtttacggctgtcgaactcgccaaagatcgctctggcagaaaaattcgccg
 D  A  A  V  Y  G  C  R  T  R  Q  R  S  L  W  Q  K  N  S  P
cctgcggattcttcagtcgtcaaggagtacgcaccacctcatttagagtcgcagtttgaa
 P  A  D  S  S  V  V  K  E  Y  A  P  P  H  L  E  S  Q  F  E
gcaatgtaccgtatgttccgtccctgcctcgtccaacgtgacgagcacgtgaacgcttca
 A  M  Y  R  M  F  R  P  C  L  V  Q  R  D  E  H  V  N  A  S
gccattctgcaaggctgccgaggcagtggcaagaccttgctggtgaatcagtgctttcag
 A  I  L  Q  G  C  R  G  S  G  K  T  L  L  V  N  Q  C  F  Q
gctttgcgagatgagctagatgcacagtctcacgccacaccaccctttcgagtcgtgcgc
 A  L  R  D  E  L  D  A  Q  S  H  A  T  P  P  F  R  V  V  R
gtccacgggttgctcgtgccgggtgacaatgttgcactggtcgtacgagaaatgctaaac
 V  H  G  L  L  V  P  G  D  N  V  A  L  V  V  R  E  M  L  N
cagctttccaacgcagcattcgaggagagtcaaacagatctggaacgacagcaacaacaa
 Q  L  S  N  A  A  F  E  E  S  Q  T  D  L  E  R  Q  Q  Q  Q
gaacaacaggaaatggaatacaatgccgatgaaccattatccaaacgtgccaaatcttct
 E  Q  Q  E  M  E  Y  N  A  D  E  P  L  S  K  R  A  K  S  S
cgaaatcgactggaacgcttcttccagctacagtcgcaaaaggcatcgttcgtcaacaat
 R  N  R  L  E  R  F  F  Q  L  Q  S  Q  K  A  S  F  V  N  N
attcaacttttcaacgaagtgttgcagattgccaaggacgacaatattcctatcgtaatt
 I  Q  L  F  N  E  V  L  Q  I  A  K  D  D  N  I  P  I  V  I
gttctggatgagctcgatctttttctgggacaacagaaagcgacggcaaattctttggat
 V  L  D  E  L  D  L  F  L  G  Q  Q  K  A  T  A  N  S  L  D
agtggtttgacggagcataaagatcgacaactgttactttatcaccttctcgatcgcgta
 S  G  L  T  E  H  K  D  R  Q  L  L  L  Y  H  L  L  D  R  V
gcttccgacggttcatctgtgtgctttgtcggtatgacctgccatctaggcctcatcggt
 A  S  D  G  S  S  V  C  F  V  G  M  T  C  H  L  G  L  I  G
ttactggaaaaacgaattcgtagtcgagccgagggtacatctcaatttcttcaatttgga
 L  L  E  K  R  I  R  S  R  A  E  G  T  S  Q  F  L  Q  F  G
ccgtgcacgtccatcggcaaacttcgcgatattgtgtcgtgcaagttggaatcgacacga
 P  C  T  S  I  G  K  L  R  D  I  V  S  C  K  L  E  S  T  R
aaagaccactatgatgcggcttatcagccatcgctttaccacgaaattcgcgcactcctg
 K  D  H  Y  D  A  A  Y  Q  P  S  L  Y  H  E  I  R  A  L  L
accaaacctgcagaagacgacaatcatcccacaaggagacgagtttataatgctttcgtg
 T  K  P  A  E  D  D  N  H  P  T  R  R  R  V  Y  N  A  F  V
cgggattgcatgttgggtcaagatgttcgctggttcagccgcgtcttgacattcgcgatc
 R  D  C  M  L  G  Q  D  V  R  W  F  S  R  V  L  T  F  A  I
acaaactatcggatggattgcttggccgccatggaaggtgatgctagctcatcggcatct
 T  N  Y  R  M  D  C  L  A  A  M  E  G  D  A  S  S  S  A  S
ccgattccaacatttgatgccgaatacctcttggaagcattggccgatatgggaggatcc
 P  I  P  T  F  D  A  E  Y  L  L  E  A  L  A  D  M  G  G  S
ttggcatctgaatcgggagaacgcgacctggtgatagtagaggatgtggctgtcgatccc
 L  A  S  E  S  G  E  R  D  L  V  I  V  E  D  V  A  V  D  P
cgtatgcaagccttgcgagatttgtcgggcccgcaagcagcgctgatattggctacgagg
 R  M  Q  A  L  R  D  L  S  G  P  Q  A  A  L  I  L  A  T  R
cgaattctatcccgggacaatcgccgggagcaaagcaacgccgttttgacgcttgaacgc
 R  I  L  S  R  D  N  R  R  E  Q  S  N  A  V  L  T  L  E  R
atgttggaagagtaccttggctcgtacaaaggtagttcaaatcggtattccaaacaagtt
 M  L  E  E  Y  L  G  S  Y  K  G  S  S  N  R  Y  S  K  Q  V
ctgaccaaggcttttgttgacttgcttagtgttggcctgctacgtcccgcttcagatcat
 L  T  K  A  F  V  D  L  L  S  V  G  L  L  R  P  A  S  D  H
agcggtggggcacccctgcagtaccaacatgatcgttccgtcttggatttggagccgtct
 S  G  G  A  P  L  Q  Y  Q  H  D  R  S  V  L  D  L  E  P  S
cttttgcaagctgttcctttgcagctaacggttgacgtgtatcgcgaattggaaggagcg
 L  L  Q  A  V  P  L  Q  L  T  V  D  V  Y  R  E  L  E  G  A
atagctcagaacattatgggttgttcgactgctctccgagagtgggggcgaaagatgaac
 I  A  Q  N  I  M  G  C  S  T  A  L  R  E  W  G  R  K  M  N
tag
 -
```

**Fig. S5: DNA and amino acid sequence information of native ORC4 from *Phaeodactylum tricornutum*.**

The DNA sequence, translation, and genome coordinates of ORC4 from *Phaeodactylum tricornutum* in the genome database (assembly ASM15095v2) and the DNA sequence, translation of ORC4 single nucleotide polymorphisms used for cloning related to expression of FLAG-tag-fused ORC4 in transformants for chromatin immunoprecipitation sequencing.

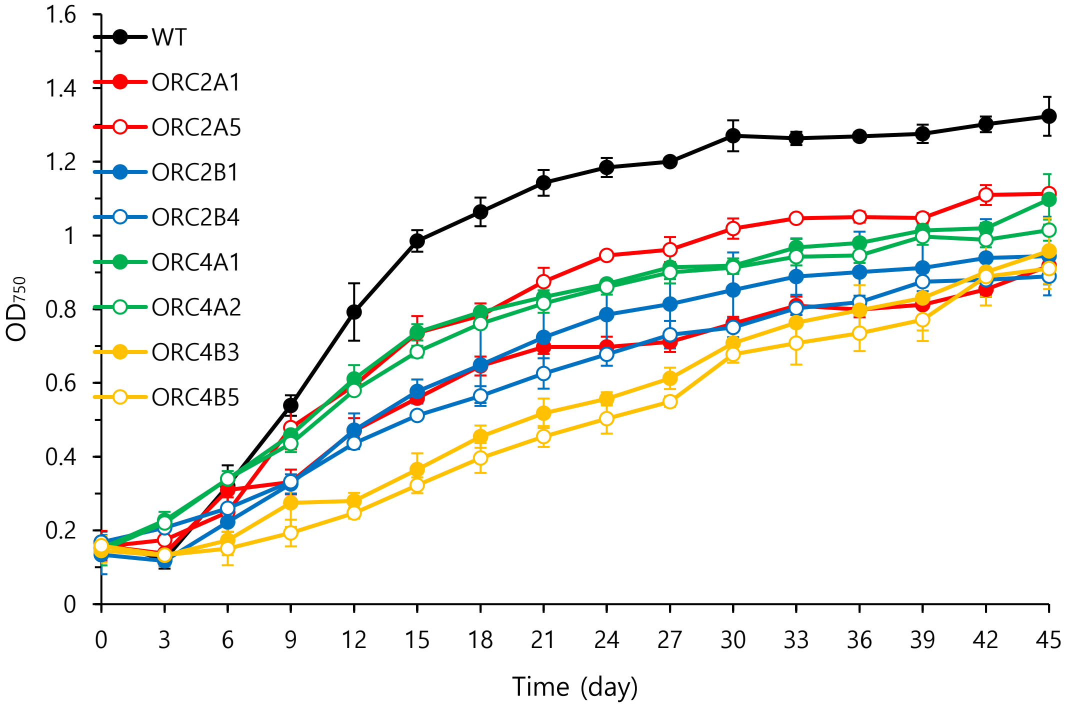


**Fig. S6: Growth patterns of wild type and transformant *Phaeodactylum tricornutum* strains.**

Under the same culture conditions (f/2 medium, shaker: 130 rpm, temperature: 20°C, light: 40 µmol photons $m^{-2}$ $s^{-1}$), the transformant showed more inhibited growth than the wild type did.

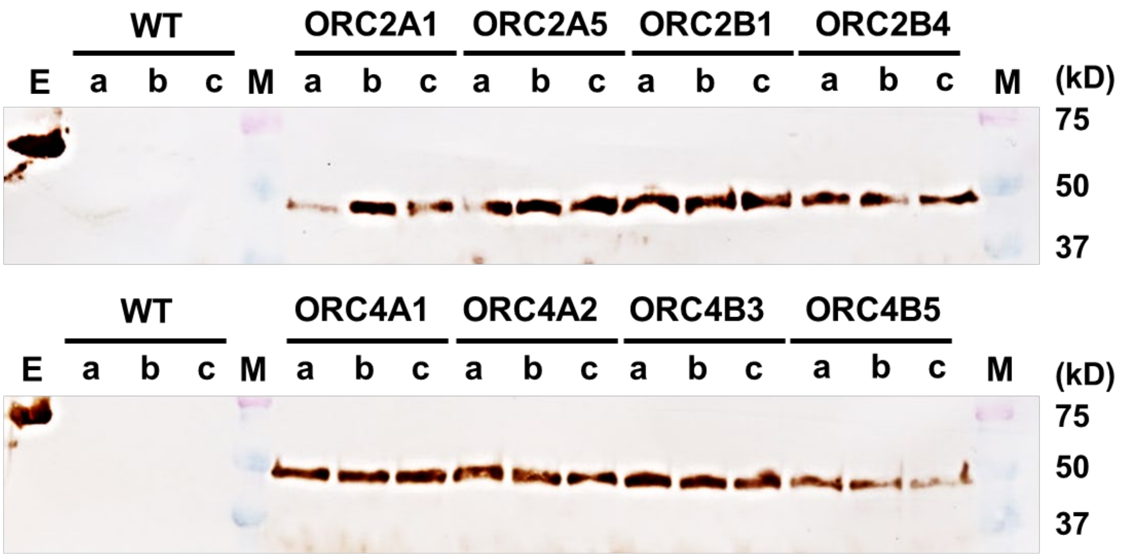


**Fig. S7: Western blotting results visualising the expression level of *Phaeodactylum tricornutum* transformants during growth.**

While the transformants were cultured, expression of the FLAG-tag-fused ORC subunit was confirmed. Based on the optical density value measured at 750 nm, the transformants expressed the FLAG-tag-fused ORC subunit in all sections ([a]0.3–0.4, [b]0.5–0.6, and [c]0.7–0.8). WT: wild type, E: Epitope Tag Protein Marker Lysate, M: Protein Dual Color Standards ladder.

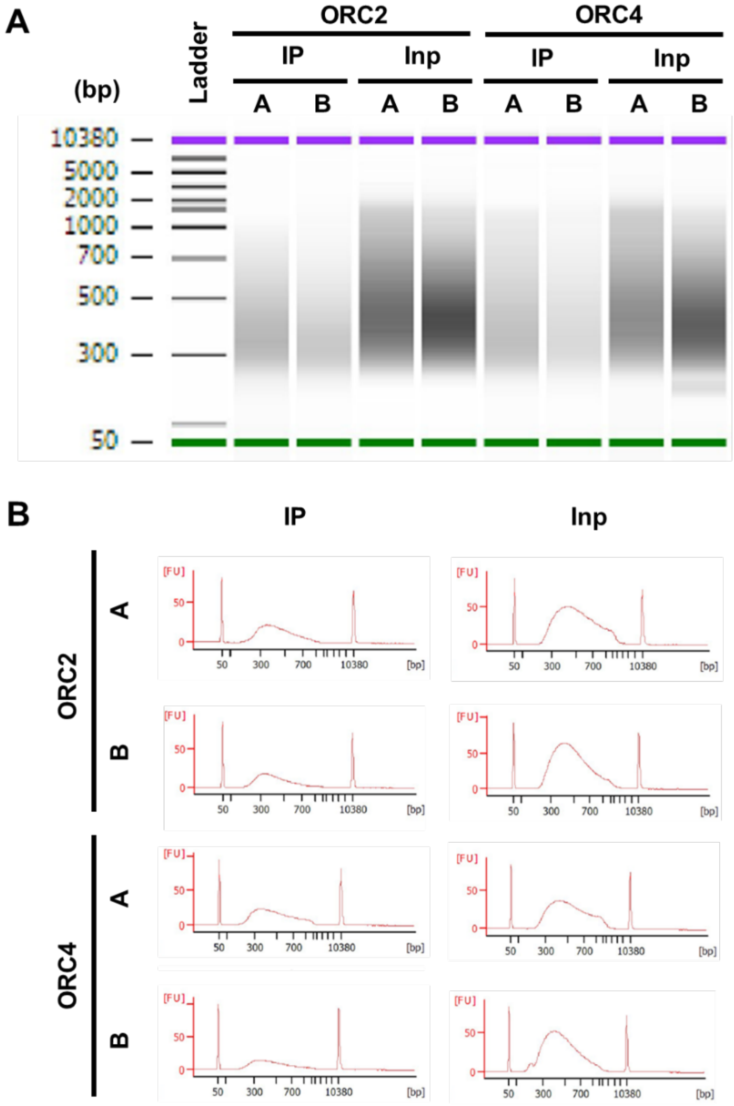


**Fig. S8: Quality of library samples.**

The concentration and fragment length of library samples were visualised via Bioanalyzer Systems. (A) Gel images of the DNA fragments included in the library samples and (B) length distribution of the DNA fragments are shown. Inp: input sample (negative control), IP: immunoprecipitation sample.

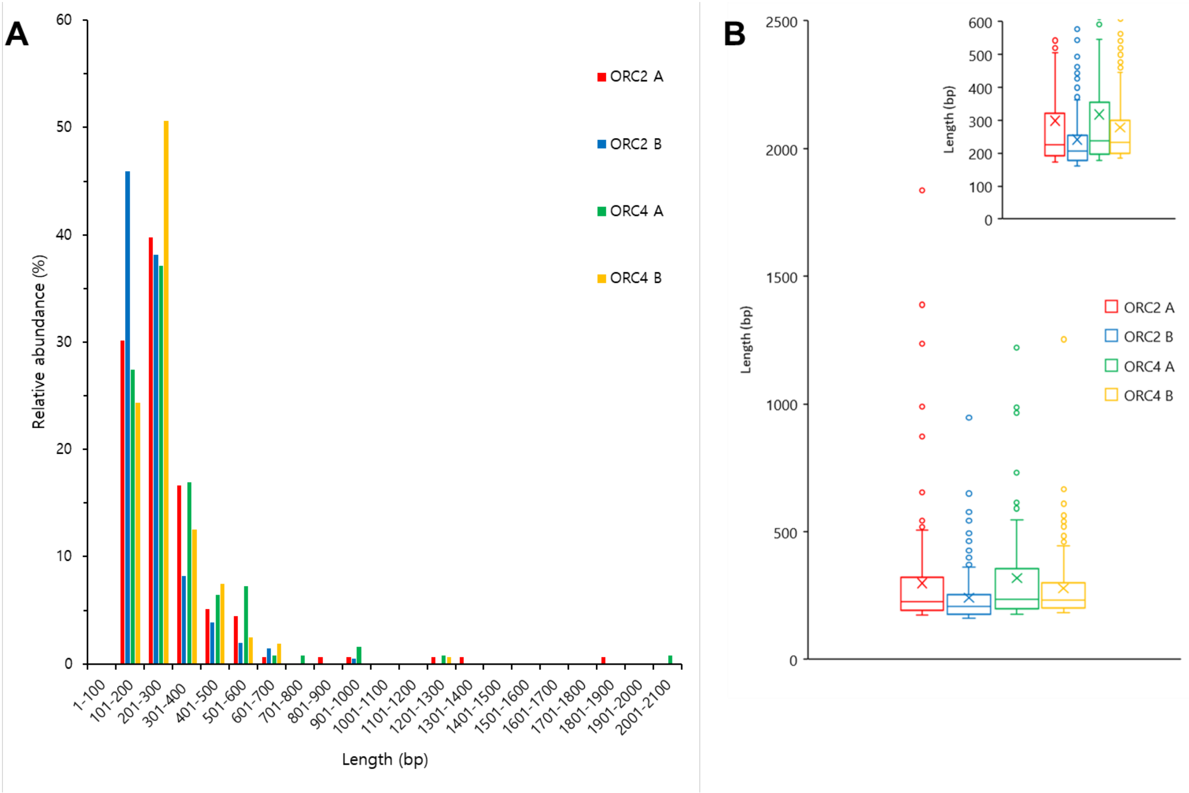


**Fig. S9: Length of sites screened using ORC2 and ORC4.**

Via chromatin immunoprecipitation sequencing, the binding sites of FLAG-tag-fused ORC2 and ORC4 were screened, and information on the length of the binding sites obtained. (A) The length distribution of the screened sites was visualised by grouping them in 10-bp increments from 1–2100 bp, and (B) statistical information about the length of the sites presented in a box-and-whisker plot.

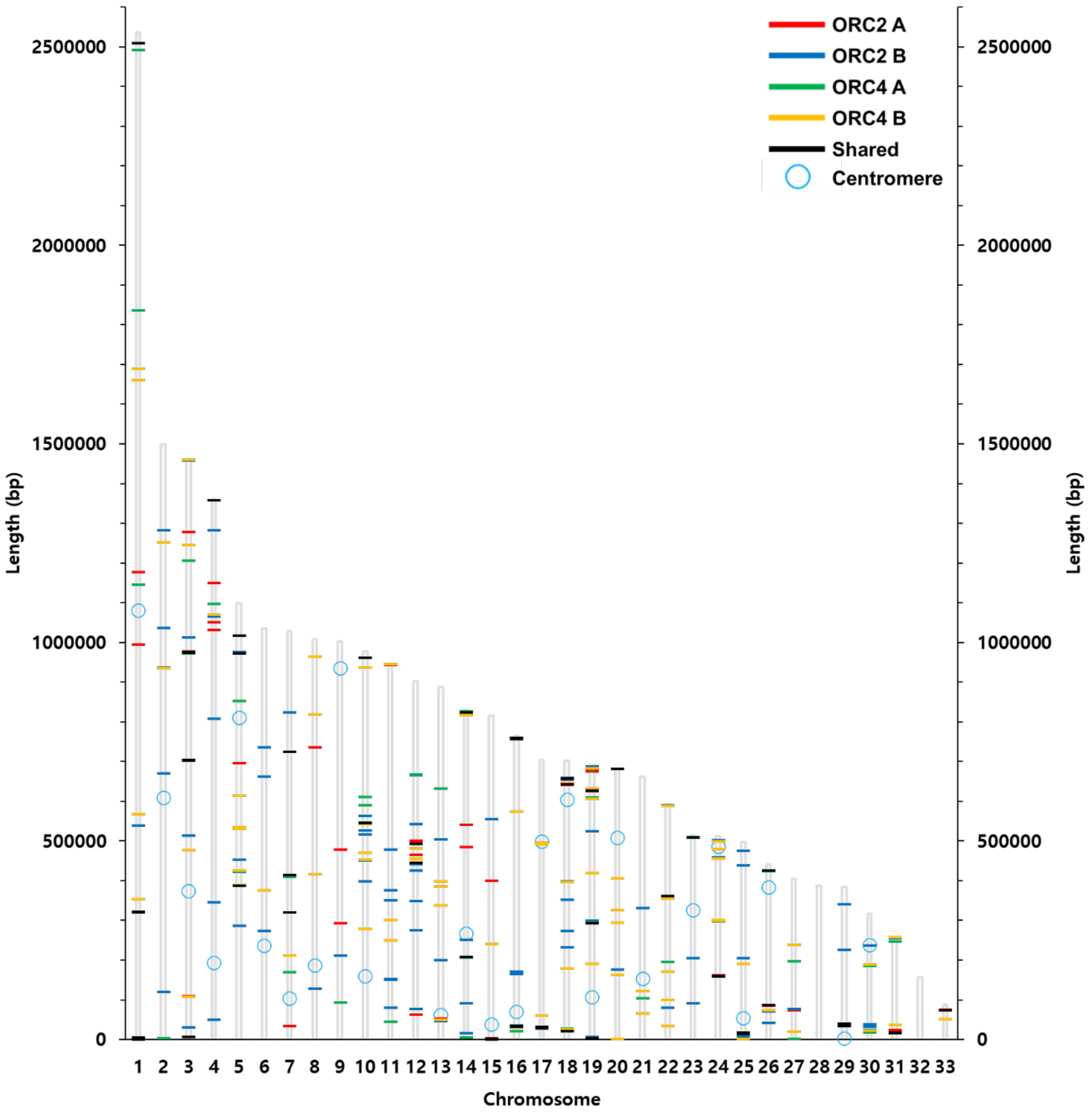


**Fig. S10: Distribution of screened sites on the genome of *Phaeodactylum tricornutum*.**

The locations of sites screened using ORC2 A (red lines), ORC2 B (blue lines), ORC4 A (green lines), and ORC4 B (orange lines) are indicated on the genome of *Phaeodactylum tricornutum*. The locations of shared sites are indicated by black lines, and those of centromeres indicated by light blue circles (Diner *et al.*, 2017). There were 18 ranges within 100 kbp with more than five sites clustered together (Chr 1, 0–100 kbp: 10 sites; Chr 3, 900–1000 kbp: 5 sites; Chr 10, 500–600 kbp: 7 sites; Chr 12, 400–500 kbp: 11 sites; Chr 15, 0–100 kbp: 6 sites; Chr 16, 0–100 kbp: 8 sites; Chr 16, 700–764.225 kbp: 8 sites; Chr 17, 0–100 kbp: 7 sites; Chr 18, 600–700 kbp: 10 sites; Chr 19, 200–300 kbp: 5 sites; Chr 19, 600–700 kbp: 13 sites; Chr 24, 100–200 kbp: 7 sites; Chr 24, 410–510 kbp: 5 sites; Chr 25, 0–100 kbp: 14 sites; Chr 26, 0–100 kbp: 7 sites; Chr 29, 0–100 kbp: 6 sites; Chr 30, 0–100 kbp: 6 sites; Chr 31, 0–100 kbp: 7 sites).

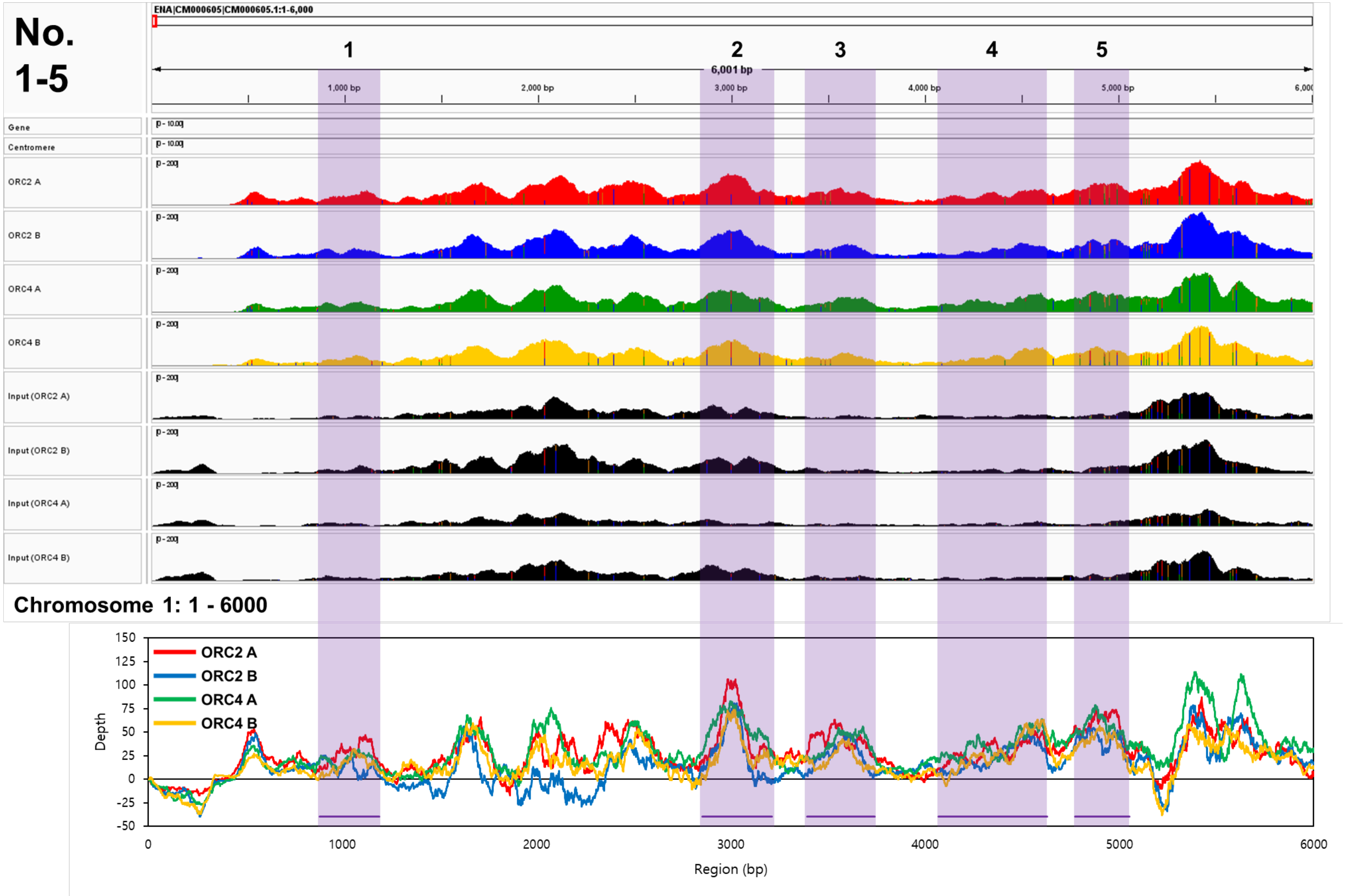

No. 1-5
ENA|CM000605|CM000605.1:1-6,000
1
2
3
4
5
6,001 bp
1,000 bp
2,000 bp
3,000 bp
4,000 bp
5,000 bp
Gene
Centromere
ORC2 A
ORC2 B
ORC4 A
ORC4 B
Input (ORC2 A)
Input (ORC2 B)
Input (ORC4 A)
Input (ORC4 B)
Chromosome 1: 1 - 6000
ORC2 A
ORC2 B
ORC4 A
ORC4 B
Depth
Region (bp)

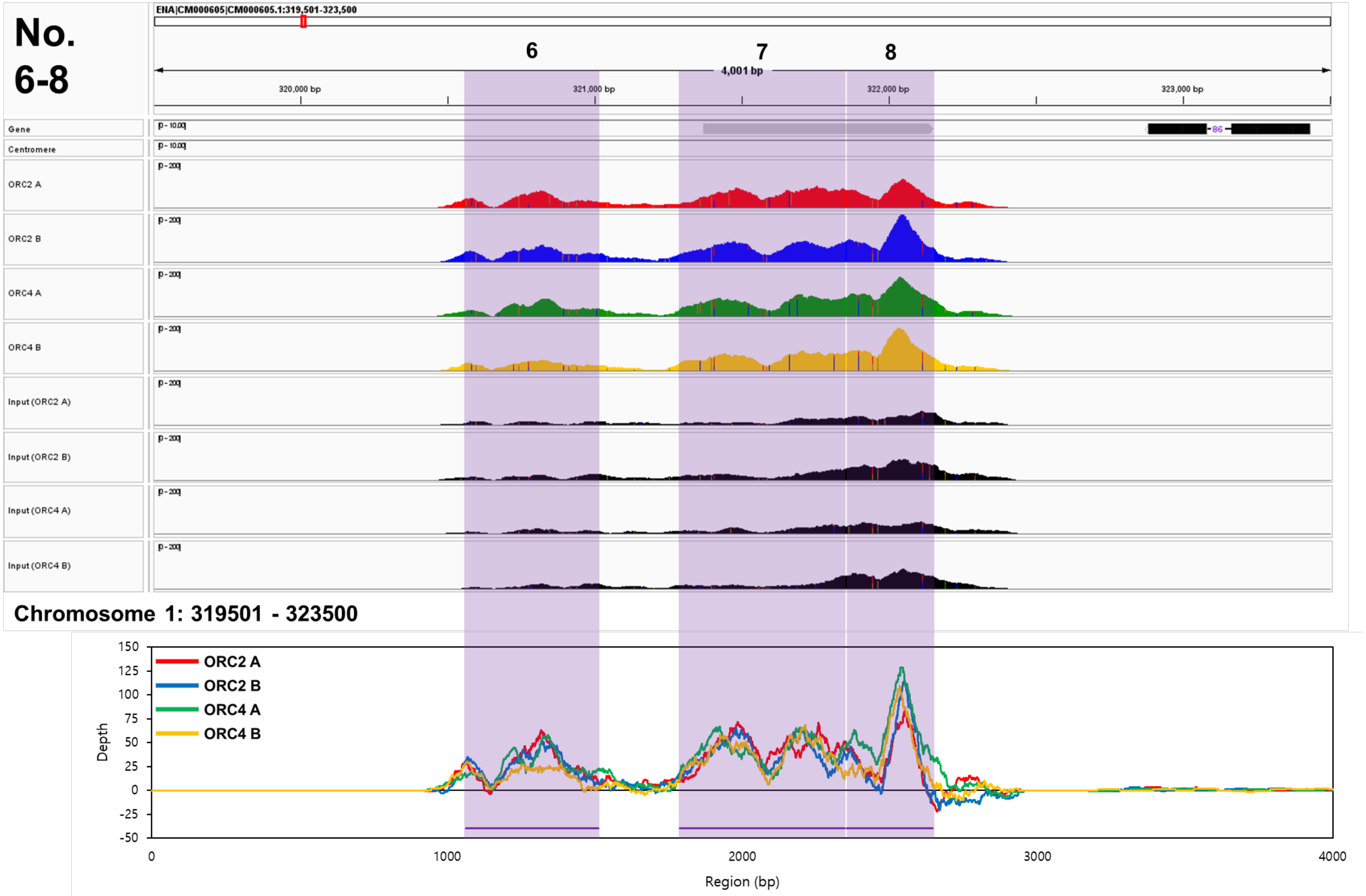
No.
6-8
ENA|CM000605|CM000605.1:319,501-323,500
6
7
8
4,001 bp
320,000 bp
321,000 bp
322,000 bp
323,000 bp
Gene
Centromere
ORC2 A
ORC2 B
ORC4 A
ORC4 B
Input (ORC2 A)
Input (ORC2 B)
Input (ORC4 A)
Input (ORC4 B)
Chromosome 1: 319501 - 323500
ORC2 A
ORC2 B
ORC4 A
ORC4 B
Depth
150
125
100
75
50
25
0
-25
-50
0
1000
2000
3000
4000
Region (bp)

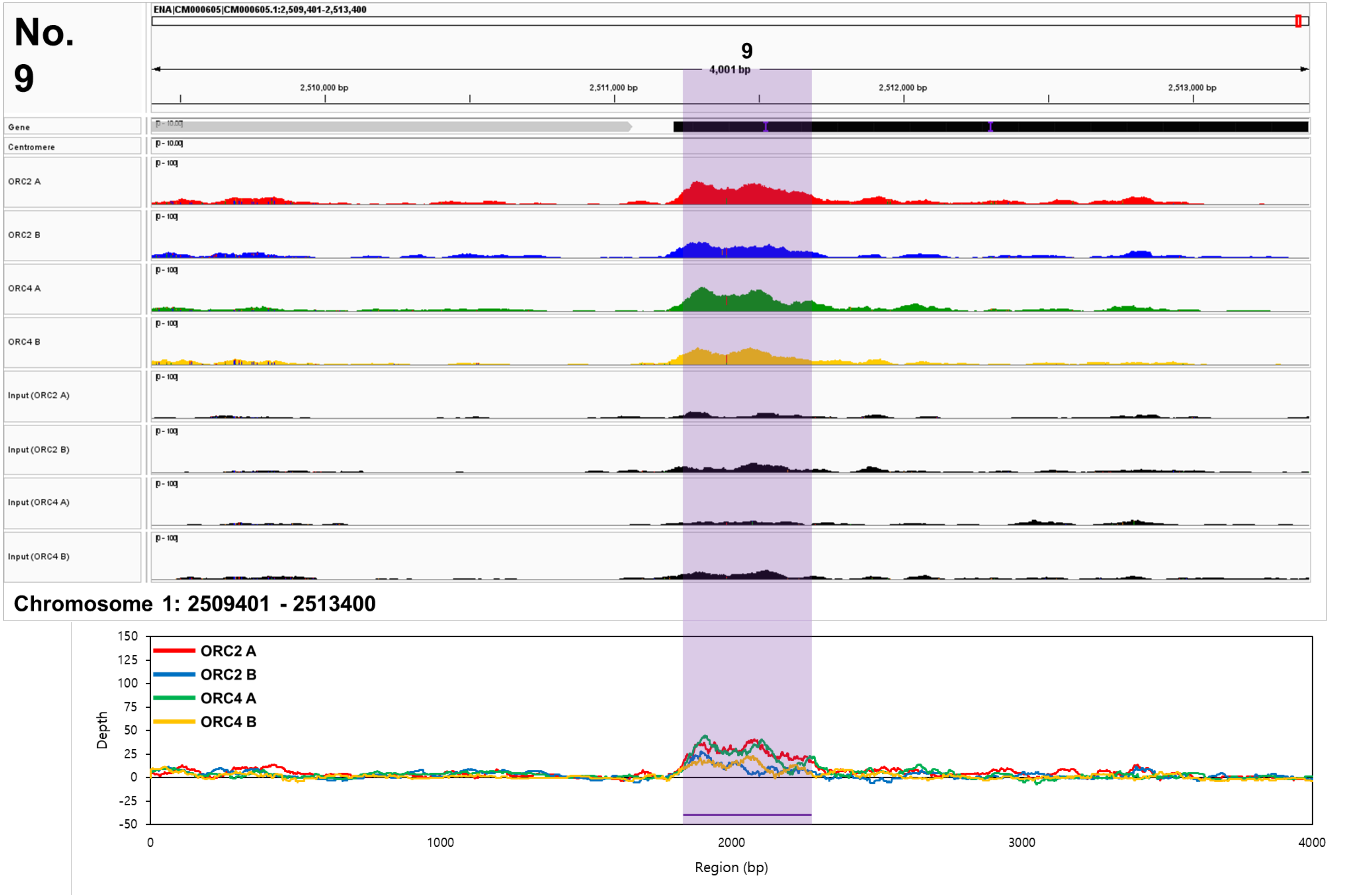
No. 9
ENA|CM000605|CM000605.1:2,509,401-2,513,400
9
4,001 bp
2,510,000 bp
2,511,000 bp
2,512,000 bp
2,513,000 bp
Gene
Centromere
ORC2 A
ORC2 B
ORC4 A
ORC4 B
Input (ORC2 A)
Input (ORC2 B)
Input (ORC4 A)
Input (ORC4 B)
Chromosome 1: 2509401 - 2513400
ORC2 A
ORC2 B
ORC4 A
ORC4 B
Depth
Region (bp)

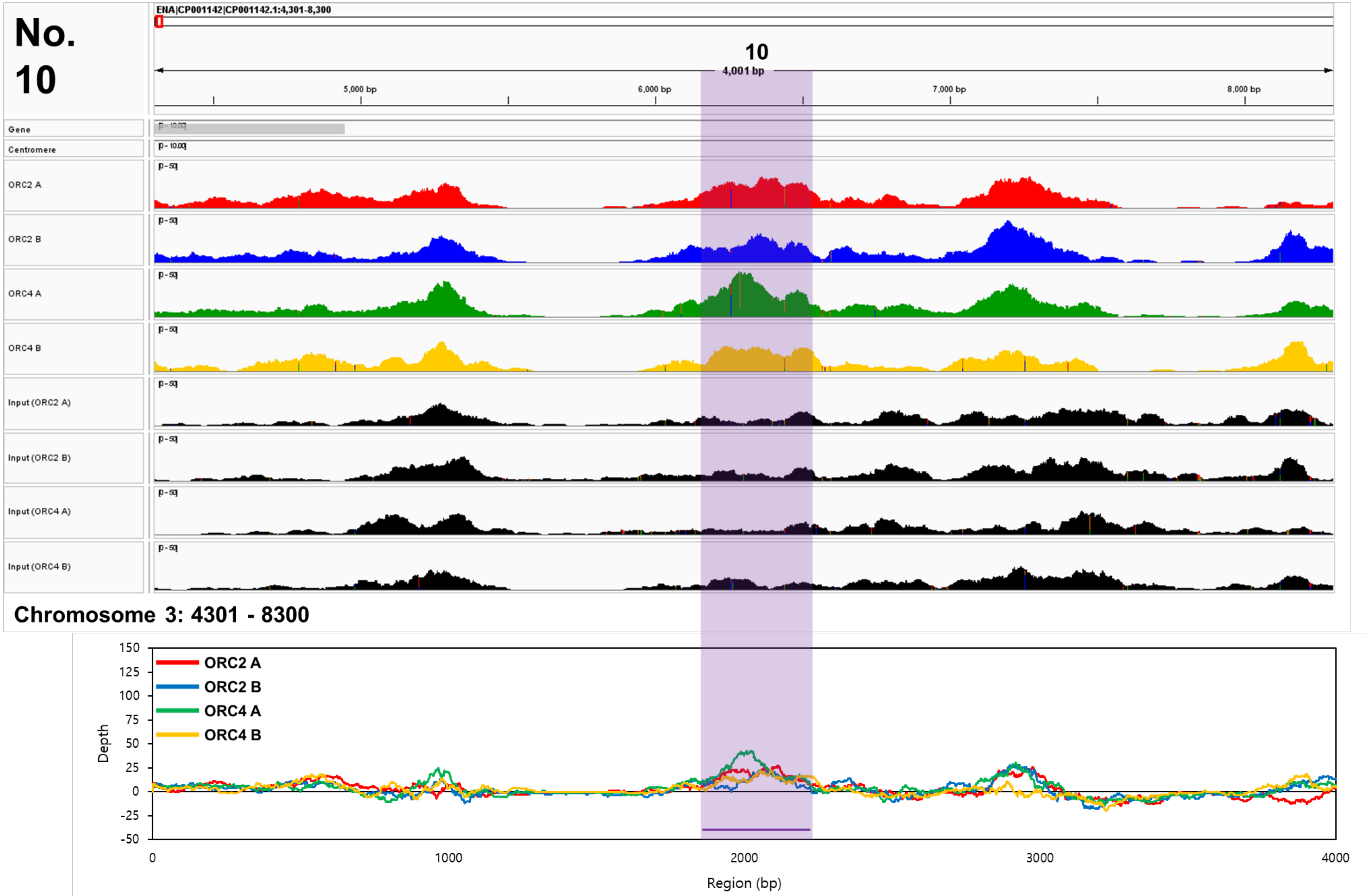
No. 10
ENA|CP001142|CP001142.1:4,301-8,300
10
4,001 bp
5,000 bp
6,000 bp
7,000 bp
8,000 bp
Gene
Centromere
ORC2 A
ORC2 B
ORC4 A
ORC4 B
Input (ORC2 A)
Input (ORC2 B)
Input (ORC4 A)
Input (ORC4 B)
Chromosome 3: 4301 - 8300
ORC2 A
ORC2 B
ORC4 A
ORC4 B
Depth
Region (bp)

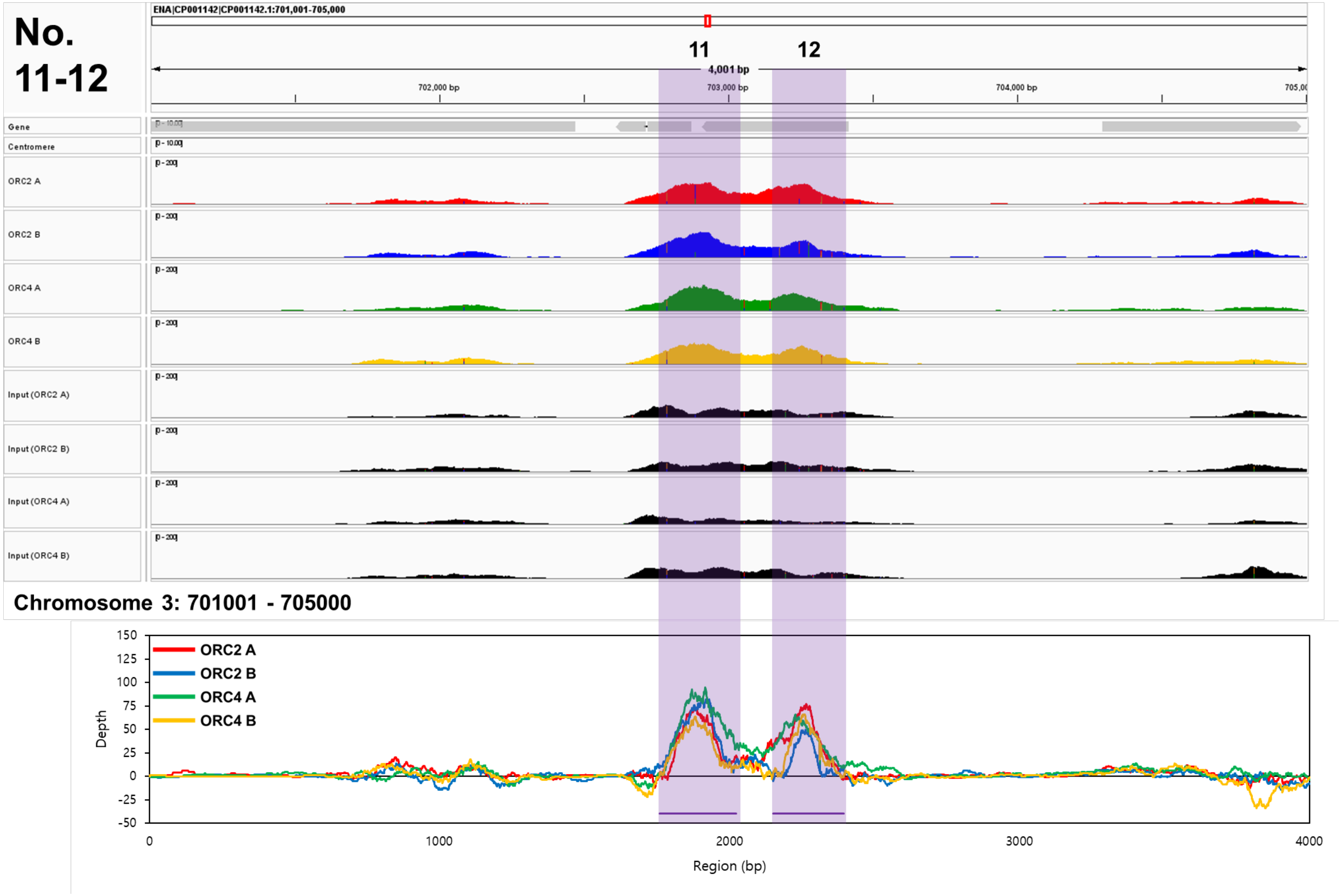
No. 11-12
ENA|CP001142|CP001142.1:701,001-705,000
11
12
4,001 bp
702,000 bp
703,000 bp
704,000 bp
Gene
Centromere
ORC2 A
ORC2 B
ORC4 A
ORC4 B
Input (ORC2 A)
Input (ORC2 B)
Input (ORC4 A)
Input (ORC4 B)
Chromosome 3: 701001 - 705000
ORC2 A
ORC2 B
ORC4 A
ORC4 B
Depth
Region (bp)
150
125
100
75
50
25
0
-25
-50
0
1000
2000
3000
4000

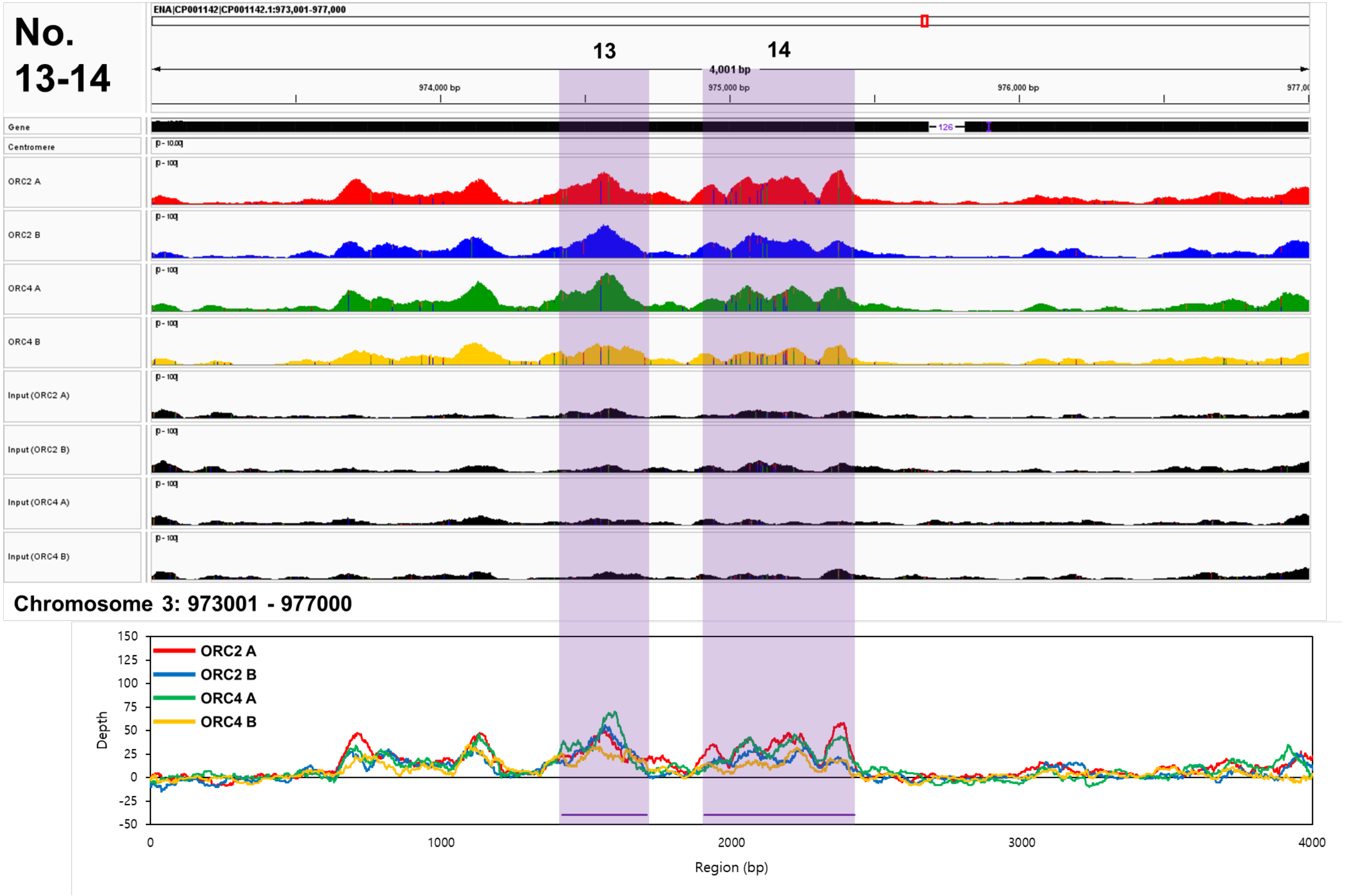
No. 13-14
ENA|CP001142|CP001142.1:973,001-977,000
13
14
4,001 bp
974,000 bp
975,000 bp
976,000 bp
Gene
Centromere
ORC2 A
ORC2 B
ORC4 A
ORC4 B
Input (ORC2 A)
Input (ORC2 B)
Input (ORC4 A)
Input (ORC4 B)
Chromosome 3: 973001 - 977000
ORC2 A
ORC2 B
ORC4 A
ORC4 B
Depth
Region (bp)
150
125
100
75
50
25
0
-25
-50
0
1000
2000
3000
4000

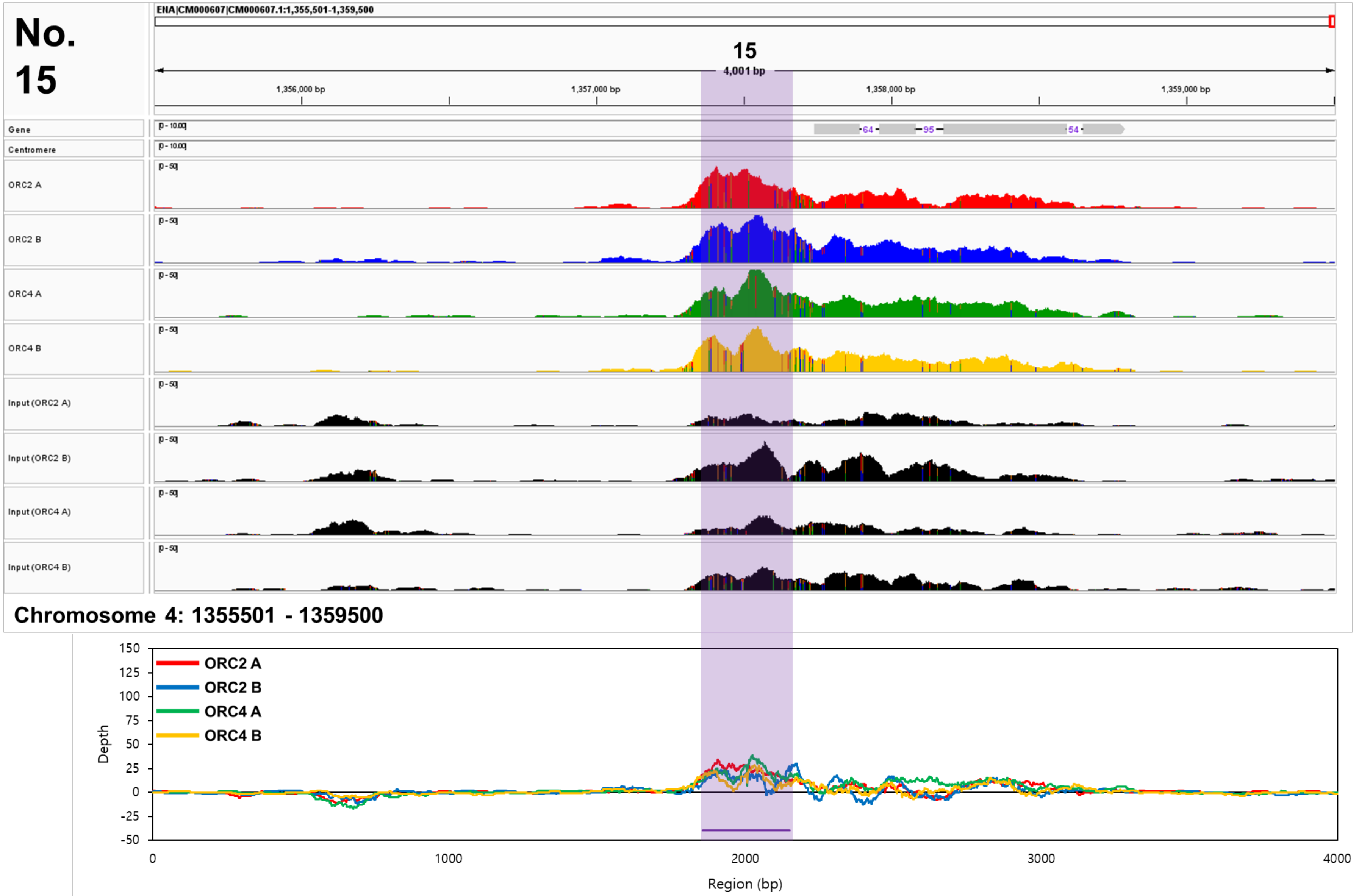
No. 15
ENA|CM000607|CM000607.1:1,355,501-1,359,500
15
4,001 bp
1,356,000 bp
1,357,000 bp
1,358,000 bp
1,359,000 bp
Gene
Centromere
ORC2 A
ORC2 B
ORC4 A
ORC4 B
Input (ORC2 A)
Input (ORC2 B)
Input (ORC4 A)
Input (ORC4 B)
Chromosome 4: 1355501 - 1359500
ORC2 A
ORC2 B
ORC4 A
ORC4 B
Depth
Region (bp)

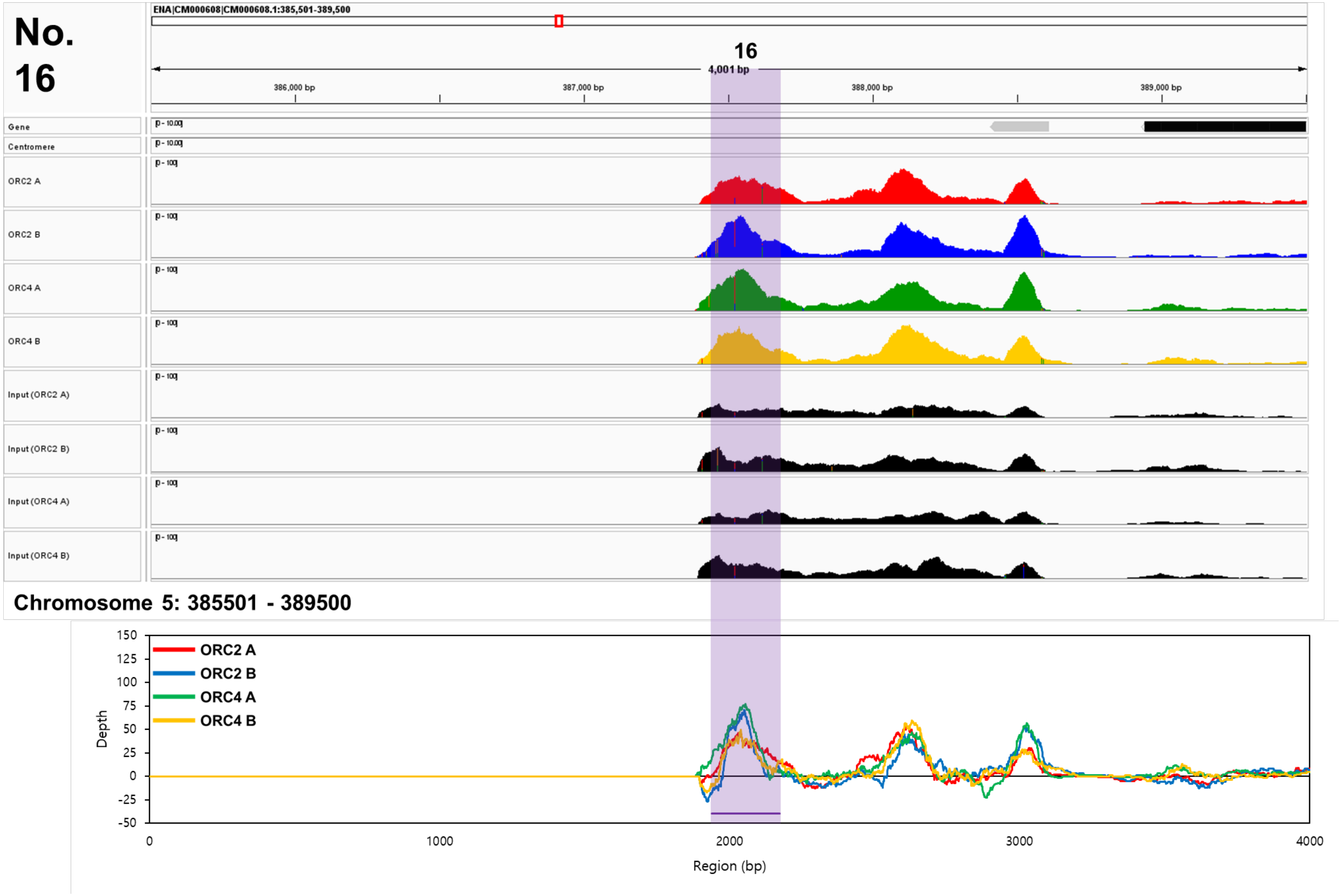
No. 16
ENA|CM000608|CM000608.1:385,501-389,500
16
4,001 bp
386,000 bp
387,000 bp
388,000 bp
389,000 bp
Gene
Centromere
ORC2 A
ORC2 B
ORC4 A
ORC4 B
Input (ORC2 A)
Input (ORC2 B)
Input (ORC4 A)
Input (ORC4 B)
Chromosome 5: 385501 - 389500
ORC2 A
ORC2 B
ORC4 A
ORC4 B
Depth
Region (bp)

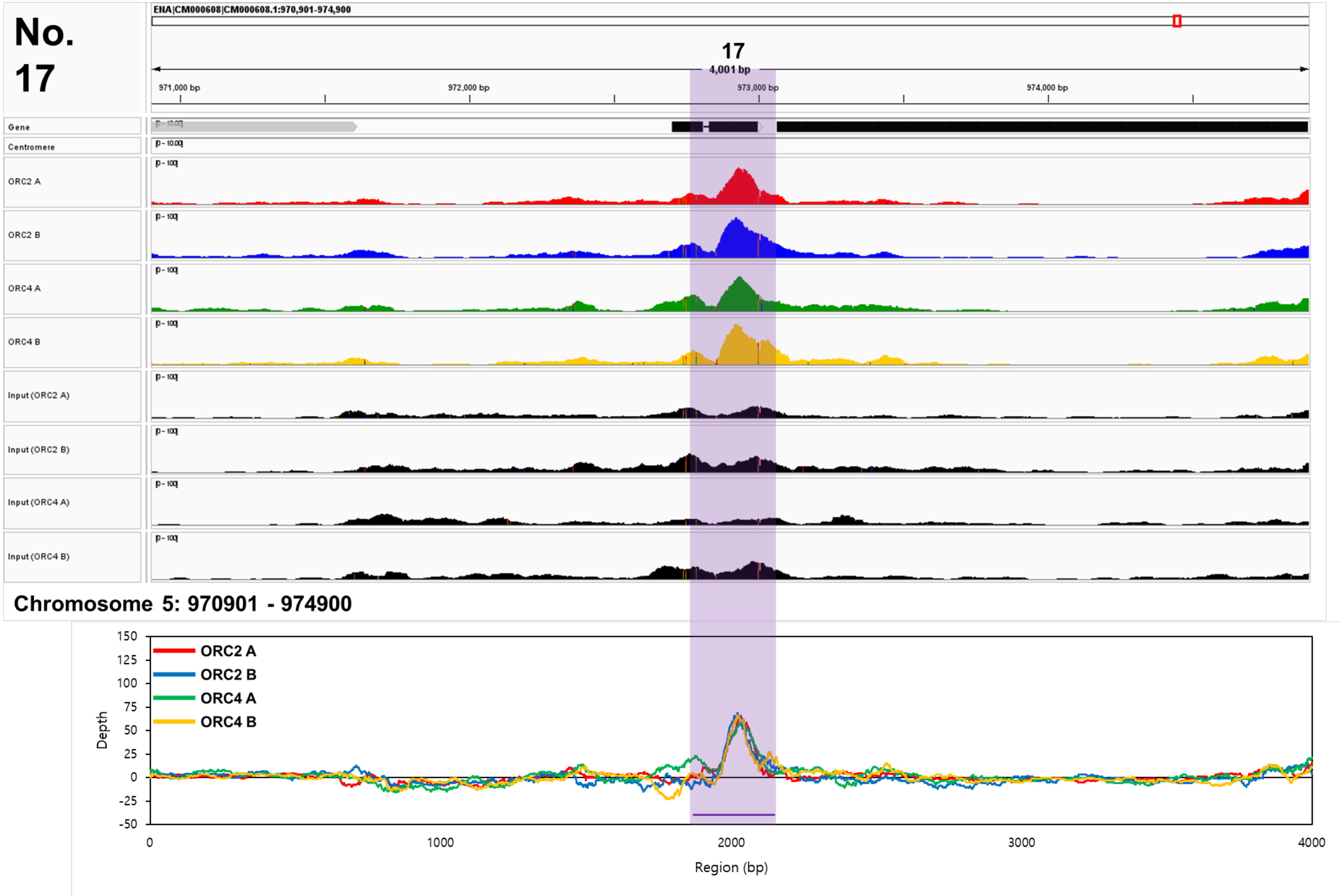

No. 17
ENA|CM000608|CM000608.1:970,901-974,900
17
4,001 bp
971,000 bp
972,000 bp
973,000 bp
974,000 bp
Gene
Centromere
ORC2 A
ORC2 B
ORC4 A
ORC4 B
Input (ORC2 A)
Input (ORC2 B)
Input (ORC4 A)
Input (ORC4 B)
Chromosome 5: 970901 - 974900
ORC2 A
ORC2 B
ORC4 A
ORC4 B
Depth
Region (bp)

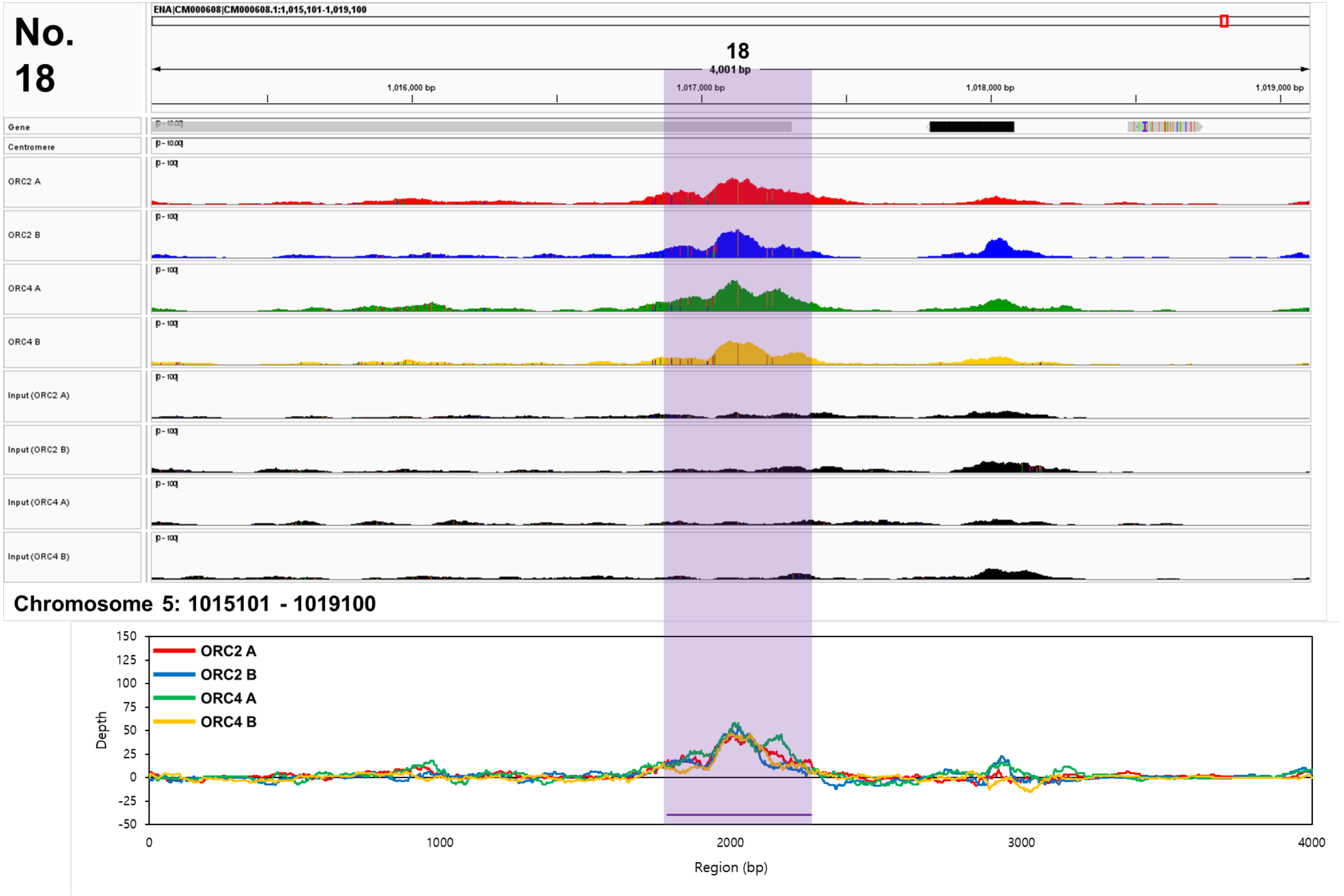
No. 18
ENA|CM000608|CM000608.1:1,015,101-1,019,100
18
4,001 bp
1,016,000 bp
1,017,000 bp
1,018,000 bp
1,019,000 bp
Gene
Centromere
ORC2 A
ORC2 B
ORC4 A
ORC4 B
Input (ORC2 A)
Input (ORC2 B)
Input (ORC4 A)
Input (ORC4 B)
Chromosome 5: 1015101 - 1019100
ORC2 A
ORC2 B
ORC4 A
ORC4 B
Depth
Region (bp)

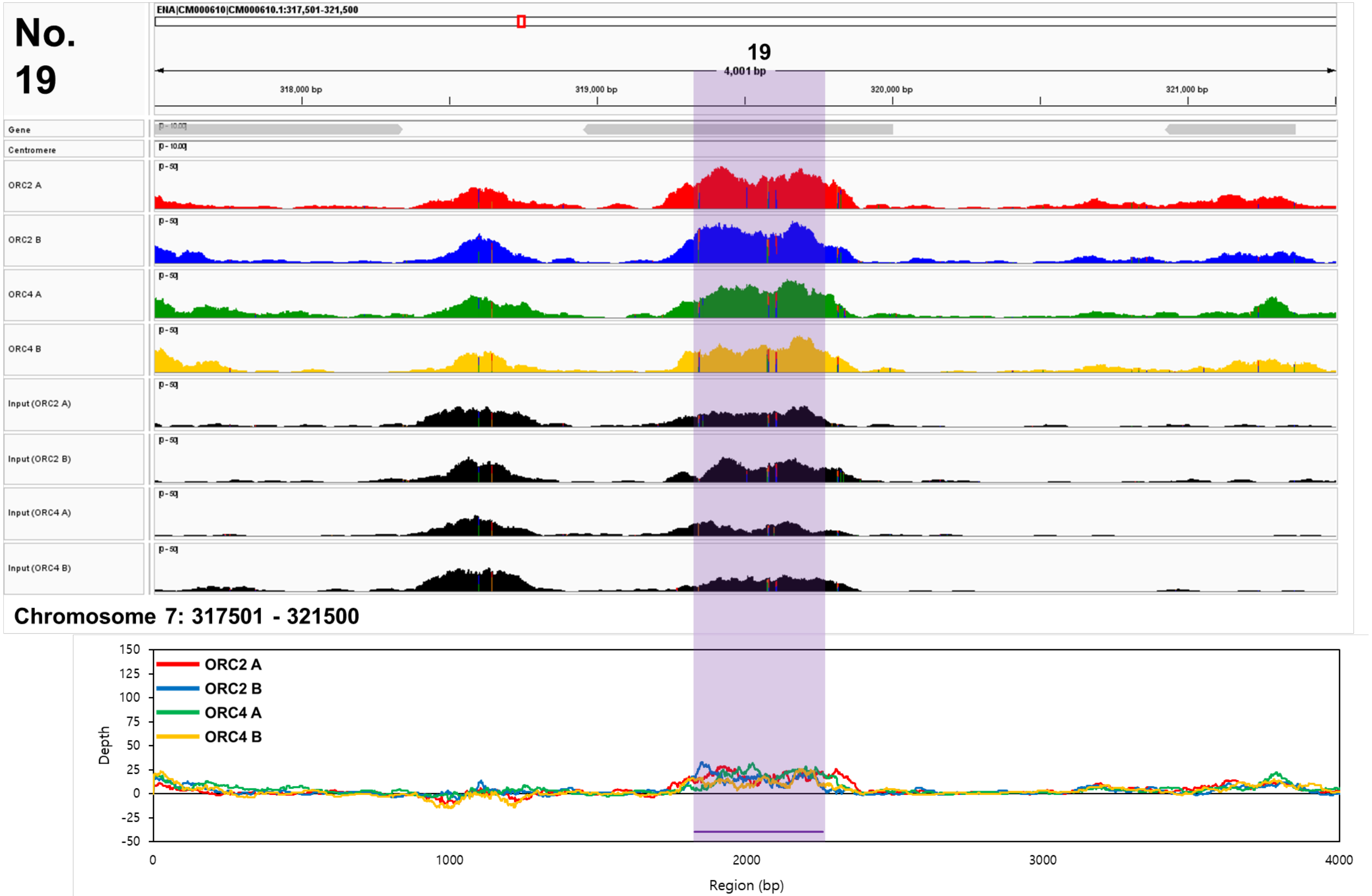
No. 19
ENA|CM000610|CM000610.1:317,501-321,500
19
4,001 bp
318,000 bp
319,000 bp
320,000 bp
321,000 bp
Gene
Centromere
ORC2 A
ORC2 B
ORC4 A
ORC4 B
Input (ORC2 A)
Input (ORC2 B)
Input (ORC4 A)
Input (ORC4 B)
Chromosome 7: 317501 - 321500
ORC2 A
ORC2 B
ORC4 A
ORC4 B
Depth
Region (bp)

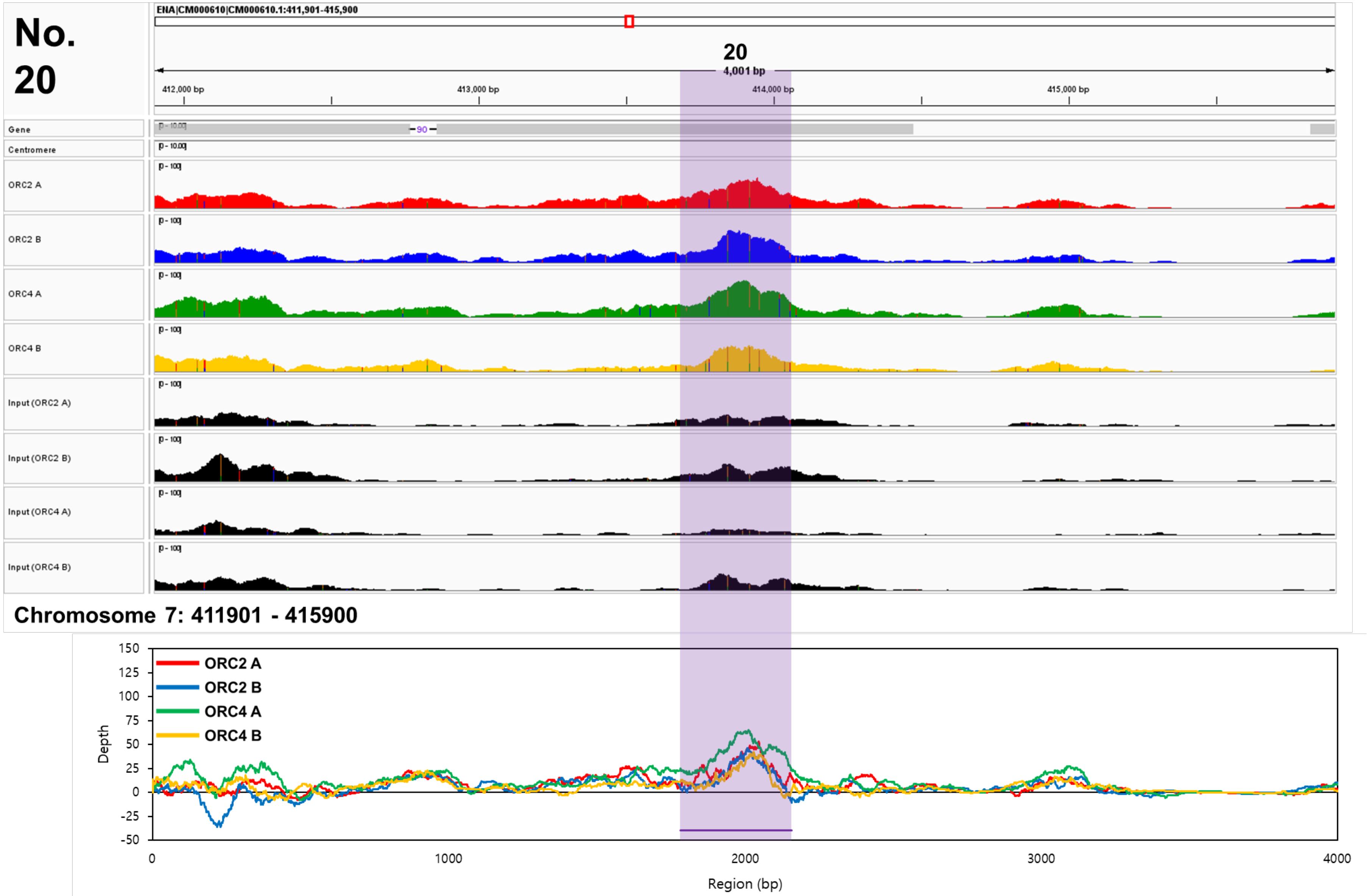
No. 20
ENA|CM000610|CM000610.1:411,901-415,900
20
4,001 bp
412,000 bp
413,000 bp
414,000 bp
415,000 bp
Gene
Centromere
ORC2 A
ORC2 B
ORC4 A
ORC4 B
Input (ORC2 A)
Input (ORC2 B)
Input (ORC4 A)
Input (ORC4 B)
Chromosome 7: 411901 - 415900
ORC2 A
ORC2 B
ORC4 A
ORC4 B
Depth
Region (bp)

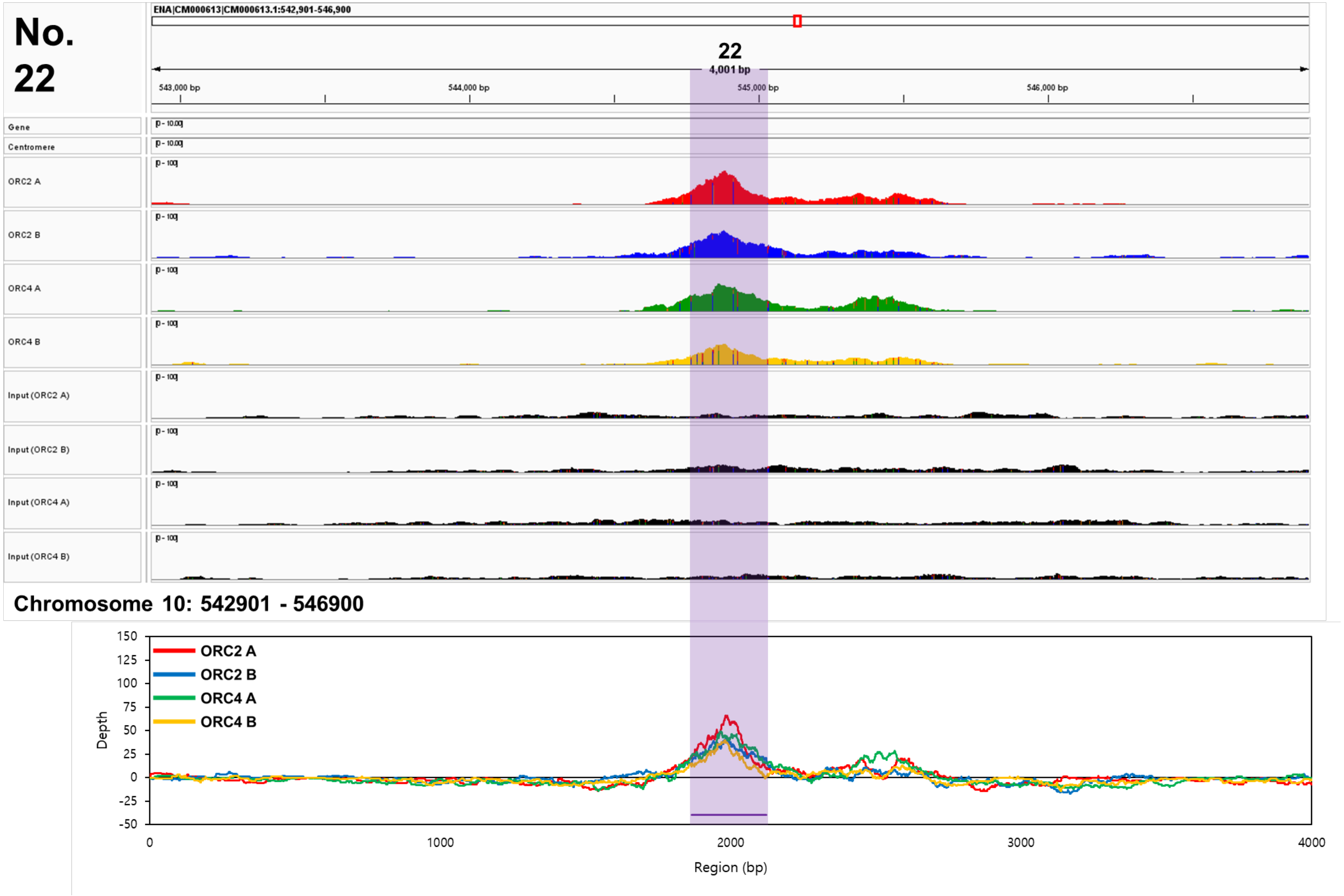

No. 22
ENA|CM000613|CM000613.1:542,901-546,900
22
4,001 bp
543,000 bp
544,000 bp
545,000 bp
546,000 bp
Gene
Centromere
ORC2 A
ORC2 B
ORC4 A
ORC4 B
Input (ORC2 A)
Input (ORC2 B)
Input (ORC4 A)
Input (ORC4 B)
Chromosome 10: 542901 - 546900
ORC2 A
ORC2 B
ORC4 A
ORC4 B
Depth
Region (bp)

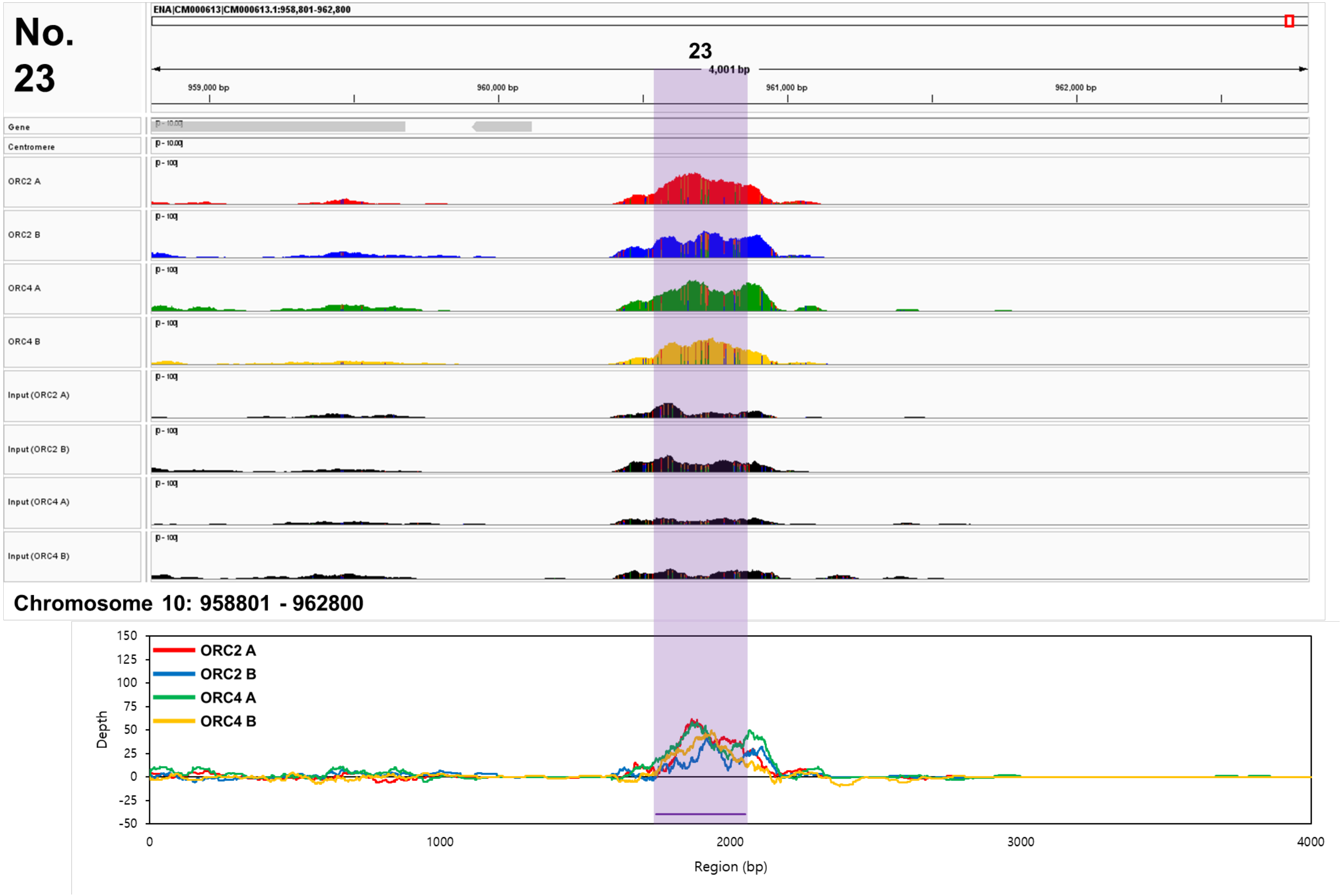
No. 23
ENA|CM000613|CM000613.1:958,801-962,800
23
4,001 bp
959,000 bp
960,000 bp
961,000 bp
962,000 bp
Gene
Centromere
ORC2 A
ORC2 B
ORC4 A
ORC4 B
Input (ORC2 A)
Input (ORC2 B)
Input (ORC4 A)
Input (ORC4 B)
Chromosome 10: 958801 - 962800
ORC2 A
ORC2 B
ORC4 A
ORC4 B
Depth
Region (bp)

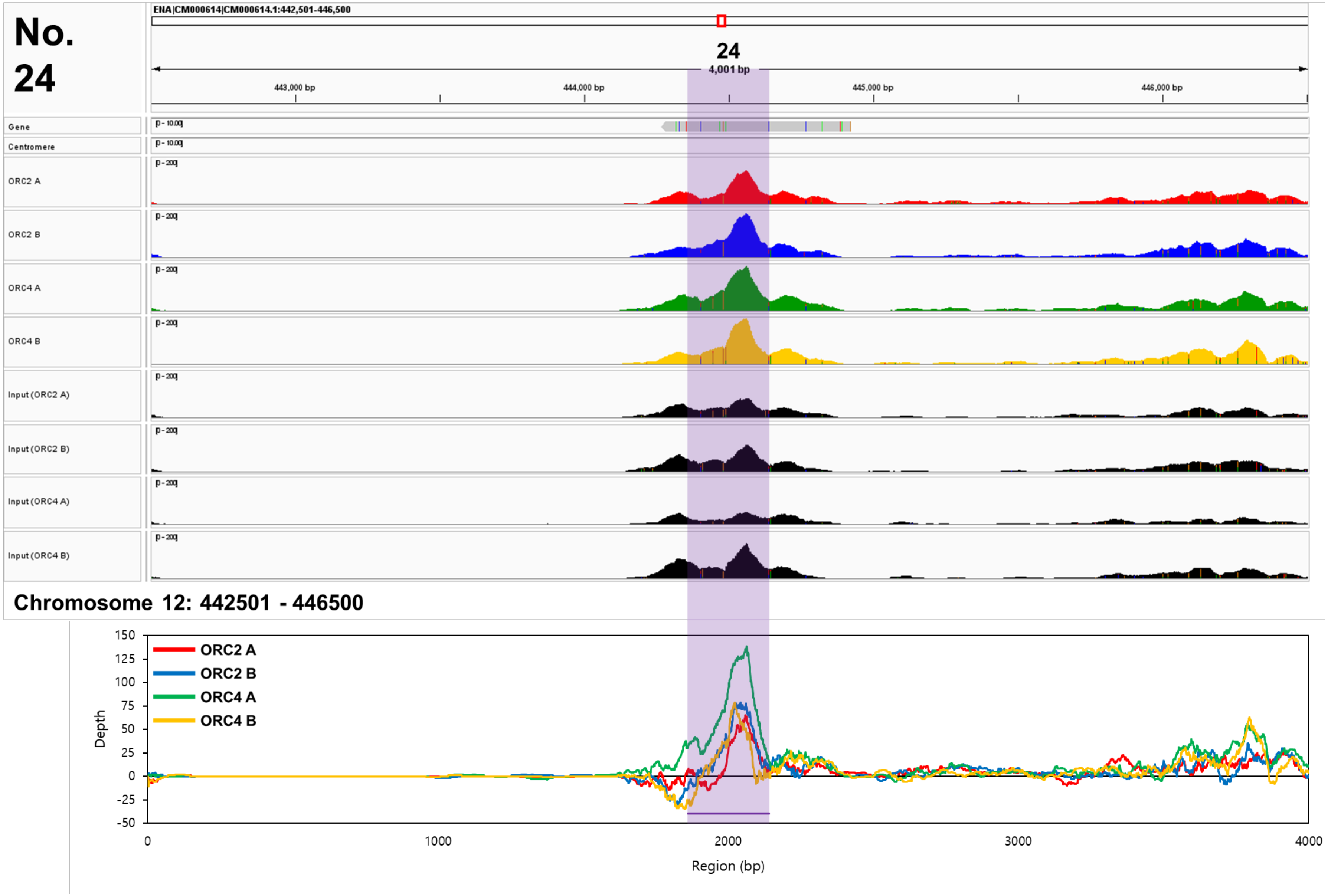

No. 24
ENA|CM000614|CM000614.1:442,501-446,500
24
4,001 bp
443,000 bp
444,000 bp
445,000 bp
446,000 bp
Gene
Centromere
ORC2 A
ORC2 B
ORC4 A
ORC4 B
Input (ORC2 A)
Input (ORC2 B)
Input (ORC4 A)
Input (ORC4 B)
Chromosome 12: 442501 - 446500
ORC2 A
ORC2 B
ORC4 A
ORC4 B
Depth
Region (bp)

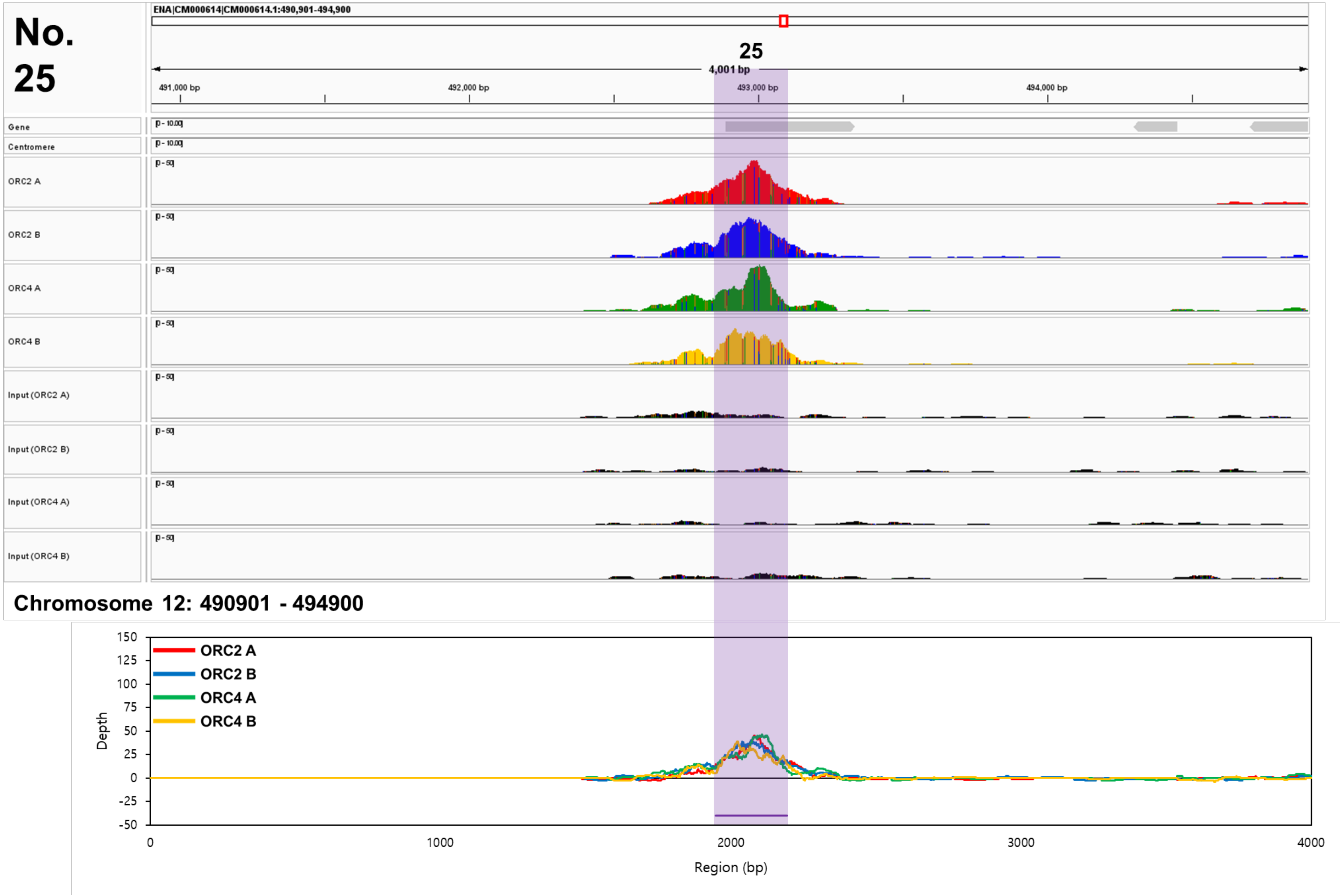
No. 25
ENA|CM000614|CM000614.1:490,901-494,900
25
4,001 bp
491,000 bp
492,000 bp
493,000 bp
494,000 bp
Gene
Centromere
ORC2 A
ORC2 B
ORC4 A
ORC4 B
Input (ORC2 A)
Input (ORC2 B)
Input (ORC4 A)
Input (ORC4 B)
Chromosome 12: 490901 - 494900
ORC2 A
ORC2 B
ORC4 A
ORC4 B
Depth
Region (bp)

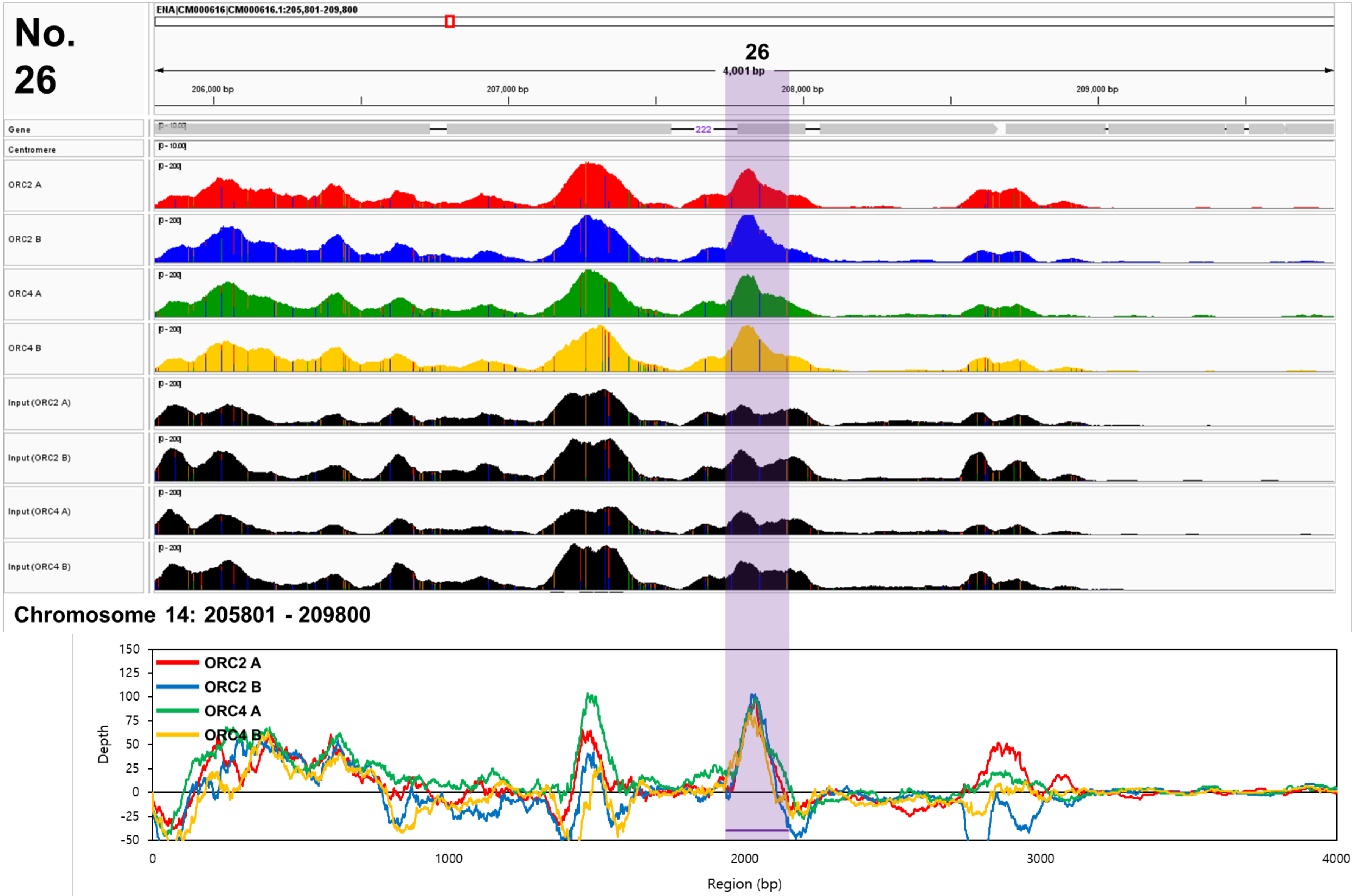
No. 26
ENA|CM000616|CM000616.1:205,801-209,800
26
4,001 bp
206,000 bp
207,000 bp
208,000 bp
209,000 bp
Gene
Centromere
ORC2 A
ORC2 B
ORC4 A
ORC4 B
Input (ORC2 A)
Input (ORC2 B)
Input (ORC4 A)
Input (ORC4 B)
222
Chromosome 14: 205801 - 209800
ORC2 A
ORC2 B
ORC4 A
ORC4 B
Depth
Region (bp)
150
125
100
75
50
25
0
-25
-50
0
1000
2000
3000
4000

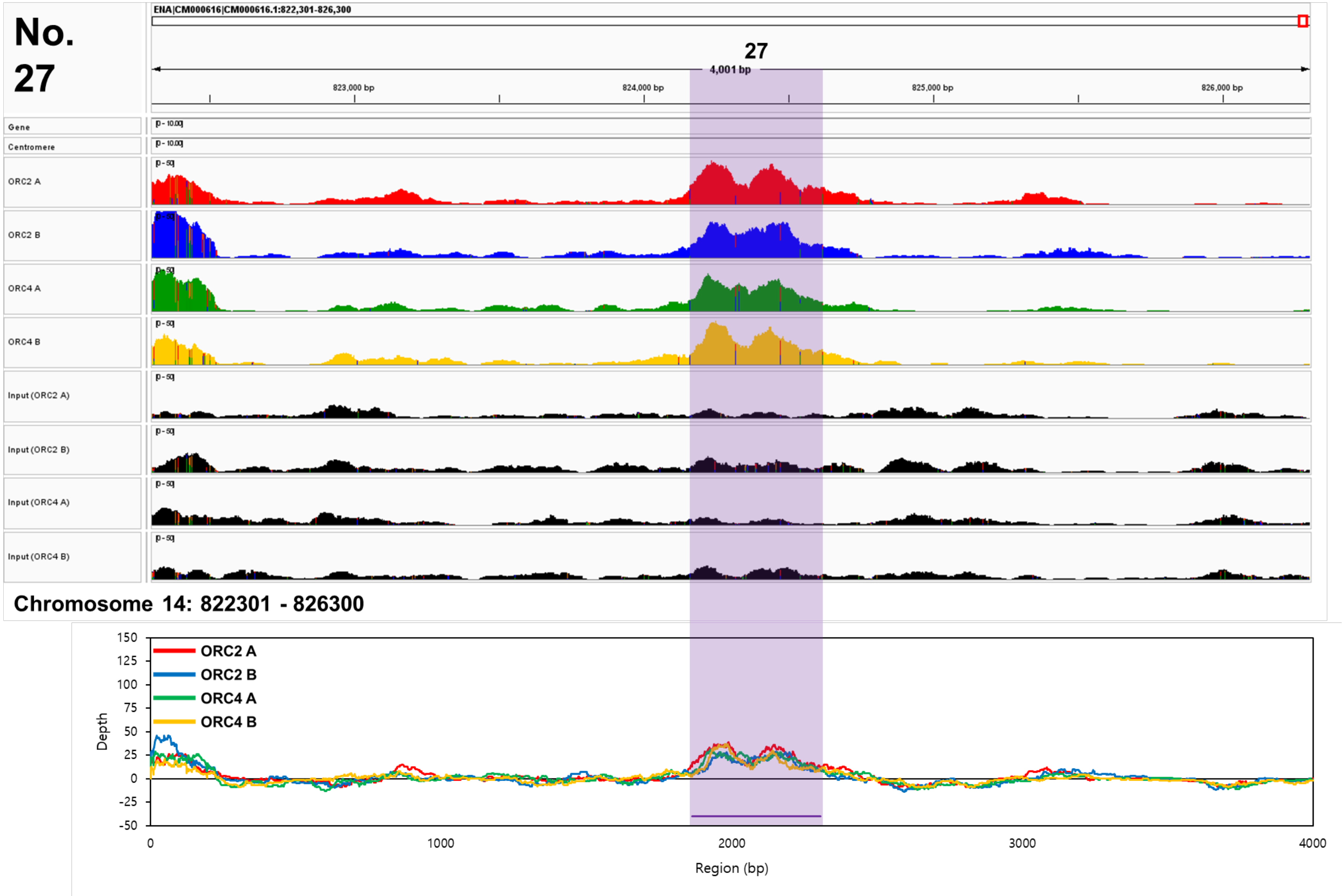
No.
27
ENA|CM000616|CM000616.1:822,301-826,300
27
4,001 bp
823,000 bp
824,000 bp
825,000 bp
826,000 bp
Gene
Centromere
ORC2 A
ORC2 B
ORC4 A
ORC4 B
Input (ORC2 A)
Input (ORC2 B)
Input (ORC4 A)
Input (ORC4 B)
Chromosome 14: 822301 - 826300
ORC2 A
ORC2 B
ORC4 A
ORC4 B
Depth
Region (bp)

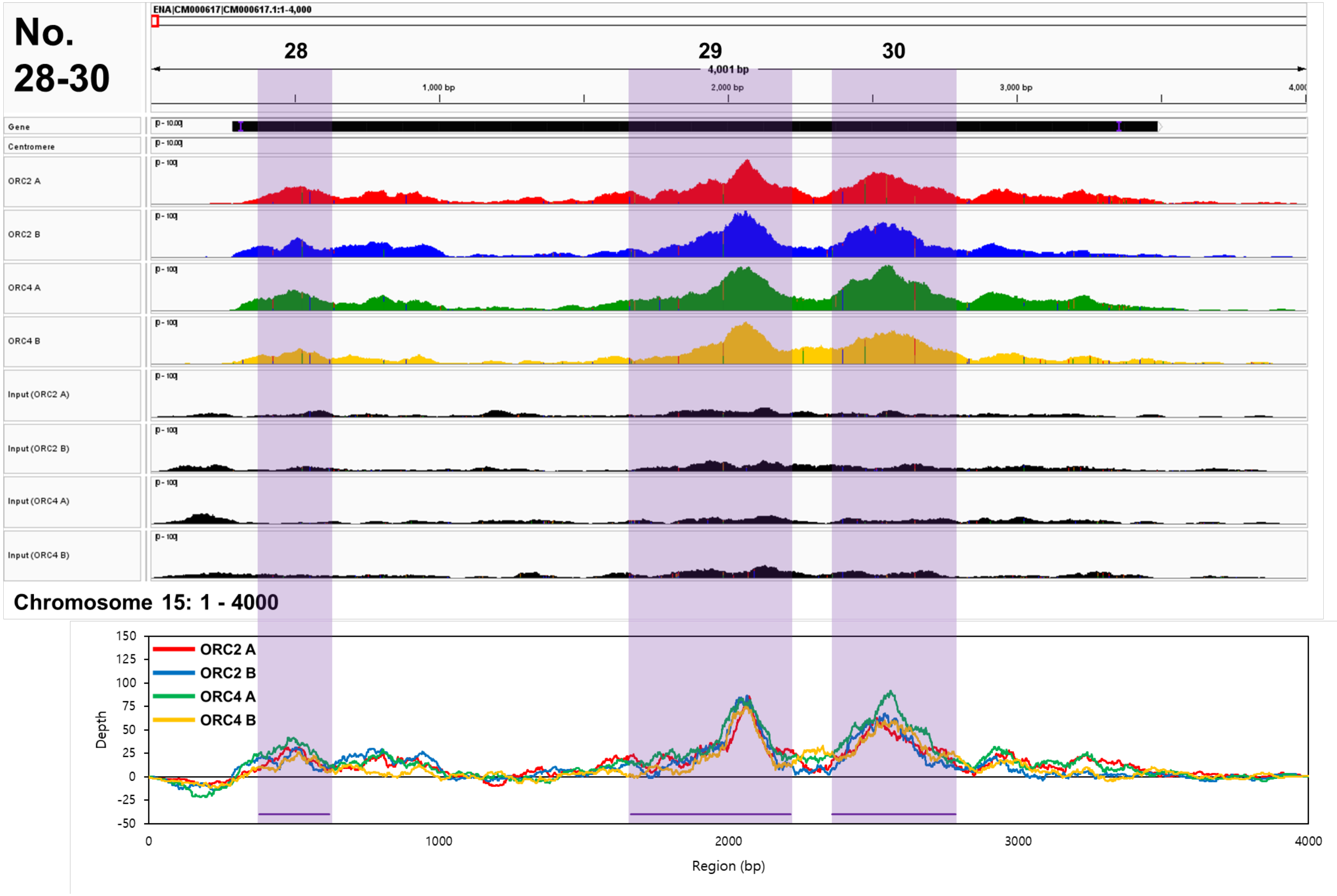
No.
28-30
ENA|CM000617|CM000617.1:1-4,000
28
29
30
4,001 bp
1,000 bp
2,000 bp
3,000 bp
Gene
Centromere
ORC2 A
ORC2 B
ORC4 A
ORC4 B
Input (ORC2 A)
Input (ORC2 B)
Input (ORC4 A)
Input (ORC4 B)
Chromosome 15: 1 - 4000
ORC2 A
ORC2 B
ORC4 A
ORC4 B
Depth
Region (bp)
150
125
100
75
50
25
0
-25
-50
0
1000
2000
3000
4000

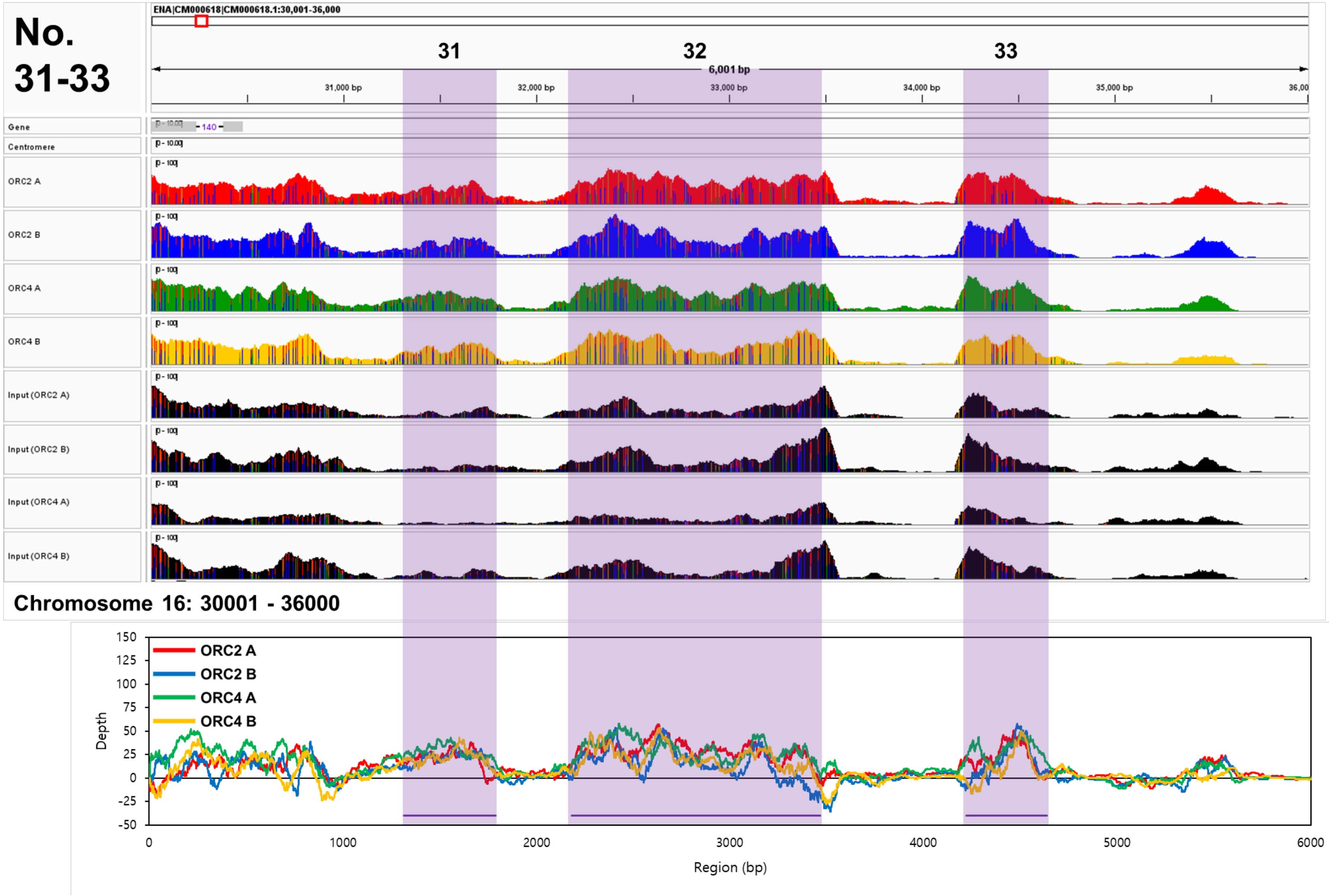
No. 31-33
ENA|CM000618|CM000618.1:30,001-36,000
31
32
33
6,001 bp
31,000 bp
32,000 bp
33,000 bp
34,000 bp
35,000 bp
Gene
Centromere
ORC2 A
ORC2 B
ORC4 A
ORC4 B
Input (ORC2 A)
Input (ORC2 B)
Input (ORC4 A)
Input (ORC4 B)
Chromosome 16: 30001 - 36000
ORC2 A
ORC2 B
ORC4 A
ORC4 B
Depth
150
125
100
75
50
25
0
-25
-50
0
1000
2000
3000
4000
5000
6000
Region (bp)

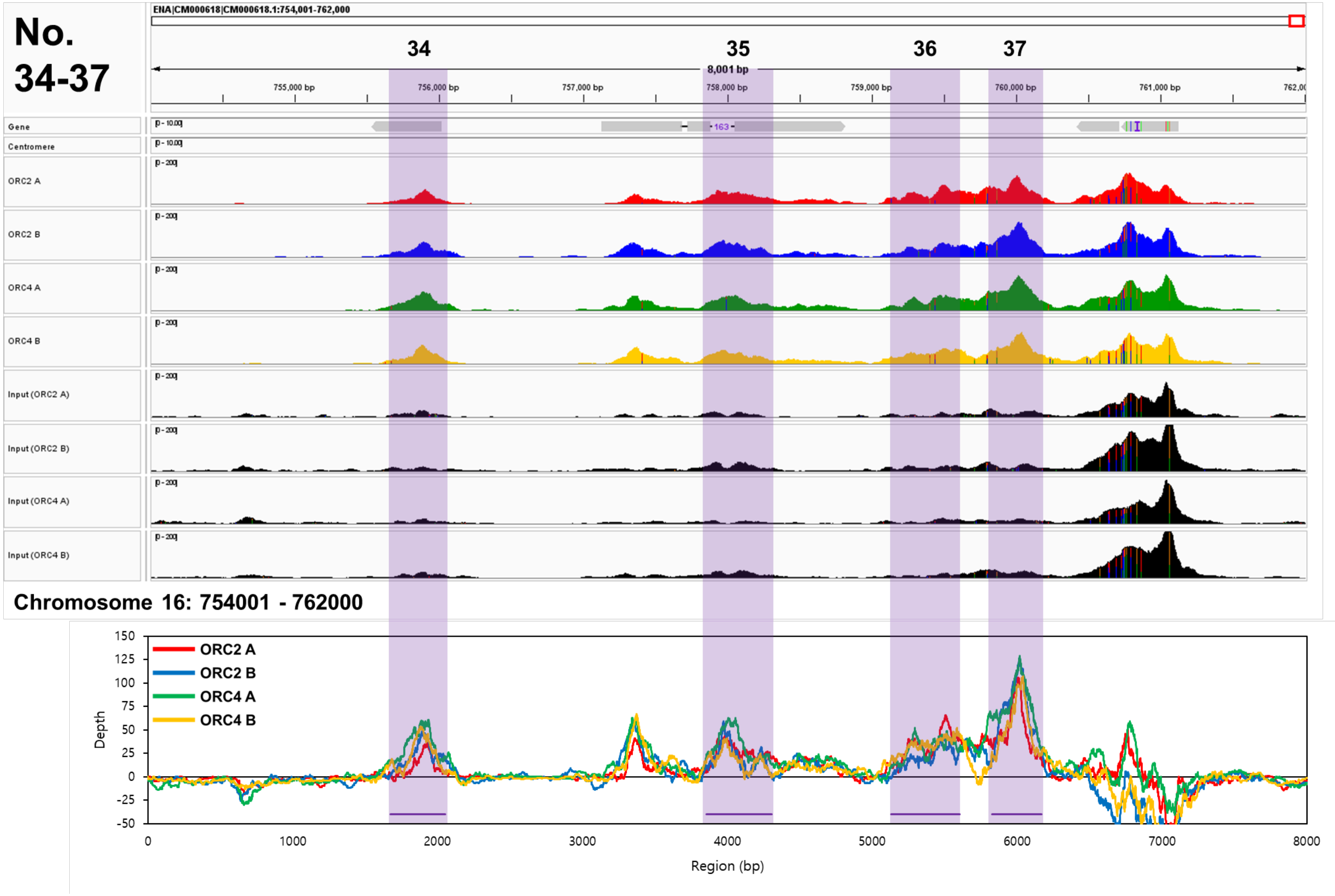

No.
34-37
ENA|CM000618|CM000618.1:754,001-762,000
34
35
36
37
8,001 bp
755,000 bp
756,000 bp
757,000 bp
758,000 bp
759,000 bp
760,000 bp
761,000 bp
Gene
Centromere
ORC2 A
ORC2 B
ORC4 A
ORC4 B
Input (ORC2 A)
Input (ORC2 B)
Input (ORC4 A)
Input (ORC4 B)
Chromosome 16: 754001 - 762000
ORC2 A
ORC2 B
ORC4 A
ORC4 B
Depth
Region (bp)

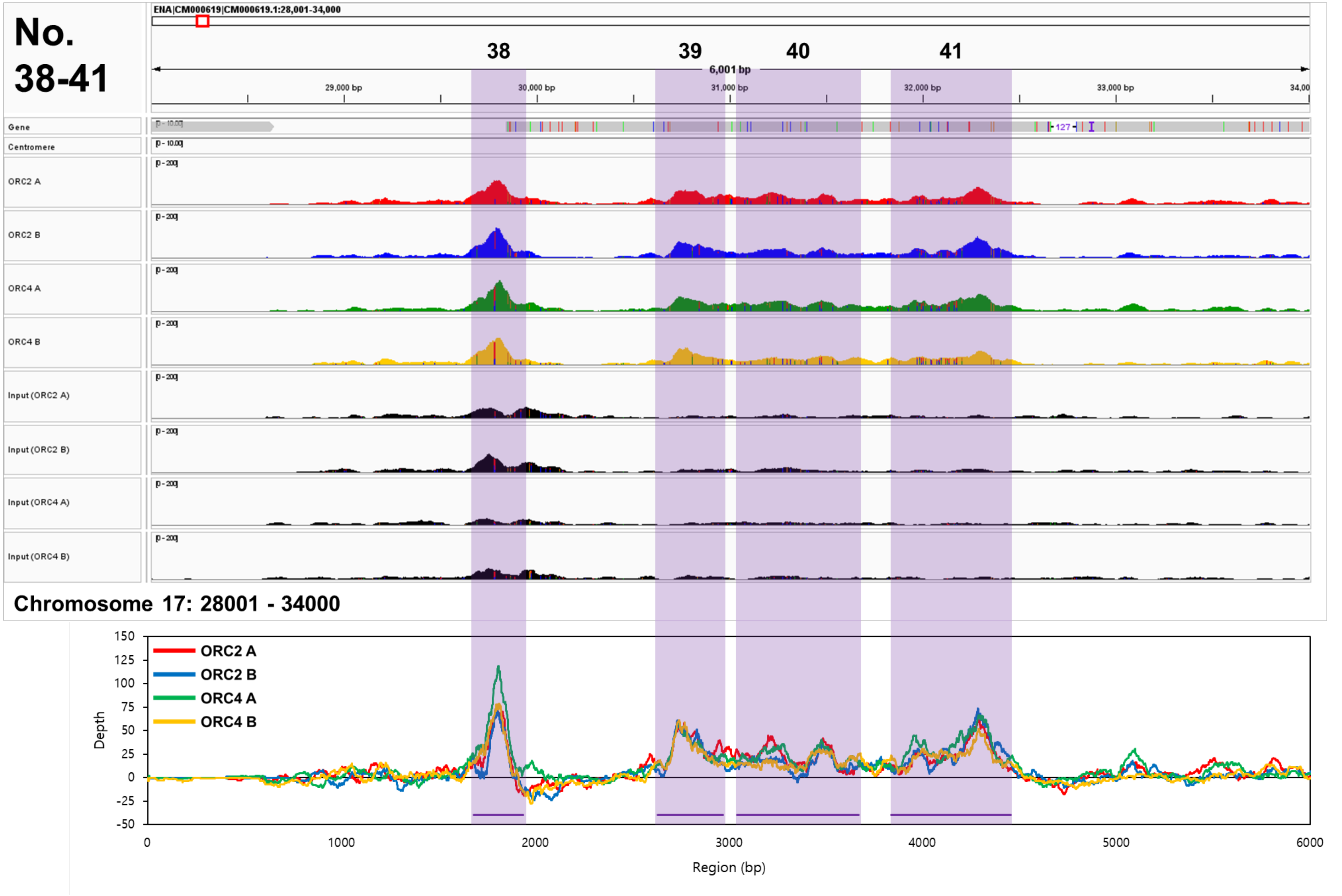
No. 38-41
ENA|CM000619|CM000619.1:28,001-34,000
38
39
40
41
6,001 bp
29,000 bp
30,000 bp
31,000 bp
32,000 bp
33,000 bp
Gene
Centromere
ORC2 A
ORC2 B
ORC4 A
ORC4 B
Input (ORC2 A)
Input (ORC2 B)
Input (ORC4 A)
Input (ORC4 B)
Chromosome 17: 28001 - 34000
ORC2 A
ORC2 B
ORC4 A
ORC4 B
Depth
Region (bp)

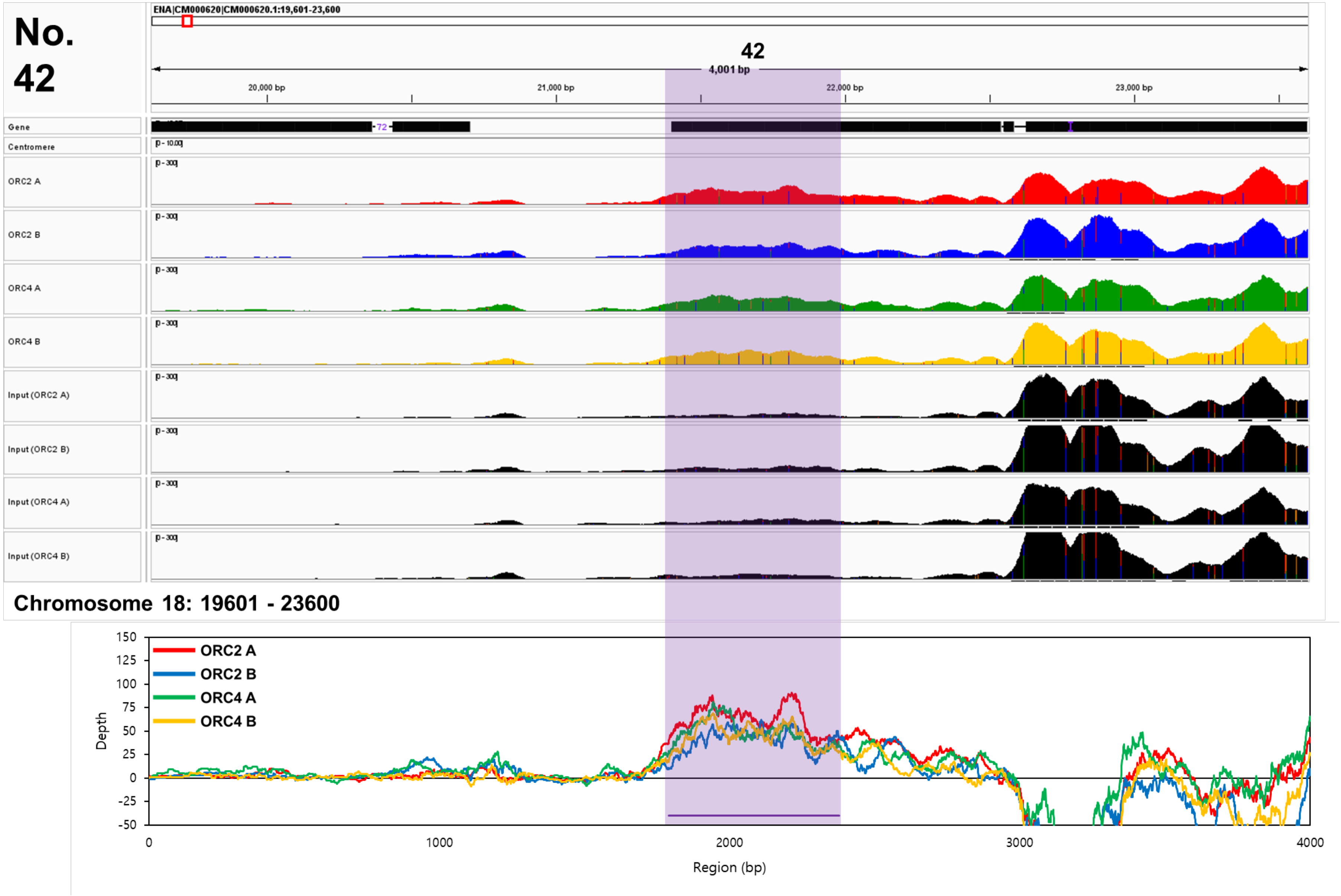
No.
42
ENA|CM000620|CM000620.1:19,601-23,600
42
4,001 bp
20,000 bp
21,000 bp
22,000 bp
23,000 bp
Gene
Centromere
ORC2 A
ORC2 B
ORC4 A
ORC4 B
Input (ORC2 A)
Input (ORC2 B)
Input (ORC4 A)
Input (ORC4 B)
Chromosome 18: 19601 - 23600
ORC2 A
ORC2 B
ORC4 A
ORC4 B
Depth
Region (bp)

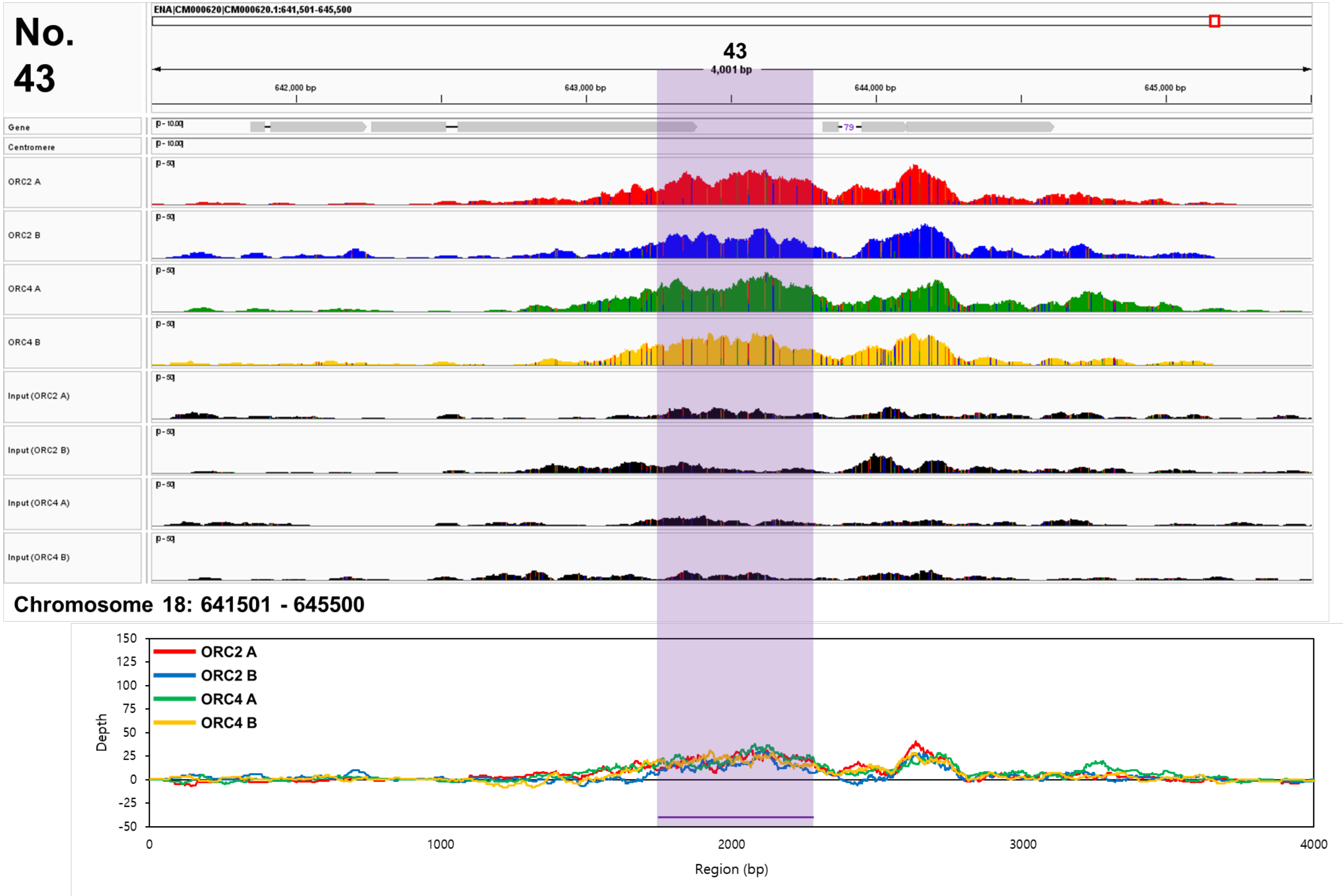
No. 43
ENA|CM000620|CM000620.1:641,501-645,500
43
4,001 bp
642,000 bp
643,000 bp
644,000 bp
645,000 bp
Gene
Centromere
ORC2 A
ORC2 B
ORC4 A
ORC4 B
Input (ORC2 A)
Input (ORC2 B)
Input (ORC4 A)
Input (ORC4 B)
Chromosome 18: 641501 - 645500
ORC2 A
ORC2 B
ORC4 A
ORC4 B
Depth
Region (bp)

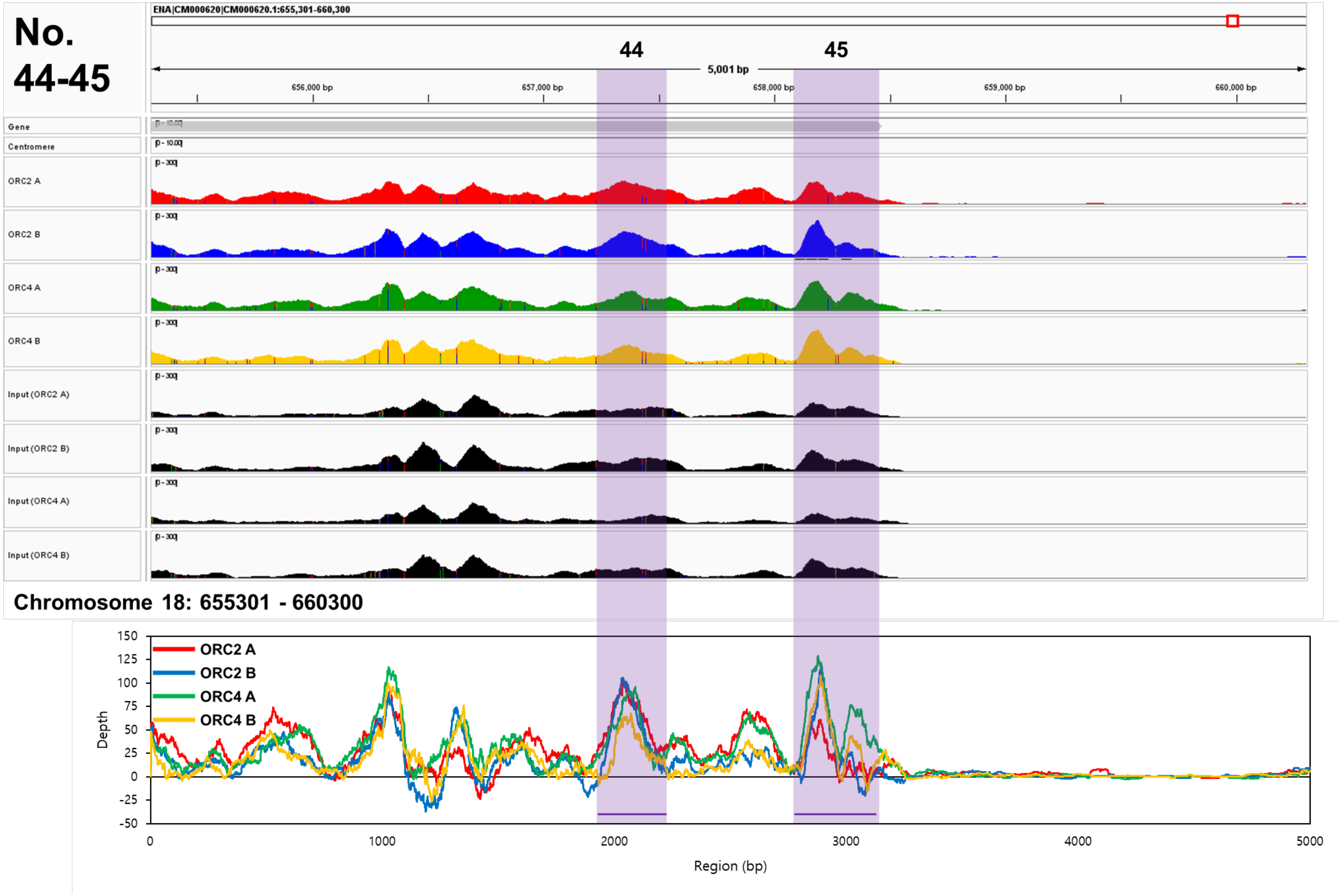

No. 44-45
ENA|CM000620|CM000620.1:655,301-660,300
44
45
5,001 bp
656,000 bp
657,000 bp
658,000 bp
659,000 bp
660,000 bp
Gene
Centromere
ORC2 A
ORC2 B
ORC4 A
ORC4 B
Input (ORC2 A)
Input (ORC2 B)
Input (ORC4 A)
Input (ORC4 B)
Chromosome 18: 655301 - 660300
ORC2 A
ORC2 B
ORC4 A
ORC4 B
Depth
Region (bp)

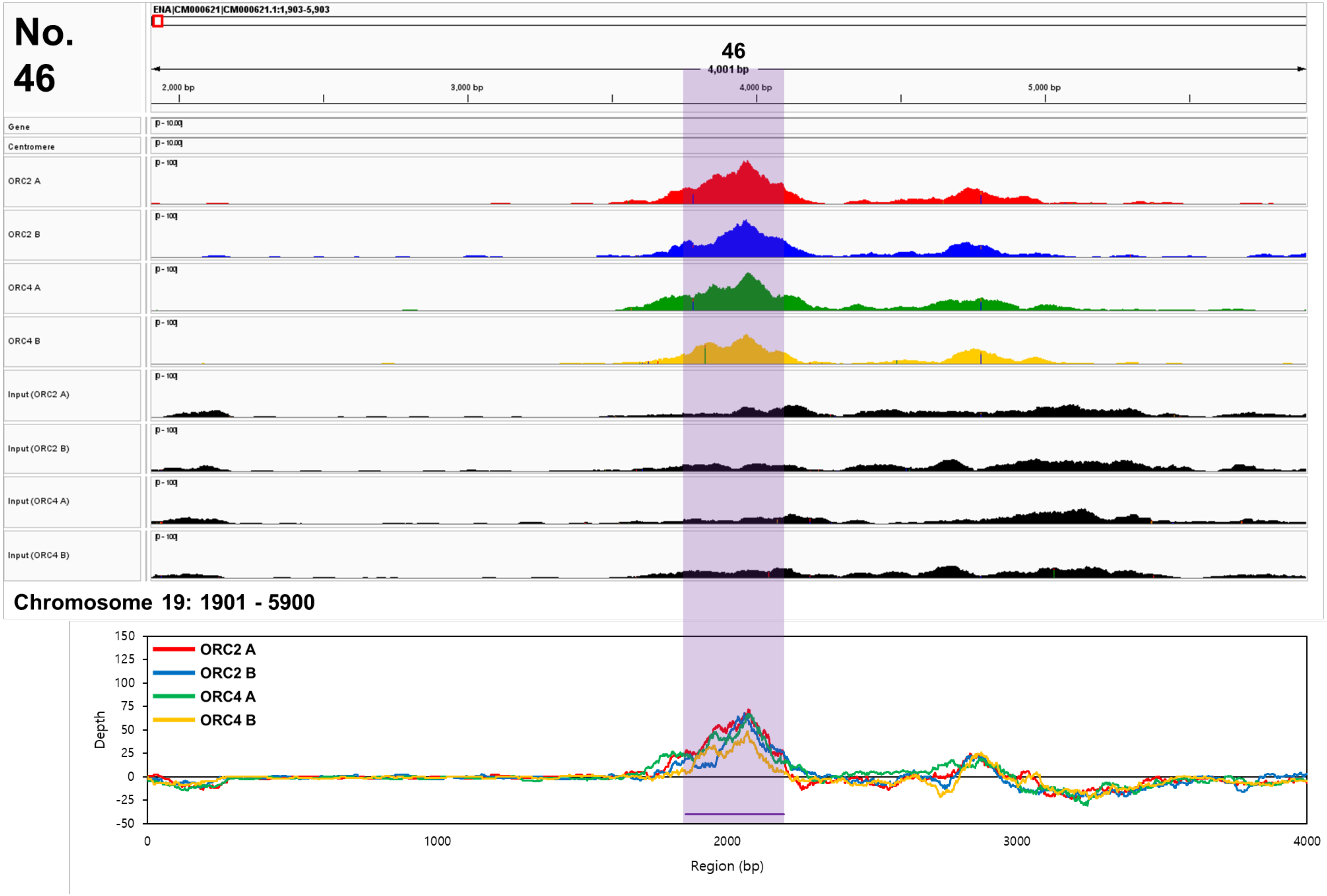
No. 46
ENA|CM000621|CM000621.1:1,903-5,903
46
4,001 bp
2,000 bp
3,000 bp
4,000 bp
5,000 bp
Gene
Centromere
ORC2 A
ORC2 B
ORC4 A
ORC4 B
Input (ORC2 A)
Input (ORC2 B)
Input (ORC4 A)
Input (ORC4 B)
Chromosome 19: 1901 - 5900
ORC2 A
ORC2 B
ORC4 A
ORC4 B
Depth
Region (bp)

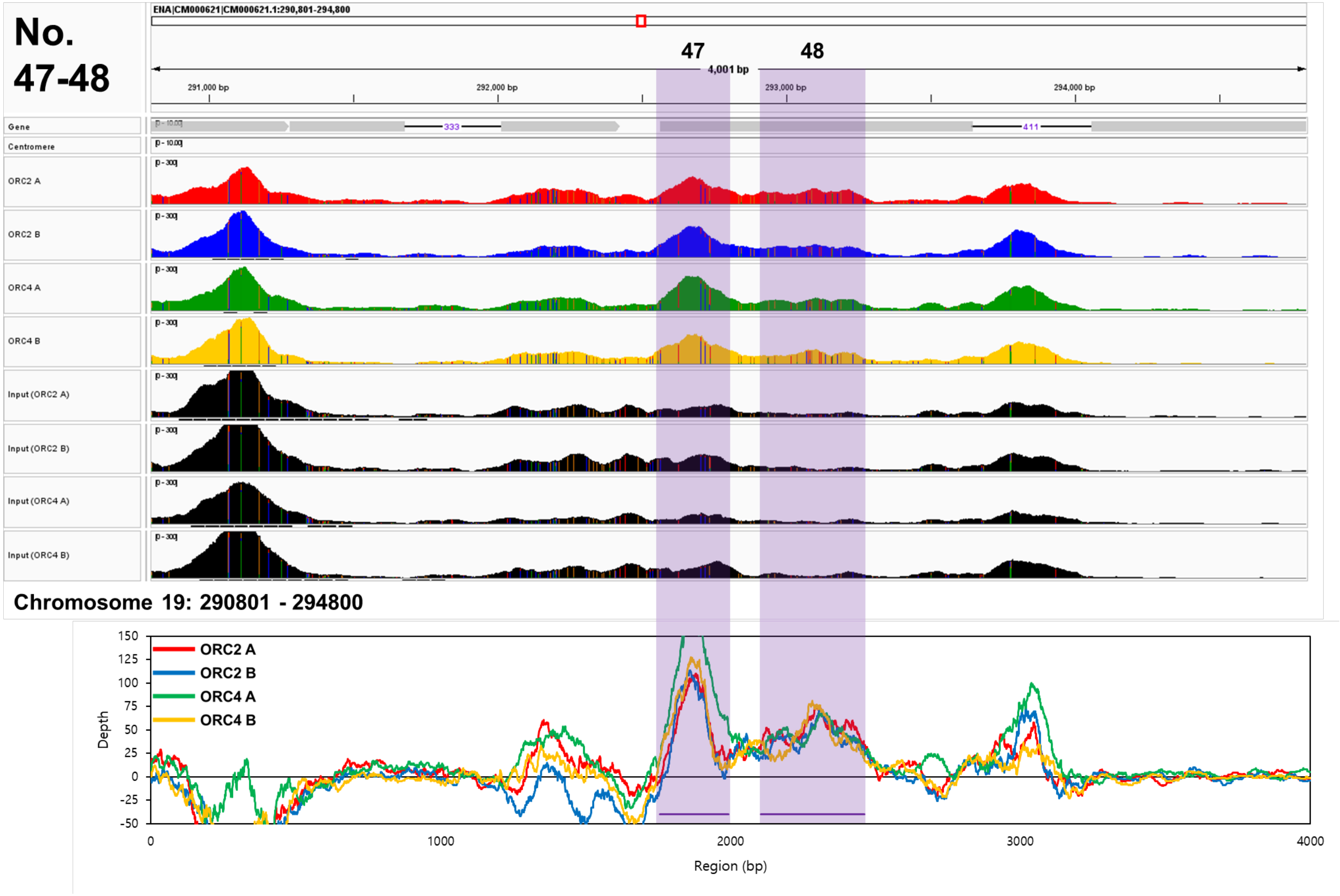
No.
47-48
ENA|CM000621|CM000621.1:290,801-294,800
47
48
4,001 bp
291,000 bp
292,000 bp
293,000 bp
294,000 bp
Gene
Centromere
333
411
ORC2 A
ORC2 B
ORC4 A
ORC4 B
Input (ORC2 A)
Input (ORC2 B)
Input (ORC4 A)
Input (ORC4 B)
Chromosome 19: 290801 - 294800
Depth
Region (bp)
150
125
100
75
50
25
0
-25
-50
0
1000
2000
3000
4000

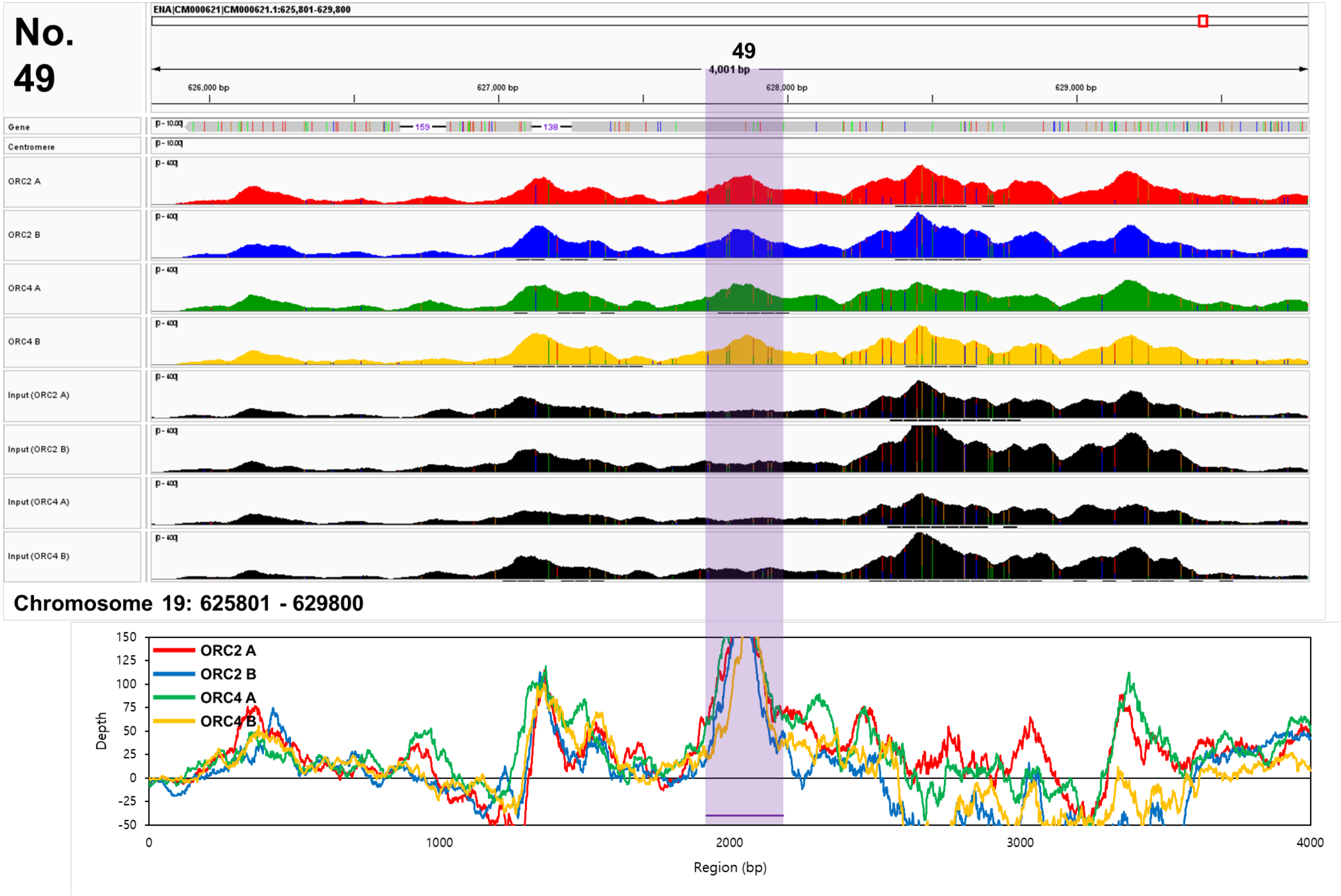
No. 49
ENA|CM000621|CM000621.1:625,801-629,800
49
4,001 bp
626,000 bp
627,000 bp
628,000 bp
629,000 bp
Gene
159
138
Centromere
ORC2 A
ORC2 B
ORC4 A
ORC4 B
Input (ORC2 A)
Input (ORC2 B)
Input (ORC4 A)
Input (ORC4 B)
Chromosome 19: 625801 - 629800
ORC2 A
ORC2 B
ORC4 A
ORC4 B
Depth
Region (bp)
150
125
100
75
50
25
0
-25
-50
0
1000
2000
3000
4000

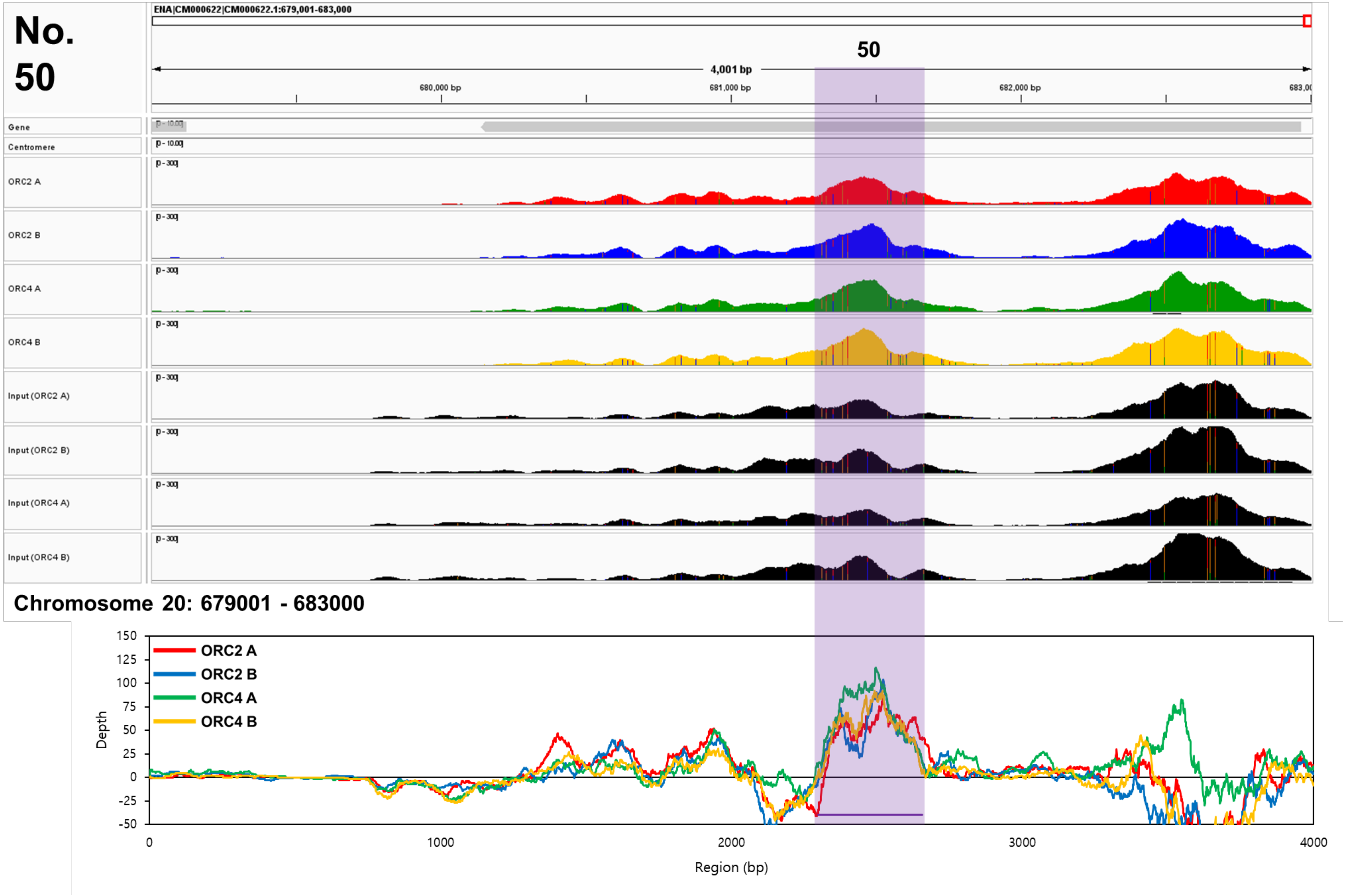

No. 50
ENA|CM000622|CM000622.1:679,001-683,000
50
4,001 bp
680,000 bp
681,000 bp
682,000 bp
Gene
Centromere
ORC2 A
ORC2 B
ORC4 A
ORC4 B
Input (ORC2 A)
Input (ORC2 B)
Input (ORC4 A)
Input (ORC4 B)
Chromosome 20: 679001 - 683000
ORC2 A
ORC2 B
ORC4 A
ORC4 B
Depth
Region (bp)

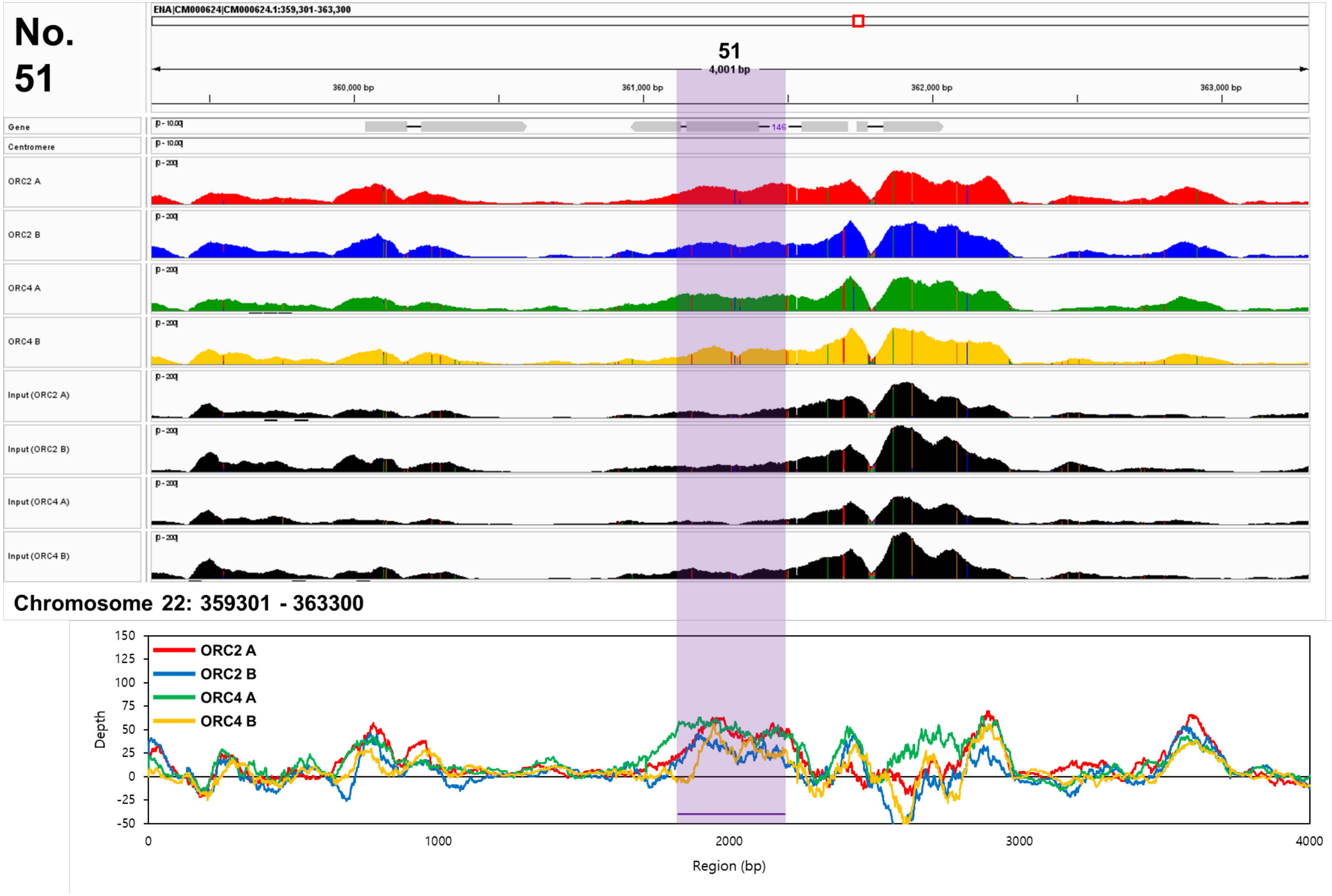
No. 51
ENA|CM000624|CM000624.1:359,301-363,300
51
4,001 bp
360,000 bp
361,000 bp
362,000 bp
363,000 bp
Gene
Centromere
ORC2 A
ORC2 B
ORC4 A
ORC4 B
Input (ORC2 A)
Input (ORC2 B)
Input (ORC4 A)
Input (ORC4 B)
Chromosome 22: 359301 - 363300
ORC2 A
ORC2 B
ORC4 A
ORC4 B
Depth
Region (bp)

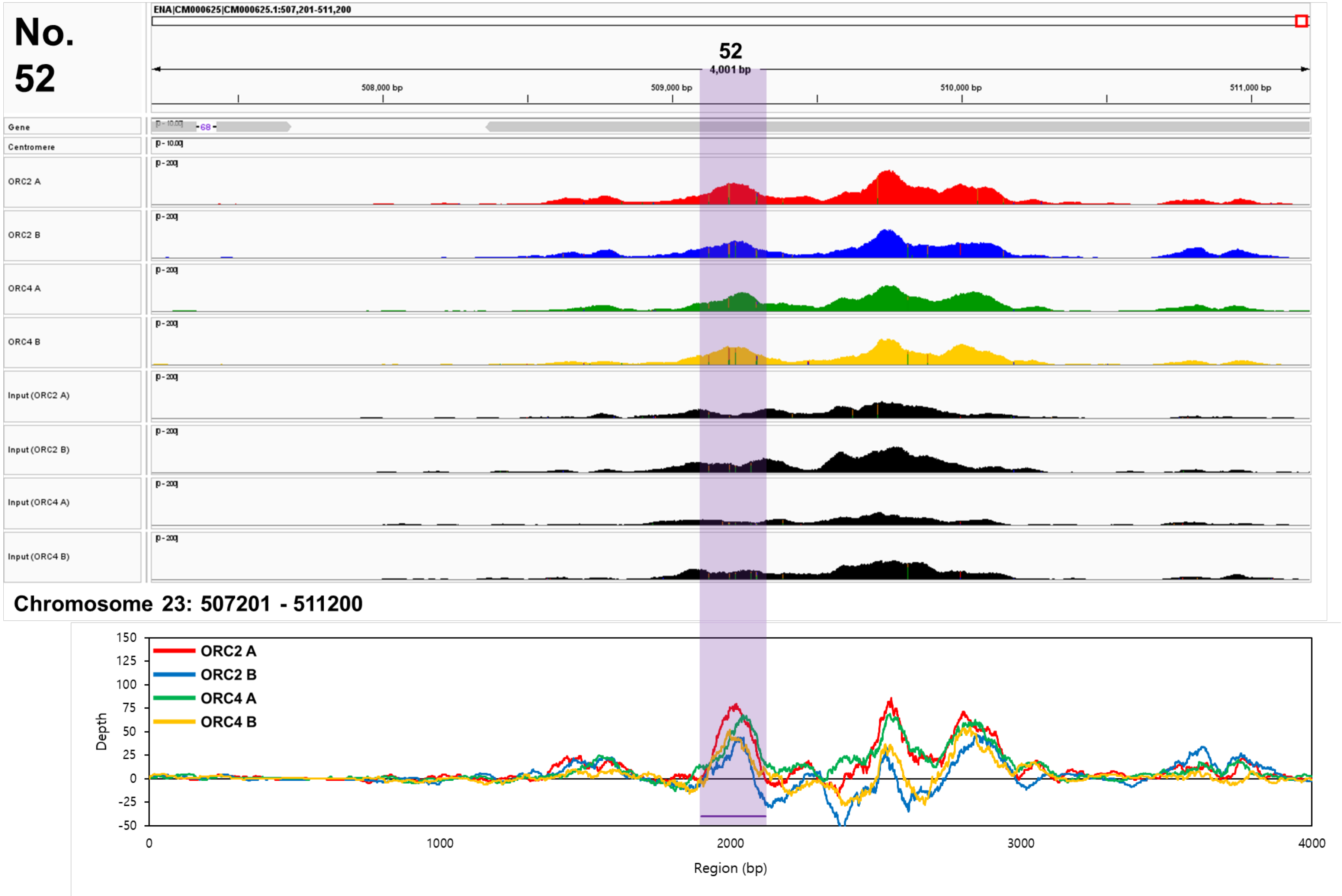
No. 52
ENA|CM000625|CM000625.1:507,201-511,200
52
4,001 bp
508,000 bp
509,000 bp
510,000 bp
511,000 bp
Gene
Centromere
ORC2 A
ORC2 B
ORC4 A
ORC4 B
Input (ORC2 A)
Input (ORC2 B)
Input (ORC4 A)
Input (ORC4 B)
Chromosome 23: 507201 - 511200
ORC2 A
ORC2 B
ORC4 A
ORC4 B
Depth
Region (bp)

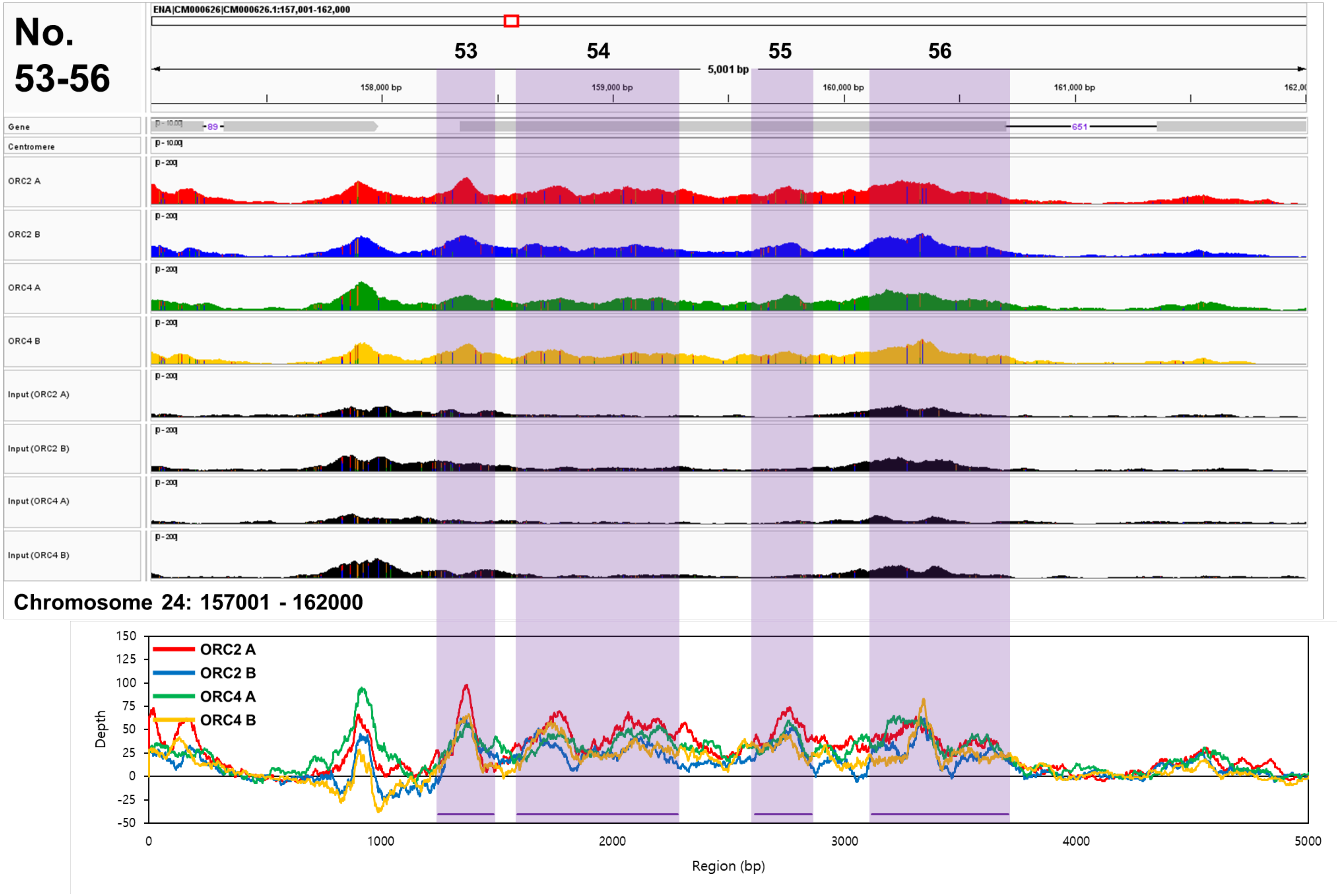
No.
53-56
ENA|CM000626|CM000626.1:157,001-162,000
53
54
55
56
5,001 bp
158,000 bp
159,000 bp
160,000 bp
161,000 bp
Gene
Centromere
ORC2 A
ORC2 B
ORC4 A
ORC4 B
Input (ORC2 A)
Input (ORC2 B)
Input (ORC4 A)
Input (ORC4 B)
Chromosome 24: 157001 - 162000
ORC2 A
ORC2 B
ORC4 A
ORC4 B
Depth
Region (bp)

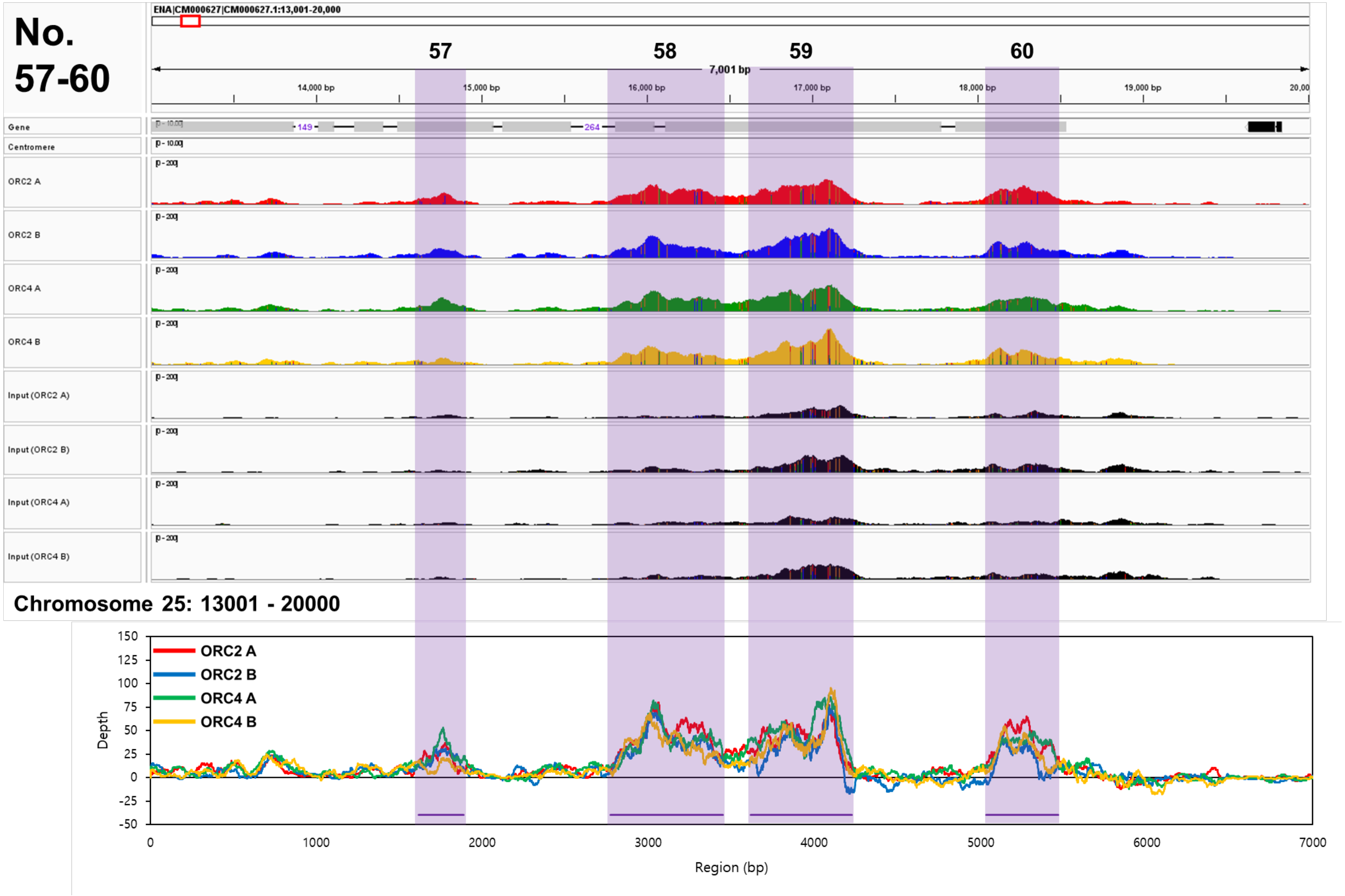
No.
57-60
ENA|CM000627|CM000627.1:13,001-20,000
57
58
59
60
7,001 bp
14,000 bp
15,000 bp
16,000 bp
17,000 bp
18,000 bp
19,000 bp
Gene
Centromere
ORC2 A
ORC2 B
ORC4 A
ORC4 B
Input (ORC2 A)
Input (ORC2 B)
Input (ORC4 A)
Input (ORC4 B)
Chromosome 25: 13001 - 20000
ORC2 A
ORC2 B
ORC4 A
ORC4 B
Depth
Region (bp)

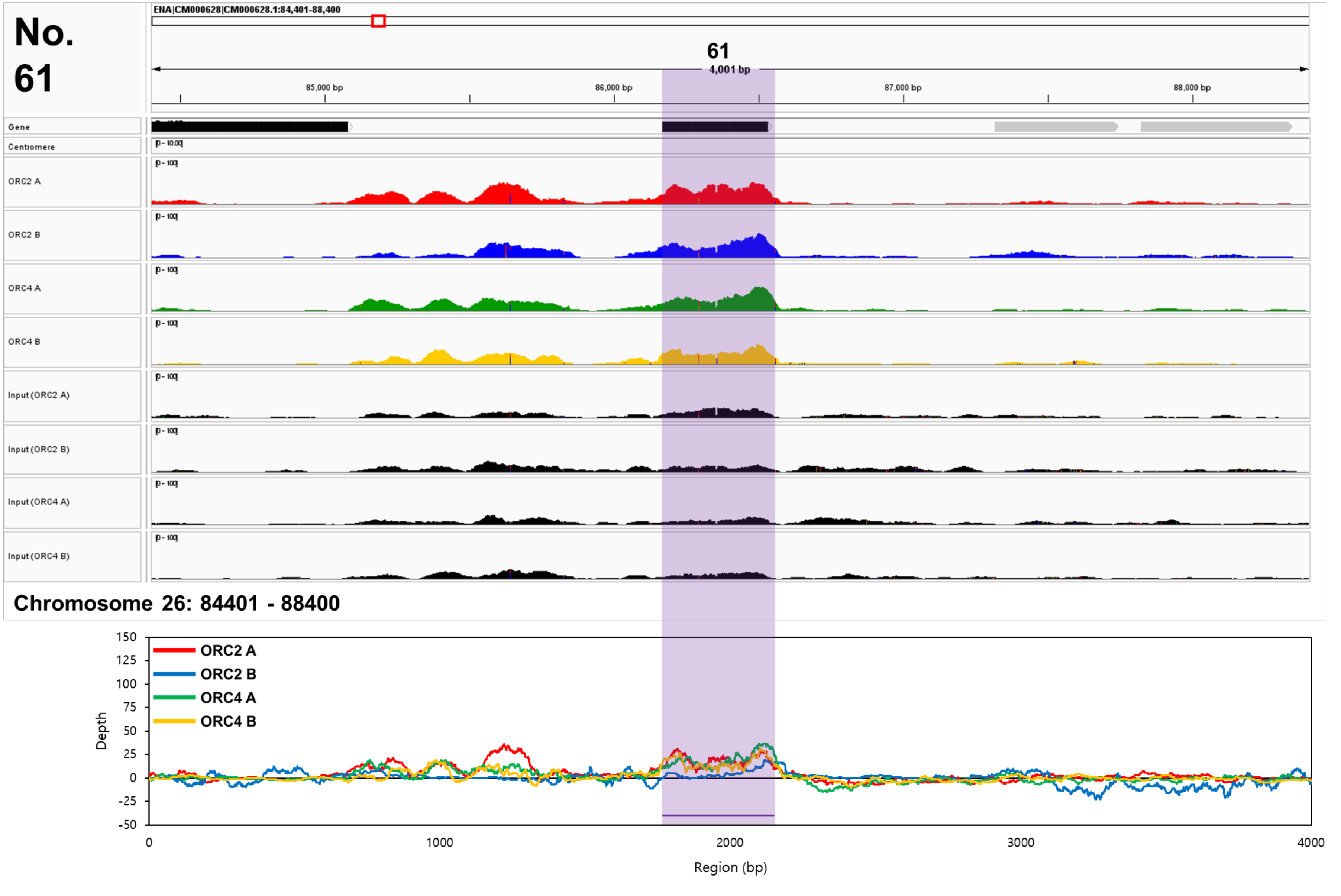
No. 61
ENA|CM000628|CM000628.1:84,401-88,400
61
4,001 bp
85,000 bp
86,000 bp
87,000 bp
88,000 bp
Gene
Centromere
ORC2 A
ORC2 B
ORC4 A
ORC4 B
Input (ORC2 A)
Input (ORC2 B)
Input (ORC4 A)
Input (ORC4 B)
Chromosome 26: 84401 - 88400
ORC2 A
ORC2 B
ORC4 A
ORC4 B
Depth
Region (bp)

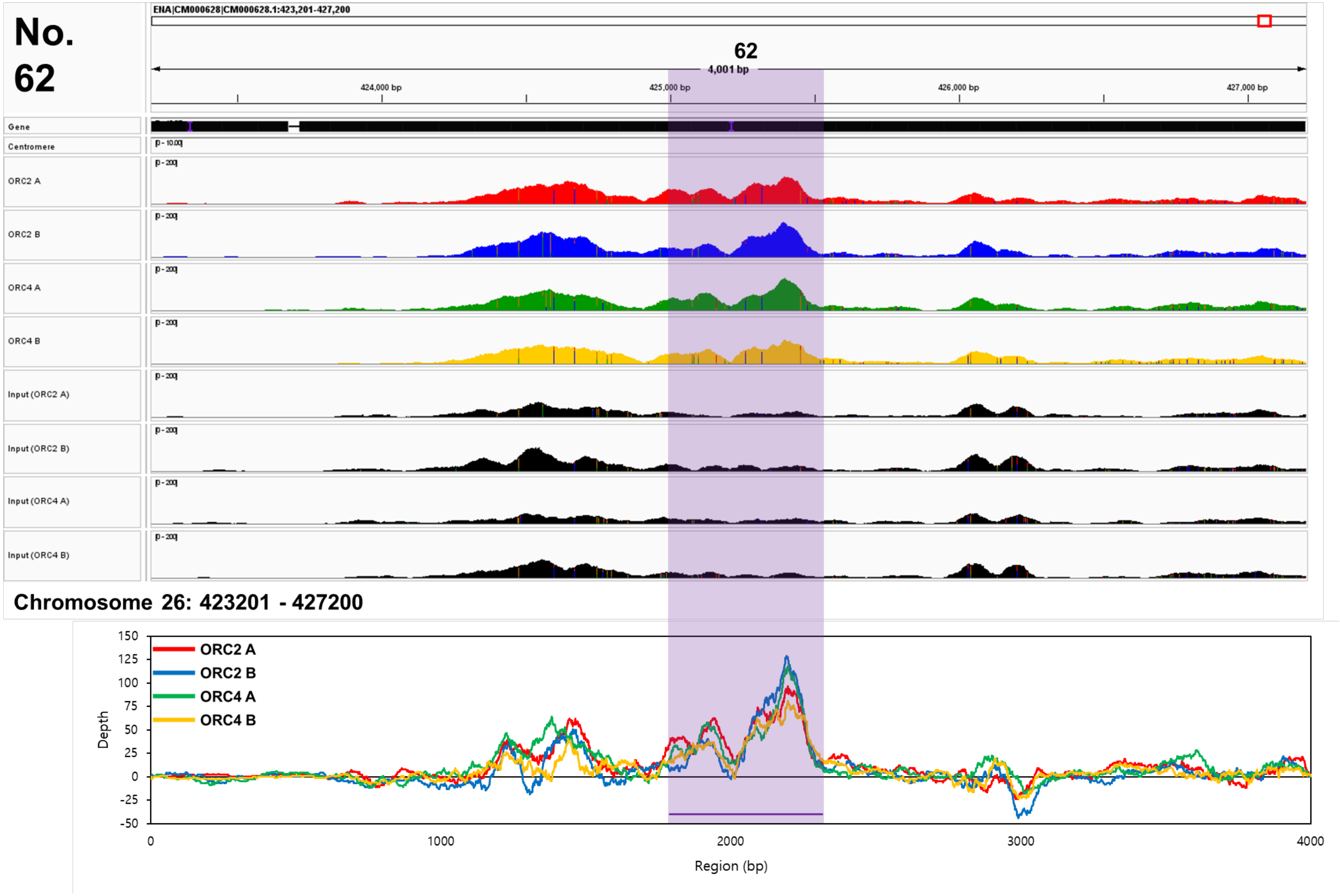
No. 62
ENA|CM000628|CM000628.1:423,201-427,200
62
4,001 bp
424,000 bp
425,000 bp
426,000 bp
427,000 bp
Gene
Centromere
ORC2 A
ORC2 B
ORC4 A
ORC4 B
Input (ORC2 A)
Input (ORC2 B)
Input (ORC4 A)
Input (ORC4 B)
Chromosome 26: 423201 - 427200
ORC2 A
ORC2 B
ORC4 A
ORC4 B
Depth
Region (bp)

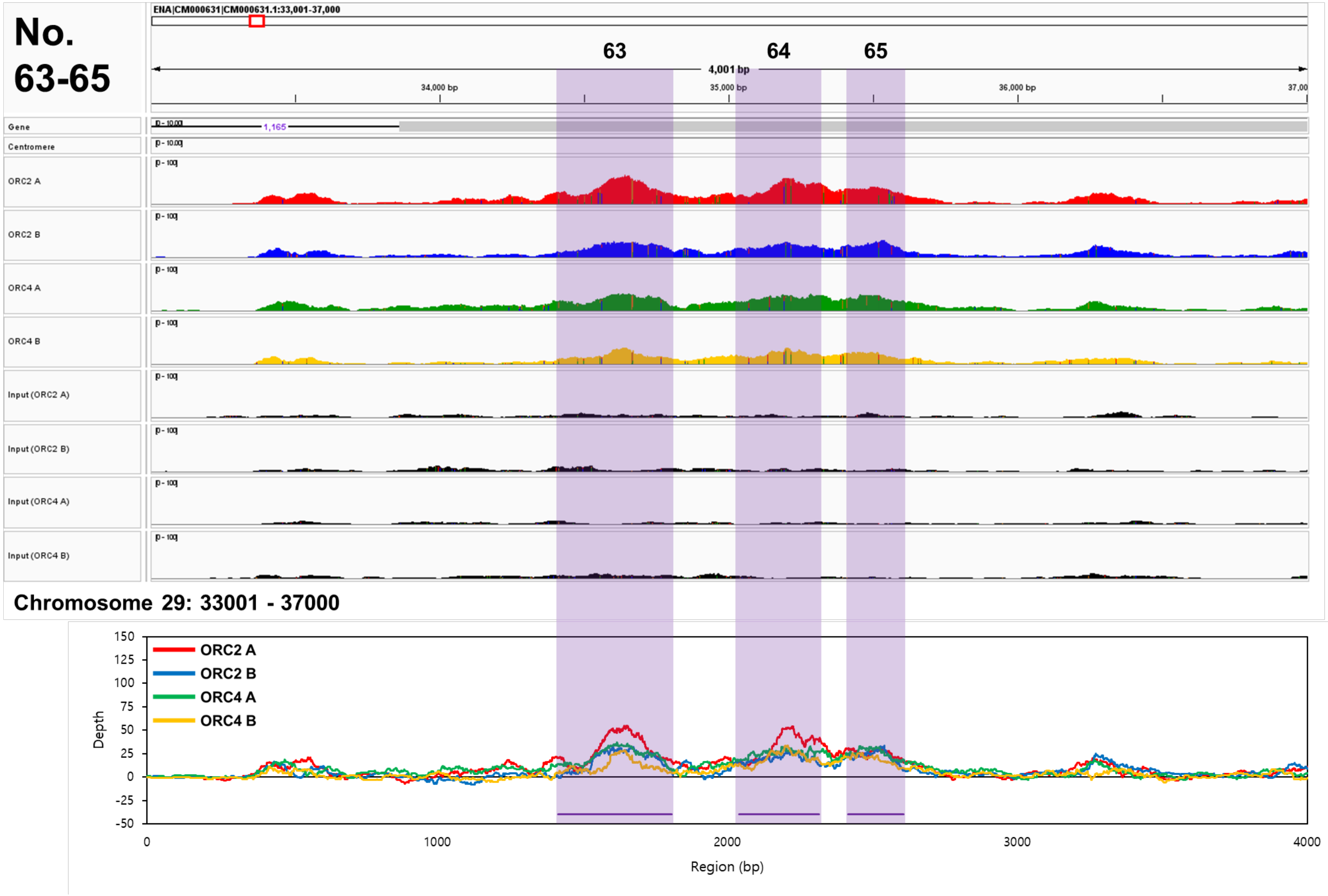
No. 63-65
ENA|CM000631|CM000631.1:33,001-37,000
63
64
65
4,001 bp
34,000 bp
35,000 bp
36,000 bp
Gene
Centromere
ORC2 A
ORC2 B
ORC4 A
ORC4 B
Input (ORC2 A)
Input (ORC2 B)
Input (ORC4 A)
Input (ORC4 B)
Chromosome 29: 33001 - 37000
ORC2 A
ORC2 B
ORC4 A
ORC4 B
Depth
Region (bp)

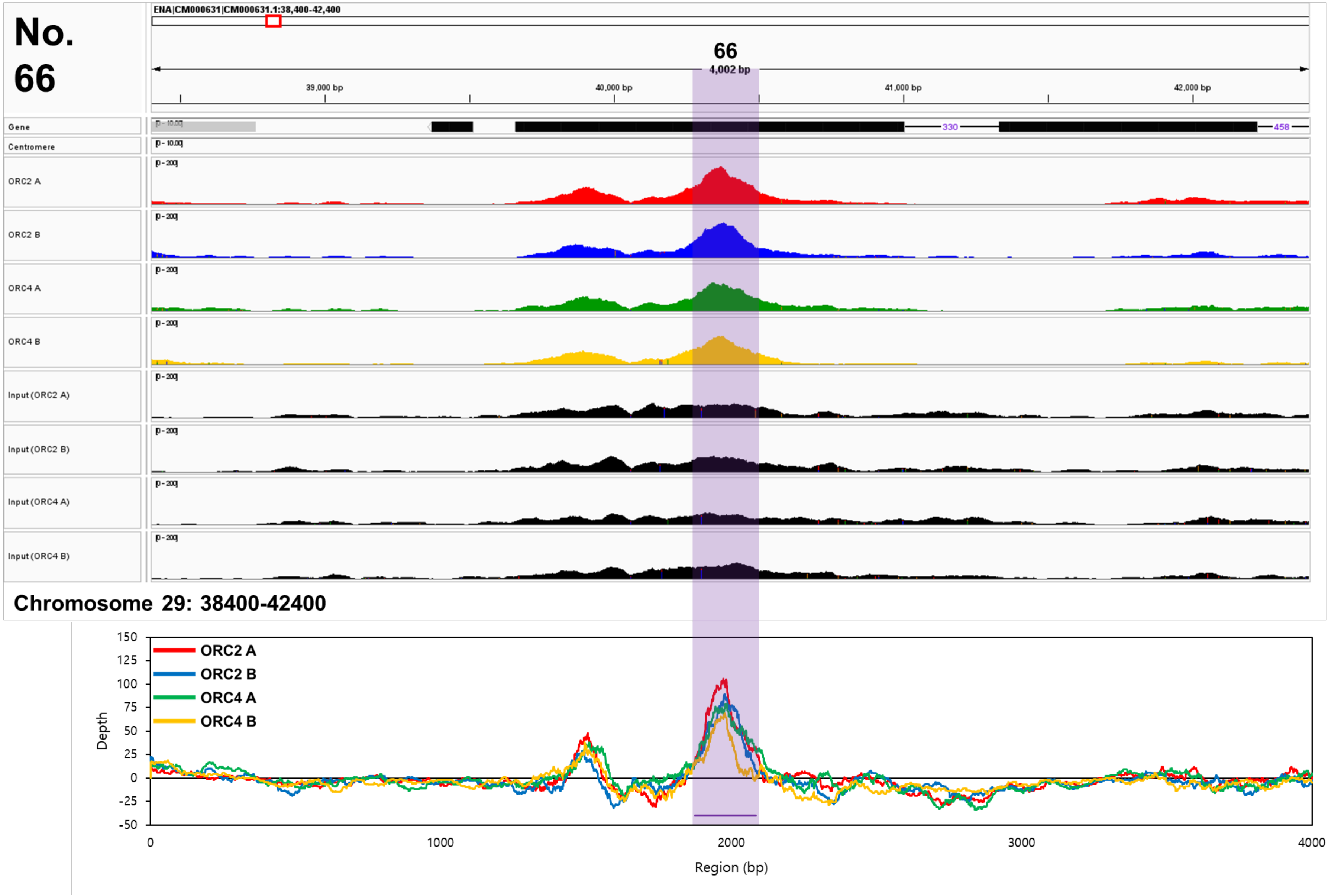
No. 66
ENA|CM000631|CM000631.1:38,400-42,400
66
4,002 bp
39,000 bp
40,000 bp
41,000 bp
42,000 bp
Gene
Centromere
ORC2 A
ORC2 B
ORC4 A
ORC4 B
Input (ORC2 A)
Input (ORC2 B)
Input (ORC4 A)
Input (ORC4 B)
Chromosome 29: 38400-42400
ORC2 A
ORC2 B
ORC4 A
ORC4 B
Depth
Region (bp)

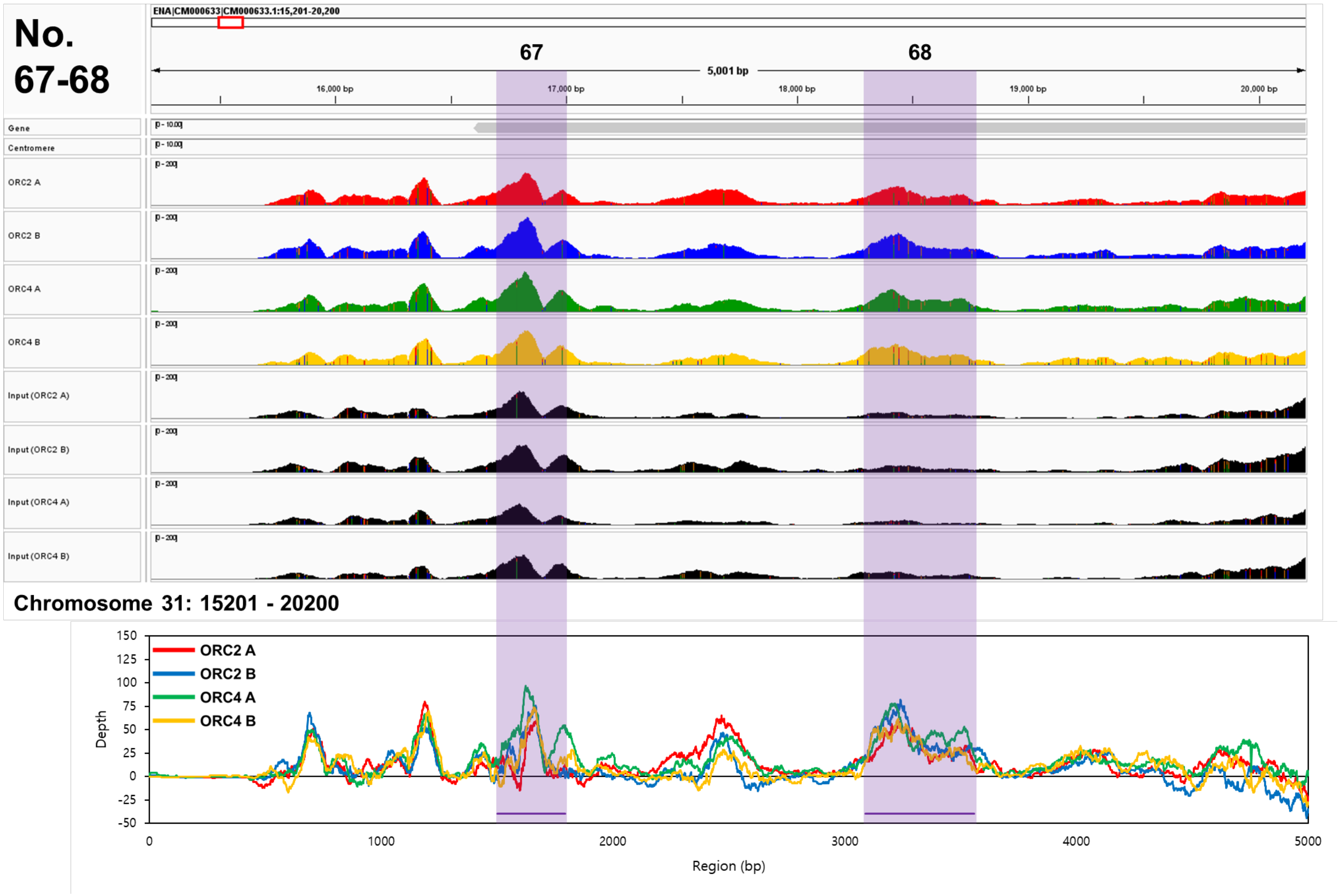
No.
67-68
ENA|CM000633|CM000633.1:15,201-20,200
67
68
5,001 bp
16,000 bp
17,000 bp
18,000 bp
19,000 bp
20,000 bp
Gene
Centromere
ORC2 A
ORC2 B
ORC4 A
ORC4 B
Input (ORC2 A)
Input (ORC2 B)
Input (ORC4 A)
Input (ORC4 B)
Chromosome 31: 15201 - 20200
ORC2 A
ORC2 B
ORC4 A
ORC4 B
Depth
Region (bp)

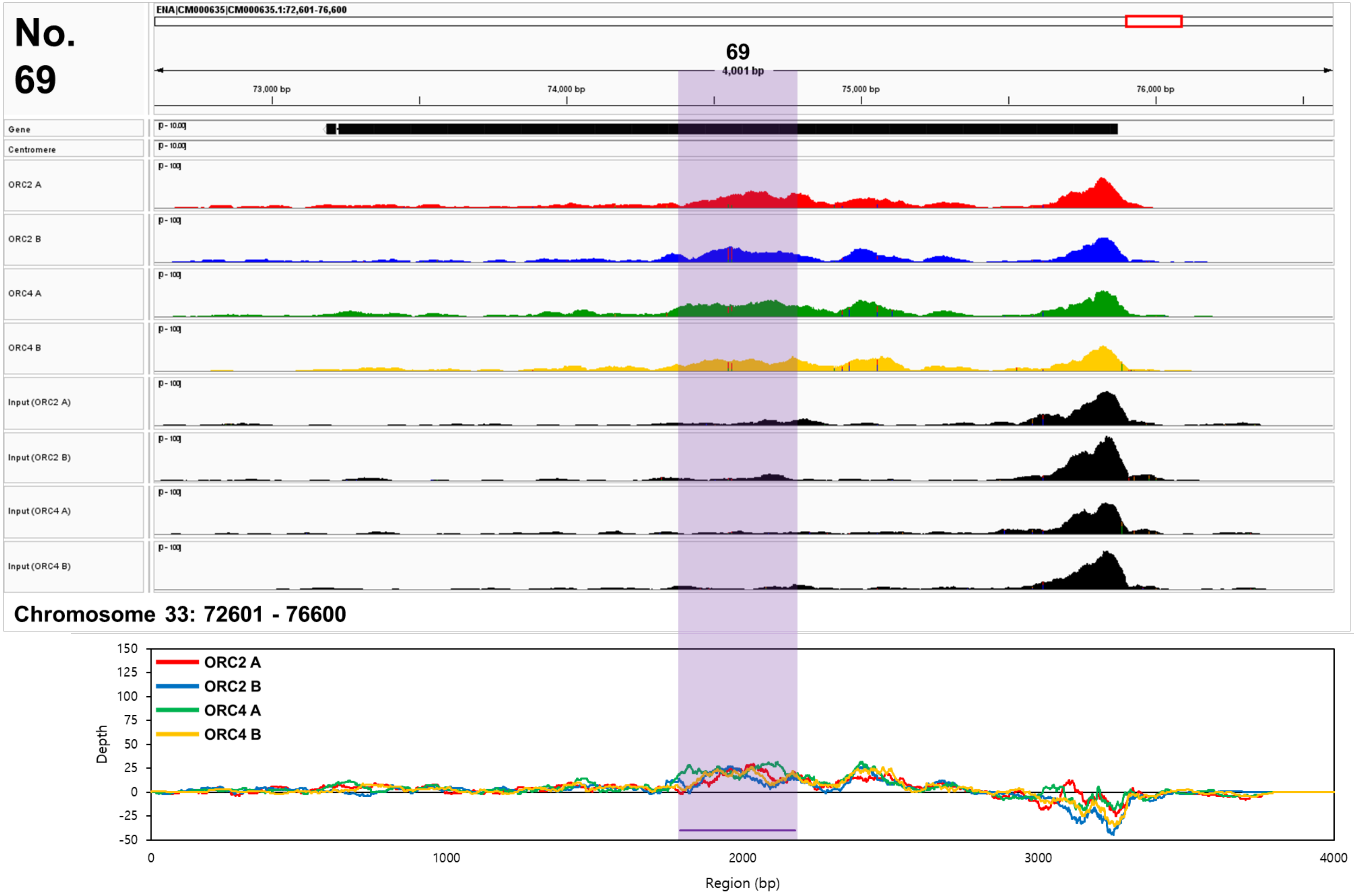
No. 69
ENA|CM000635|CM000635.1:72,601-76,600
69
4,001 bp
73,000 bp
74,000 bp
75,000 bp
76,000 bp
Gene
Centromere
ORC2 A
ORC2 B
ORC4 A
ORC4 B
Input (ORC2 A)
Input (ORC2 B)
Input (ORC4 A)
Input (ORC4 B)
Chromosome 33: 72601 - 76600
ORC2 A
ORC2 B
ORC4 A
ORC4 B
Depth
Region (bp)
150
125
100
75
50
25
0
-25
-50
0
1000
2000
3000
4000

**Fig. S11: Peak data and depth values at the locations of 68 shared sites (site numbers 1–20 and 22–69) on the genome of *Phaeodactylum tricornutum*.**

The results obtained from ORC2 A (red peaks and lines), ORC2 B (blue peaks and lines), ORC4 A (green peaks and lines), and ORC4 B (orange peaks and lines) were visualised through peak (IGV image) and depth images, and shared sites that are autonomously replicating sequence candidates indicated by purple lines over the peak and depth images.

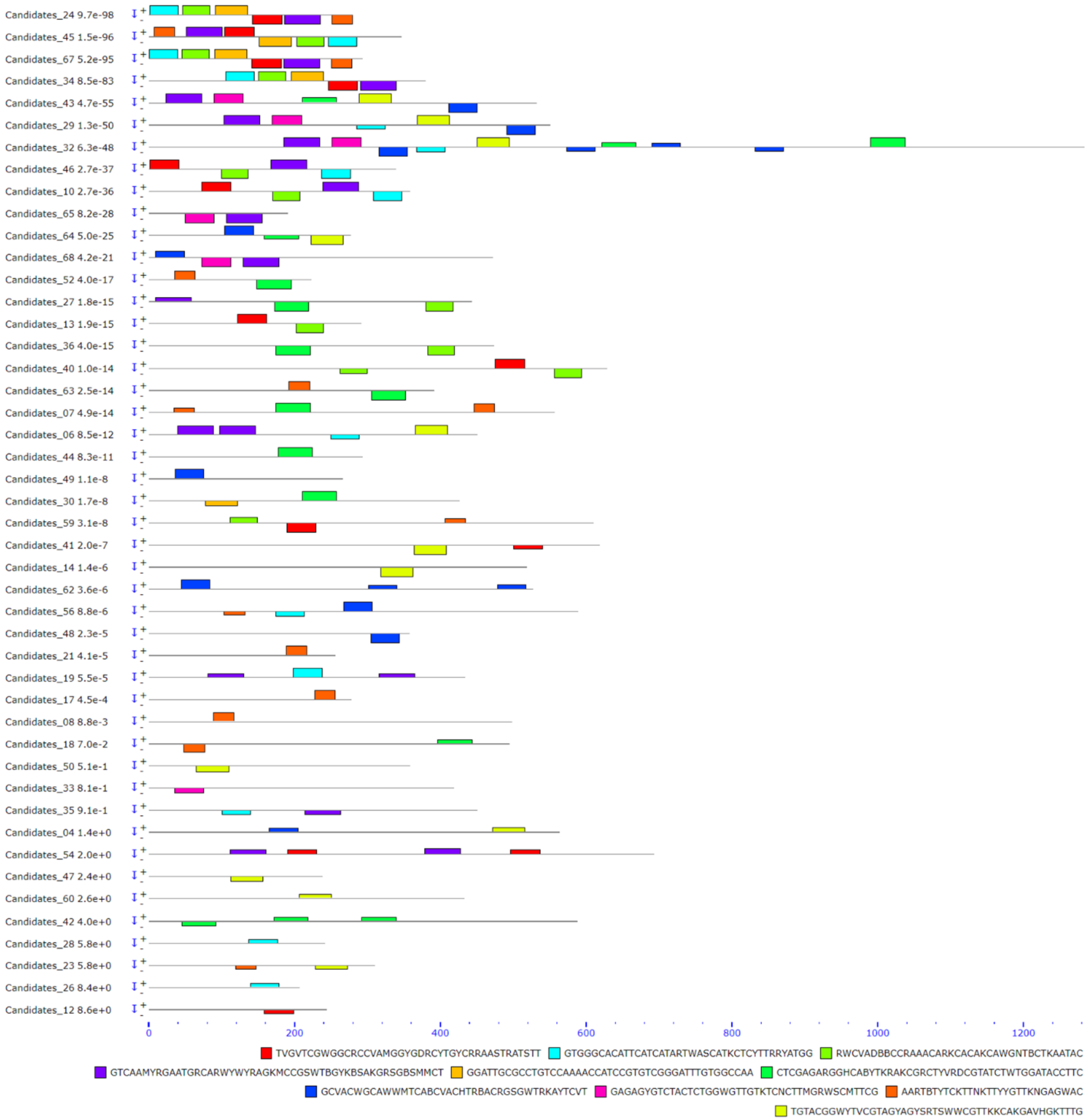
Candidates_24 9.7e-98
Candidates_45 1.5e-96
Candidates_67 5.2e-95
Candidates_34 8.5e-83
Candidates_43 4.7e-55
Candidates_29 1.3e-50
Candidates_32 6.3e-48
Candidates_46 2.7e-37
Candidates_10 2.7e-36
Candidates_65 8.2e-28
Candidates_64 5.0e-25
Candidates_68 4.2e-21
Candidates_52 4.0e-17
Candidates_27 1.8e-15
Candidates_13 1.9e-15
Candidates_36 4.0e-15
Candidates_40 1.0e-14
Candidates_63 2.5e-14
Candidates_07 4.9e-14
Candidates_06 8.5e-12
Candidates_44 8.3e-11
Candidates_49 1.1e-8
Candidates_30 1.7e-8
Candidates_59 3.1e-8
Candidates_41 2.0e-7
Candidates_14 1.4e-6
Candidates_62 3.6e-6
Candidates_56 8.8e-6
Candidates_48 2.3e-5
Candidates_21 4.1e-5
Candidates_19 5.5e-5
Candidates_17 4.5e-4
Candidates_08 8.8e-3
Candidates_18 7.0e-2
Candidates_50 5.1e-1
Candidates_33 8.1e-1
Candidates_35 9.1e-1
Candidates_04 1.4e+0
Candidates_54 2.0e+0
Candidates_47 2.4e+0
Candidates_60 2.6e+0
Candidates_42 4.0e+0
Candidates_28 5.8e+0
Candidates_23 5.8e+0
Candidates_26 8.4e+0
Candidates_12 8.6e+0
0
200
400
600
800
1000
1200
TVGVTCGWGGCRCCVAMGGYGDRCYTGYCRRAASTRATSTT
GTGGGCACATTCATCATARTWASCATKCTCYTTRRYATGG
RWCVADBBCCRAAACARKCACAKCAWGNTBCTKAATAC
GTCAAMYRGAATGRCARWYWYRAGKMCCGSWTBGYKBSAKGRSGBSMMCT
GGATTGCGCCTGTCCAAAACCATCCGTGTCGGGATTTGTGGCCAA
CTCGAGARGGHCABYTKRAKCGRCTYVRDCGTATCTWTGGATACCTTC
GCVACWGCAWWMTCABCVACHTRBACRGSGWTRKAYTCVT
GAGAGYGTCTACTCTGGWGTTGTKTCNCTTMGRWSCMTTCG
AARTBTYTCKTTNKTTYYGTTKNGAGWAC
TGTACGGWYTVCGTAGYAGYSRTSWWCGTTKKCAKGAVHGKTTTG

1. TVGVTCGWGGCRCCVAMGGYGDRCYTGYCRRAASTRATSTT

| Name | Strand | Start | *p*-value | | Sites | |
|---|---|---|---|---|---|---|
| 67. Candidates_67 | - | 142 | 6.47e-23 | GCGCACTACT | TGGGTCGTGGCACCCAAGGCGAGCCTGTCAAAACTAATCTT | GTGTGTTTGG |
| 24. Candidates_24 | - | 143 | 6.47e-23 | GCGCACTACT | TGGGTCGTGGCACCCAAGGCGAGCCTGTCAAAACTAATCTT | GTGTGTTTGG |
| 45. Candidates_45 | + | 105 | 1.73e-22 | GCGCACTACT | TGGGTCGTGGCACCCAAGGCGAGCCTGTCGAAACTAATCTT | GTGTGTTTGG |
| 34. Candidates_34 | - | 247 | 3.25e-20 | GCGCACTACT | TGGGTCGTGGCACCCAAGGCAAGCCTGTCAAAACTAATCTT | GTGTGTTTGG |
| 40. Candidates_40 | + | 476 | 2.83e-16 | TCCTCCCTCA | TCGACAGAGGCGCAAACGGCGGACTTGCCGGAAGCGATGTT | AAAATCCTTA |
| 13. Candidates_13 | + | 122 | 2.83e-16 | TCCTCCCTCA | TCGACAGAGGCGCAAACGGCGGACTTGCCGGAAGCGATGTT | AAAATCCTTA |
| 59. Candidates_59 | - | 190 | 3.41e-15 | TCTGCACTTG | TCGATCGCGGTGCCAACGGTGGACTTGCGGGGAGTGATGTT | ACCGTACTGC |
| 46. Candidates_46 | + | 2 | 9.19e-15 | G | AAGCTGTACGCACCGCAGTTGTACTGGCCAGAACAAAACAT | GATGGTACAG |
| 10. Candidates_10 | + | 73 | 9.19e-15 | ACTGCTGGAG | AAGCTGTACGCACCGCAGTTGTACTGGCCAGAACAAAACAT | GATGGTACAG |

2. GTGGGCACATTCATCATARTWASCATKCTCYTTRRYATGG

| Name | Strand | Start | *p*-value | | Sites | |
|---|---|---|---|---|---|---|
| 67. Candidates_67 | + | 1 | 8.05e-25 | | GTGGGCACATTCATCATAGTTACCATGCTCCTTAGCATGG | CAACGATCGA |
| 45. Candidates_45 | - | 247 | 8.05e-25 | ACACGGCACC | GTGGGCACATTCATCATAGTTACCATGCTCCTTAGCATGG | CAACGATCGA |
| 34. Candidates_34 | + | 106 | 8.05e-25 | ACACGGCACC | GTGGGCACATTCATCATAGTTACCATGCTCCTTAGCATGG | CTACGGTCAA |
| 24. Candidates_24 | + | 2 | 8.05e-25 | C | GTGGGCACATTCATCATAGTTACCATGCTCCTTAGCATGG | CAAYGATCGA |
| 46. Candidates_46 | - | 238 | 6.65e-17 | TGACAGCACA | GTGGGTACCTGCAGCCAAATAAGGGTTCTGTTTGATACGG | GTCGACGGAT |
| 10. Candidates_10 | - | 309 | 6.65e-17 | TGACAGCACA | GTGGGTACCTGCAGCCAAATAAGGGTTCTGTTTGATACGG | GTCGACGGAT |
| 19. Candidates_19 | + | 199 | 3.52e-15 | CTCCCCCAAA | GGCTCCAAAGTCATCATCATCAGCATTCTCGTTGTTATCG | TCAGCCTCAT |

3. RWCVADBBCCRAAACARKCACAKCAWGNTBCTKAATAC

| | Name | Strand | Start | *p*-value | Sites | | |
|---|---|---|---|---|---|---|---|
| 34. | Candidates_34 | + | 151 | 2.69e-21 | CATGGCTACG | GTCAAGTCCCAAAACAGTCACATCATGTTTCTGAATAC | TCAGCTGGGA |
| 67. | Candidates_67 | + | 46 | 1.71e-20 | CATGGCAACG | ATCGAGTCCCGAAACAGTCACATCATGTTTCTGAATAC | TCAACTGGGA |
| 45. | Candidates_45 | - | 204 | 1.71e-20 | CATGGCAACG | ATCGAGTCCCGAAACAGTCACATCATGTTTCTGAATAC | TCAACTGGGA |
| 24. | Candidates_24 | + | 47 | 1.71e-20 | CATGGCAAYG | ATCGAGTCCCGAAACAGTCACATCATGTTTCTGAATAC | TCAGCTGGGA |
| 40. | Candidates_40 | - | 557 | 6.01e-15 | CAGCGGCGGT | GACAATGTCCAAATCAGGCAAAGTATGGTCATTAATAC | CCGTGATGCT |
| 13. | Candidates_13 | - | 203 | 6.01e-15 | CAGCGGCGGT | GACAATGTCCAAATCAGGCAAAGTATGGTCATTAATAC | CCGTGATGCT |
| 27. | Candidates_27 | - | 381 | 2.55e-13 | GGAGTAAGTC | GTCCTTGTCCCGGACAATGCCTTCAAGATCCTGAATTK | GGCGGCGGAG |
| 46. | Candidates_46 | - | 100 | 2.64e-13 | AACCAATACC | AACCAACGCCAATACAAGCACAGGGAGCTGCTTCACAG | CCCAAAGAAG |
| 10. | Candidates_10 | - | 171 | 2.64e-13 | AACCAATACC | AACCAACGCCAATACAAGCACAGGGAGCTGCTTCACAG | CCCAAAGAAG |
| 36. | Candidates_36 | - | 383 | 2.74e-13 | GGAGTAAGTC | GTCCTTGTCCCGGACAATGCCTTCAAGATCCTGAATTT | GGCGGCGGAG |

4. GTCAAMYRGAATGRCARWYWYRAGKMCCGSWTBGYKBSAKGRSGBSMMCT

| | Name | Strand | Start | *p*-value | Sites | | |
|---|---|---|---|---|---|---|---|
| 45. | Candidates_45 | + | 52 | 9.33e-21 | AGTGGGGAGT | GTCAACTGGAATGACAGTTTTGAGTACCGGTTCGCGCGATGGCGCGCACT | ACTTGGGTCG |
| 24. | Candidates_24 | - | 187 | 9.33e-21 | AGTGGGGAGT | GTCAACTGGAATGACAGTTTTGAGTACCGGTTCGCGCGATGGCGCGCACT | ACTTGGGTCG |
| 29. | Candidates_29 | + | 104 | 1.77e-20 | CGTATATGAC | GTCAAGCATGATGGCAGACACAAGGCCAGGATGGTGGCAGGTGGTCACCT | AACCCCAGTA |
| 34. | Candidates_34 | - | 291 | 4.49e-19 | ATCAAATCAT | GTCAACTGGAATGACAGTTTTGAGTACTGGTTTGCGTGATGGCGCGCACT | ACTTGGGTCG |
| 67. | Candidates_67 | - | 186 | 2.13e-18 | AGTGGGGAGT | GTCAACTGGAATGACAGTTTTGAKTACCGGTTCGCRCGATGGCGCGCACT | ACTTGGGTCG |
| 65. | Candidates_65 | - | 107 | 9.64e-18 | GGTCTACGAT | GTTAAACATGATGGTAGACACAAGGCCCGTATGGTTGCTGGAGGTCACCT | WACCCCCGTG |
| 43. | Candidates_43 | + | 24 | 7.63e-17 | TGTCTATGAT | GTTAAACATGACGGTAGACACAAGGCTCGCATGGTTGCTGGAGGTCACCT | TACCCCTGTG |
| 46. | Candidates_46 | + | 168 | 2.79e-15 | TGTCTACAGT | GACAACAAGAACAGCAACAACAACGACAGCAATGGGTGACGATGGCAACG | GTCAAACCGG |
| 10. | Candidates_10 | + | 239 | 1.38e-14 | TGTCTACAGT | GACAACAAGAACAGCAACAACAACGACAGCAATGGGTGACAATGGCAACG | GTCAAACCGG |
| 68. | Candidates_68 | - | 130 | 2.08e-14 | CGTGTTTGAC | GTGAAACAGTCACTCAAATACAAGGCCCGGCTCGTCGCCGGTGGTCACAT | GACAGCGCCT |
| 32. | Candidates_32 | + | 186 | 1.48e-13 | YGTATTTGAT | GTCAAGCACAGTCTCAAACGAAAGGCCCGCCTAGTTGCYGGAGGGCAYAC | TACAGCGCCR |

5. GGATTGCGCCTGTCCAAAACCATCCGTGTCGGGATTTGTGGCCAA

| Name | Strand | Start | *p*-value | Sites |
|---|---|---|---|---|
| 67. Candidates_67 | + | 91 | 8.55e-28 | TACTCAACTG GGATTGCGCCTGTCCAAAACCATCCGTGTCGGGATTTGTGGCCAA ACACACAAGA |
| 45. Candidates_45 | - | 152 | 8.55e-28 | TACTCAACTG GGATTGCGYCTGTCCAAAACCATCCGTGTCGGGATTTGTGGCCAA ACACACAAGA |
| 34. Candidates_34 | + | 196 | 8.55e-28 | TACTCAGCTG GGATTGCGCCTGTCCAAAACCATCCGTGTCGGGATTTGTGGCCAA ACACACAAGA |
| 24. Candidates_24 | + | 92 | 8.55e-28 | TACTCAGCTG GGATTGCGCCTGTCCAAAACCATCCGTGTCGGGATTTGTGGCCAA ACACACAAGA |

6. CTCGAGARGGHCABYTKRAKCGRCTYVRDCGTATCTWTGGATACCTTC

| Name | Strand | Start | *p*-value | Sites |
|---|---|---|---|---|
| 44. Candidates_44 | + | 178 | 6.93e-22 | CGTGCTGCCC CTCGAGAAGGACATTTGAATCGYCTTCGTCGTATCATYGGATACCTTC GTCGTTACCC |
| 52. Candidates_52 | - | 149 | 3.33e-21 | CGTGTTGCTC CCCGAGAAGGCCACCTTGATCGCCTTAAGCGTATGTACGGATACCTTC GTAAGATGAA |
| 63. Candidates_63 | - | 306 | 3.71e-19 | CGTGTTGCTC CCCRACAAGGTCATCTYGATCGACTCAAACGTATGTATGGGTACCTTC GTAAGATGAA |
| 32. Candidates_32 | + | 991 | 2.42e-18 | CGTGYGGCTC CACGTAAGGGACATCTTGAGAGACTCAAGCGTATCATCGGATACCTAC GACACTACCC |
| 7. Candidates_07 | + | 175 | 3.35e-18 | ACTGCGTGCC CTCGAAAGGGACATTTGGAGCGACTTCTGAGGGTCTTTGGATATCTGA AGAAGCGACC |
| 30. Candidates_30 | + | 211 | 1.04e-17 | CGTGTTGCGC CTCGCGAGGGCCACCTAGATCGACTCAAAAGGATCTATGGTTACGTTA GGAAGATGAA |
| 36. Candidates_36 | - | 175 | 4.37e-17 | GTTCGTTTTC CTCTAGAAGTTGTGTAGATGGGGACTGGTCGTCTCTTTGGAGAACTTC CACGTACGGG |
| 27. Candidates_27 | - | 173 | 4.37e-17 | GTTCGTTTTC CTCTAGAAGTTGTGTAGATGGGGACTGGTCGTCTCTTTGGAGAACTTC CACGTACGGG |

7. GCVACWGCAWWMTCABCVACHTRBACRGSGWTRKAYTCVT

| Name | Strand | Start | *p*-value | Sites |
|---|---|---|---|---|
| 64. Candidates_64 | + | 105 | 3.38e-20 | GTCATGCGCT GCGACAGCAATATCATCGACATACACGGCGATGTACTCGT AAACACCTCG |
| 29. Candidates_29 | - | 492 | 7.50e-18 | ATCATGGGCT GCCACTGCAATGTCATCAACATACACAGCAATATACTCAT ACACACCGGA |
| 62. Candidates_62 | + | 45 | 1.01e-16 | CAACTTGTTT GCAACGGCATACTCAGCGACTTCGACAGGGTTAGATTCCT TCAGRTCTTT |
| 49. Candidates_49 | + | 37 | 1.13e-16 | AAGCTTGTTA GCAACTGCATATTCAGCAACTTGCACAGGGTTTGATTCCT TAAGGTTGGC |
| 43. Candidates_43 | - | 412 | 2.70e-16 | ATCATGGGCT GCAATAGCAATATCGTCGACGTAGACGGCAATGTACTCAT AGACGCCTTG |
| 48. Candidates_48 | - | 305 | 8.04e-16 | AAGTTTGTTT GCTACCGCATACTCGGCCACTTGTACGGGGTTAGATTCCT TGAGGTTAGC |
| 56. Candidates_56 | + | 268 | 1.17e-12 | CAACAGCTGA GCCACAGCAAGAGGCCCCAACTTTACGGCGTTCTACACGA AAGAGAGTAG |
| 32. Candidates_32 | - | 316 | 3.14e-12 | TTCCAAATAT GCGACTGCAATRTCACCAACCCAGGTGTCGAGATYGTTGA GTTCTCCTGC |

8. GAGAGYGTCTACTCTGGWGTTGTKTCNCTTMGRWSCMTTCG

| Name | Strand | Start | *p*-value | | Sites | |
|---|---|---|---|---|---|---|
| 43. Candidates_43 | + | 90 | 4.22e-25 | TGTGCCCCCT | GAGAGCGTCTACTCTGGAGTTGTGTCTCTTCGAAGCCTTCG | TATTGTTGTG |
| 65. Candidates_65 | - | 50 | 4.65e-24 | CGTGCCCCCT | GAGAGCGTCTACTCTGGAGTTGTGTCCCTTCGAAGCCTTCG | CATTGTCGTG |
| 68. Candidates_68 | - | 73 | 4.65e-20 | GCCTCCAAAG | GACAGTGTTTACTCCGGTGTTGTTTCTCTTCGGTCCATTCG | TCTCGCTATT |
| 29. Candidates_29 | + | 170 | 6.68e-20 | AGTACCAAAT | GAGAGTGTCTATTCAGGAGTCGTTTCACTCAGAAGCCTTCG | AATTGTCATA |
| 32. Candidates_32 | + | 252 | 3.52e-18 | GCCRCCGAAG | GATAGCGTATACTCTGGTGTGGTYTCGCTKAGGTCYATCCG | GTTRGCACTG |

9. AARTBTYTCKTTNKTTYYGTTKNGAGWAC

| Name | Strand | Start | *p*-value | | Sites | |
|---|---|---|---|---|---|---|
| 67. Candidates_67 | - | 251 | 6.68e-14 | AGGTGAGACG | AAATCTCCCGTTCGTTCTGTTTCGAGTAC | CTCACAGTGG |
| 45. Candidates_45 | + | 8 | 6.68e-14 | TGAGACG | AAATCTCCCGTTCGTTCTGTTTCGAGTAC | CTCACAGTGG |
| 24. Candidates_24 | - | 252 | 6.68e-14 | | AAATCTCCCGTTCGTTCTGTTTCGAGTAC | CTCACAGTGG |
| 21. Candidates_21 | + | 189 | 1.05e-12 | CCAAACCCAT | AAGTGKTTCTTTTTTTTCGTTTTGAGAAC | CGCAAGAACC |
| 17. Candidates_17 | + | 228 | 1.05e-12 | CCAAACCCAT | AAGTGKTTCTTTTTTTTCGTTTTGAGAAC | CGCAAGAACC |
| 63. Candidates_63 | + | 193 | 1.94e-10 | CGTGGGGAAC | AAGTTCCTCGACGTTTCCGTATATGGAAC | GTGCCCATTC |
| 8. Candidates_08 | + | 89 | 1.94e-10 | GCGTTCCATA | ACGGGTCTTTTGATTTTCGTCGGGCGTAC | CCCTGTGTTT |
| 7. Candidates_07 | + | 447 | 1.94e-10 | GCGTTCCATA | ACGGGTCTTTTGATTTTCGTCGGGCGTAC | CCCTGTGTTT |
| 52. Candidates_52 | + | 36 | 7.71e-10 | CGTGGGGAAC | AAGTTCTTCCACATTTCCGTAGATGGAAC | GGGCCCATTC |
| 18. Candidates_18 | - | 49 | 1.51e-9 | TGCCCCGGCT | ACATCTGTTGTTGCCTCTGTTGCTAGAAG | TACTCTGGAA |

10. TGTACGGWYTVCGTAGYAGYSRTSWWCGTTKKCAKGAVHGKTTTG

| Name | Strand | Start | *p*-value | | Sites | |
|---|---|---|---|---|---|---|
| 43. Candidates_43 | + | 289 | 4.10e-21 | AACAAGGCCT | TGTACGGTTTACGTAGCAGTGGRCTTCGTTGGCATGAGAGGTTTG | CCGATACGCT |
| 64. Candidates_64 | - | 223 | 9.45e-19 | AGCAAAGCTC | TATACGGTTTGCGTAGCAGTGGTCTCCGTTGGCATGAGAGGTTCG | CAGACACACT |
| 29. Candidates_29 | + | 369 | 4.52e-17 | AACAAGGCCC | TGTACGGACTCAGGAGTAGCGGATTGCGCTGGCACGAGCGTTTTG | CGGATACCCT |
| 32. Candidates_32 | + | 451 | 1.92e-16 | GAAAAGGCYC | TGTACGGGCTCCGYACTAGTGGCGCACGTTTCCATGAACGRTTGG | CTGACACCCT |
| 41. Candidates_41 | - | 365 | 4.49e-16 | ATAGACCTCA | CGAGCGAATTGCGTAGTGGCCATGATGGTTTTCTGGACTGTGTYG | GCGGAGACCC |
| 14. Candidates_14 | - | 319 | 6.25e-16 | ATAGACCTCA | CGAGCAAATTGCGTAGTGGCCATGATGGTTTTCTGGACTGTGTTG | GCGGAGACCC |
| 6. Candidates_06 | + | 366 | 2.85e-15 | CGAAAAGCGC | TGTACGGCCTACGATCGAGCTCGGAACGTTGGCATGCACATTTTG | CGGATACCCT |

**Fig. S12: Information on motifs predicted from the 69 shared sites.**

Motifs were predicted using the MEME Suite, and the locations of predicted motifs displayed on sequences of the shared sites. The sequence information of each predicted motif was aligned and organised.

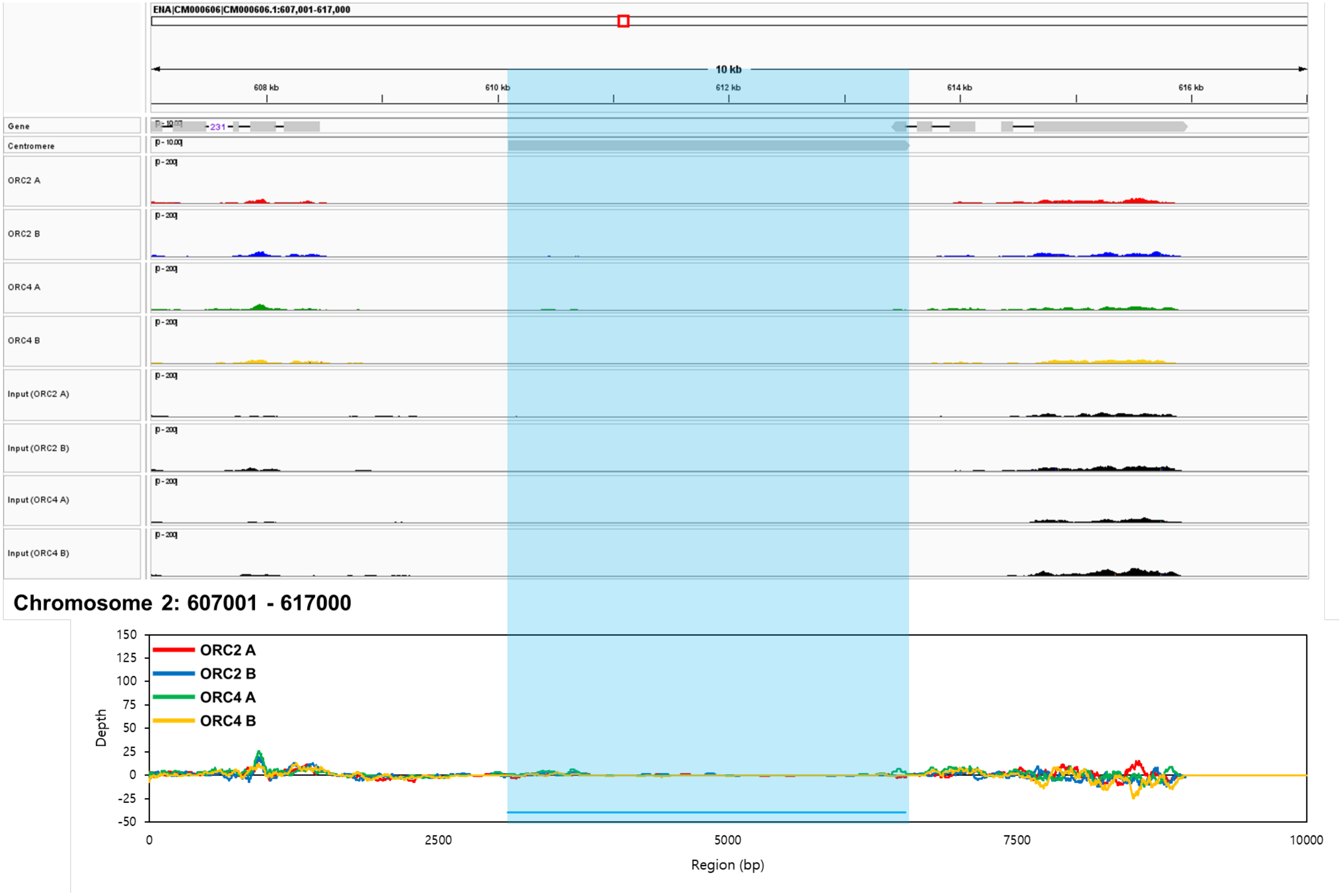
ENA|CM000606|CM000606.1:607,001-617,000
10 kb
608 kb
610 kb
612 kb
614 kb
616 kb
Gene
Centromere
ORC2 A
ORC2 B
ORC4 A
ORC4 B
Input (ORC2 A)
Input (ORC2 B)
Input (ORC4 A)
Input (ORC4 B)
Chromosome 2: 607001 - 617000
ORC2 A
ORC2 B
ORC4 A
ORC4 B
Depth
Region (bp)
150
125
100
75
50
25
0
-25
-50
0
2500
5000
7500
10000

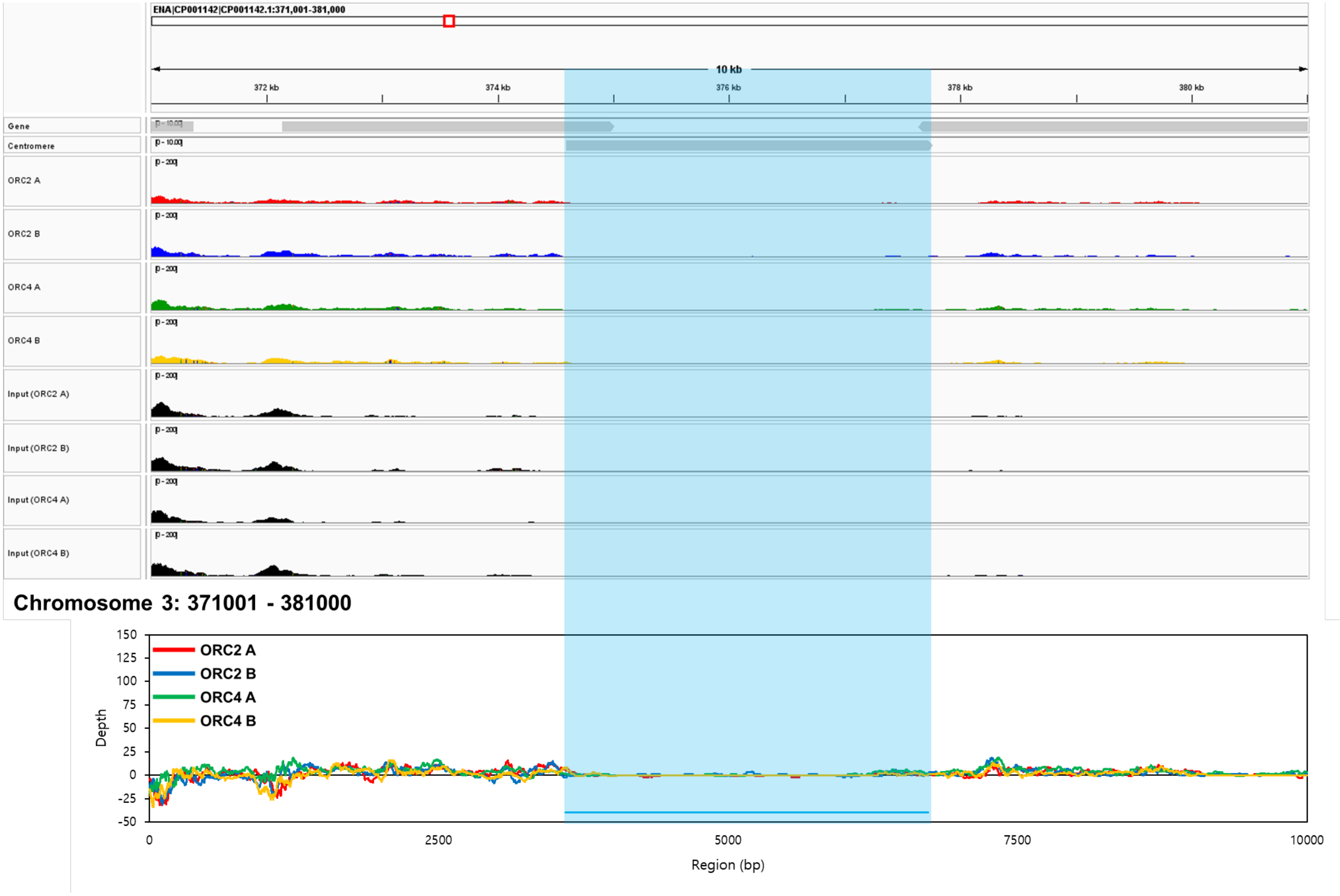

ENA|CP001142|CP001142.1:371,001-381,000
10 kb
372 kb
374 kb
376 kb
378 kb
380 kb
Gene
Centromere
ORC2 A
ORC2 B
ORC4 A
ORC4 B
Input (ORC2 A)
Input (ORC2 B)
Input (ORC4 A)
Input (ORC4 B)
Chromosome 3: 371001 - 381000
ORC2 A
ORC2 B
ORC4 A
ORC4 B
Depth
Region (bp)

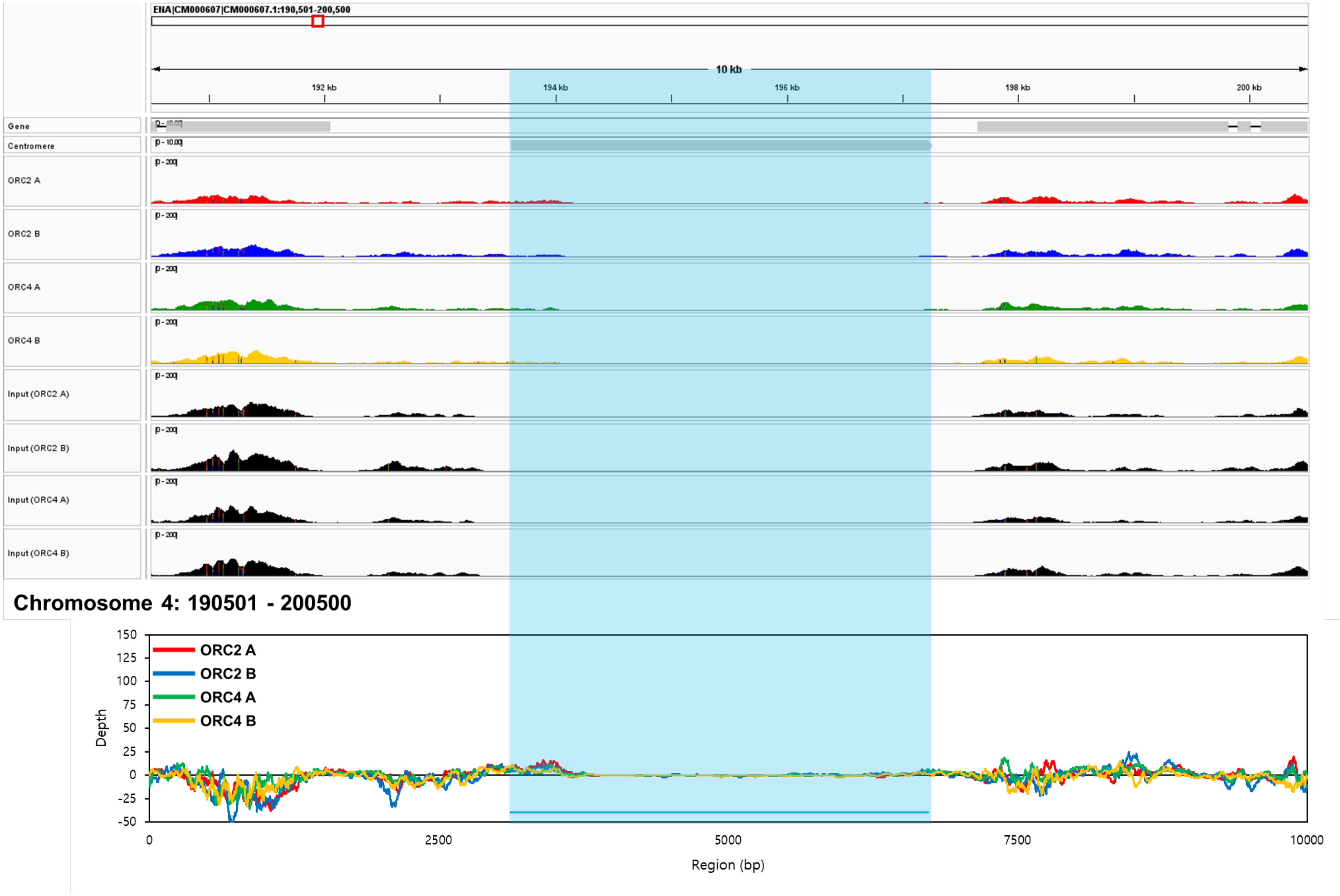
ENA|CM000607|CM000607.1:190,501-200,500
10 kb
192 kb
194 kb
196 kb
198 kb
200 kb
Gene
Centromere
ORC2 A
ORC2 B
ORC4 A
ORC4 B
Input (ORC2 A)
Input (ORC2 B)
Input (ORC4 A)
Input (ORC4 B)
Chromosome 4: 190501 - 200500
ORC2 A
ORC2 B
ORC4 A
ORC4 B
Depth
150
125
100
75
50
25
0
-25
-50
0
2500
5000
7500
10000
Region (bp)

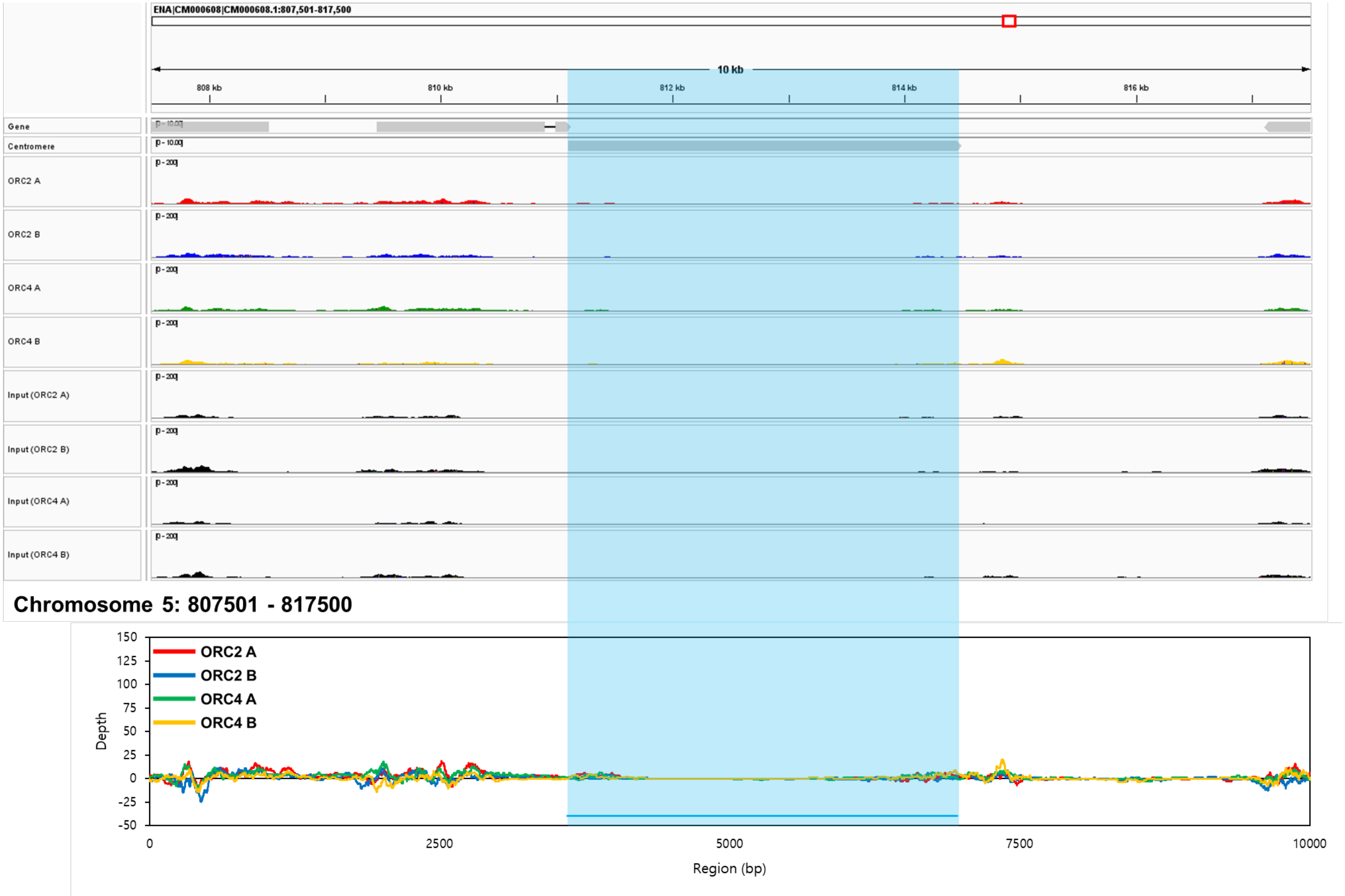
ENA|CM000608|CM000608.1:807,501-817,500
10 kb
808 kb
810 kb
812 kb
814 kb
816 kb
Gene
Centromere
ORC2 A
ORC2 B
ORC4 A
ORC4 B
Input (ORC2 A)
Input (ORC2 B)
Input (ORC4 A)
Input (ORC4 B)
Chromosome 5: 807501 - 817500
Depth
Region (bp)

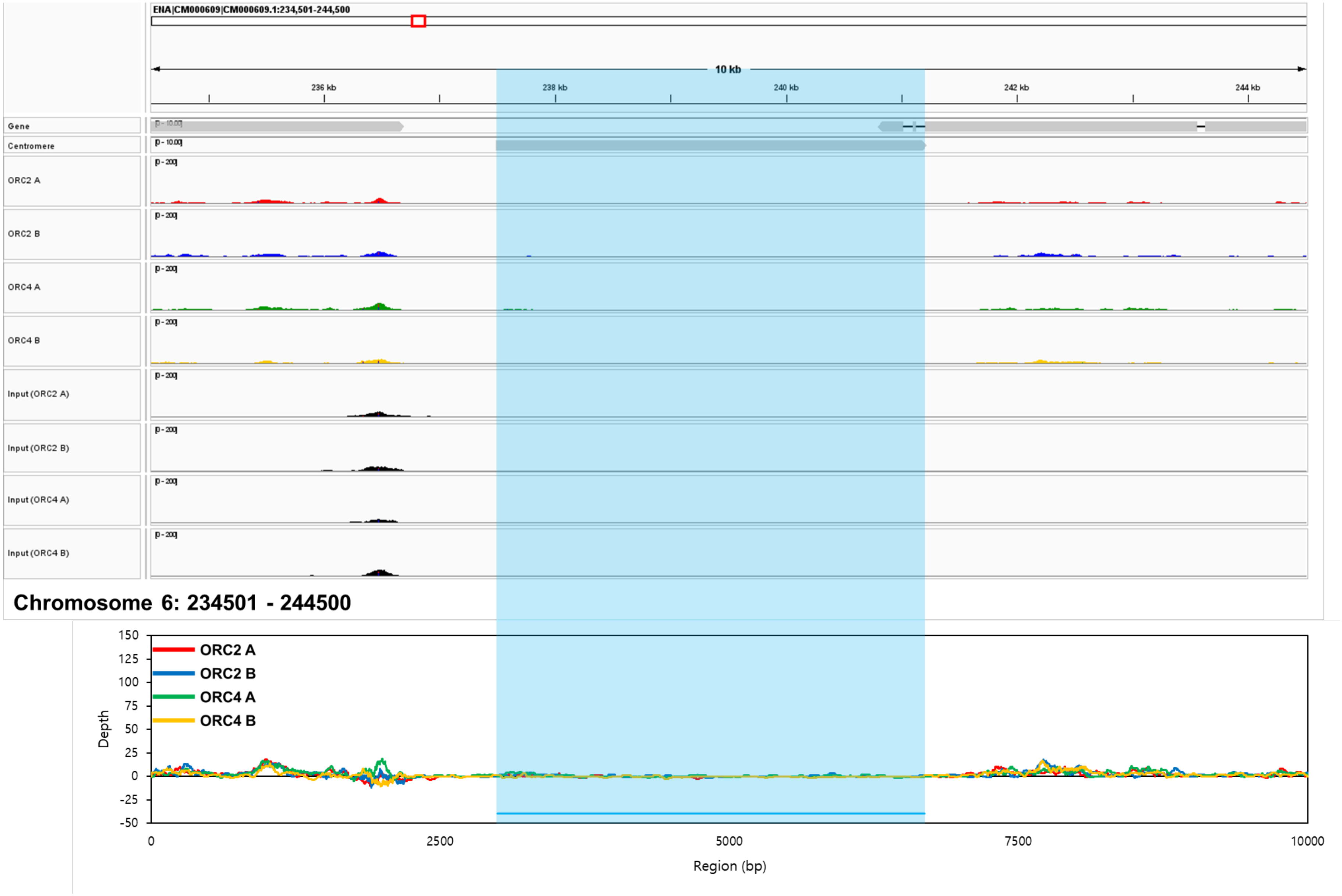
ENA|CM000609|CM000609.1:234,501-244,500
10 kb
236 kb
238 kb
240 kb
242 kb
244 kb
Gene
Centromere
ORC2 A
ORC2 B
ORC4 A
ORC4 B
Input (ORC2 A)
Input (ORC2 B)
Input (ORC4 A)
Input (ORC4 B)
Chromosome 6: 234501 - 244500
ORC2 A
ORC2 B
ORC4 A
ORC4 B
Depth
Region (bp)
150
125
100
75
50
25
0
-25
-50
0
2500
5000
7500
10000

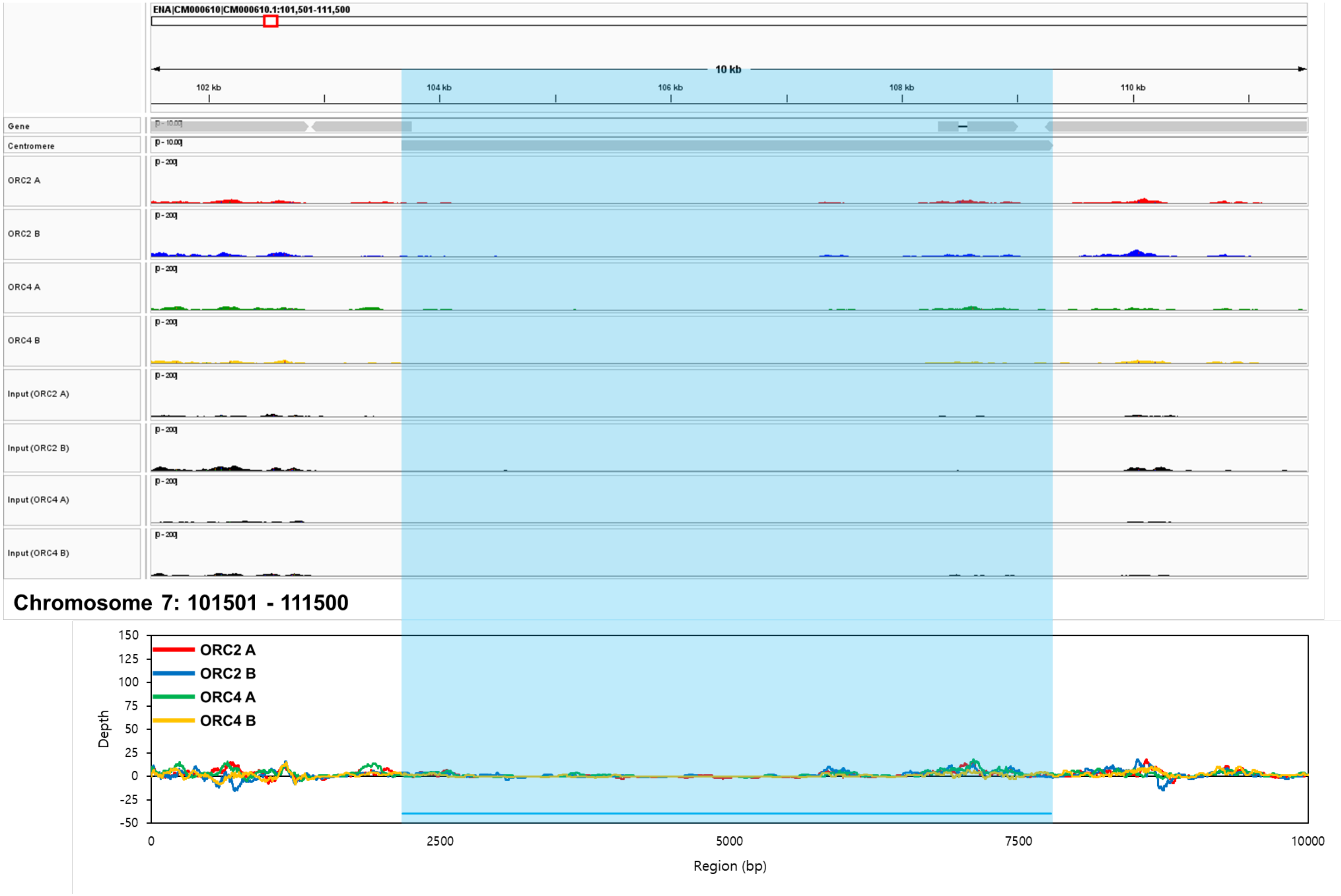

ENA|CM000610|CM000610.1:101,501-111,500
10 kb
102 kb
104 kb
106 kb
108 kb
110 kb
Gene
Centromere
ORC2 A
ORC2 B
ORC4 A
ORC4 B
Input (ORC2 A)
Input (ORC2 B)
Input (ORC4 A)
Input (ORC4 B)
Chromosome 7: 101501 - 111500
ORC2 A
ORC2 B
ORC4 A
ORC4 B
Depth
Region (bp)

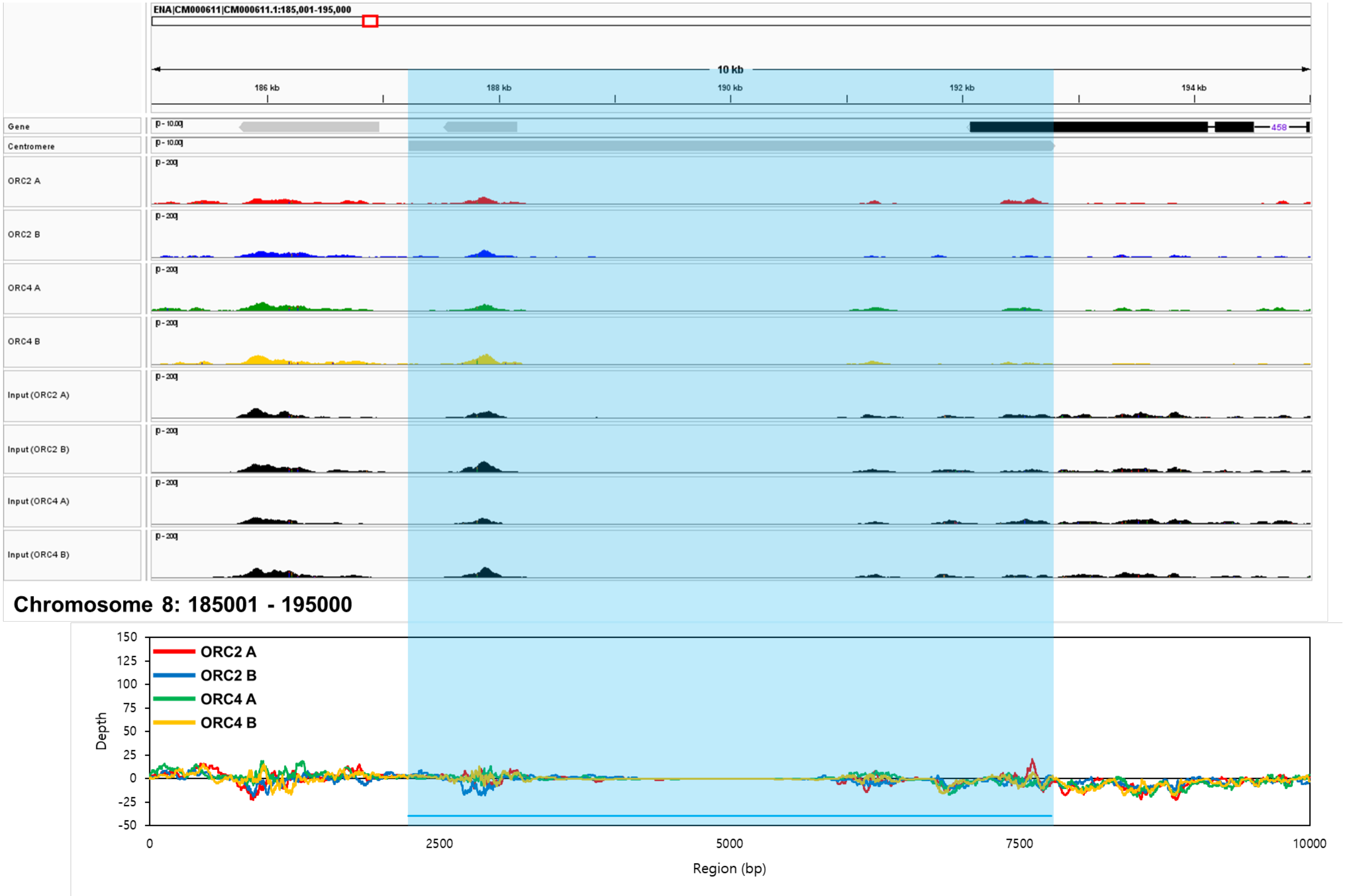
ENA|CM000611|CM000611.1:185,001-195,000
10 kb
186 kb
188 kb
190 kb
192 kb
194 kb
Gene
Centromere
ORC2 A
ORC2 B
ORC4 A
ORC4 B
Input (ORC2 A)
Input (ORC2 B)
Input (ORC4 A)
Input (ORC4 B)
Chromosome 8: 185001 - 195000
ORC2 A
ORC2 B
ORC4 A
ORC4 B
Depth
Region (bp)

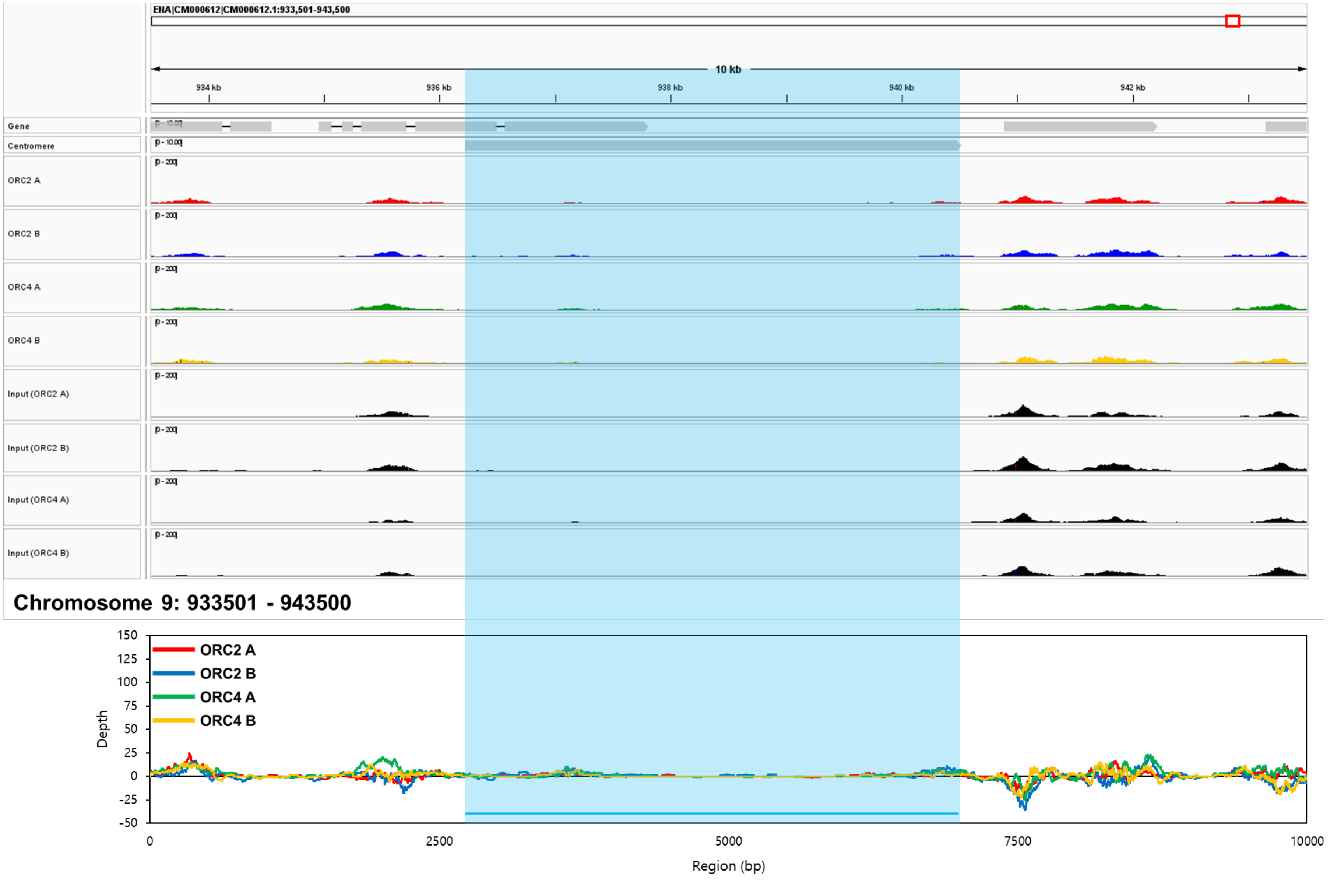
ENA|CM000612|CM000612.1:933,501-943,500
10 kb
934 kb
936 kb
938 kb
940 kb
942 kb
Gene
Centromere
ORC2 A
ORC2 B
ORC4 A
ORC4 B
Input (ORC2 A)
Input (ORC2 B)
Input (ORC4 A)
Input (ORC4 B)
Chromosome 9: 933501 - 943500
ORC2 A
ORC2 B
ORC4 A
ORC4 B
Depth
150
125
100
75
50
25
0
-25
-50
0
2500
5000
7500
10000
Region (bp)

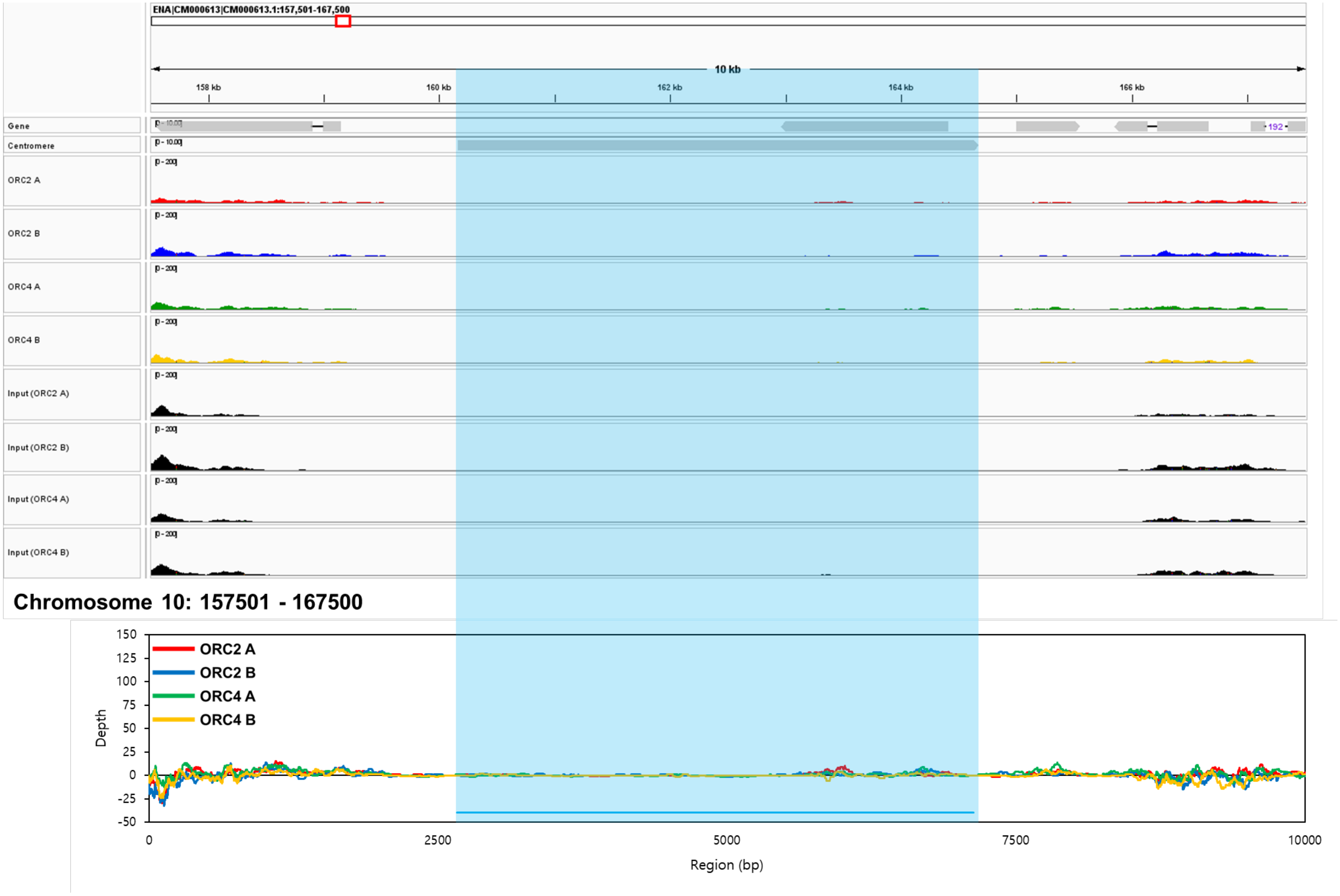
ENA|CM000613|CM000613.1:157,501-167,500
10 kb
158 kb
160 kb
162 kb
164 kb
166 kb
Gene
Centromere
ORC2 A
ORC2 B
ORC4 A
ORC4 B
Input (ORC2 A)
Input (ORC2 B)
Input (ORC4 A)
Input (ORC4 B)
Chromosome 10: 157501 - 167500
ORC2 A
ORC2 B
ORC4 A
ORC4 B
Depth
150
125
100
75
50
25
0
-25
-50
0
2500
5000
7500
10000
Region (bp)

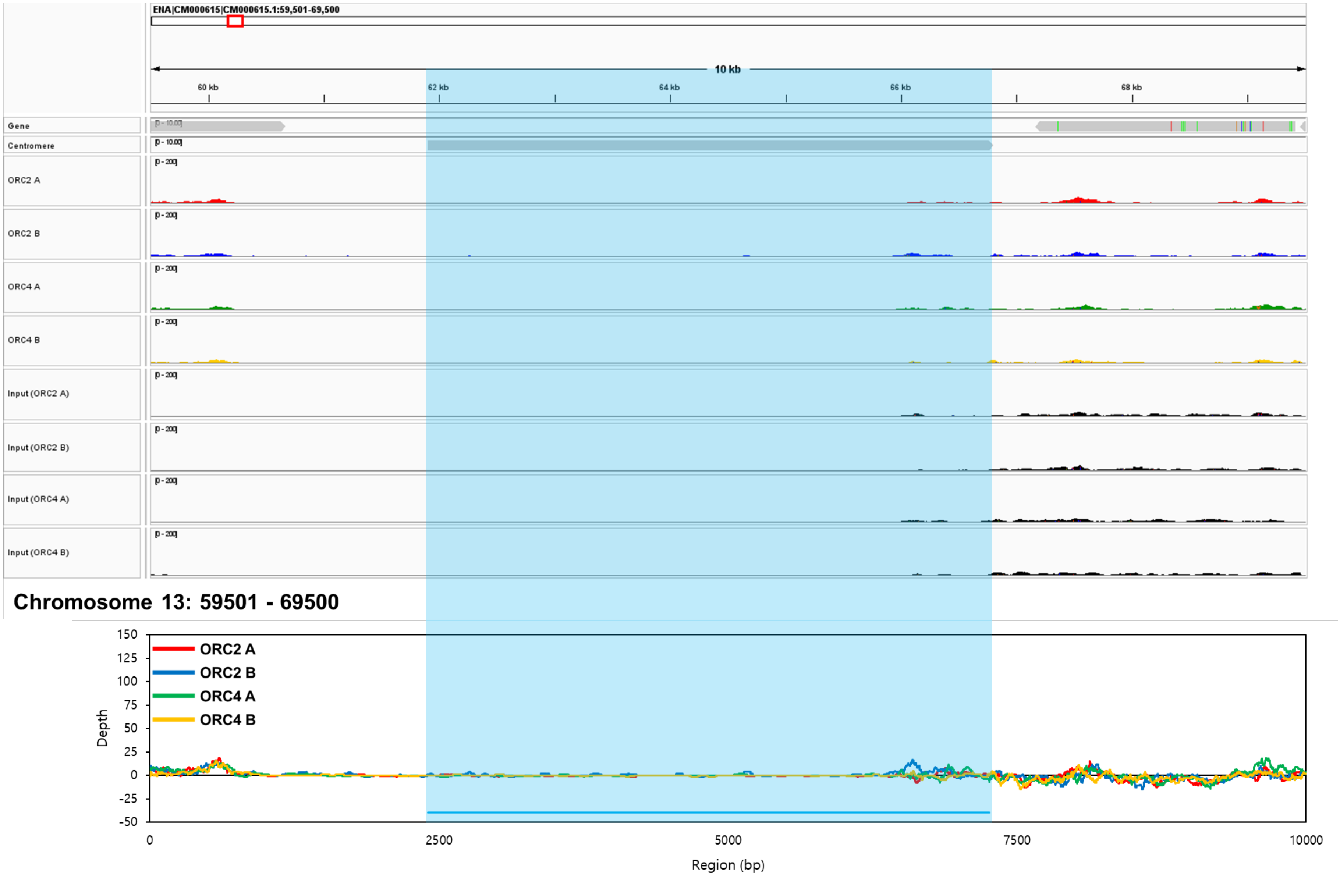
ENA|CM000615|CM000615.1:59,501-69,500
10 kb
60 kb
62 kb
64 kb
66 kb
68 kb
Gene
Centromere
ORC2 A
ORC2 B
ORC4 A
ORC4 B
Input (ORC2 A)
Input (ORC2 B)
Input (ORC4 A)
Input (ORC4 B)
Chromosome 13: 59501 - 69500
ORC2 A
ORC2 B
ORC4 A
ORC4 B
Depth
150
125
100
75
50
25
0
-25
-50
0
2500
5000
7500
10000
Region (bp)

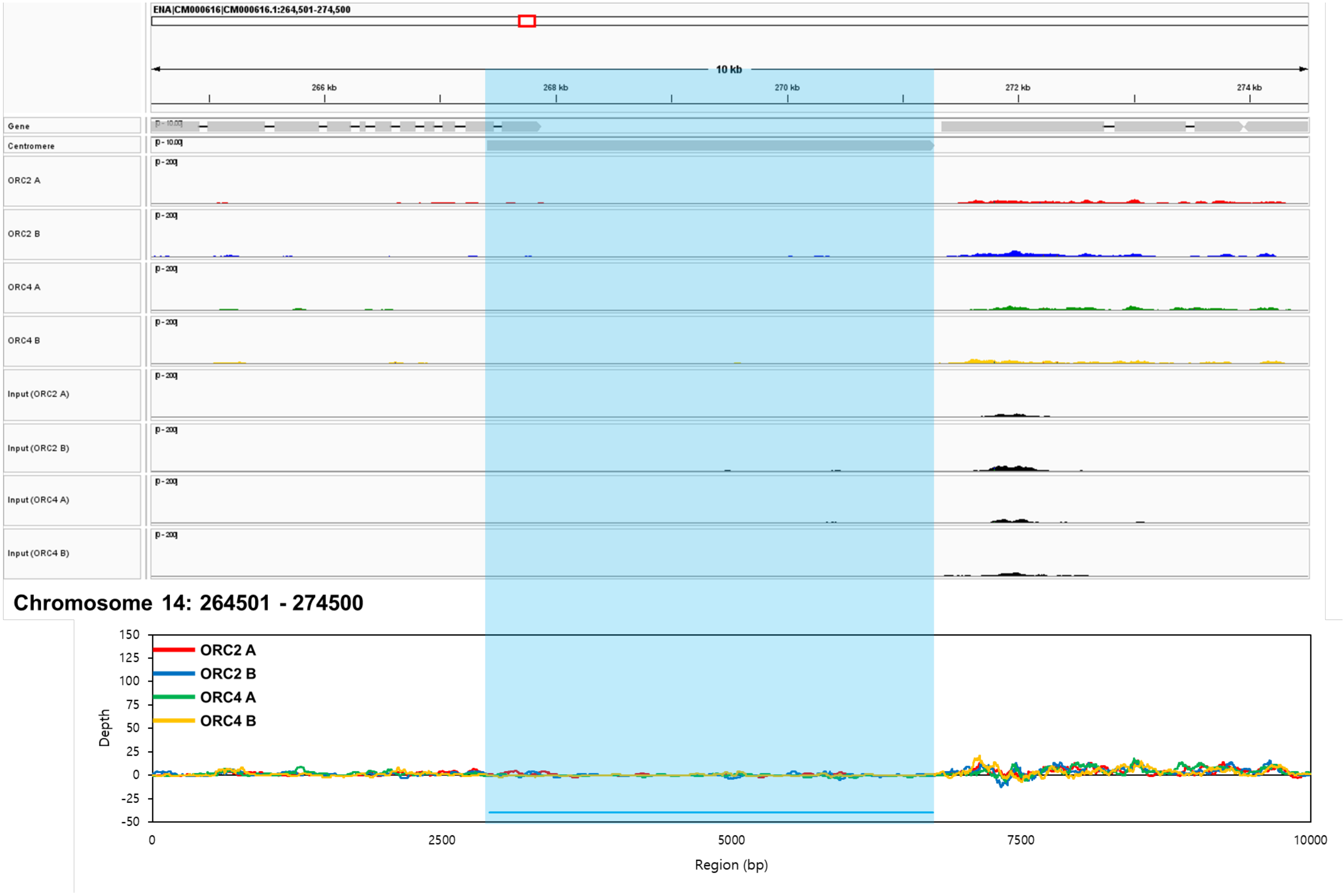
ENA|CM000616|CM000616.1:264,501-274,500
10 kb
266 kb
268 kb
270 kb
272 kb
274 kb
Gene
Centromere
ORC2 A
ORC2 B
ORC4 A
ORC4 B
Input (ORC2 A)
Input (ORC2 B)
Input (ORC4 A)
Input (ORC4 B)
Chromosome 14: 264501 - 274500
ORC2 A
ORC2 B
ORC4 A
ORC4 B
Depth
150
125
100
75
50
25
0
-25
-50
0
2500
5000
7500
10000
Region (bp)

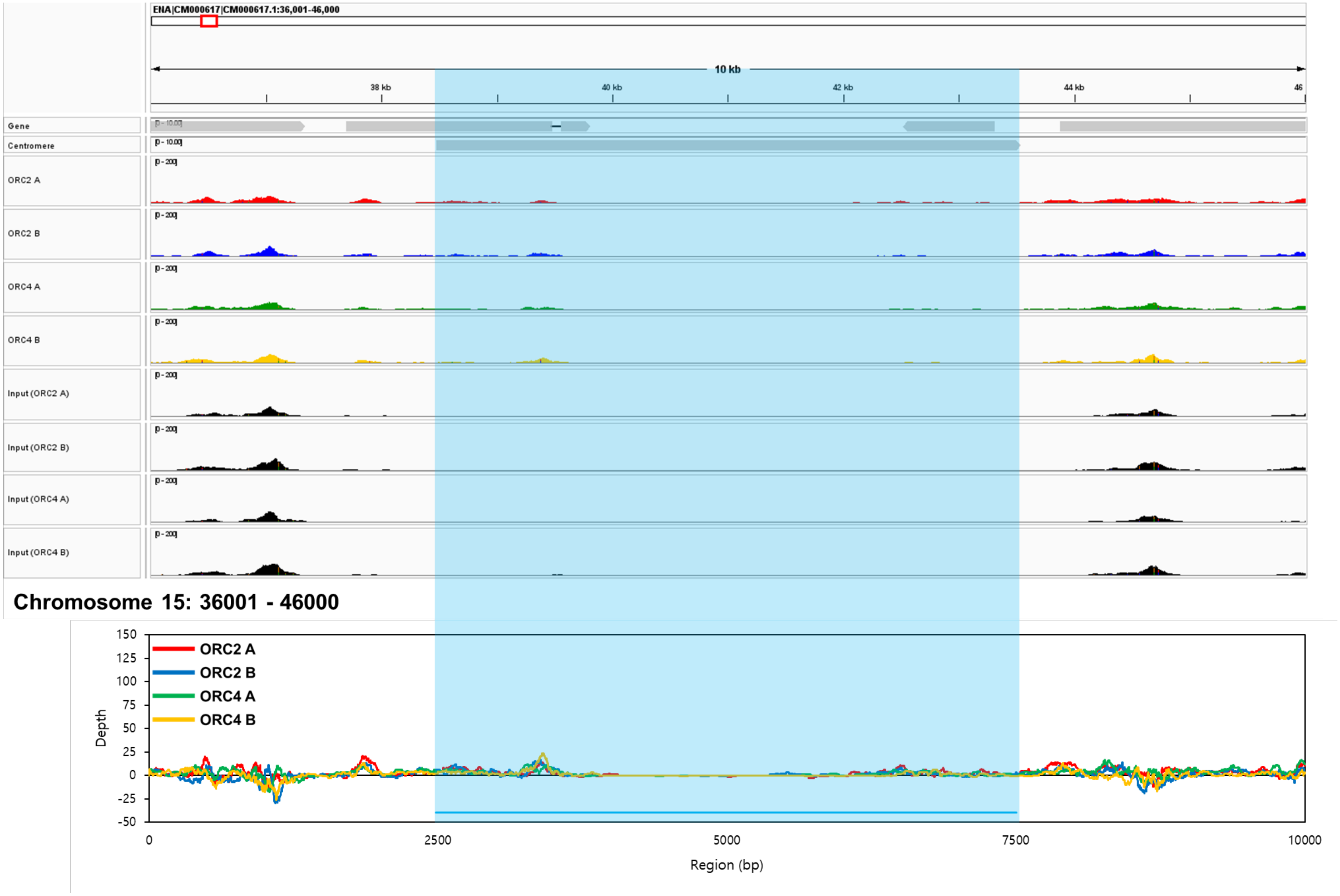
ENA|CM000617|CM000617.1:36,001-46,000
10 kb
38 kb
40 kb
42 kb
44 kb
46
Gene
Centromere
ORC2 A
ORC2 B
ORC4 A
ORC4 B
Input (ORC2 A)
Input (ORC2 B)
Input (ORC4 A)
Input (ORC4 B)
Chromosome 15: 36001 - 46000
ORC2 A
ORC2 B
ORC4 A
ORC4 B
Depth
150
125
100
75
50
25
0
-25
-50
0
2500
5000
7500
10000
Region (bp)

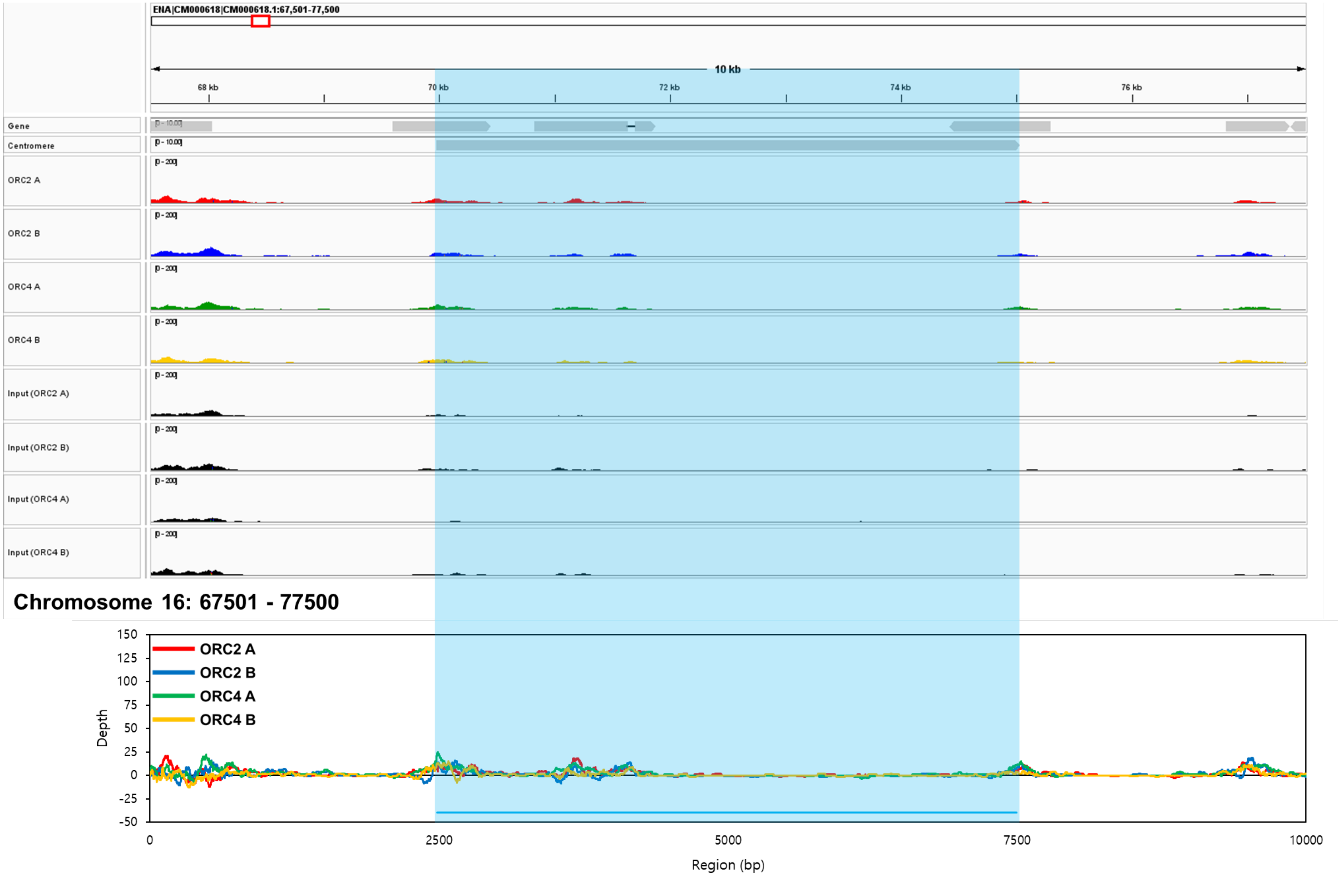
ENA|CM000618|CM000618.1:67,501-77,500
10 kb
68 kb
70 kb
72 kb
74 kb
76 kb
Gene
Centromere
ORC2 A
ORC2 B
ORC4 A
ORC4 B
Input (ORC2 A)
Input (ORC2 B)
Input (ORC4 A)
Input (ORC4 B)
Chromosome 16: 67501 - 77500
ORC2 A
ORC2 B
ORC4 A
ORC4 B
Depth
Region (bp)

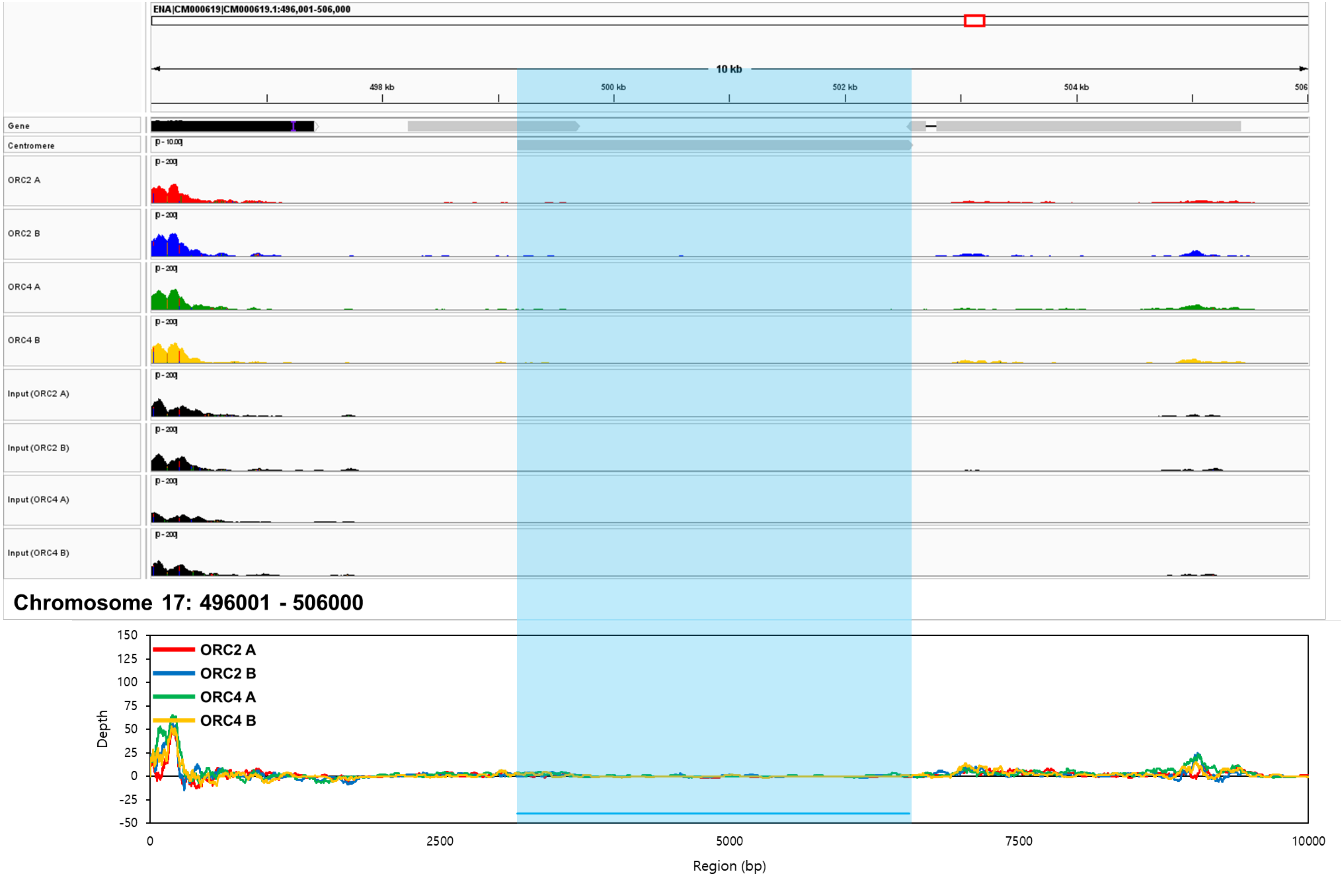
ENA|CM000619|CM000619.1:496,001-506,000
10 kb
498 kb
500 kb
502 kb
504 kb
506
Gene
Centromere
ORC2 A
ORC2 B
ORC4 A
ORC4 B
Input (ORC2 A)
Input (ORC2 B)
Input (ORC4 A)
Input (ORC4 B)
Chromosome 17: 496001 - 506000
Depth
Region (bp)
ORC2 A
ORC2 B
ORC4 A
ORC4 B
150
125
100
75
50
25
0
-25
-50
0
2500
5000
7500
10000

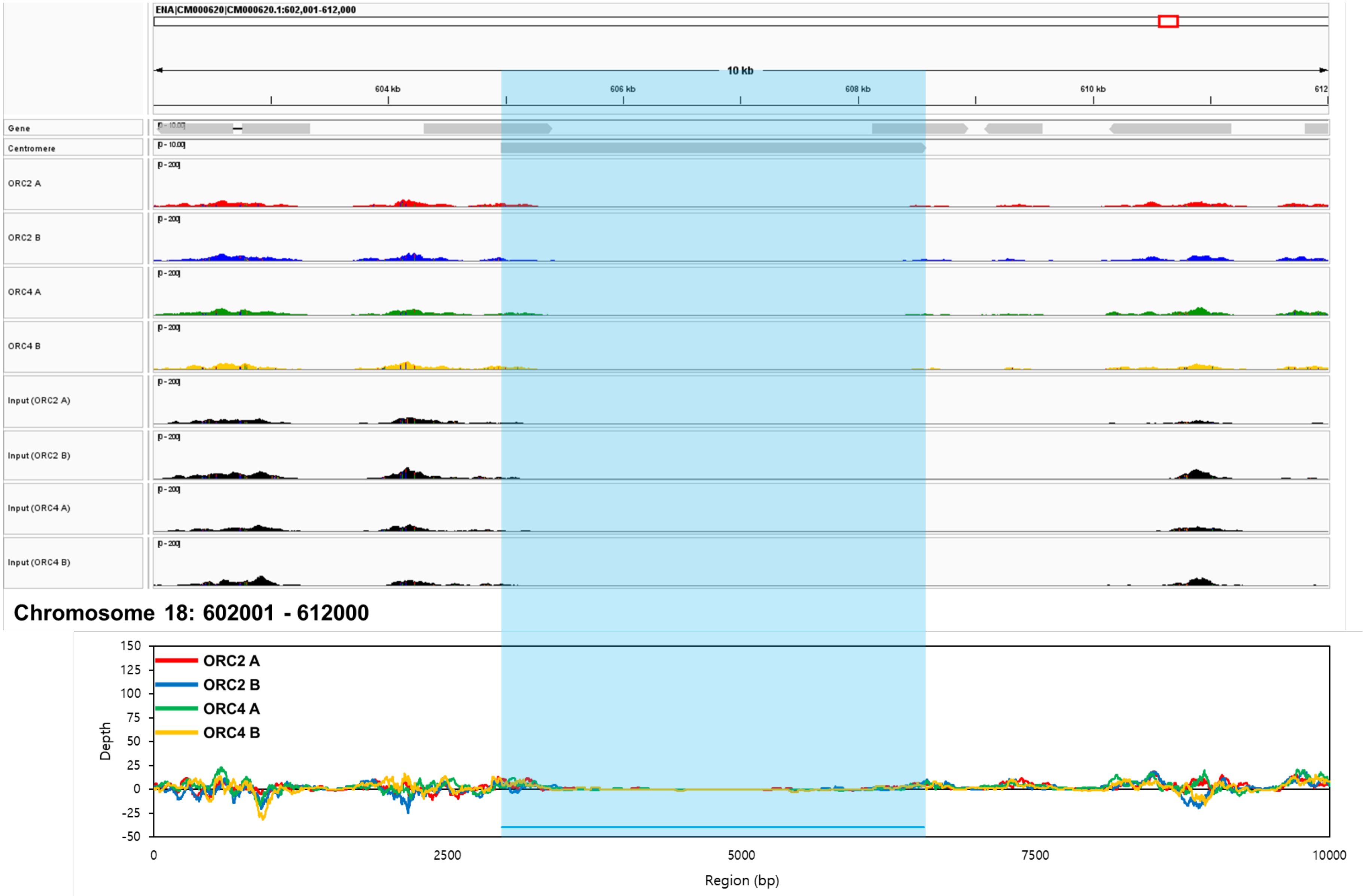
ENA|CM000620|CM000620.1:602,001-612,000
10 kb
604 kb
606 kb
608 kb
610 kb
612
Gene
Centromere
ORC2 A
ORC2 B
ORC4 A
ORC4 B
Input (ORC2 A)
Input (ORC2 B)
Input (ORC4 A)
Input (ORC4 B)
Chromosome 18: 602001 - 612000
ORC2 A
ORC2 B
ORC4 A
ORC4 B
Depth
Region (bp)

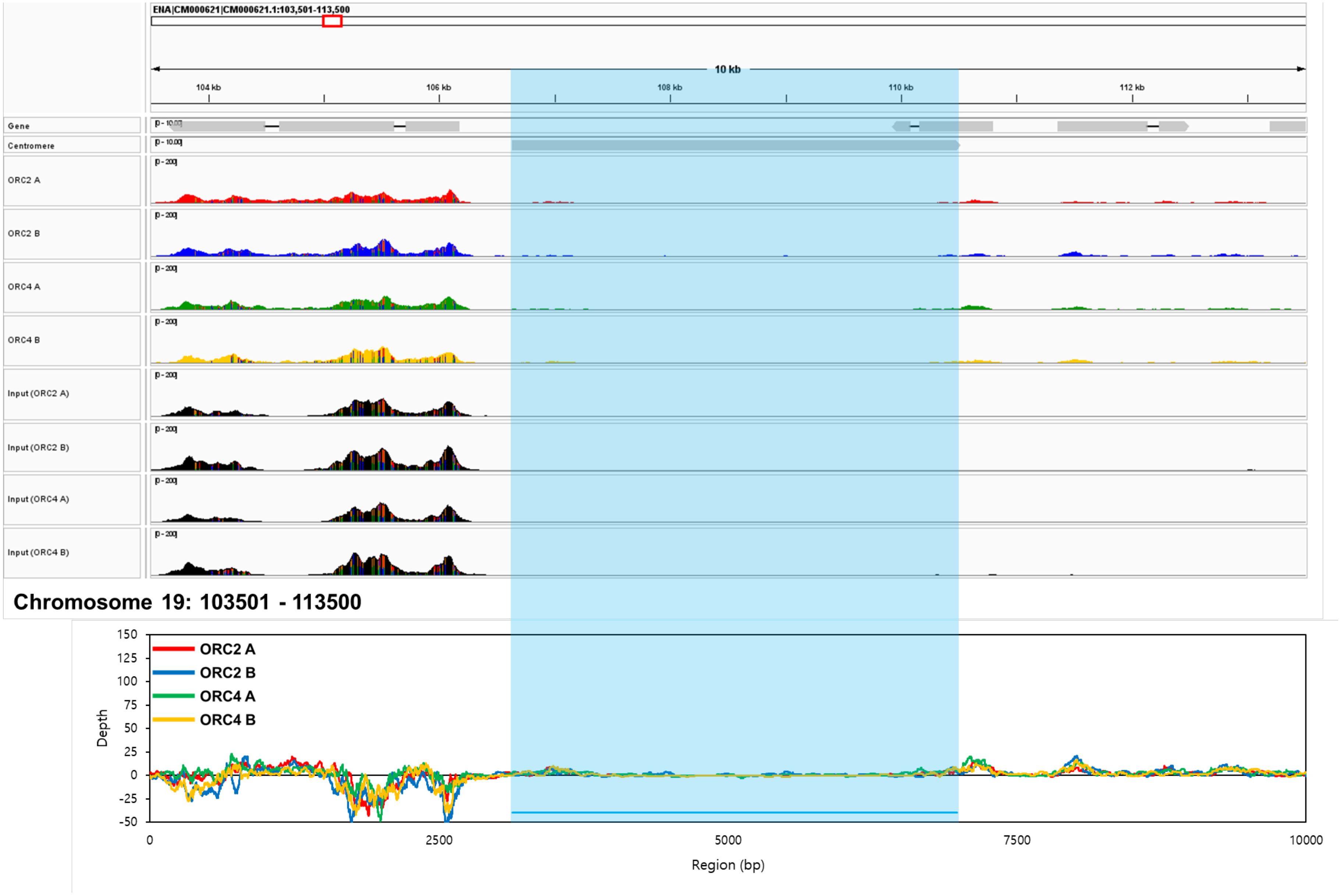
ENA|CM000621|CM000621.1:103,501-113,500
10 kb
104 kb
106 kb
108 kb
110 kb
112 kb
Gene
Centromere
ORC2 A
ORC2 B
ORC4 A
ORC4 B
Input (ORC2 A)
Input (ORC2 B)
Input (ORC4 A)
Input (ORC4 B)
Chromosome 19: 103501 - 113500
150
125
100
75
50
25
0
-25
-50
Depth
ORC2 A
ORC2 B
ORC4 A
ORC4 B
0
2500
5000
7500
10000
Region (bp)

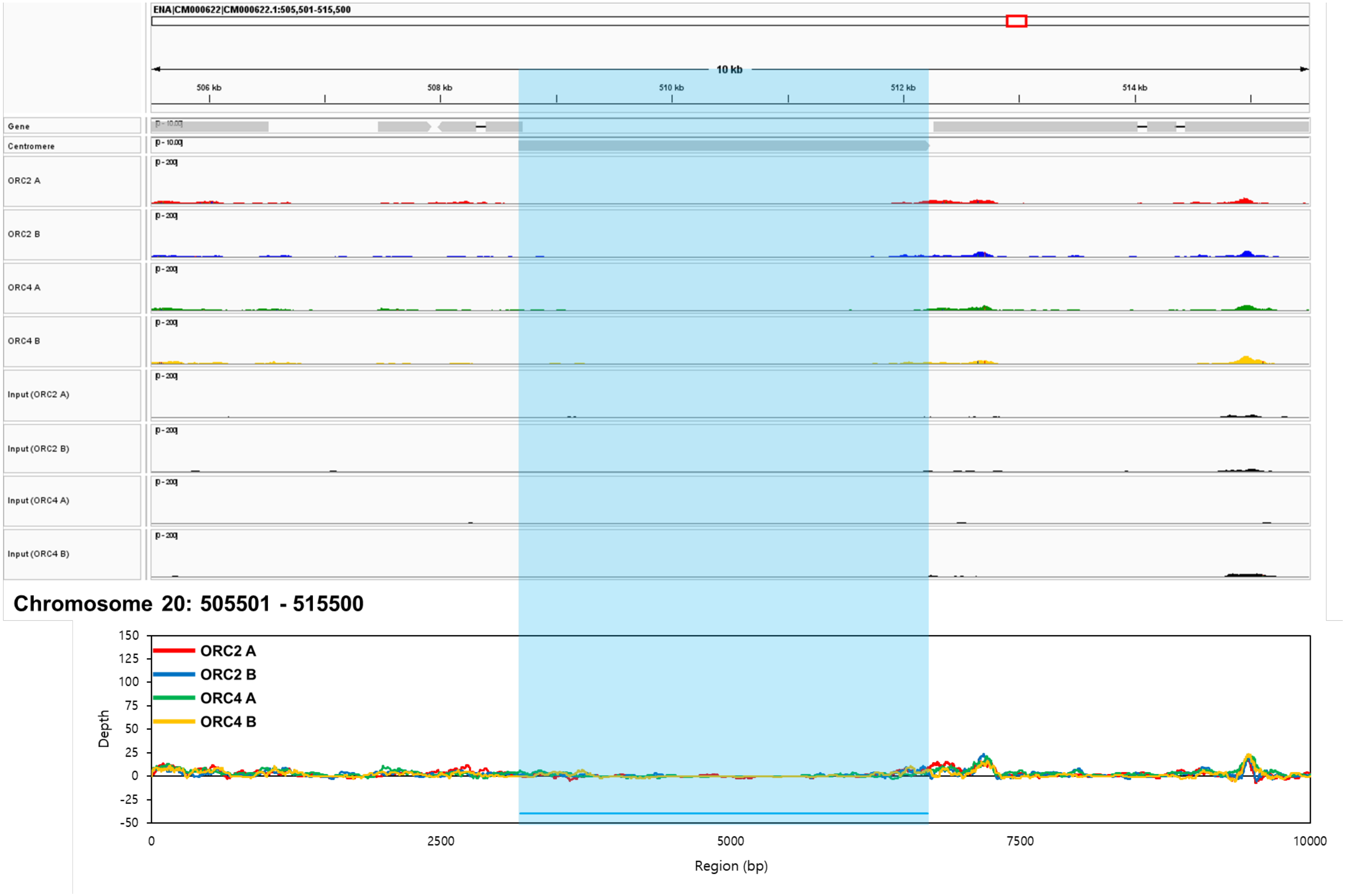
ENA|CM000622|CM000622.1:505,501-515,500
10 kb
506 kb
508 kb
510 kb
512 kb
514 kb
Gene
Centromere
ORC2 A
ORC2 B
ORC4 A
ORC4 B
Input (ORC2 A)
Input (ORC2 B)
Input (ORC4 A)
Input (ORC4 B)
Chromosome 20: 505501 - 515500
ORC2 A
ORC2 B
ORC4 A
ORC4 B
Depth
Region (bp)
150
125
100
75
50
25
0
-25
-50
0
2500
5000
7500
10000

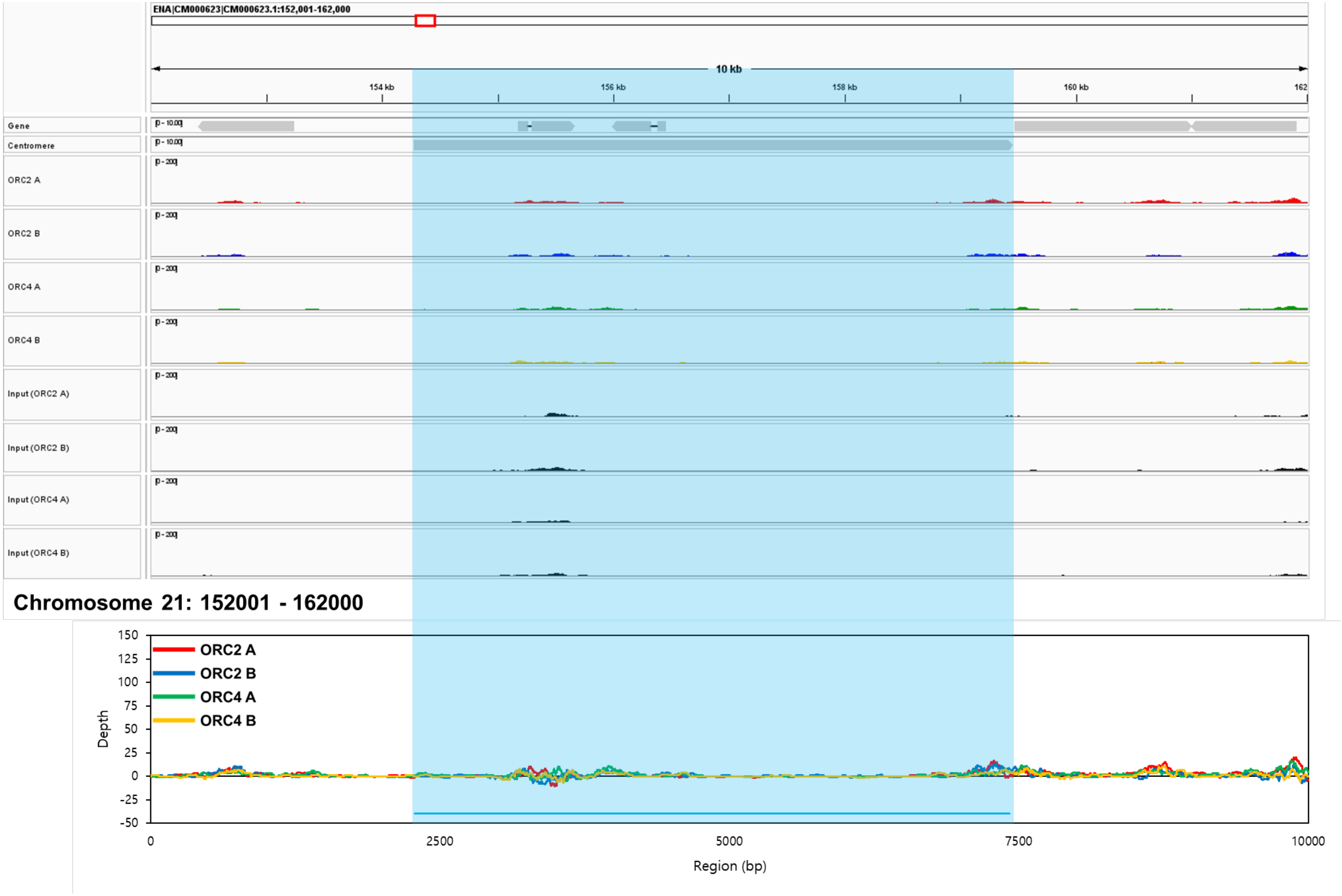
ENA|CM000623|CM000623.1:152,001-162,000
10 kb
154 kb
156 kb
158 kb
160 kb
162
Gene
Centromere
ORC2 A
ORC2 B
ORC4 A
ORC4 B
Input (ORC2 A)
Input (ORC2 B)
Input (ORC4 A)
Input (ORC4 B)
Chromosome 21: 152001 - 162000
ORC2 A
ORC2 B
ORC4 A
ORC4 B
Depth
Region (bp)
150
125
100
75
50
25
0
-25
-50
0
2500
5000
7500
10000

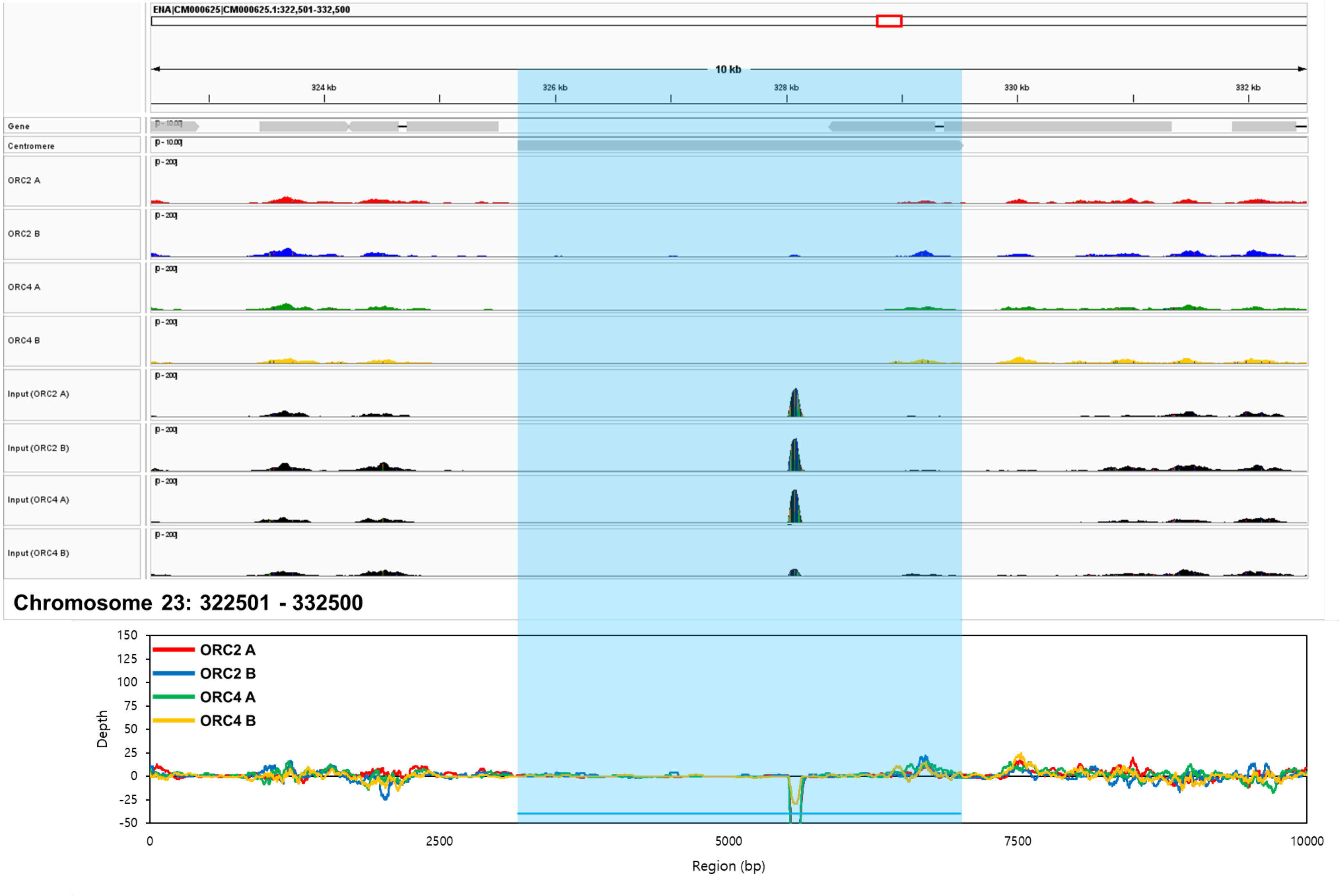
ENA|CM000625|CM000625.1:322,501-332,500
10 kb
324 kb
326 kb
328 kb
330 kb
332 kb
Gene
Centromere
ORC2 A
ORC2 B
ORC4 A
ORC4 B
Input (ORC2 A)
Input (ORC2 B)
Input (ORC4 A)
Input (ORC4 B)
Chromosome 23: 322501 - 332500
ORC2 A
ORC2 B
ORC4 A
ORC4 B
Depth
Region (bp)

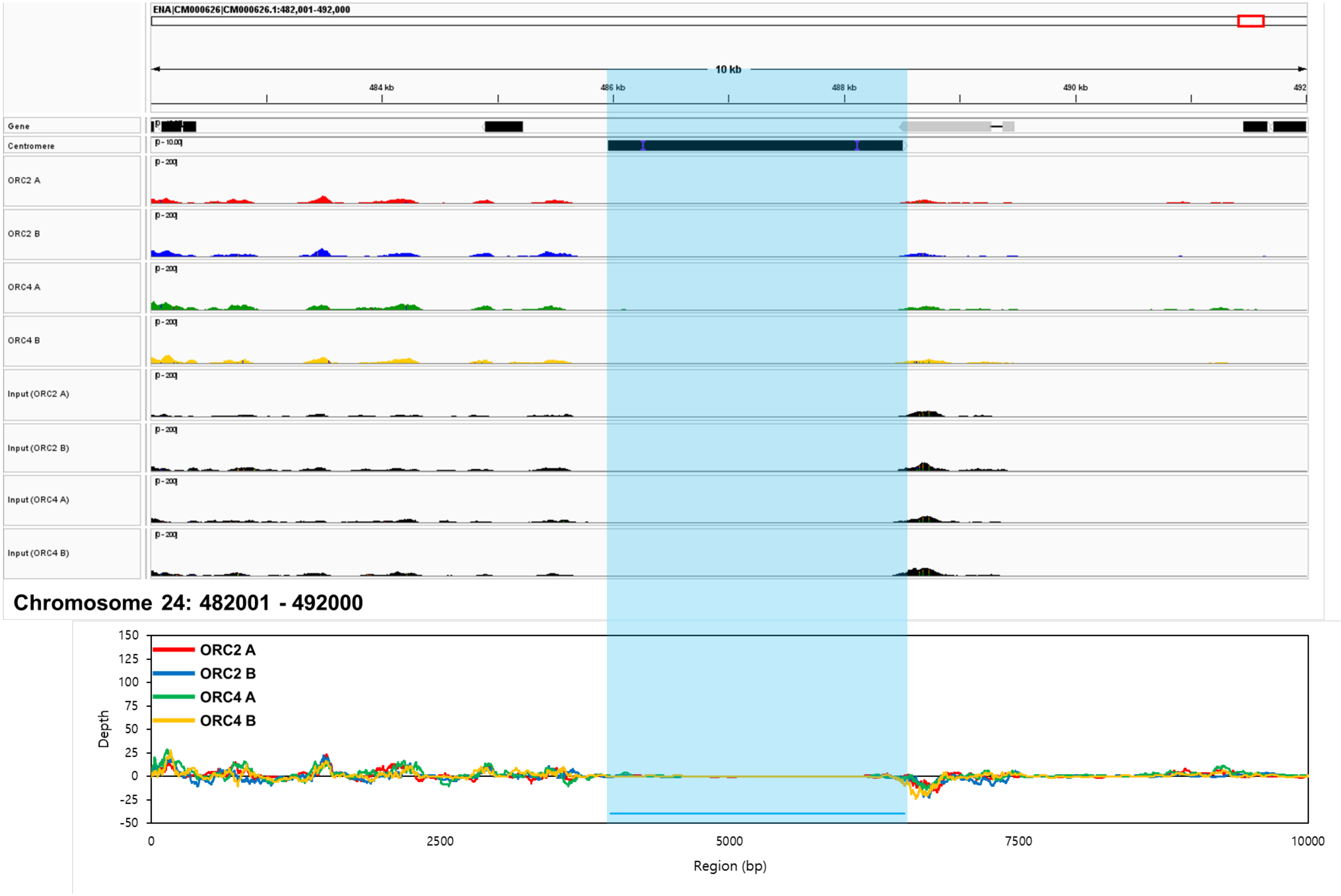
ENA|CM000626|CM000626.1:482,001-492,000
10 kb
484 kb
486 kb
488 kb
490 kb
492
Gene
Centromere
ORC2 A
ORC2 B
ORC4 A
ORC4 B
Input (ORC2 A)
Input (ORC2 B)
Input (ORC4 A)
Input (ORC4 B)
Chromosome 24: 482001 - 492000
Depth
Region (bp)
ORC2 A
ORC2 B
ORC4 A
ORC4 B
150
125
100
75
50
25
0
-25
-50
0
2500
5000
7500
10000

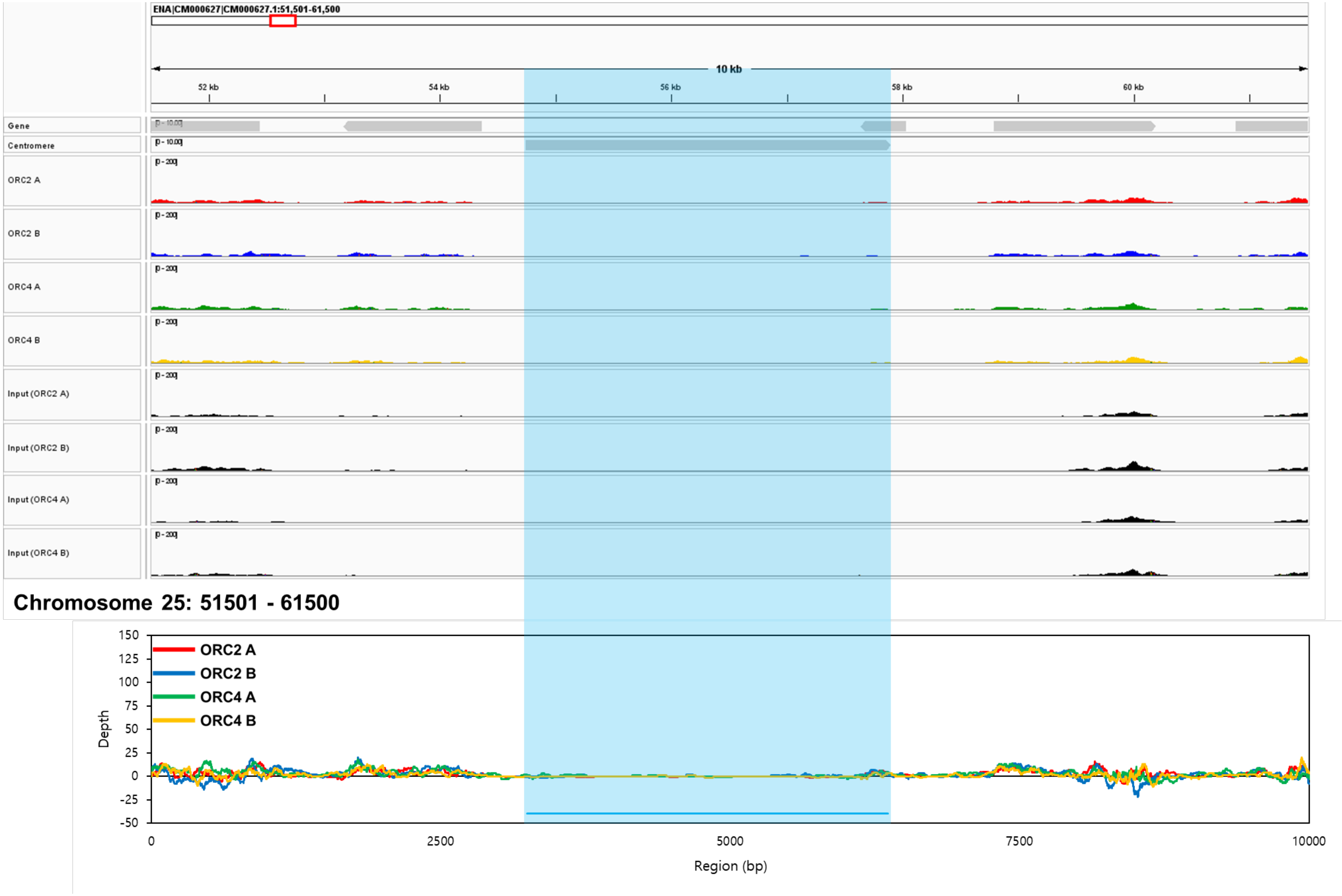
ENA|CM000627|CM000627.1:51,501-61,500
10 kb
52 kb
54 kb
56 kb
58 kb
60 kb
Gene
Centromere
ORC2 A
ORC2 B
ORC4 A
ORC4 B
Input (ORC2 A)
Input (ORC2 B)
Input (ORC4 A)
Input (ORC4 B)
Chromosome 25: 51501 - 61500
ORC2 A
ORC2 B
ORC4 A
ORC4 B
Depth
Region (bp)

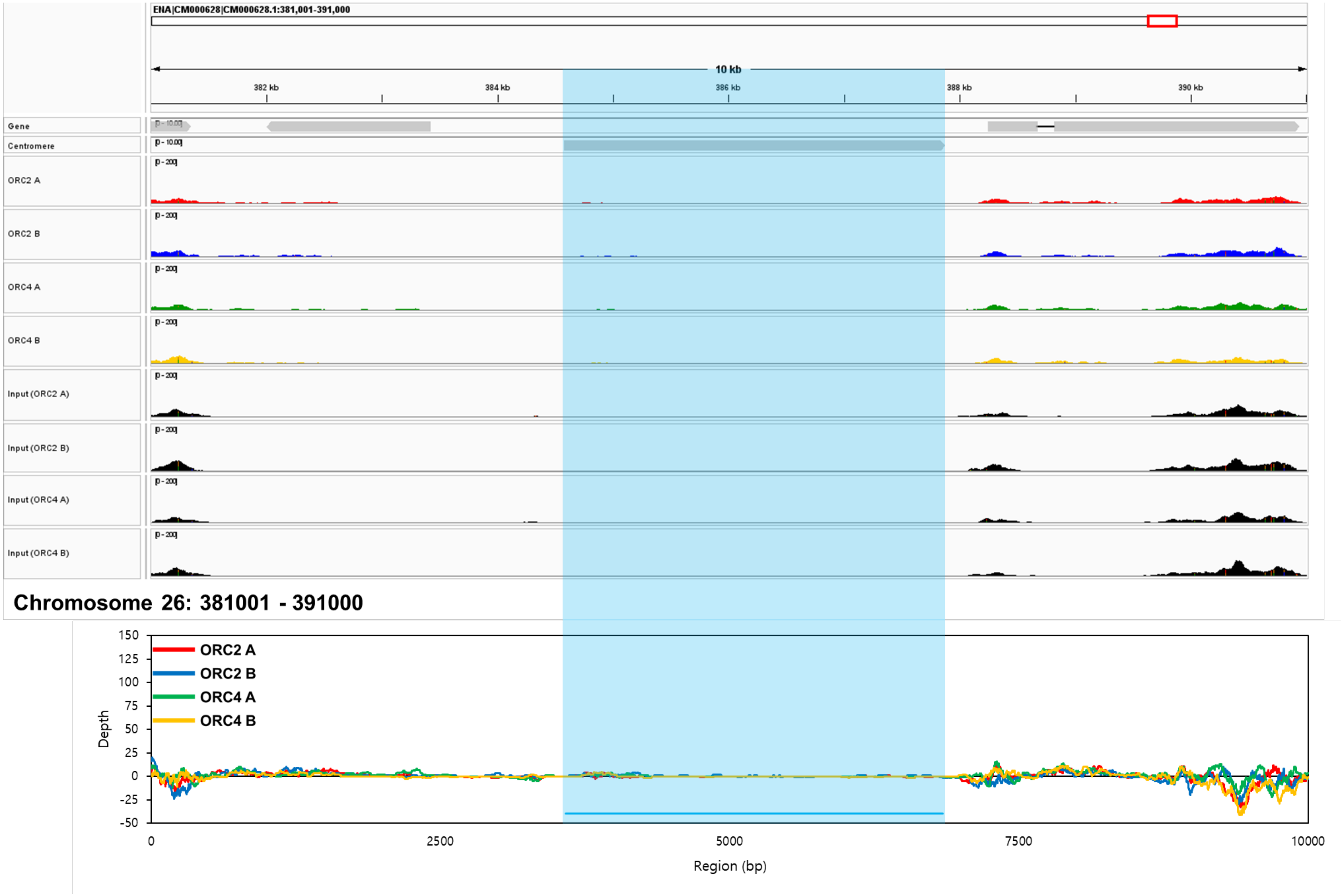
ENA|CM000628|CM000628.1:381,001-391,000
10 kb
382 kb
384 kb
386 kb
388 kb
390 kb
Gene
Centromere
ORC2 A
ORC2 B
ORC4 A
ORC4 B
Input (ORC2 A)
Input (ORC2 B)
Input (ORC4 A)
Input (ORC4 B)
Chromosome 26: 381001 - 391000
Depth
Region (bp)
ORC2 A
ORC2 B
ORC4 A
ORC4 B
150
125
100
75
50
25
0
-25
-50
0
2500
5000
7500
10000

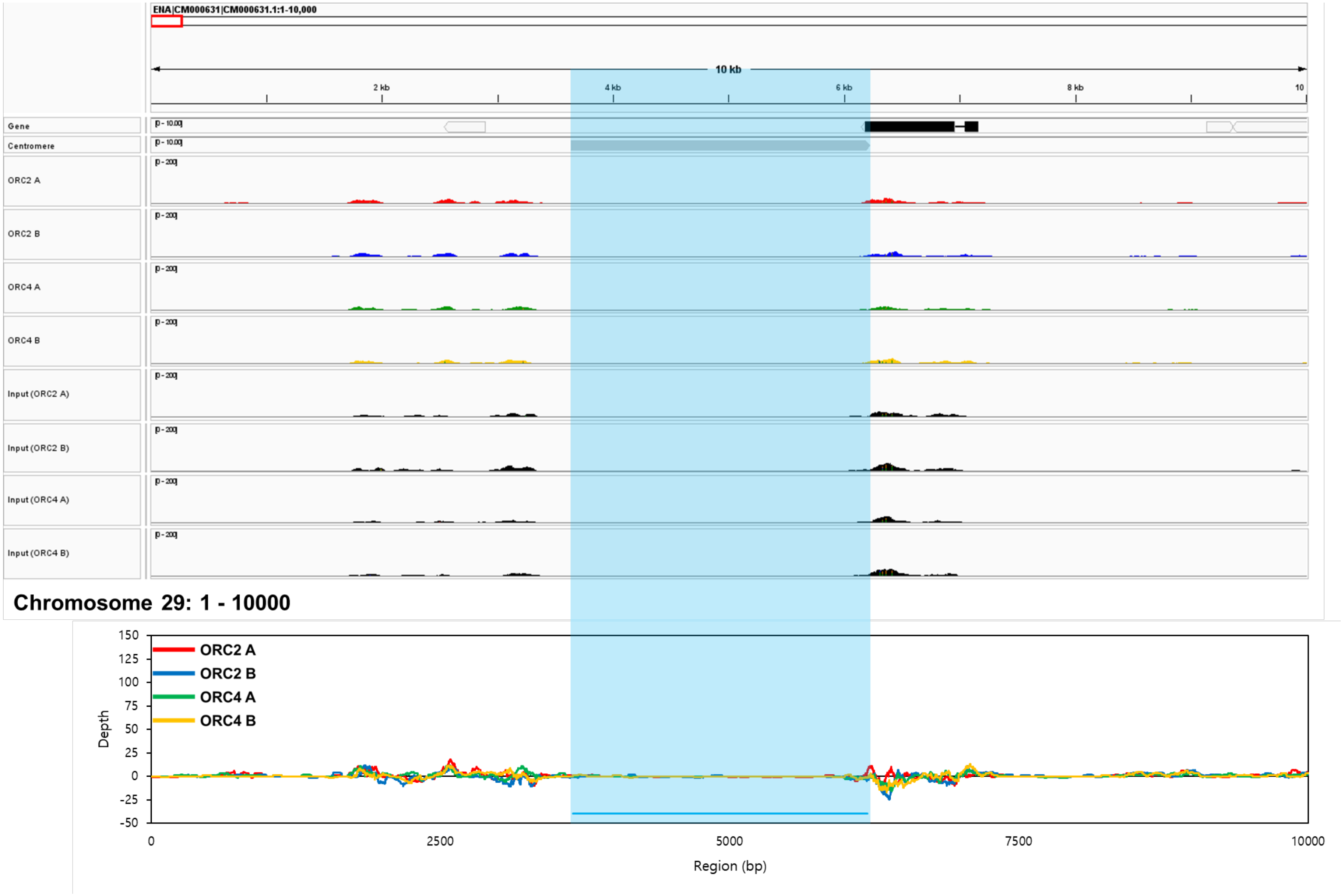
ENA|CM000631|CM000631.1:1-10,000
10 kb
2 kb
4 kb
6 kb
8 kb
10
Gene
Centromere
ORC2 A
ORC2 B
ORC4 A
ORC4 B
Input (ORC2 A)
Input (ORC2 B)
Input (ORC4 A)
Input (ORC4 B)
Chromosome 29: 1 - 10000
ORC2 A
ORC2 B
ORC4 A
ORC4 B
Depth
Region (bp)
150
125
100
75
50
25
0
-25
-50
0
2500
5000
7500
10000

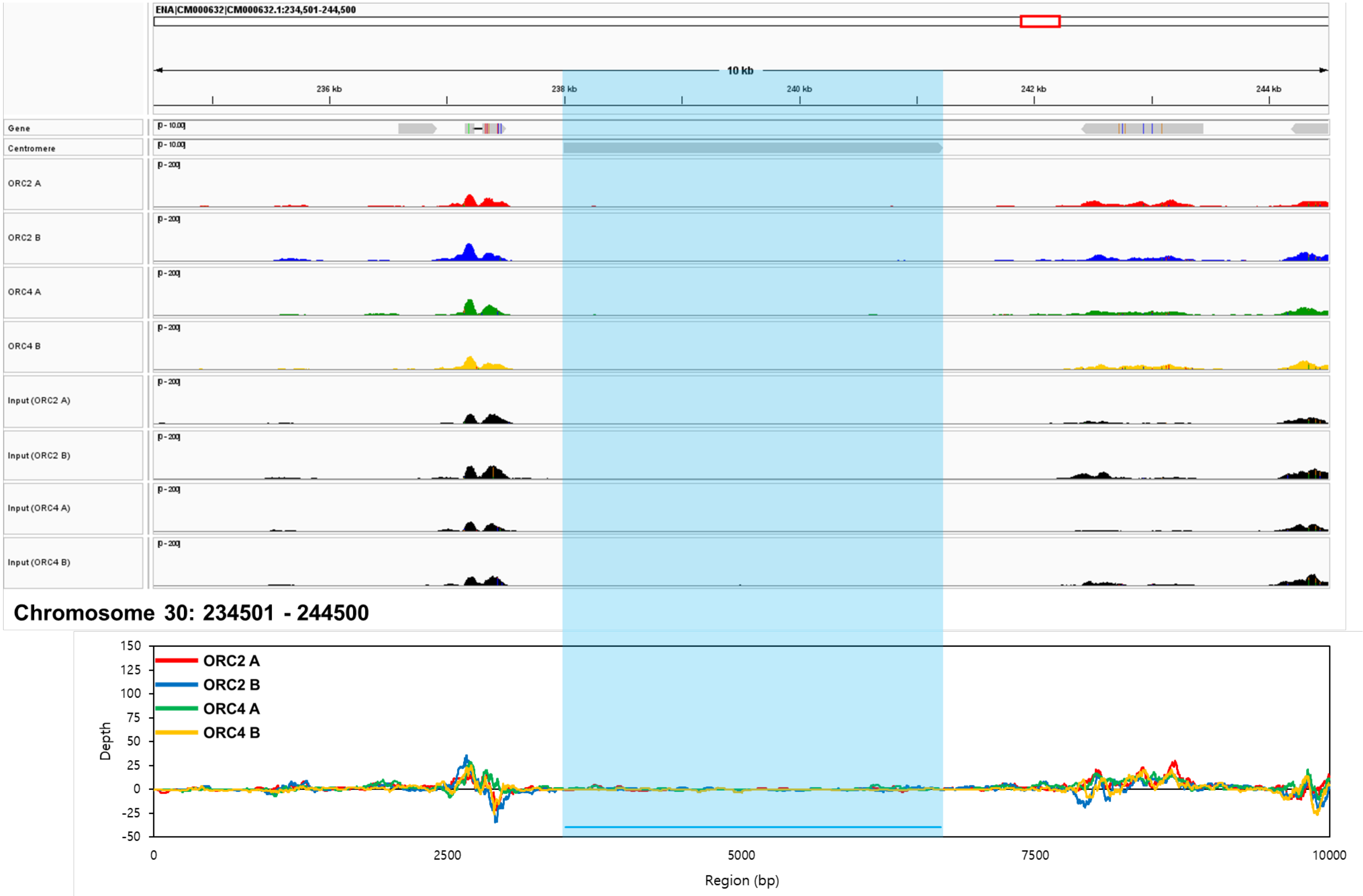


**Fig. S13: Peak data and depth values at the centromere (chromosomes 2–10, 13–21, 23–26, and 29–30) region and surroundings of *Phaeodactylum tricornutum*.**

The results obtained from ORC2 A (red peaks and lines), ORC2 B (blue peaks and lines), ORC4 A (green peaks and lines), and ORC4 B (orange peaks and lines) were visualised through peak (IGV image) and depth images, and the centromere regions (Diner *et al.,* 2017) indicated by light blue lines over the peak and depth images.

**Table S1.** Sequences of primers used in this study.

| Purpose | Target | Primer name | Sequence (5'-3') |
|---|---|---|---|
| Amplification of vector | pPha-Plasmid | pPha_Fw | AAGCTTCAGAAGCGTGCTATCG |
| | | pPha_Rev | GAATTCCGTTCGCACAAGTG |
| Amplification of *Phaeodactylum tricornutum* ORC subunit gene fused with FLAG-tag for insert fragment | ORC2 | ORC2_Fw | GTGCGAACGGAATTCATGGATTATAAAGATGATGATGATAAAATGGAGACAGAGATT |
| | | ORC2_Rev | ACGCTTCTGAAGCTTTCAATGTTTACGTGG |
| | ORC4 | ORC4_Fw | GTGCGAACGGAATTCATGGATTATAAAGATGATGATGATAAAATGACTGTTGTTCCG |
| | | ORC4_Rev | ACGCTTCTGAAGCTTCTAGTTCATCTTTCG |

The sequences that compose the FLAG-tag were displayed using underlining.

| | | | |
|---|---|---|---|
| Sequencing of inserts introduced into vectors | ORC2 and ORC4 | PtNRpseq_Fw | GGTGTTCACAATTTTGCCTCCTC |
| | ORC2 | ORC2_seq_Fw01 | CCTAAAGCGTAGAGCTTCAA |
| | | ORC2_seq_Fw02 | AATTCGCGCGGCGGGAGCTG |
| | | ORC2_seq_Fw03 | CGTCCGTATGTAGAGGAACT |
| | ORC4 | ORC4_seq_Fw01 | CAGGCTTTGCGAGATGAGCT |
| | | ORC4_seq_Fw02 | TGTGTGCTTTGTCGGTATGA |
| | | ORC4_seq_Fw03 | TAGAGGATGTGGCTGTCGAT |

**Table S2.** BLAST output of each SNP against the genome database of *Phaeodactylum tricornutum*.

| Gene | SNP type | DNA | | | | Protein | | | |
|---|---|---|---|---|---|---|---|---|---|
| | | Length (bp) | Overlap (%) | Number of base substitution (bp) | Sequence similarity (%) | Length (aa) | Overlap (%) | Number of amino acid replacement (aa) | Sequence similarity (%) |
| ORC2 | Type A | 1767 | 100 | 6 | 99 | 588 | 100 | 2 | 99 |
| | Type B | 1767 | 100 | 2 | 99 | 588 | 100 | 2 | 99 |
| ORC4 | Type A | 1803 | 100 | 3 | 99 | 600 | 100 | 0 | 100 |
| | Type B | 1803 | 100 | 6 | 99 | 600 | 100 | 4 | 99 |

**Table S3.** Distribution of peak called sites by the ChIP-Seq experiments on the *Phaeodactylum tricornutum* genome.

| Chromosome | ORC2A | ORC2B | ORC4A | ORC4B | ORC2A/ORC2B[a] | ORC2A/ORC4A[b] | ORC2A/ORC4B[c] | ORC2B/ORC4A[d] | ORC2B/ORC4B[e] | ORC4A/ORC4B[f] | ORC2A/ORC2B/ORC4A[g] | ORC2A/ORC2B/ORC4B[h] | ORC2A/ORC4A/ORC4B[i] | ORC2B/ORC4A/ORC4B[j] | ORC2A/ORC2B/ORC4A/ORC4B[k] | Total |
|---|---|---|---|---|---|---|---|---|---|---|---|---|---|---|---|---|
| 1 | 2 | 1 | 2 | 4 | 0 | 2 | 1 | 0 | 0 | 0 | 0 | 0 | 0 | 1 | 9 | 22 |
| 2 | 0 | 5 | 0 | 1 | 0 | 0 | 0 | 0 | 0 | 0 | 1 | 0 | 1 | 0 | 0 | 8 |
| 3 | 3 | 5 | 2 | 5 | 2 | 0 | 0 | 0 | 0 | 0 | 0 | 0 | 1 | 0 | 5 | 23 |
| 4 | 3 | 5 | 1 | 1 | 0 | 0 | 0 | 0 | 0 | 0 | 0 | 0 | 0 | 0 | 1 | 11 |
| 5 | 1 | 4 | 1 | 3 | 0 | 0 | 1 | 0 | 0 | 0 | 1 | 1 | 0 | 0 | 3 | 15 |
| 6 | 0 | 3 | 0 | 0 | 0 | 0 | 0 | 0 | 1 | 0 | 0 | 0 | 0 | 0 | 0 | 4 |
| 7 | 1 | 1 | 1 | 1 | 0 | 0 | 0 | 2 | 0 | 0 | 0 | 0 | 0 | 0 | 3 | 9 |
| 8 | 1 | 1 | 0 | 3 | 0 | 0 | 0 | 0 | 0 | 0 | 0 | 0 | 0 | 0 | 0 | 5 |
| 9 | 2 | 1 | 1 | 0 | 0 | 0 | 0 | 0 | 0 | 0 | 0 | 0 | 0 | 0 | 0 | 4 |
| 10 | 0 | 6 | 3 | 4 | 0 | 0 | 0 | 0 | 1 | 0 | 0 | 0 | 0 | 0 | 2 | 16 |
| 11 | 0 | 6 | 1 | 2 | 0 | 0 | 0 | 0 | 0 | 0 | 0 | 1 | 0 | 0 | 0 | 10 |
| 12 | 5 | 7 | 0 | 1 | 0 | 1 | 0 | 0 | 0 | 0 | 0 | 0 | 1 | 1 | 2 | 18 |
| 13 | 2 | 3 | 1 | 3 | 0 | 0 | 1 | 0 | 0 | 0 | 0 | 0 | 0 | 0 | 0 | 10 |
| 14 | 3 | 3 | 2 | 1 | 0 | 0 | 0 | 0 | 0 | 0 | 2 | 0 | 0 | 0 | 2 | 13 |
| 15 | 3 | 2 | 0 | 1 | 0 | 0 | 0 | 0 | 0 | 0 | 0 | 0 | 0 | 0 | 3 | 9 |
| 16 | 1 | 4 | 1 | 1 | 1 | 0 | 0 | 0 | 0 | 2 | 0 | 0 | 1 | 0 | 7 | 18 |

| | | | | | | | | | | | | | | | | |
|---|---|---|---|---|---|---|---|---|---|---|---|---|---|---|---|---|
| 17 | 0 | 1 | 0 | 2 | 0 | 0 | 1 | 0 | 0 | 0 | 0 | 0 | 0 | 1 | 4 | 9 |
| 18 | 4 | 4 | 0 | 2 | 1 | 0 | 0 | 1 | 0 | 0 | 0 | 2 | 0 | 1 | 4 | 19 |
| 19 | 3 | 5 | 1 | 2 | 2 | 0 | 0 | 1 | 0 | 1 | 0 | 3 | 0 | 1 | 4 | 23 |
| 20 | 0 | 1 | 0 | 1 | 0 | 0 | 0 | 0 | 0 | 2 | 0 | 1 | 0 | 1 | 1 | 7 |
| 21 | 0 | 1 | 1 | 2 | 0 | 0 | 0 | 0 | 0 | 0 | 0 | 0 | 0 | 0 | 0 | 4 |
| 22 | 2 | 2 | 1 | 3 | 0 | 0 | 1 | 0 | 1 | 0 | 0 | 0 | 1 | 0 | 1 | 12 |
| 23 | 0 | 3 | 1 | 0 | 0 | 1 | 0 | 0 | 0 | 0 | 0 | 0 | 0 | 0 | 1 | 6 |
| 24 | 1 | 1 | 1 | 1 | 1 | 0 | 1 | 1 | 0 | 1 | 1 | 2 | 0 | 0 | 4 | 15 |
| 25 | 1 | 6 | 6 | 1 | 0 | 0 | 0 | 0 | 0 | 0 | 0 | 0 | 0 | 1 | 4 | 19 |
| 26 | 2 | 1 | 1 | 0 | 1 | 0 | 1 | 0 | 0 | 0 | 1 | 0 | 0 | 0 | 2 | 9 |
| 27 | 1 | 1 | 1 | 1 | 0 | 0 | 0 | 1 | 1 | 0 | 0 | 0 | 0 | 0 | 0 | 6 |
| 28 | 0 | 0 | 0 | 0 | 0 | 0 | 0 | 0 | 0 | 0 | 0 | 0 | 0 | 0 | 0 | 0 |
| 29 | 0 | 1 | 1 | 0 | 2 | 0 | 0 | 0 | 1 | 0 | 0 | 0 | 0 | 0 | 4 | 9 |
| 30 | 0 | 2 | 2 | 1 | 2 | 0 | 0 | 0 | 0 | 1 | 1 | 0 | 0 | 0 | 0 | 9 |
| 31 | 2 | 0 | 0 | 1 | 1 | 1 | 1 | 0 | 0 | 0 | 1 | 2 | 0 | 0 | 2 | 11 |
| 32 | 0 | 0 | 0 | 0 | 0 | 0 | 0 | 0 | 0 | 0 | 0 | 0 | 0 | 0 | 0 | 0 |
| 33 | 0 | 0 | 0 | 0 | 0 | 0 | 0 | 0 | 0 | 1 | 0 | 0 | 0 | 0 | 1 | 2 |
| Total | 43 | 86 | 32 | 48 | 13 | 5 | 8 | 6 | 5 | 8 | 8 | 12 | 5 | 7 | 69 | 355 |

[a]: Sites shared only between the results of ORC2A and ORC2B
[b]: Sites shared only between the results of ORC2A and ORC4A
[c]: Sites shared only between the results of ORC2A and ORC4B
[d]: Sites shared only between the results of ORC2B and ORC4A
[e]: Sites shared only between the results of ORC2B and ORC4B

[f]: Sites shared only between the results of ORC4A and ORC4B
[g]: Sites shared only between the results of ORC2A, ORC2B, and ORC4A
[h]: Sites shared only between the results of ORC2A, ORC2B, and ORC4B
[i]: Sites shared only between the results of ORC2A, ORC4A, and ORC4B
[j]: Sites shared only between the results of ORC2B, ORC4A, and ORC4B
[k]: Sites shared only between the results of ORC2A, ORC2B, ORC4A, and ORC4B
